# Evolutionary history of dimethylsulfoniopropionate (DMSP) demethylation enzyme DmdA in marine bacteria

**DOI:** 10.1101/766360

**Authors:** Laura Hernández, Alberto Vicens, Luis Enrique Eguiarte, Valeria Souza, Valerie De Anda, José M. González

## Abstract

Dimethylsulfoniopropionate (DMSP), an osmolyte produced by oceanic phytoplankton, is predominantly degraded by bacteria belonging to the *Roseobacter* lineage and other marine *Alphaproteobacteria* via DMSP-dependent demethylase A protein (DmdA). To date, the evolutionary history of DmdA gene family is unclear. Some studies indicate a common ancestry between DmdA and GcvT gene families and a co-evolution between *Roseobacter* and the DMSP-producing-phytoplankton around 250 million years ago (Mya). In this work, we analyzed the evolution of DmdA under three possible evolutionary scenarios: 1) a recent common ancestor of DmdA and GcvT, 2) a coevolution between *Roseobacter* and the DMSP-producing-phytoplankton, and 3) pre-adapted enzymes to DMSP prior to *Roseobacter* origin. Our analyses indicate that DmdA is a new gene family originated from GcvT genes by duplication and functional divergence driven by positive selection before a coevolution between *Roseobacter* and phytoplankton. Our data suggest that *Roseobacter* acquired *dmdA* by horizontal gene transfer prior to exposition to an environment with higher DMSP. Here, we propose that the ancestor that carried the DMSP demethylation pathway genes evolved in the Archean, and was exposed to a higher concentration of DMSP in a sulfur rich atmosphere and anoxic ocean, compared to recent *Roseobacter* ecoparalogs (copies performing the same function under different conditions), which should be adapted to lower concentrations of DMSP.

## INTRODUCTION

Dimethylsulfoniopropionate (DMSP) is an osmolyte synthesized by oceanic phytoplankton (Galinski, 1995; Yoch, 2002). This molecule became abundant in the oceans 250 million years ago (Mya), coinciding with the expansion and diversification of dinoflagellates (Bullock et al., 2017). Since then, it has played an important role in the biogeochemistry of sulfur cycle on Earth (Lovelock, 1983). DMSP is the main precursor of the climate-relevant gas dimethylsulfide (DMS; Reisch et al., 2011). In marine ecosystems, DMSP is rapidly degraded by different bacterial communities (González et al., 1999), and some strains seem to be very efficient and even become dependent on its degradation (Tripp et al., 2008). In fact, DMSP supports up to 13% of the bacterial carbon demand in surface waters, making it one of the most significant substrates for bacterioplankton (Kiene et al., 1999; González et al., 1999). *Candidatus* Pelagibacter ubique (SAR11), dominant in the bacterioplankton and especially in surface waters, can only use sulfur atoms derived organic molecules, such as DMSP (Tripp et al., 2008). In the case of *Ruegeria pomeroyi* DSS-3, a model organism for DMSP studies, the turnover rate of DMSP transformation depends on salinity conditions (Salgado et al., 2014).

The first step in the degradation of DMSP involves two competing pathways, cleavage and demethylation. The DMSP cleavage pathway metabolizes DMSP with the release of DMS (Kiene et al., 1999), a step catalyzed by a number of enzymes (Curson et al., 2011). In the alternative pathway, DMSP is first demethylated by a DMSP-dependent demethylase A protein (DmdA; Howard et al., 2006). Compared to the DMS-releasing pathway, *dmdA* is the most frequent gene in the genomes of oceanic bacteria (Newton et al., 2010). The DmdA enzyme was originally annotated as a glycine cleavage T-protein (GcvT) in the model bacteria *R. pomeroyi* (Reisch et al., 2011a), although it forms a separate clade from the known GcvTs (*gcvT, gcvH, gcvP* and *gcvT-C*) (Bullock et al., 2017). Despite their structural similarity which might indicate a common ancestry, DmdA and GcvT are mechanistically distinct (Schuller et al., 2012). DmdA produces 5-methyl-THF from DMSP as the result of a redox-neutral methyl transfer while GcvT converts glycine to 5,10-methylene-THF (Reisch et al., 2008).

Nearly all known DMSP-catabolizing bacteria belong to the phylum *Proteobacteria* with DmdA orthologs found in most of the sequenced members of the *Rhodobacteraceae* family, as well as strains of SAR11, SAR324, SAR116 and in marine *Gammaproteobacteria* (González et al., 1999; González, 2003; Howard et al., 2006; Bürgmann et al., 2007; Reisch et al., 2008; González et al., 2019). This phylogenetic distribution suggests an expansion of *dmdA* through horizontal gene transfer events (HGT) between different lineages of bacteria, presumably through viruses (Raina et al., 2010). Since the genome expansion of *Roseobacter* coincides with the diversification of the dinoflagellates and coccolithophores around 250 Mya (Luo et al., 2013; Luo & Moran, 2014; Bullock et al., 2017) it has been suggested a co-evolutionary event between *Roseobacter* and the DMSP-producing-phytoplankton (González et al., 1999; Zubkov et al., 2001; Moran et al., 2007; Bullock et al., 2017). Under this scenario, the enzymes of the DMSP demethylation pathway could have evolved within the last 250 Mya, as phytoplankton responded to the marine catastrophe at the end of the Permian with the diversification of dinoflagellates that produce DMSP and *Roseobacter* clade expanding by using DMSP as its main sulfur source. Despite this hypothesis, there is a lack of knowledge about the main evolutionary events that lead the DMSP adaptation in *Roseobacte*r.

In terms of production, the biosynthesis of DMSP has been reported in marine heterotrophic bacteria, such as the *Alphaproteobacteria*, i.e. *Labrenzia aggregata* (Curson et al., 2017). Since a common ancestor within the *Roseobacter* originated in the Archean, more than 2 billion years ago (Kumar et al., 2017), the *Roseobacter* and other *Alphaproteobacteria* might have been exposed to this DMSP early (Reisch et al. 2011a,b). According to this hypothesis, the DMSP demethylation and the cleavage pathways arose by the evolution of enzymes that were already present in bacterial genomes and adapted in response to the wide availability of DMSP. As mentioned earlier, *Alphaproteobacteria* in the SAR11 seems to thrive at the expense of organic sulfur compounds, such as DMSP and has a common ancestor that lived ca. 826 Mya, at the end of the Precambrian (Luo et al., 2013). We would then expect a common ancestor of the DmdA gene family during the early Proterozoic Mya and that the functional divergence between DmdA and GcvT gene families was driven by both functional constraints and widespread HGT. Probably in the Huronian snowball earth, a period of planetary crisis where the greatest microbial diversity took refuge in the shallow seas close to the equator (Tang, Thomas, & Xia, n.d.).

Here, we analyzed the evolutionary history of the DmdA gene family in marine *Proteobacteria* by considering three evolutionary scenarios: 1) a recent common ancestry of DmdA and GcvT, 2) a coevolution between *Roseobacter* and the DMSP-producing-phytoplankton, and 3) pre-adapted enzymes to DMSP prior to *Roseobacter* origin. We first analyzed if convergent, independent or HGT-based evolution can explain the presence of *dmdA* genes in different bacterial lineages of SAR11, SAR116 and *Rhodobacteraceae*. Then, we inferred the most recent common ancestor (MRCA) of the DmdA gene family, the timing of its origin and any duplication events. We also reconstructed the ancestral forms of DmdA enzymes to infer the most likely ecological conditions where DmdA thrive. We provide insights into their function by analyzing DmdA structural evolution. Finally, we examined how natural selection could have driven the divergence of the DmdA gene family. Our results indicate that *dmdA* appeared before the origin of *Roseobacter* clade and the conditions of the late Permian created by eukaryotic phytoplankton. Therefore, DmdA is an adapted version of enzyme that evolved in response to the availability of DMSP.

## METHODS

### Data mining

*DmdA* orthologs and *dmdA* homologs were collected from a set of 771 genomes manually curated and hosted in the MarRef database (Klemetsen et al., 2018). The sequences were obtained as described by González et al. (2019). The DmdA homologs included were obtained using a HMM designed for DmdA orthologs (González et al., 2019), with a relaxed maximum e-value (e-50). A total of 204 sequences from 184 genomes were used to infer the evolutionary history of DmdA gene family (Supplementary Table 1).

### Phylogenetic tree reconstruction and topology tests

The phylogenetic tree of the DmdA protein sequences included DmdA orthologs and DmdA homologs (called non-DmdA). The sequences were aligned using MUSCLE (Edgar, 2004). Regions poorly aligned or with gaps were removed using TrimAl (Capella-Gutiérrez et al., 2009) with parameters set to a minimum overlap of 0.55 and a percent of good positions to 60. Best-fit evolutionary model was selected based on the results of the package ProtTest 3 (Darriba et al., 2011) to determine the best-fit model for maximum likelihood (ML) and Bayesian inference (BI).

For the maximum likelihood analysis, PhyML v3.0 (Guindon et al., 2010) or RaxML v7.2.6 (Stamatakis, 2006) were used to generate 100 ML bootstrap trees, using the Le Gascuel (LG) model with a discrete gamma distribution (+G) with four rate categories, as this was the model with the lowest Akaike information criterion and Bayesian information criterion score. For the Bayesian analysis, trees were constructed using the PhyloBayes program (Lartillot & Philippe, 2004, 2006; Lartillot et al., 2007) with the CAT model that integrates heterogeneity of amino acid composition across sites of a protein alignment. In this case, two chains were run in parallel and checked for convergence using the tracecomp and bpcomp scripts provided in PhyloBayes. As an alternative, we computed a phylogenetic tree using a Bayesian inference implemented in BEAST2 program which was run with relaxed clock model and Birth Death tree prior (Bouckaert et al., 2014). Finally, we used R v3.6.1 (R Core Team, 2017) with phangorn v2.5.5 (Schliep, 2011) to perform consensus unrooted tree.

We ran several topology tests to establish whether the trees generated using the ML and BI methods provided an equivalent explanation for the two main groups, i.e., the non-DmdA and DmdA clades. For this analysis, the topologies were compared with the TOPD/FMTS software v4.6 (Puigbo et al., 2007). A random average split distance of 100 trees was also created to check if the differences observed were more likely to have been generated by chance.

### Horizontal gene transfer (HGT) test and GC content analysis

Two approaches were used to detect HGT. First, a phylogenetic incongruence analysis (Ravenhall, Škunca, Lassalle, & Dessimoz, 2015) through three topology tests, the Kishino-Hasegawa (KH) (Kishino & Hasegawa, 1989), the Shimodaira-Hasewaga (SH) (Shimodaira & Hasegawa, 1999) and the approximately unbiased (AU) (Shimodaira, 2002), implemented in the IQ-TREE software v1.5.5 (Nguyen et al., 2015). Two topologies were tested, the ML topology obtained for the species tree of the genomes here analyzed, and the ML phylogeny of DmdA. To construct the species tree, ribosomal protein 16 small subunit (RPS16) sequences were collected from the MarRef database (Klemetsen et al., 2018), one for each genome (Supplementary Table 1).

The GC content variation was studied to identify genes that have a different percentage of GC content at the third position of codons with respect to the neighboring genomic regions. The EPIC-CoGe browser (Nelson et al., 2018) was used to visualize the genomes and sequences and look for genes that use different codons with respect to the rest of the genomic dataset (data are available under permission as “ULL-microevolution” on https://genomevolution.org/).

### Molecular dating

We first tested for heterogeneities in the substitution rates of the genes using a likelihood ratio test (LRT) (Felsenstein, 1981) with the ML-inferred tree. Likelihoods’ values were estimated using baseml in PAML v4.8 (Yang, 2007) under rate constant and rate variable models and used to compute the likelihood ratio test (LRT) statistic according to the following equation:

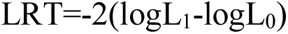

where L_1_ is the unconstrained (nonclock) likelihood value, and L_0_ is the likelihood value obtained under the rate constancy assumption. LRT is distributed approximately as a chi-square random variable with (m-2) degrees of freedom (df), m being the number of branches/parameters.

To conduct a molecular dating analysis with BEAST 2 (Bouckaert et al., 2014), two independent MCMC tree searches were run for 50 million generations, with a sampling frequency of 1000 generations over codon alignment obtained, as we explain in the next section. The GTR substitution model with a gamma shape parameter and a proportion of invariants (GTR + G + I), was selected with PartitionFinder software v2.1.1 (Lanfear et al., 2016) based on the Bayesian Information Criterion (Darriba et al., 2012), applied with a Birth Death tree prior (Gernhard, 2008) and an uncorrelated relaxed clock log-normal. The molecular clock was calibrated using information from the TimeTree database (Hedges et al., 2006, 2015; Kumar et al., 2017). We used the dates of the most recent common ancestor of (1) the *Alpha-* and *Gammaproteobacteria* (2480 Mya), (2) the *Halobacteriales* (455 Mya) (Supplementary Fig 1-3), and (3) the SAR11 (826 Mya) (Luo et al., 2013). A log-normal prior distribution on the calibrated nodes centered at the values mentioned above was specified with 20 standard deviations and constrained to be monophyletic. Convergence of the stationary distribution was checked by visual inspection of plotted posterior estimates in Tracer v1.6 (Rambaut, & Drummond, 2013) to ensure effective sample sizes (ESSs) of parameters were >> 200, as recommended by the authors. After discarding the first 15% trees as burn-in, the samples were summarized in the maximum clade credibility tree using TreeAnnotator v1.6.1 (Rambaut, & Drummond, 2002) with a PP limit of 0.5 and summarizing mean node heights. Means and 95 % higher posterior densities (HPDs) of age estimates are obtained from the combined outputs using Tracer v1.6. The results were visualized using FigTree v.1.4.3 (Rambaut, 2009).

### Maximum likelihood tests of positive selection

To measure the strength and mode of natural selection during the evolution of DmdA gene family, the ratio of non-synonymous (dN) to synonymous substitutions (dS) (ω=dN/dS) was calculated in CodeML implemented in the suite Phylogenetic Analysis by Maximum Likelihood (PAML package v4.8) (Yang, 2007).

CodeML requires an alignment of coding sequences, and a phylogenetic tree. DNA alignment was achieved by MUSCLE (Edgar, 2004) implemented in MEGA-CC v7.0.26 (Kumar et al., 2016) and poorly aligned segments were eliminated with Gblocks under defaults parameters (Castresana, 2000). The phylogenetic tree was built using ML with PhyML v3.0 (Guindon et al., 2010) as described above and a nucleotide substitution model selected by jModelTest (Darriba et al., 2012). DAMBE (Xia, 2001) was also used to check for saturation of nucleotide substitutions using a plot of the number of transitions and transversions for each pairwise comparison against the genetic distance calculated with the F84 model of nucleotide substitution (Huelsenbeck & Rannala, 1997), which allows different equilibrium nucleotide frequencies and a transition rate-transversion rate bias. Multiple sequence alignments with similar characteristics (i.e., showing saturation of nucleotide substitutions) were then analyzed with CodeML (Yang, 2007).

Three sets of models were used (site-specific, branch-specific and branch-site models) to detect pervasive and episodic selection during the evolution of *dmdA* orthologs. Likelihood-ratio tests (LRTs) were used to compare models, and significant results (p-value<0.05) were determined contrasting with a chi-square distribution (chisq) (Anisimova et al., 2001).

In the site-specific analysis, we tested for variability of selection (type and magnitude) across the codons of the gene using three pairs of nested models. The first pair includes M0 (just one dN/dS ratio) and M3 (“K” discrete categories of dN/dS) and has four degrees of freedom (df). The second pair of models considers M1a (just two classes of sites, purifying [dN/dS<1] and neutral selection [dN/dS=1]) and M2a (the same as M1a adding a third class of sites dedicated to positive selection [dN/dS>1]), this has two df. Finally, the third pair of models comprised M7 (a beta distribution that allows dN/dS to vary among the interval [0,1]) and M8 (adds an extra discrete category to M7 with dN/dS>1), with two df. Whereas M0 vs M3 test for evidence of dN/dS variation across sites, M1a vs M2a and M7 vs M8 test for the presence of sites under positive selection (dN/dS > 1).

Using three branch models (Yang, 1998), we tested for variation of selection over evolutionary time. The null model (M0) assumes that all branches evolve at the same rate, therefore, there is only one value of dN/dS for all the branches of the tree. The two-ratio model allows two dN/dS values, one value for all *Roseobacter* lineages (we called this group A) and another for the rest of branches (named group B). The free-ratio model, allows one dN/dS value for each branch. Null and two-ratio model are compared by LRT with one df but null and free-ratio model are compared with 36 df.

For the last set of models, we identified sites that have been under positive selection at a particular point of evolution using branch-site models, in which dN/dS can vary among sites and among branches (Zhang, 2005). We computed two models: a null model, in which the “foreground branch” may have different proportions of sites under neutral selection to the “background branches”, and an alternative model in which the “foreground branch” may have a proportion of sites under positive selection. We compare these models for each terminal branch with a LRT of one df. For each branch-site analysis, we applied the Bonferroni correction for multiple testing.

In site and branch-site tests, we identified sites under positive selection as those with Bayes Empirical Bayes (BEB) posterior probability above the 0.95 (Yang, 2005). We also checked for convergence of the parameter estimates in PAML by carrying out at least two runs for each tree and starting the analysis with different ω (0.2, 1, 1.2 and 2). In addition, to test for convergent selection in several lineages, we ran at Branch-site analysis selecting as “foreground branches” all those under positive selection in a previous analysis.

### Analysis of functional divergence

Divergent selection is indicated by different ω’s values among paralogous clades. We tested whether selective pressures diverged following duplication that led to *dmdA* and non-*dmdA* genes (Bielawski & Yang, 2004). We compared the M3 model, which accounts for ω variation among sites but not among branches or clades, with a model allowing a fraction of sites to have different ω between two clades of a phylogeny (clade model D). We also tested M0 and M3 models and we used a posterior BEB probability above the 0.95 to identify sites evolving under divergent selective pressures. We checked for convergence of the parameter estimates in PAML by carrying out at least two runs for the tree and starting the analysis with different ω (0.1, 0.25, 2, 3 and 4).

Finally, we applied two branch-site models (as described above) to test dN/dS differences on the branches representing the ancestral lineages of the DmdA and non-DmdA clades (see results) (Supplementary Fig 25). We considered the ancestral sequences from DmdA and non-DmdA clades as foreground branches in two different models.

### Reconstruction of ancestral DmdA sequence

To reconstruct the ancient conditions where *dmdA* gene prospered, we inferred the ancestral sequences of the DmdA node using the FastML web server (Ashkenazy et al., 2012) and then computed estimated physico-chemical properties on predecessor sequence using Compute ProtParam tool from Expasy – SIB Bioinformatics Resource Portal (Gasteiger et al., 2005). Moreover, we also reconstructed the ancestral sequence of the non-DmdA node, as well as the ancestral sequence of both the DmdA, and the non-DmdA families. FastML was run considering the alignment of proteins and the ML phylogenetic tree for those DmdA orthologs or homologs inferred as we explained above. Posterior amino acid probabilities at each site were calculated using the Le Gascuel (LG) matrix (Le & Gascuel, 2008) and Gamma distribution. Both marginal and joint probability reconstructions were performed. Protein sequences resulting from marginal reconstructions were used to predict tertiary structure (see below) as well as to identify family domains using Pfam v32 (Finn et al., 2010).

### Protein tertiary structure analysis

Predicted three-dimensional structures of protein sequences were examined by Iterative Threading ASSEmbly Refinement (I-TASSER) (Roy et al., 2010; Yang et al., 2015). First, I-TASSER uses local meta-threading-server (LOMETS) (Wu & Zhang, 2007) to identify templates for the query sequence in a non-redundant Protein Data Bank (PDB) structure library. Then, the top-ranked template hits obtained are selected for the 3D model simulations. To evaluate positively the global accuracy of the predicted model, a C-score should return between -5 and 2. At the end, top 10 structural analogs of the predicted model close to the target in the PDB (Berman et al., 2000) are generated using TM-align (Zhang, 2005). The TM-score value scales the structural similarity between two proteins, and should return 1 if a perfect match between two structures is found. A TM-score value higher than 0.5 suggests that the proteins belong to the same fold family.

We used PyMol v1.7.4 (DeLano, 2002) to visualize the 3D structure of the proteins and to map the positively selected sites onto the 3D structure of DmdA (pdb: 3tfh).

## RESULTS

### Phylogenetic tree for DmdA family

We identify a total of 204 DmdA protein sequences out of 150 curated genomes, and reconstruct their evolutionary relationships by Bayesian Inference (BI) (Fig 1) and Maximum Likelihood (ML) (Supplementary Fig 4). Unrooted trees in TOPD-FMTS indicated that split distances did not exceed 0.19, indicating that the phylogenetic reconstruction is robust, with minor variations in alignment filtering and methods for inferring topologies (Supplementary Table 2).

**Fig 1.**
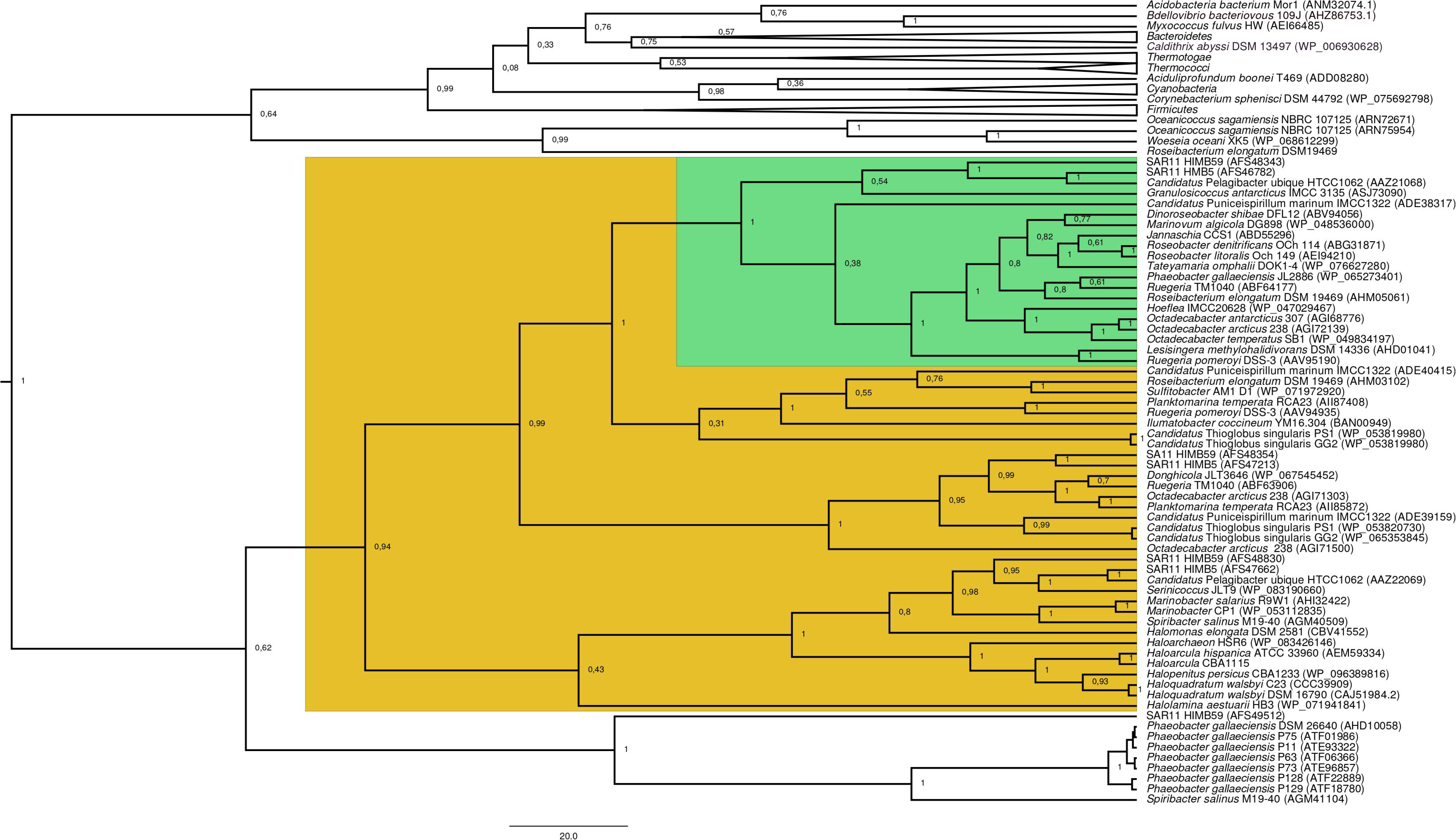
GcvT phylogenetic tree based on 20 DmdA orthologs protein sequences and 184 DmdA homologs using Beast and the same parameters set for molecular dating but with 100 million generations. DmdA sequences are indicated with green color and closer homologs for those with yellow color. Tip labels include a maximum e-value < e-50.

The BI tree (Fig 1) shows a main duplication between two lineages. The larger phylogenetic group comprises genes from *Bacteroidetes*, while the smaller group includes genes from *Alphaproteobacteria*. We focused on this smaller group as it includes the DmdA sequences (Fig 1; green color) and the closest homologs to DmdA (Fig 1; yellow color).

Using phylogenetic analyses including DmdA orthologs and DmdA homologs close to those (the limit to select closer homologs was set to a maximum e-value of e-80) we resolve the position of the first DmdA sequences isolated from two marine bacterial species, *R. pomeroyi* (AAV95190.1) and *Ca.* P. ubique (AAZ21068.1). In addition, the inclusion of DmdA homologs allowed to resolve a robust phylogenetic relationship of DmdA gene family (Fig 2). We detected a clear separation between DmdA and putative non-DmdA families. Indeed, the four DmdA family trees constructed using different methods compared in TOPD-FMTS using split distances (Supplementary Table 3) and unrooted trees (Supplementary Fig 5) agreed with this result. The average split distance was 0.60, indicating that the trees were neither identical (split difference=0) nor completely different (1). A random split distance was calculated to analyze whether the split distances were significantly different. Because the random split distance resulted in a value close to 1 (0.988), our observations are unlikely to be given by chance.

**Fig 2.**
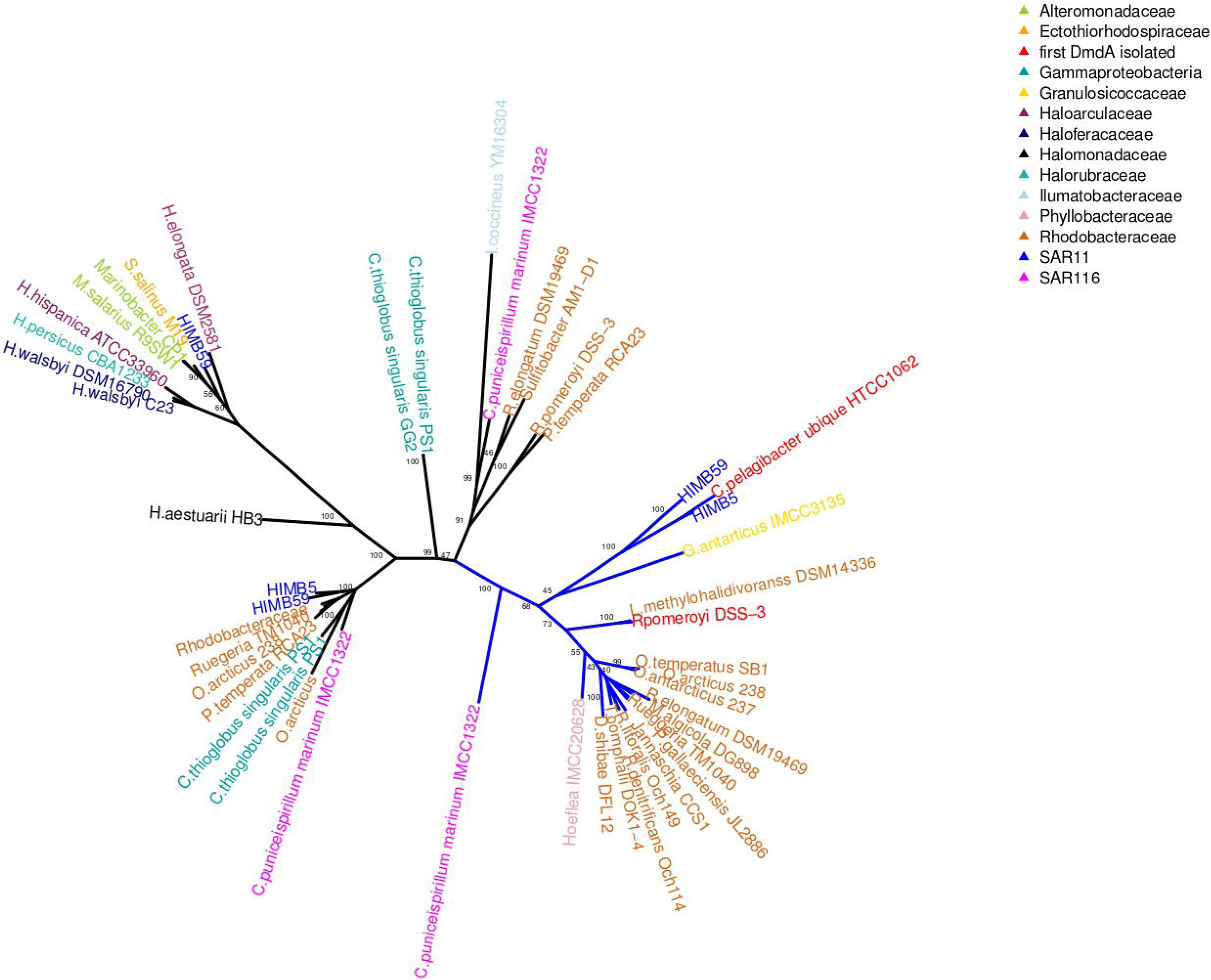
Phylogenetic tree of DmdA based on 20 DmdA orthologs protein sequences and 28 DmdA homologs (more information in Supplementary Table 1) using RaxML. A non-parametric bootstrap is shown to establish the support for the clades. DmdA sequences are indicated with blue branch. Tip labels show color for first dmdA gene identified or taxonomy classification. Tip labels include a maximum e-value <e-80.

To identify HGT and duplication events, we constructed a proxy for the species tree of the genomes considered here by using a set of small subunit ribosomal protein (see Material and Methods).

Given this (proxy) species tree (Supplementary Fig 6), the positions of many sequences on the DmdA tree are better explained as cases of HGT (Supplementary Fig 6; Fig 3) with high statistical support. We then tested whether the topology for a common set of taxa within the DmdA family (Supplementary Fig 7) were similar to that of the species tree (Supplementary Fig 8). We found significant differences (at an alpha of 0.01) between the topology of DmdA group and that of the proxy species tree (Table 1); this incongruence between phylogenies is true irrespective of the test used (Kishino-Hasegawa, Shimodaira-Hasewaga and unbiased tests). From these results we conclude that the phylogenetic relationships within each DmdA group are different to those of the species tree, strongly supporting a HGT-based evolution of DmdA family (Supplementary Fig 9).

**Fig 3.**
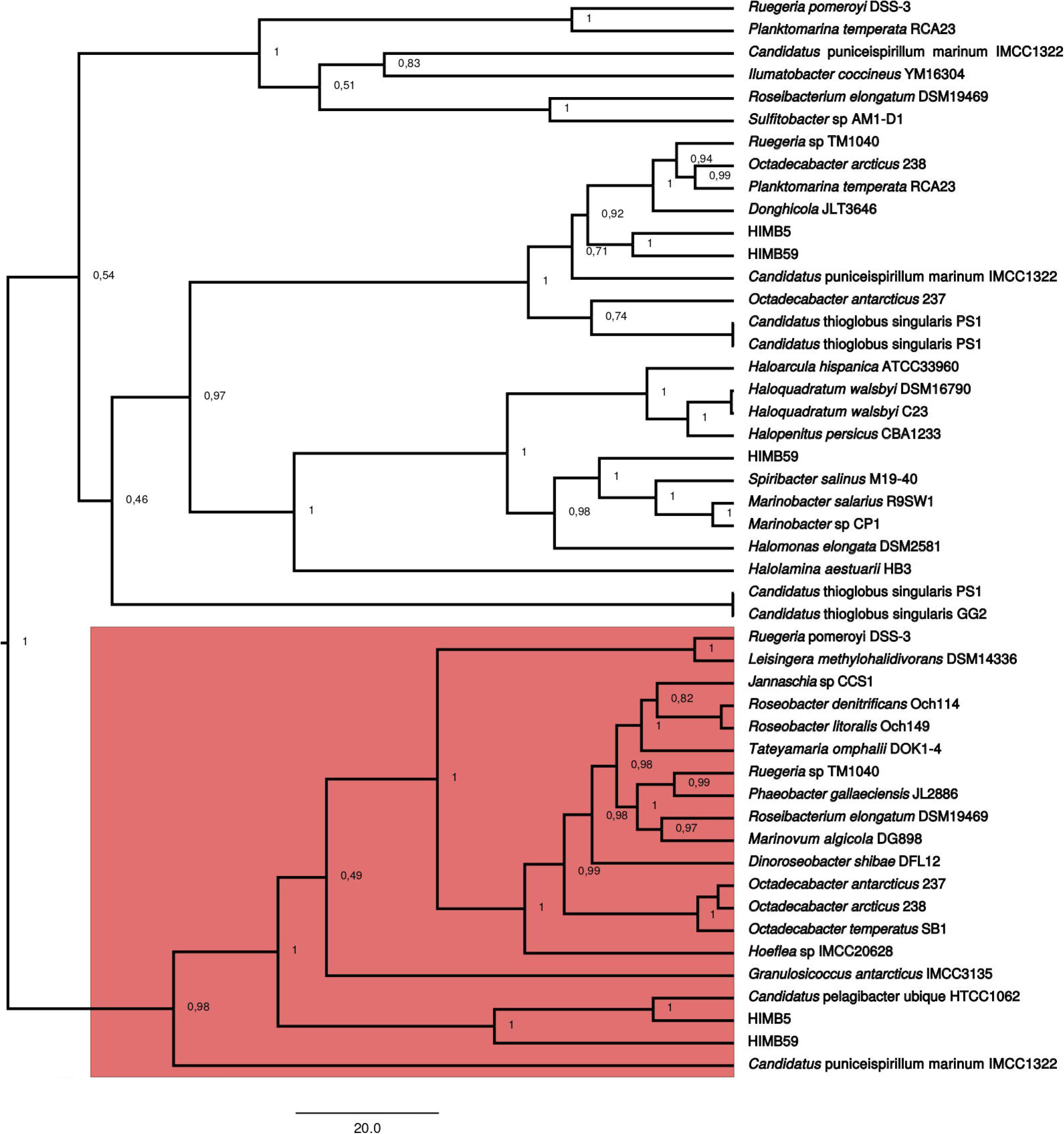
Phylogenetic tree of DmdA based on 20 DmdA orthologs protein sequences and 28 DmdA homologs using BEAST2. Bayesian posterior probabilities (PP) is shown to establish the support for the clades. Red color indicates DmdA clade.

**Table 1.** Topology tests of DmdA phylogenetic tree with respect to species tree.

| Group | pKH* | pSH* | pAU* |
| --- | --- | --- | --- |
| DmdA family | 0.0010 | 0.0010 | 0.0001 |

\*p-values under the Kishino-Hasegawa (KH) test, the Shimodaira-Hasegawa (SH) test and the approximately unbiased (AU) test, respectively.

Moreover, we found many genes that use different codons than the neighboring genomic regions. These genes are inferred as having been horizontally transferred given their (G+C) wobble content (Supplementary Table 1), supporting an HGT-based evolution of DmdA family (Supplementary Fig 9).

### Structural modeling

The structure for DmdA orthologs inferred on the protein sequences by Iterative Threading ASSEmbly Refinement (I-TASSER) were threaded onto the known structure of DMSP-dependent demethylase A protein (PDB accession: 3tfhA) with a C-score<= 2 (Table 2). However, the predicted models for DmdA homologs were threaded onto two types of known structure; DmdA orthologs, and the structure of the mature form of rat dimethylglycine dehydrogenase (DmgdH) (PDB accession, 4ps9sA) with a C-score < 2 except for the sequence with accession number AEM59334.1, which shows a C-score > 2 (Supplementary Fig 10a, Supplementary Data 1).

**Table 2.**
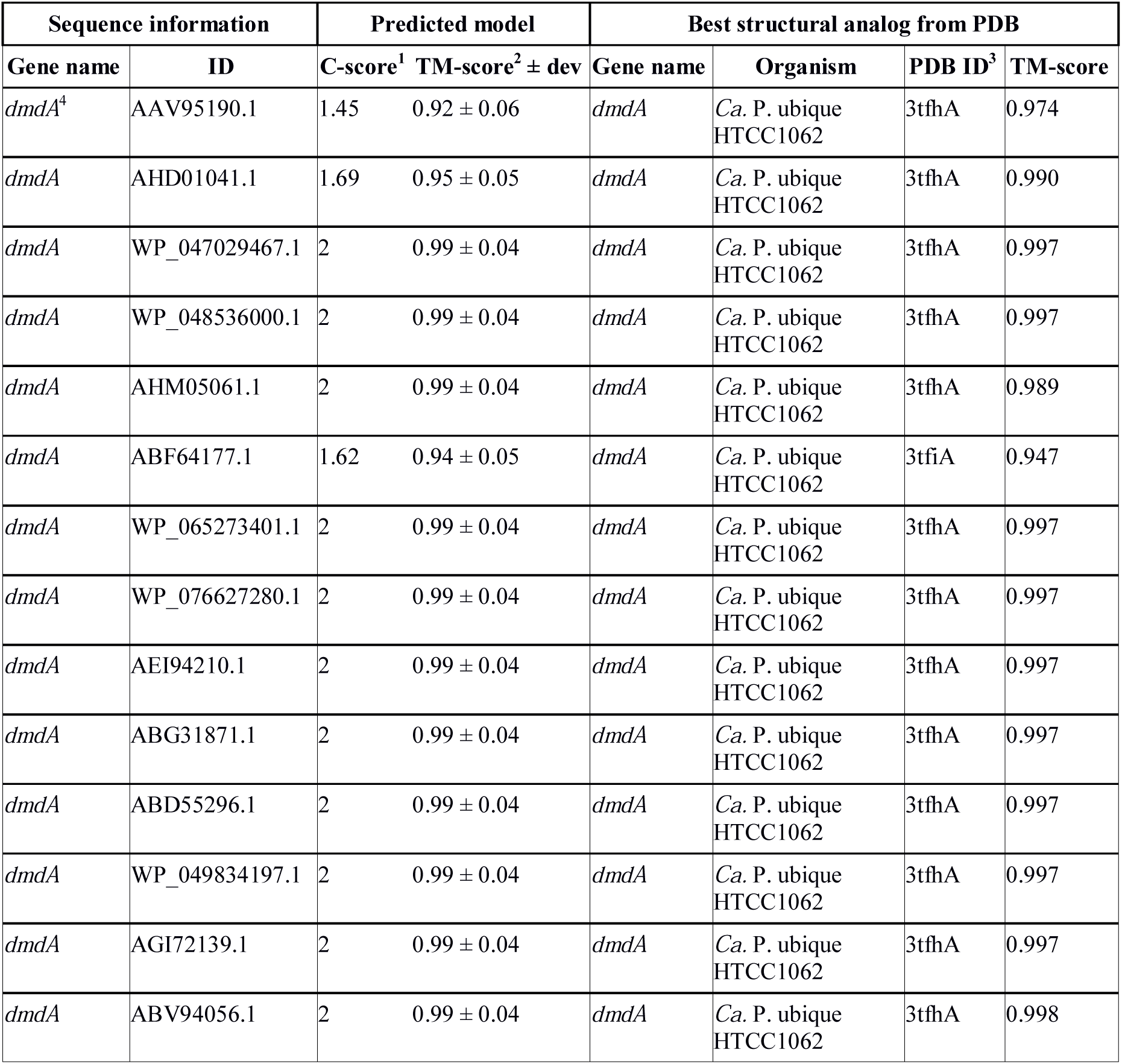
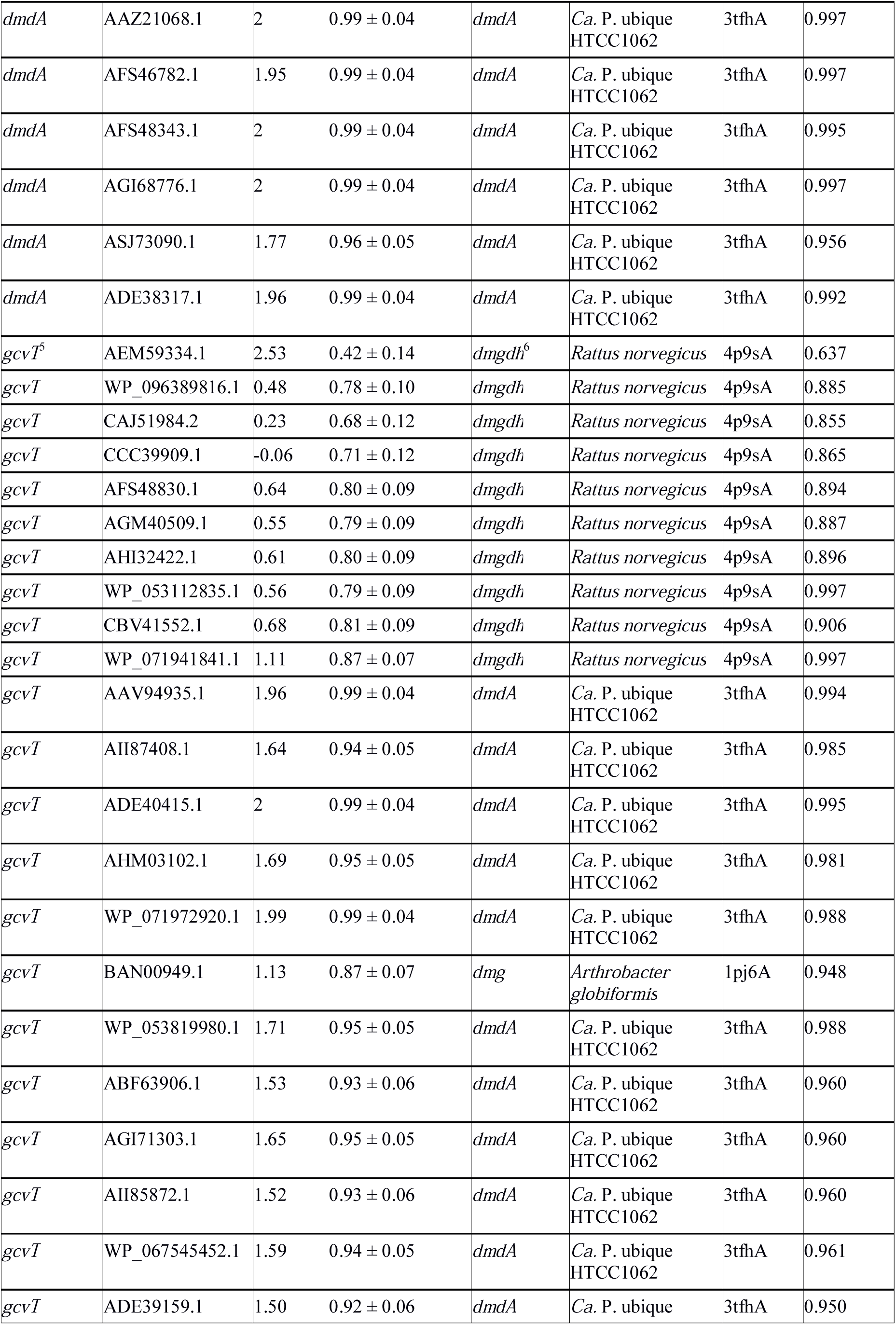
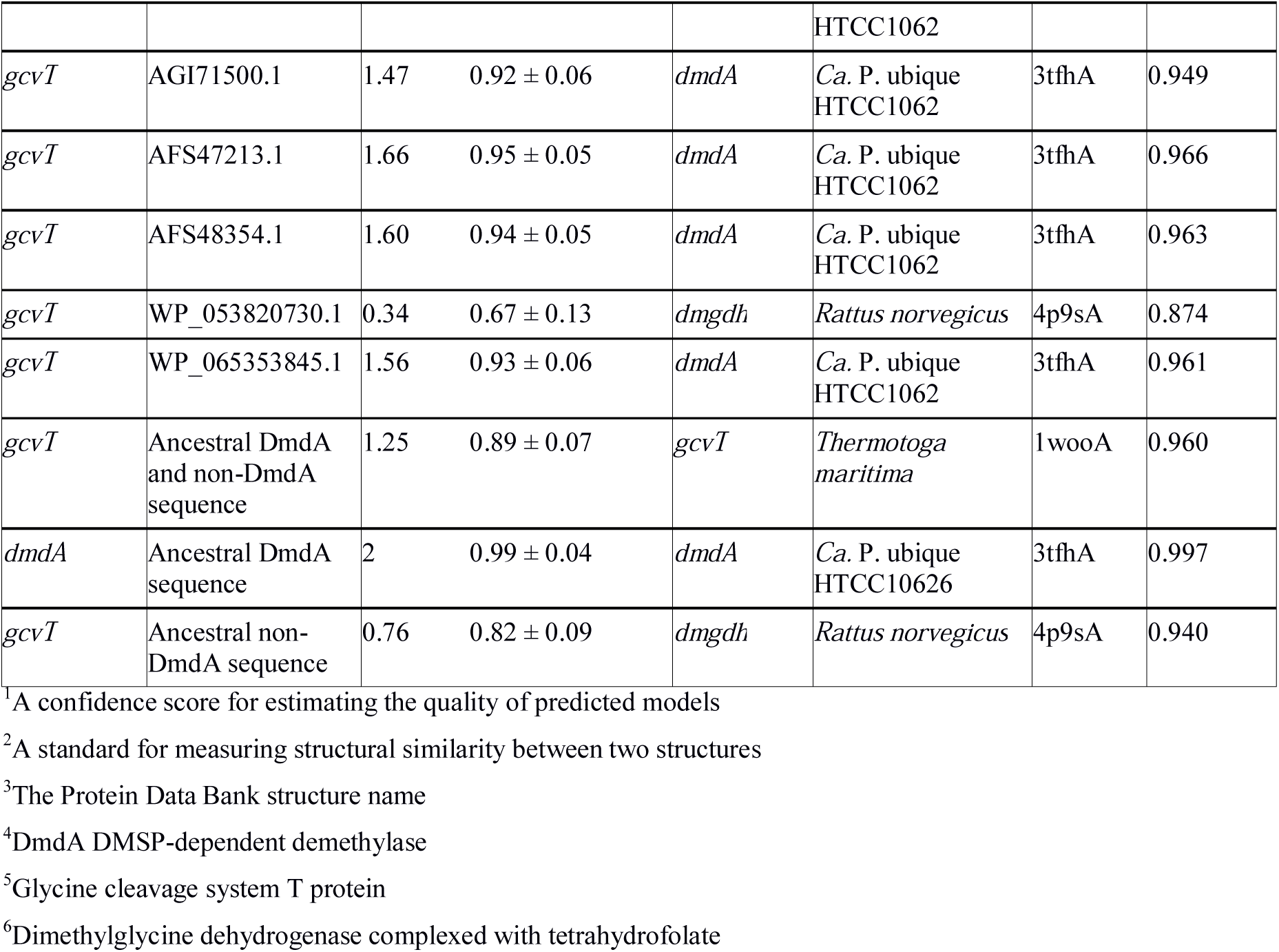
Structural model predicted by I-TASSER for each sequence used in the evolutionary study of DmdA gene family and the best identified structural analogs in PDB by TM-align.

We clustered sequences with a putative DmgdH structure in a separate group using principal component analysis (Supplementary Fig 11). There is a clear 3D-structure coincidence between DmdA clade (red color in Supplementary Fig 10a) and the majority of lineages from non-DmdA clade (orange color in Supplementary Fig 10a) as well as a conserved folate-binding domain (Supplementary Fig 10b: 99S, 178E and 180Y). However, in the alignment we found a pattern of conserved residues coherent with phylogeny results (Supplementary Fig 10a, Supplementary Fig 10b), where non-DmdA clade is formed by three subclades, one of them with DmgdH tertiary structure. Indeed, key residue for DMSP specific interaction is shown in clades with DmdA tertiary structure (Supplementary Fig 10b: W171) but not in a clade with DmgdH tertiary structure (Supplementary Fig 10b: F171).

### Molecular dating

The log likelihood test (LRT) detected heterogeneity in the substitution rates of *dmdA* orthologs and *dmdA* homologs genes (Fig 2) (log L_0_=-29,827.108; log L_1_= -29,546.053; degrees of freedom = 46; x^2^ = 562.11; P<0.001), thus rejecting the hypothesis of a strict molecular clock. This finding validates the use of relaxed molecular clock approach to estimate the node ages throughout Bayesian analysis (see Methods for details). We observed that the marginal densities for each run of the divergence time estimate analysis were nearly identical, pointing that the runs converged on the same stationary distributions. In all runs, the marginal densities for the standard deviation hyperparameter of the uncorrelated log-normal relaxed clock model were quite different from the prior, with no significant density at zero and with a coefficient of variation around 0.2. Analyses using three different calibrated prior dates showed not discrepancies in the final divergence time estimates (Table 3).

**Table 3.** Divergence time estimates in million years ago (Mya), and node 95% highest posterior density (HPD) interval for the clades of the most recent common ancestor (MRCA) of *Halobacteriales*, SAR11 and *Alphaproteobacteria* from each set of calibration priors.

| Taxonomic group of MRCA | Clade | Age | 95% HPD |
| --- | --- | --- | --- |
| *Halobacteriales* (455) | Mrca1 | 438 | 311.1 – 572.3 |
| SAR11 (826) | Mrca2 | 827.5 | 588.3 – 1089.8 |
| *Alphaproteobacteria* (2480) | Mrca3 | 2118.6 | 1543 – 2717.1 |

The time estimates for the MRCA of each gene family (Table 3 and Fig 4) indicate that the most recent common ancestor of DmdA gene family occurred in the late Archean, around 2,400 Mya, after a gene duplication event. Also, a duplication within the DmdA lineage generated a separated SAR11 and *Roseobacter* DmdA lineage in the early Precambrian ca. 1,894 Mya (Fig 4: red arrow). *Ca.* P. ubique HTCC1062 within the first cluster and *R. pomeroyi* DSS-3 within the second cluster, resulted from a duplication around 300 Mya (Fig 4: blue arrow). However, a higher number of duplication events took place in the second cluster. Thus the number of paralogous genes comprising the *Roseobacter* DmdA family is larger than in SAR11 (Fig 4).

**Fig 4.**
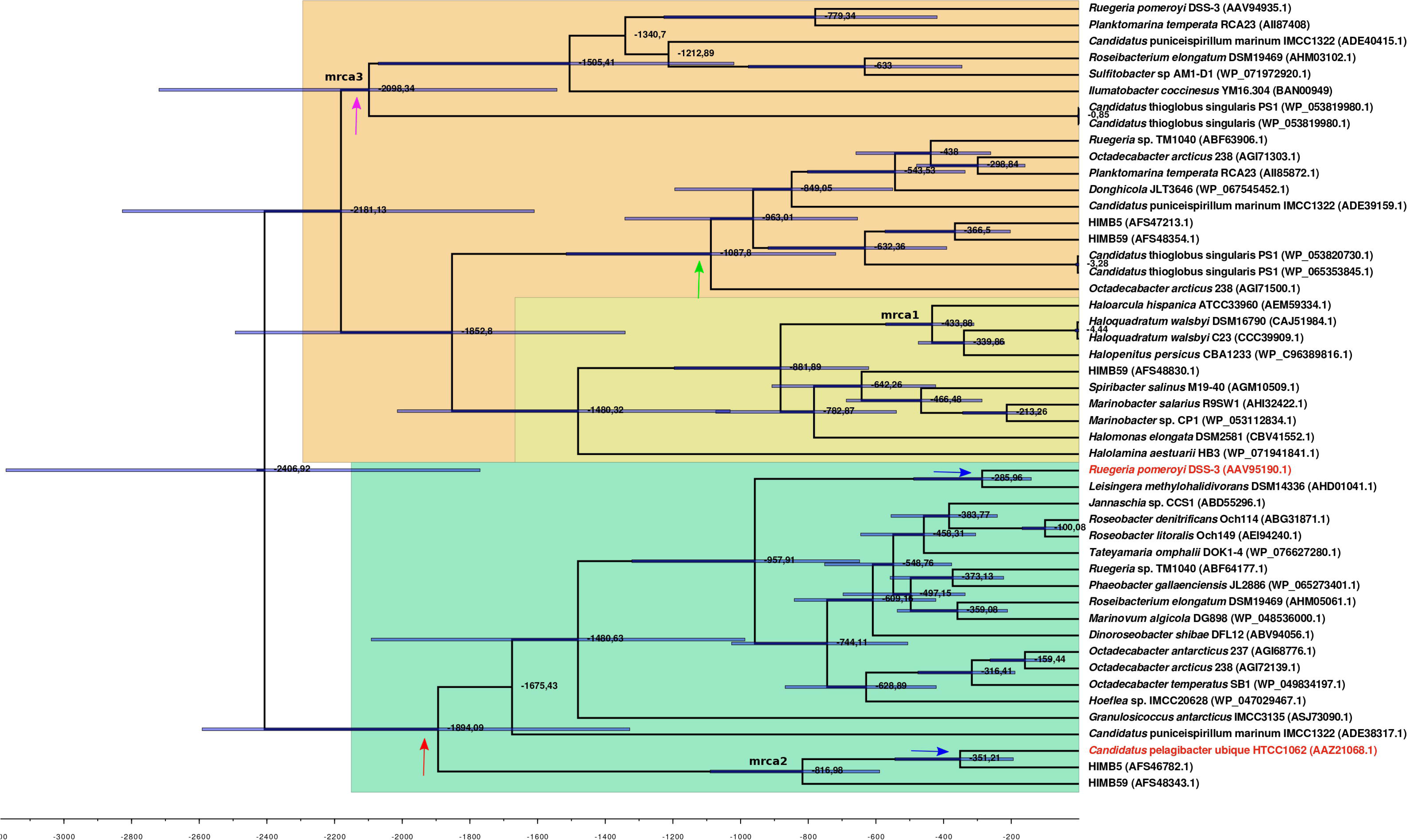
(Upper) BEAST divergence time estimates from *dmdA* and non-*dmdA* genes under uncorrelated relaxed clock model and Birth-death tree model. Nodes are at mean divergence times and gray bars represent 95% HPD of node age. Nodes used as calibrated priors in BEAST analysis are marked as mrca1, mrca2 and mrca3 as well as colored. (Lower) Absolute time scale in Ma. Arrows indicate duplication events occurred 1894 Mya (red), 300 Mya (blue) and 1000 Mya (green).

We detected two duplication events within the putative non-DmdA clade (Fig 4; orange color); showing that the gene families were originated through old duplication events. One duplication involving the DmgdH family (Fig 4 dark yellow color; Table 2) occurred 1,480 Mya and another duplication 1,000 Mya (Fig 4: green arrow), with tertiary structure similar to the DmdA from *Ca.* P. ubique. The other event of duplication took place during the Huronian glaciation, around 2100 Mya (Fig 4: violet arrow).

## Reconstruction of ancestral DmdA sequence

Our analysis was focused on the reconstruction of the ancestral sequences of the DmdA clade, the non-DmdA clade as well as the ancestral sequence of both the DmdA and non-DmdA clades.

FastML inferred the 100 most likely ancestral sequences of the DmdA family. We observed that the same sequences were always inferred. Indeed, the difference in log-likelihood between the most likely ancestral sequence at this node (N1; Supplementary Fig 12) and the 100th most likely sequence was only 0.105, indicating that both sequences are almost as likely to reflect the “true” ancestral sequence. That ancestral protein contains both PF01571 (GCV_T) and PF08669 (GCV_T_C) domains, found in the DmdA orthologs and it is nearly identical to *Ca.* P. ubique HTCC1062 DmdA sequence. Moreover, PSI-BLAST search confirmed that the ancestral sequence in node 1 close to DmdA genes hosted in EMBL-EBI databases (Supplementary Fig 13) and the structure for *Ca*. P. ubique apoenzyme DmdA was the closest analog to our predicted models (Table 2; Supplementary Data 1). Inferred physico-chemical properties are identical between *Ca*. P. ubique and the DmdA ancestral sequence (Supplementary Table 4).

On the other hand, the ancestral sequence inferred for non-DmdA family (N1; Supplementary Fig 14) and the ancestral sequence previous to functional divergence (N1; Supplementary Fig 15) contains only the PF01571 domain. That domain was located onto the known structure of T-protein of the Glycine Cleavage System (PDB accession: 1wooA) with a C-score= 1.25 (Table 2; Supplementary Data 1) in the case of the ancestral DmdA and non-DmdA sequence. However, the ancestral sequence for non-DmdA was better threaded onto the known structure of mature form of rat DmgdH (PDB accession: 4p9sA) with a C-score= 0.76 (Table 2; Supplementary Data 1).

### Detection of positive selection on *dmdA* sequences

To infer how natural selection has influenced on the evolutionary history of DmdA gene family, we used an alignment of the 20 sequences clustered as *dmdA* orthologs. The phylogenetic tree for these sequences was constructed by ML using the symmetrical model (SYM) with a discrete gamma distribution.

The average dN/dS value for the *dmdA* gene was 0.085, suggesting that this gene evolved under strong negative (purifying) selection. Then, we analyzed dN/dS variation across the codons in the gene, comparing M0 and M3 models through a LRT. The M3 model had better fit to the data than the M0 model (chisq= 775.387, p-value< 0.01). All codons in the gene are under strong purifying selection with dN/dS <1 (Fig 5), suggesting the importance of this sulfur pathway for the cells. In accordance with this, the LTRs designed to detect codons under positive selection were not significant (M1 vs M2, chisq= 0 and p-value = 1, and M7 vs M8, chisq = 1.459 and p-value = 0.482). Hence, we did not detect sites in *dmdA* subjected to positive selection (Supplementary Fig 17).

**Fig 5.**
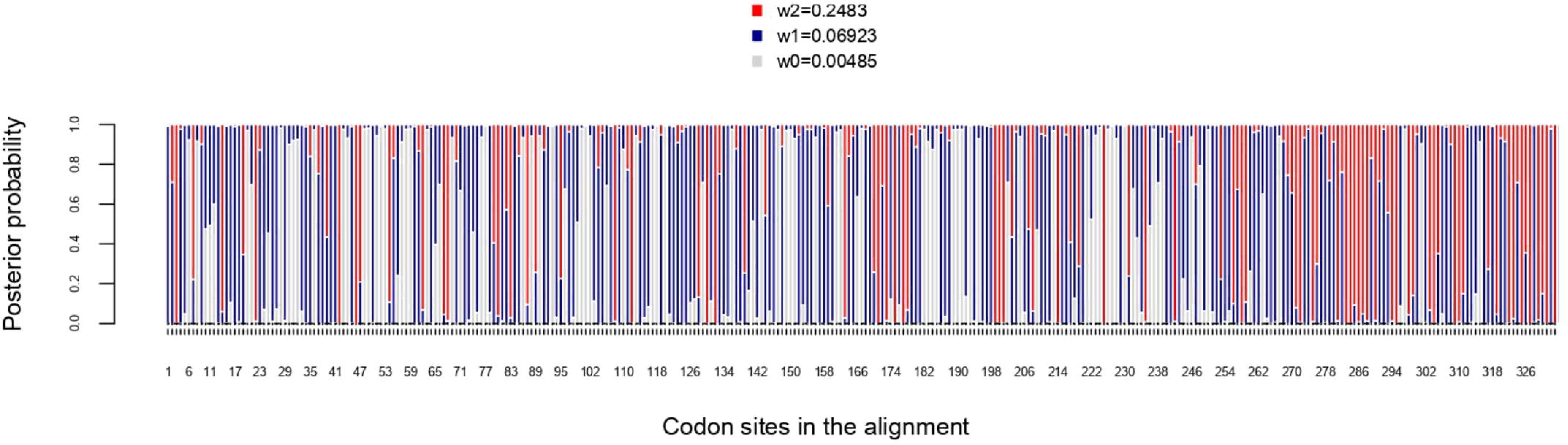
Posterior probabilities for dN/dS categories under the M3 model. Grey, red and blue bars depict the three dN/dS categories (values for each category are provide in the key). Sites that are mostly grey denote codons under strong purifying selection, whereas those predominantly red show codons under weaker purifying selection. Red, blue and grey colors indicate codon sites with ω_2_= 0.2483, ω_1_=0.06923 and ω_0_=0.00485, respectively.

We tested the variation in the intensity of selection over evolutionary time. A two-ratio model comparing the *Roseobacter* with the rest of lineages (Supplementary Fig 18) fits better the data, as the LRT was 23.777 and p-value < 0.01 (Table 4). dN/dS value in *Roseobacter* (ω_1_: 0.0767) was significantly lower than in the remaining branches (ω_2_: 0.1494), suggesting stronger purifying selection on *dmdA* in *Roseobacter*. When we tested the intensity of selection over evolutionary time using the free-ratio model (Table 4), we found changes in the selection pressure from the branches which defines the separation of SAR11 and *Roseobacter* DmdA gene families (Supplementary Fig 19: branches from nodes 21 to 23). In particular, we observed a dN/dS value > 1 in the branch connecting nodes 21-23. We also identified some more recent branches (connecting nodes 25-26 and 28-29) for which dN/dS >> 1 was estimated (Supplementary Fig 19).

**Table 4.** Parameters of branch-models.

| Model | $\omega_1$ | $\omega_2$ | $-\ln L^1$ | LRT<sup>2</sup> | P-value |
| --- | --- | --- | --- | --- | --- |
| One $\omega$ (one-ratio) | 0.08518 | NA | -14580.019867 | NA | NA |
| Two $\omega$ (two-ratio) | 0.0767 | 0.1494 | -14568.131038 | 23.777658 | 0.0 |
| 38 $\omega$ (free-ratio) | * | * | -14428.881747 | 302.27624 | 0 |

\* $\omega$ values are shown in Supplementary Fig 19.
<sup>1</sup>Log-likelihood score under the model
<sup>2</sup>Likelihood ratio test

Finally, we applied the two branch-site models to test for sites under selection on the individual lineages associated with *dmdA* (Supplementary Fig 20). Four sequences (WP_047029467, AHM05061.1, ABV94056.1, AFS48343.1) had a significant LRT after correcting for multiple testing (Table 5), suggesting episodic positive selection on these lineages (Supplementary Fig 20). It should be highlighted that three selected sites are shared by at least two lineages (Table 5; Fig 6). One shared site is located next to the GcvT domain (152 K; Supplementary Fig 21), and two shared sites are closed to conserved positions (17E; 87Y; Supplementary Fig 21). The residue 87Y is adjacent to the conserved interaction site with THF (88Y; Supplementary Fig 21). Interestingly, since the selected lineages are separated in the tree, the adaptive mutations seem to have occurred through three parallel independent changes (Supplementary Fig 22).

**Fig 6.**
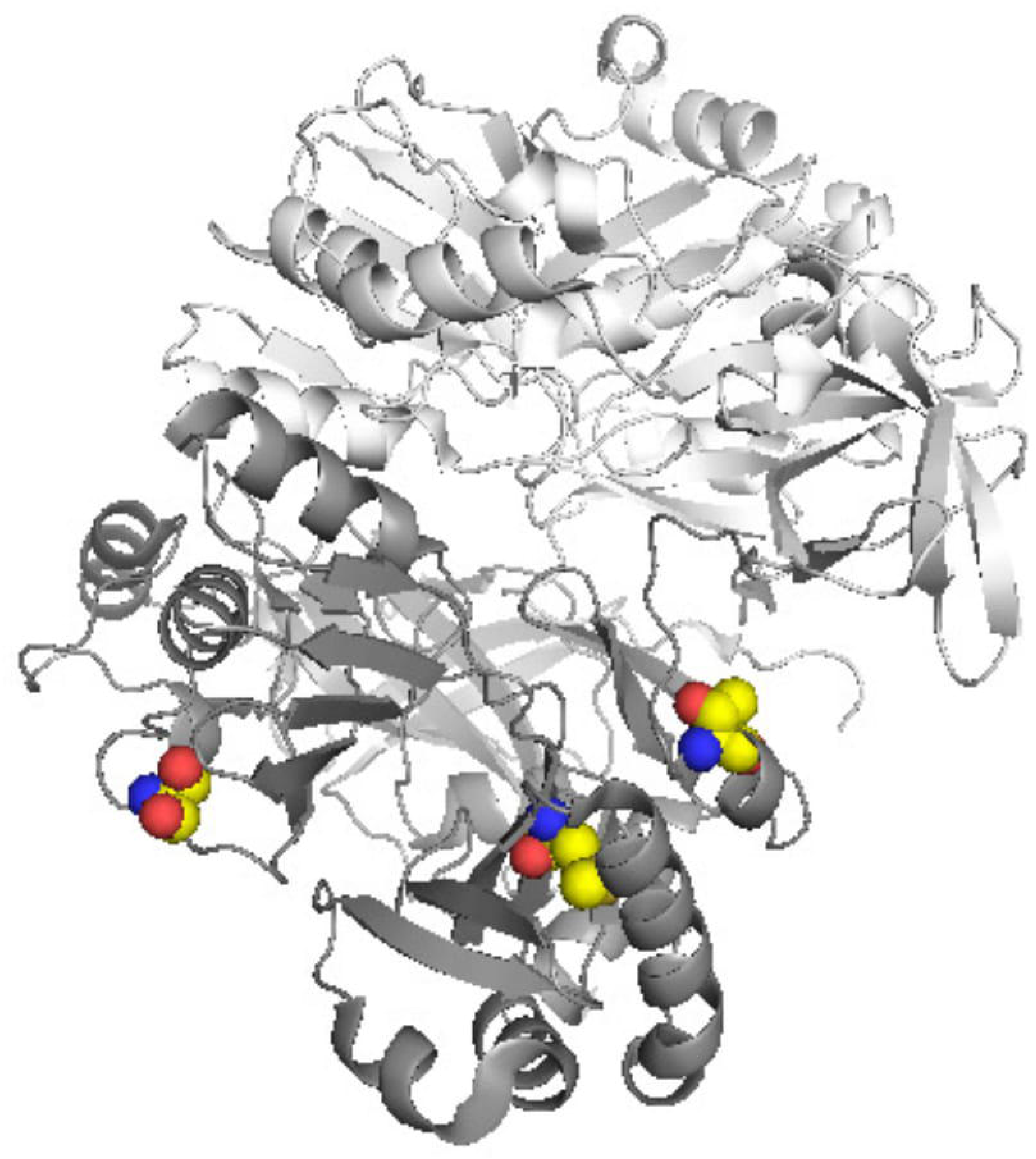
Tertiary structure of DmdA (PBD: 3tfh) with sites under episodic positive selection mapped in yellow color through Pymol.

**Table 5.** Parameters of PAML branch-site models.

| Branch | Ho (-lnL)<sup>1</sup> | Ha (-lnL)<sup>2</sup> | LRT<sup>3</sup> | P-value<sup>4</sup> | CorrectedP-value<sup>5</sup> | Pos. Selected sites* (BEB>0.95) |
| --- | --- | --- | --- | --- | --- | --- |
| ADE38317.1 | -14465.244 | -14463.099 | 4.290 | 0.038 | 0.767 | NA |
| AAV95190.1 | -14476.763 | -14476.763 | 0 | 1 | 1 | NA |
| AHD01041.1 | -14476.763 | -14476.763 | 0 | 1 | 1 | NA |
| WP_047029467.1 | -14476.763 | -14437.565 | 78.397 | 0.00 | 0.00 | 7V; **17E**; 47H; 65D; 68Y; **87Y**; 89A; **152K**; 157M; 163N; 203V; 279G; 290P; 319T; 320H |
| WP_048536000.1 | -14476.763 | -14476.763 | 0 | 1 | 1 | NA |
| AHM05061.1 | -14466.948 | -14460.844 | 12.206 | 0.000 | 0.000 | **17E** ; **152K**; 178E; 285V |
| ABF64177.1 | -14476.763 | -14476.763 | 0 | 1 | 1 | NA |
| WP_065273401.1 | -14476.763 | 14476.763 | 0 | 1 | 1 | NA |
| WP_076627280.1 | -14476.763 | 14476.763 | 0 | 1 | 1 | NA |
| AEI94210.1 | -14476.763 | -14476.763 | 0 | 1 | 1 | NA |
| ABG31871.1 | -14476.763 | -14476.763 | 0 | 1 | 1 | NA |
| ABD55296.1 | -14476.764 | -14476.764 | 0 | 1 | 1 | NA |
| WP_049834197.1 | -14476.763 | -14476.763 | 0 | 1 | 1 | NA |
| AGI72139.1 | -14476.763 | -14476.763 | 0 | 1 | 1 | NA |
| AGI68776.1 | -14476.763 | -14476.763 | 0 | 1 | 1 | NA |
| ABV94056.1 | -14462.942 | -14454.885 | 16.112 | 0.000 | 0.000 | **87Y**; **152K**; 243N; 247L; 257F |
| ASJ730990.1 | -14463.474 | -14461.176 | 4.595 | 0.032 | 0.641 | NA |
| AAZ21068.1 | -14465.122 | -14462.171 | 5.902 | 0.015 | 0.302 | NA |
| AFS46782.1 | -14467.961 | -14464.484 | 6.954 | 0.008 | 0.167 | NA |
| AFS48343.1 | -14460.566 | -14425.923 | 31.802 | 0.000 | 0.000 | 4S; 5A; 9S; 35S; 38V; 70T; 83D; 84H; 85I; 91V; 94D; 95Q; 103L; 109P; 119T; 139T; 155E; 158K; 168N; 176N; 179F; 210L; 211R; 217G; 231S; 253A; 259P; 270Q; 274V; 277S; 292N; 298T; 305S; 311C; 321T |
| Ancestral branch to the DmdA clade | -28761.935 | -28758.081 | 7.7084 | 0.005 | 0.010 | 39Q |
| Ancestral branch to the non-DmdA clade | -28770.533 | -28766.874 | 7.3182 | 0.006 | 0.013 | - |

Branch identifiers follow the nomenclature of Supplementary Fig 19
Colors show same mutation in different lineages.
\*Amino acids refer to the first sequence in the alignment: AFS48343.1
<sup>1</sup>Log-likelihood score under the model under Null model
<sup>2</sup>Log-likelihood score under alternative model
<sup>3</sup>Likelihood ratio test
<sup>4</sup>Uncorrected p-value: raw- p-value without correction for multiple testing
<sup>5</sup>p-value corrected for multiple testing by Bonferroni

### Functional divergence during the molecular evolution of DmdA sequences

We tested whether DmdA and non-DmdA gene families were subjected to different functional constrains after gene duplication (Supplementary Fig 5). We estimated the one-ratio model (M0) that yielded a value ω = 0.053 (Table 6), indicating that purifying selection dominated the evolution of these proteins. The discrete model (M3) was applied to these sequences (Table 6) and the LRTs comparing M0 and M3 indicated significant variation in selective pressure among sites (Table 6; Supplementary Fig 23).

**Table 6.**
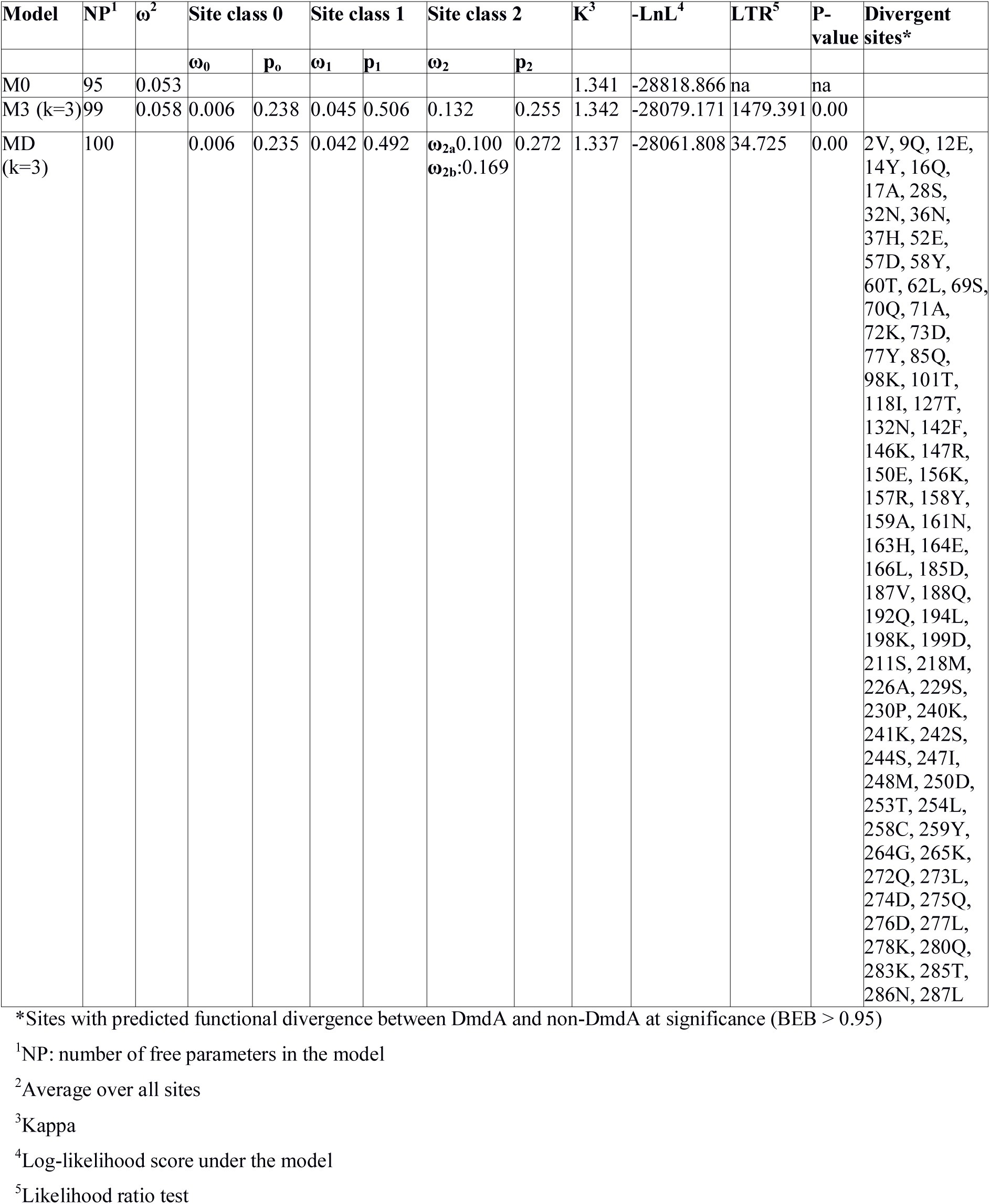
Parameter estimates of models evaluating functional divergence of DmdA and non-DmdA after gene duplication.

| Model | NP<sup>1</sup> | $\omega^2$ | Site class 0 | | Site class 1 | | Site class 2 | | K<sup>3</sup> | -LnL<sup>4</sup> | LTR<sup>5</sup> | P-value | Divergent sites* |
| --- | --- | --- | --- | --- | --- | --- | --- | --- | --- | --- | --- | --- | --- |
| | | | $\omega_0$ | $p_0$ | $\omega_1$ | $p_1$ | $\omega_2$ | $p_2$ | | | | | |
| M0 | 95 | 0.053 |  |  |  |  |  |  | 1.341 | -28818.866 | na | na |  |
| M3 (k=3) | 99 | 0.058 | 0.006 | 0.238 | 0.045 | 0.506 | 0.132 | 0.255 | 1.342 | -28079.171 | 1479.391 | 0.00 |  |
| MD (k=3) | 100 | | 0.006 | 0.235 | 0.042 | 0.492 | $\omega_{2a}$:0.100<br>$\omega_{2b}$:0.169 | 0.272 | 1.337 | -28061.808 | 34.725 | 0.00 | 2V, 9Q, 12E, 14Y, 16Q, 17A, 28S, 32N, 36N, 37H, 52E, 57D, 58Y, 60T, 62L, 69S, 70Q, 71A, 72K, 73D, 77Y, 85Q, 98K, 101T, 118I, 127T, 132N, 142F, 146K, 147R, 150E, 156K, 157R, 158Y, 159A, 161N, 163H, 164E, 166L, 185D, 187V, 188Q, 192Q, 194L, 198K, 199D, 211S, 218M, 226A, 229S, 230P, 240K, 241K, 242S, 244S, 247I, 248M, 250D, 253T, 254L, 258C, 259Y, 264G, 265K, 272Q, 273L, 274D, 275Q, 276D, 277L, 278K, 280Q, 283K, 285T, 286N, 287L |

\*Sites with predicted functional divergence between DmdA and non-DmdA at significance (BEB > 0.95)
<sup>1</sup>NP: number of free parameters in the model
<sup>2</sup>Average over all sites
<sup>3</sup>Kappa
<sup>4</sup>Log-likelihood score under the model
<sup>5</sup>Likelihood ratio test

The M3 model was compared with Model D, which accommodates both heterogeneity among sites and divergent selective pressures. The LRT was significant and supported the model D (Table 6), implying statistical evidence of functional divergence between DmdA and non-DmdA. Parameter estimates under Model D with k=3 site classes suggested that 23.6% of sites were evolving under strong purifying selection (ω = 0.006), while 26.7% of sites were evolving under much weaker selective pressure (ω = 0.04). Interestingly, a large set of sites (49.6%) were evolving under divergent selective pressures, with weaker purifying selection in the DmdA-clade (ω = 0.169) than non-DmdA-clade (ω = 0.100). We identified 77 sites evolving under divergent selective pressures between DmdA and non-DmdA (Table 6). Nineteen sites were located within the alpha helix (red tube in Supplementary Fig 24) of the secondary structure prediction and sixteen were located in the beta sheet (green arrows in Supplementary Fig 24). According to the global dN/dS estimates, for all divergent positions *dmdA* sequences seem to be more conserved than non-*dmdA* sequences. Moreover, this data is only compatible with recombination breaking linkage disequilibrium within the gene set that we observed with the HGT analysis.

Finally, we are interested in knowing if adaptive evolution has occurred in the lineages immediately following the main duplication event (Supplementary Fig 25). We applied two branch-site models to test for sites under selection on the ancestor associated with the DmdA and non-DmdA clades (Table 5). The LRT was significant for both ancestral branches (LRT > 7 and p-value < 0.05).

Nonetheless, the foreground ω for class 2 sites tended to infinite (ω=999) in both cases, indicating lack of synonymous substitutions (dS=0) in these sites. We also performed two-ratio models to estimate global ω on these branches, but both estimates tended to infinite (Supplementary Table 5), suggesting lack of synonymous substitution in the divergence of DmdA and non-DmdA ancestors. Therefore, although the fixation of only non-synonymous substitutions following gene duplication might indicate strong positive selection driving functional divergence of DmdA and non-DmdA families, we cannot confirm it with the applied tests.

## DISCUSSION

In this study we evaluated three scenarios for the evolutionary history of the DmdA gene family in marine bacteria. The results for each one are discussed separately.

### First scenario: a recent common ancestry between DmdA and GcvT

In relation to the first scenario, we found that contrary to our initial expectations, DmdA and GcvT have not a recent common ancestry, but they share an old common ancestor. However, the clear separation between DmdA and putative non-DmdA gene families that originated in the Archean ca. 2,400 Mya after a gene duplication, supports a common recent ancestry for DmdA and non-DmdA (Fig. 7; down and up). Our tertiary structure analyses indicate that they share a putative GcvT protein (EC 2.1.2.10) as their ancestor sequence. Indeed, our results agree with other studies in the case of DmdA (Reisch et al., 2008). Then, this clade seems to have originally been a GcvT (Fig. 7) as Bullock et al. (2017) suggested.

**Fig 7.**
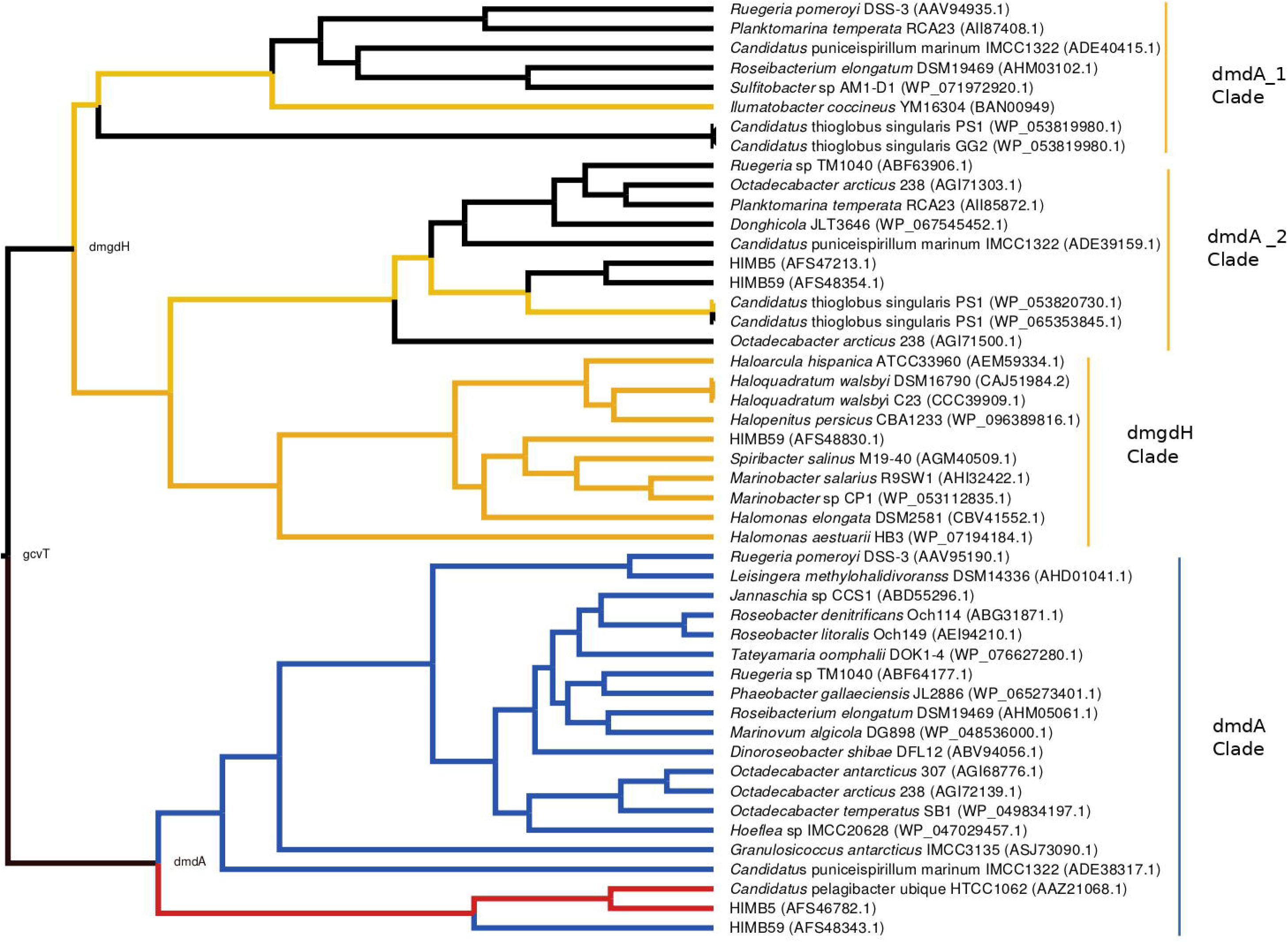
Hypothesis of DmdA evolution. BI phylogeny under uncorrelated relaxed clock model and Birth-death tree model. Node names represent the ancestral sequences reconstructed; GcvT prior to main duplication, DmdA for DmdA clade and DmgdH for non-DmdA clade. In DmdA clade, blue color represents ecoparalogs where pI is < 5.7 and they are adapted to less concentration of DMSP in comparison with DmdA paralogs (red color) which have pI => 6.5. In non-DmdA clade, yellow branches represents paralogs with DmgdH tertiary structure and black branches paralogs with DmdA tertiary structure.

The DmdA clade is a member of aminomethyltransferase (AMT/GCV_T) family with DMSP-dependent demethylase tertiary structure while non-DmdA clade includes an ancestor with a tertiary structure that better matches the dimethylglycine dehydrogenase oxidorreductase (DmgdH, EC 1.5.99.2) (Fig. 7) and members with DmdA tertiary structure. To establish structural convergence as the reason of this DmdA structure coincidence between DmdA and non-DmdA members, we used a phylogenetic approach based on reconstructing ancestral sequences of the two clades, and then to model the ancestral proteins. We determined different structural features between ancestral sequence reconstructed from DmdA and non-DmdA families. In the first case, the ancestral sequence reconstructed coincides with a DmdA tertiary structure, as well as with a DmdA sequence with physico-chemical properties inferred in this study and agree with previous ones (Reisch et al., 2008). However, the non-DmdA ancestral sequence reconstructed is a DmgdH that seems to be kept in the clade called DmgdH (Fig. 7: yellow color) as well as in some members of DmdA clades (within non-DmdA clade) where the majority of sequence gained DmdA structure (Fig. 7). Therefore, DmdA structural features seem to have emerged independently in both clades: DmdA and non-DmdA. This finding is extremely interesting, since known cases of structural convergence of proteins are rare (Zakon, 2002). Experimental assays expressing and screening the activity of the ancestral proteins at different conditions will be required to corroborate the structural convergence.

Since GcvT does not share the most recent common ancestry with DmdA, we examined the functional divergence between DmdA and non-DmdA clades to explain how natural selection could have driven the divergence of the DmdA gene family. We found 77 codon sites evolving under divergent selective pressures between DmdA and non-DmdA gene families. Structural divergence seemed to be imposed on the protein during sequence divergence, since nineteen sites were located within the alpha helix of 2D structure and sixteen in the beta sheet. Nonetheless, essential regions of the enzymes as active sites seem to be under strong purifying selection, suggesting preservation of the ancestral function. The observation that DmdA sequences have less conserved divergent sites than non-DmdA sequences, suggests that non-DmdA conserves the ancestral function, whereas DmdA evolved to acquire new functions in different environments, probably as a response to the Huronia ice ball Earth (Zhang, 2003).

### Second scenario: coevolution between *Roseobacter* and DMSP-producing-phytoplankton

In the second scenario, our data does not support the hypothesis of a co-evolution sceneario between *Roseobacter* and DMSP-producing-phytoplankton (Luo et al., 2013). On the contrary, we found an ancestor sequence of DmdA cluster similar to DmdA from a strain of *Ca.* P. ubique that diverged after a more recent duplication event, before the dinoflagellate radiation in the late Permian. This finding indicates that the enzyme activity has not changed in the course of DmdA evolution. Indeed, we found that most of the codons in DmdA clade are under purifying selection probably due to the importance of this pathway for sulfur acquisition. Nonetheless, we also detected episodic positive selection in four sequences affecting a few sites, suggesting that adaptive evolution fine-tuned the function of DmdA in *Roseobacter*. Furthermore, positively selected residues were located around the GcvT domain and close to the residue involved in conserved interaction with THF, reinforcing the idea of adaptive evolution in response to the external environment.

During the study of this scenario, we suspected that *dmdA* was acquired by HGT in *Roseobacter* and SAR11. This agrees with Luo et al., (2013) and Tang et al. (2010) which found that the expansion of *dmdA* was by HGT. Moreover, our study evidence that DmdA ancestral sequence in our phylogeny comes from a marine heterotrophic bacteria adapted to presence of DMSP in the Archean, after a HGT event from this bacteria to another linage that acquired the *dmdA* ancestral sequence. However, after the HGT events, some *dmdA* sequences have acquired similar residue changes by independent (parallel) evolution, reinforcing the idea of functional/ecological constrains. Therefore, *Rhodobacteraceae* can live in an environment where DMSP is the main source of sulfur because they acquired the DmdA ancestor sequence by HGT, prior to have been exposed to the environment in which this protein proved useful, as Luo & Moran (2014) suggested. We did not find any signal of positive selection in *Roseobacter* group, but in contrast we found episodic evolution between SAR11 sequences. Yet, as we already mentioned DMSP is part of an ancient pathway in *Alphaproteobacteria* (Bullock et al., 2017) and it could explain the ancient origin of DmdA.

On the other hand, *Roseobacter* paralogs analyzed in this study were functionally annotated as DmdA function (González et al., 2019), as they perform the same function as the original gene (DmdA ancestor). However, we found differences in predicted isoelectric point values (pI), which were inferred in this study. Then, these paralogs could be considered as ecoparalogs as Sánchez-Pérez et al (2008) proposed for their study. Isoelectric point of a protein provides an indication of its acidic nature (Oren et al., 2005) and in this case, differences in pI suggest that the proteins differ in halophilicity. We observed proteins with the highest pI values in the DmdA ancestor sequence, as well as *Ca.* P ubique sequence and this last one has a pI similar to the first (DmdA ancestor) (Fig. 7). Therefore, we deduced that DmdA ancestor was adapted to a higher concentration of salinity, which could have modulated the selection of the DMSP enzymatic degradation routes as in bacteria such as the model organism *R. pomeroyi* DSS-3 (Salgado et al., 2014). Interestingly, *R. pomeroyi* degradates more DMSP by the demethylation pathway under high salinity conditions, and then produces a high amount of MeSH (Howard et al., 2008; Magalhães et al., 2012; Salgado et al., 2014).

Given our data, we propose that the ancestor of the pathway that evolved in the Archean, was exposed to a higher concentration of DMSP in a sulfur rich atmosphere and in an anoxic ocean, compared to recent ecoparalogs which should adapt to lower concentration of DMSP (Fig 7). Indeed, the ancestral ecoparalog from which recent ecoparalogs derived (*Ca* Puniceispirilum marinum IMCC1322 or ADE38317.1 and the *Roseobacter* clade) could have undergone episodes of adaptation (the branch showed positive selection in branch-models) which would explain the change in protein stability (Pál et al., 2006). As consequence, the protein could have experimented slight reductions or loss of function.

### Third scenario: pre-adapted enzymes to DMSP prior to Roseobacter origin

In this evolutionary scenario, *Roseobacter* clade was pre-adapted to the conditions created by eukaryotic phytoplankton at the late Permian, including dinoflagellates that released vast amounts of DMSP (Bullock et al., 2017; Luo & Moran, 2014). Our analyses indicate that the *Roseobacter* ancestor has already adapted to a high DMSP before *Roseobacter* clade arose (Luo et al., 2013). Therefore, we support Reisch et al. (2011 a,b) hypothesis where DMSP demethylation pathway enzymes are adapted versions of enzymes that were already in bacterial genomes, and evolved in response to the availability of DMSP. Since the first step in DMSP demethylation is a reaction catalyzed by DMSP demethylase encoded by *dmdA* gene (Dickschat et al., 2015), DMSP adaptation could have been evolved in this gene that originated in the Archean, a time where several lineages of bacteria produced DMSP as an osmolyte or antioxidant in the presence of the early cyanobacteria, or as a cryoprotectant in the Huronian glaciation. In bacteria, a methyltransferase gene, *dysB*, is up-regulated during increased salinity, nitrogen limitation, and at low temperatures (Curson et al., 2017), conditions already predicted to stimulate DMSP production in phytoplankton and algae (Bullock, et al., 2017; Ito, et al., 2011). Afterward, those roles may have helped to drive the fine adaptation of existing enzymes for DMSP metabolism, and those adaptations came handy in the late Precambrian glaciations that allowed the radiation of algae and animals.

## CONCLUSION

In conclusion, we found that *Roseobacter* adaptation to DMSP occurred via functional diversification after duplication events of the DmdA gene and adaptations to environmental variations via ecoparalogs of intermediate divergence. Our findings suggest that salinity could have been a trigger for the adaptation to DMSP metabolism.

## AUTHOR CONTRIBUTIONS

LH conceived the study, performed the phylogenetic, molecular and protein structure analysis and wrote the paper. LH and AV performed the selection analysis. LH, LE, VS and AV interpreted findings. All authors contributed to the design of the study, manuscript revision, read and approval of the submitted version.

## ACKNOWLEDGMENTS

We would like to thank to Dr. Romain Studer from BenevolentAI for his critical role in the 3D visualization of protein and mapping sites onto the 3D structure and Dr. Buckley Iglesias from Universidad Autonóma de Madrid for his introduction to molecular dating analysis with BEAST 2. This research was supported by grant CTM2016-80095-C2 from the Spanish Ministry of Economy and Competitiveness.

## Conflict of Interest Statement

The authors declare that the research was conducted in the absence of any commercial or financial relationships that could be construed as a potential conflict of interest.

## SUPPORTING INFORMATION

### FIGURES#

**Supplementary Fig 1.** Time tree of *Alphaproteobacteria* evolution with geologic timescale. Solid circles mark nodes that map directly to the NCBI Taxonomy and the open circles indicate nodes that were created during the polytomy resolution process which is described in Hedges et al. (2015).

**Supplementary Fig 2.** Time tree of *Gammaproteobacteria* evolution with geologic timescale. Solid circles mark nodes that map directly to the NCBI Taxonomy and the open circles indicate nodes that were created during the polytomy resolution process which is described in Hedges et al. (2015).

**Supplementary Fig 3.** Time tree of *Halobacteriales* evolution with geologic timescale. Solid circles mark nodes that map directly to the NCBI Taxonomy and the open circles indicate nodes that were created during the polytomy resolution process which is described in Hedges et al. (2015).

**Supplementary Fig 4.** GcvT phylogenetic tree based on 20 DmdA ortholog protein sequences and 184 DmdA homologs using RaxML. DmdA sequences are indicated with red color and closer homologs for those with blue color. Tip labels include a maximum e-value < e-50.

**Supplementary Fig 5.** Phylogenetic trees of DmdA based on 20 DmdA ortholog protein sequences and 28 DmdA homologs using RaxML (A), Phylobayes (B), Phylip (C) and Beast (D). DmdA sequences are indicated with blue color and the first DmdA proteins identified with read color (AAV95190.1: *Ruegeria pomeroyi* DSS-3, AAZ21068.1: *Ca.* P. ubique HTCC1062). Tip labels include a maximum e-value < e-80.

**Supplementary Fig 6.** Proxy for the species tree constructed by BI and using RPS16 sequences from 35 genomes here analyzed for inferring evolutionary history of DmdA.

**Supplementary Fig 7.** DmdA tree using the common set of taxa used for the topology tests. Tree was constructed by ML for topology tests and BI for an easily visualization of phylogenetic relationships in unrooted trees.

**Supplementary Fig 8.** Proxy for the species tree using the common set of taxa used for the topology tests. Proxy was constructed by ML for topology tests and BI for an easily visualization of phylogenetic relationships in unrooted trees.

**Supplementary Fig 9.** Proxy for the species tree using the common set of taxa used for the topology tests. The blue branches denote HGT events and red arrows the direction.

**Supplementary Fig 10a.** Phylogenetic tree of DmdA based on 20 DmdA ortholog protein sequences and 28 DmdA homologs using BEAST2. Bayesian posterior probabilities (PP) is shown to establish the support for the clades. Red color denote DmdA clade, orange color indicate non-DmdA clade and yellow color DmgdH clade.

**Supplementary Fig 10b.** Multiple sequence alignment with blue color represents the highest level of conservation (100%) when the alignment is divided in the same four clades found in the Supplementary Fig 10a and Fig 4.

**Supplementary Fig 11.** Clustering sequences based on principal component analysis from Jalview v2.10. The sequences are projected along three vectors giving a 3-dimensional view of how the sequences cluster. Components are generated by an eigenvector decomposition of the matrix formed from the sum of substitution matrix scores at each aligned position between each pair of sequences – computed with blosum62 matrix. Grey color denotes sequences with putative dmgdH structure.

**Supplementary Fig 12.** DmdA phylogenetic tree with the ancestor labeling included. Internal nodes labels were inferred using FastML. N1is the oldest ancestor and from N2 to N18 are children.

**Supplementary Fig 13.** Psi-blast results for sequences similar to the DmdA ancestral protein inferred with FastML.

**Supplementary Fig 14.** Non-DmdA phylogenetic tree with the ancestor labeling included. Internal nodes labels were inferred using FastML. N1 is the oldest ancestor and from N2 to N18 are children.

**Supplementary Fig 15.** Phylogenetic tree of DmdA based on 20 DmdA ortholog protein sequences and 28 DmdA homologs with the ancestor labeling included. Internal nodes labels were inferred using FastML. N1 is the oldest ancestor and from N2 to N18 are children.

**Supplementary Fig 16.** Phylogenetic trees of *dmdA* based on 20 *dmdA* ortholog gene sequences using PhyML. A non-parametric bootstrap is shown to establish the support for the clades. Tip labels show red color for the first *dmdA* gene identified (AAV95190.1: *R. pomeroyi* DSS-3, AAZ21068.1: *Ca.* P. ubique HTCC1062).

**Supplementary Fig 17.** Posterior probabilities for dN/dS categories under the M1a model. Blue bars depict the category with the dN/dS = 1 and grey bars the category with dN/dS << 1. Sites that are grey denote codons under strong purifying selection.

**Supplementary Fig 18.** Phylogeny for *dmdA* sequences. Blue color indicates the branches from group B which are compared with the rest of branches (group A) under two-ratio models.

**Supplementary Fig 19.** Phylogeny for *dmdA* sequences constructed by ML from DNA alignment in frame. Red branches have a dN/dS value > 1. Red numbers indicate the branches. “ω” represents a dN/dS value where non-synonymous mutations are higher than synonymous mutations. Four sequences (WP_047029467, AHM05061,1, ABV94056,1, AFS48343,1) presented a significant LRT after correcting for multiple testing (green color).

**Supplementary Fig 20.** Foreground-branches tested for branch-site selection models. Red color indicates the branches of interest (foreground branches). We performed 20 tests, where only one of the branches pointed by red color was considered at a time; all other branches are corresponding to background-branches.

**Supplementary Fig 21.** Multiple sequence alignment of DmdA orthologs. Blue colors represent sites with the highest level of conservation (100%). Red squares represents sites under positive selection. The posterior probability of each site was calculated by BEB. Green asterisk indicate residues that have a conserved interaction with THF (Schuller et al. 2012).

**Supplementary Fig 22.** Parallel mutational changes detected in specific genes from different lineages. Red color identifies parallel mutational changes on specific branches of the *dmdA* phylogeny. The shared sites are under positive selection. Branch identifiers follow the nomenclature of Supplementary Fig 21.

**Supplementary Fig 23.** Posterior probabilities for dN/dS categories under the M3 model. Red and blue bars depict the categories with the highest dN/dS (values for each category are provide in the key). Sites that are mostly grey denote codons under strong purifying selection, whereas those predominantly red show codons under light purifying selection.

**Supplementary Fig 24.** Multiple sequence alignment of DmdA orthologs and DmdA homologs showing conserved regions (blue color) and codon sites evolving under divergent selective pressures (red colored columns). The secondary structure prediction using Jpred4 via Jalview is also shows for the alignment.

**Supplementary Fig 25.** Phylogeny for *dmdA* ortholog and *dmdA* homolog sequences. Ancestral branches to the DmdA clade and to non-DmdA clades, with red and blue colors respectively, are considered as foreground-branches in different branch-site selection models.

### TABLES

**Supplementary Table 1.** Data collected from MarRef database include information about sequences and genomes used in this study, taxonomy and sampling environment.

**Supplementary Table 2.** Tree comparison by TOPD/FMTS. Two randomization methods estimate that the similarity between two trees produced by BI or ML is better than random. This random analysis is repeated 100 times and the result is the mean and SD of the different repetitions.

|  | Split Distance MM |  | Split Distance random |  |
| --- | --- | --- | --- | --- |
|  | Mean | SD | Mean | SD |
| Beast vs beast | 0 | 0 | 0.9988 | 0.002 |
| Beast vs RaxML | 0.1990 | 0 | 0.9988 | 0.002 |

**Supplementary Table 3.**
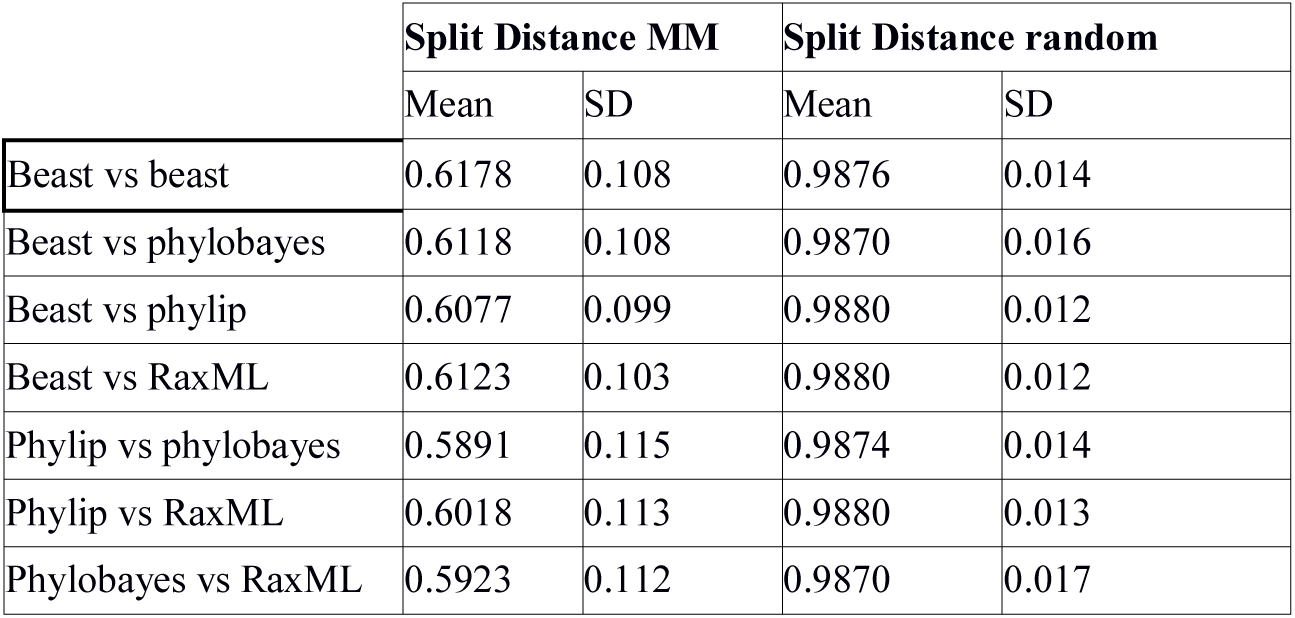
Tree comparison by TOPD/FMTS. Two randomization methods estimate that the similarity between two trees produced by BI or ML is better than random. This random analysis is repeated 100 times and the result is the mean and SD of the different repetitions.

**Supplementary Table 4.** Physico-chemical properties on predecessor and DmdA ortholog sequences inferred through Compute ProtParam tool from Expasy – SIB Bioinformatics Resource Portal.

| Taxonomy | Identification | PI<sup>1</sup> | Mw<sup>2</sup> | Instability index<sup>3</sup> |  | Aliphatic index<sup>4</sup> | Location (PSORTb v.3.0) |
| --- | --- | --- | --- | --- | --- | --- | --- |
| ASR<sup>5</sup> | Root marginal sequences of DmdA family | **6.5\*** | 41334.4 | 39.5 | stable | 91.32 | Cytoplasmic |
| *Ca. P. ubique* HTCC1062 | AAZ21068.1 | **6.47\*** | 41831.81 | 32.73 | stable | 86.1 | Cytoplasmic |
| HIMB59 | AFS48343.1 | 5.17 | 41499.43 | 39.62 | stable | 86.1 | Cytoplasmic |
| HIMB5 | AFS46782.1 | **6.99\*** | 41692.14 | 39.23 | stable | 91.9 | Cytoplasmic |
| *G. antarcticus* IMCC3135 | ASJ73090.1 | 4.91 | 43371.33 | 33.58 | stable | 92.19 | Cytoplasmic |
| *Ca. puniceispirillum marinum* IMCC1322 | ADE38317.1 | 5.55 | 41421.73 | **43.47** | **unstable** | 92.21 | Cytoplasmic |
| *L. methylohalidivorans* DSM14336 | AHD01041.1 | 4.93 | 40057.61 | 39.14 | stable | 86.57 | Cytoplasmic |
| *R. pomeroyi* DSS-3 | AAV95190.1 | 5.27 | 39895.45 | 37.59 | stable | 84.4 | Cytoplasmic |
| *Hoeflea* sp IMCC20628 | WP_047029467.1 | 4.98 | 40736.4 | 35.67 | stable | 87.91 | Cytoplasmic |
| *D. shibae* DFL12 | ABV94056.1 | 4.81 | 41294.27 | 39.05 | stable | 90.03 | Cytoplasmic |
| *O. temperatus* SB1 | WP_049834197.1 | 5.03 | 40693.65 | 27.46 | stable | 85.68 | - |
| *O. antarcticus* 307 | AGI68776.1 | 5.32 | 40692.51 | 26.43 | stable | 86.75 | Cytoplasmic |
| *O. arcticus* 238 | AGI72139.1 | 5.5 | 40570.47 | 28.03 | stable | 87.32 | Cytoplasmic |
| *R. elongatum* DSM19469 | AHM05061.1 | 5.49 | 40459.36 | 40.05 | unstable | 88.15 | Cytoplasmic |
| *M. algicola* DG898 | WP_048536000.1 | 5.12 | 40770.54 | 35.43 | stable | 85.17 | Cytoplasmic |
| *Jannaschia* sp CCS1 | ABD55296.1 | 5.05 | 40852.7 | **42.04** | **unstable** | 86.57 | Cytoplasmic |
| *P. gallaeciensis* JL2886 | WP_065273401.1 | 5.58 | 41133.97 | **40.89** | **unstable** | 82.07 | Cytoplasmic |
| *Ruegeria* sp TM1040 | ABF64177.1 | 5.78 | 42951.08 | 38.16 | stable | 82.26 | Cytoplasmic |
| *T. oomphalii* DOK1-4 | WP_076627280.1 | 5.24 | 41152 | **41.96** | **unstable** | 84.71 | Cytoplasmic |
| *R. denitrificans* Och114 | ABG31871.1 | 5.15 | 40785.38 | 30.1 | stable | 86.62 | Cytoplasmic |
| *R. litoralis* Och149 | AEI94210.1 | 5.09 | 40648.28 | 27.93 | stable | 87.71 | Cytoplasmic |
| ASR | Root marginal sequence of DmdA and non-DmdA families | 4.43 | 31948.03 | **45.46** | **unstable** | 89.16 | Cytoplasmic |
| ASR | Root marginal sequence of non-DmdA family | 4.3 | 40334.85 | **43.72** | **unstable** | 84.44 | Cytoplasmic |
| N10 | Non-DmdA tree | 4.69 | 39908.25 | **41.21** | **unstable** | 92.38 |  |
| N3 | DmdA tree | 4.85 | 41479.21 | **40.25** | **unstable** | 86.56 |  |

<sup>1</sup>Theoretical isoelectric point
<sup>2</sup>Theoretical molecular weight
<sup>3</sup>A protein whose instability index is smaller than 40 is predicted as stable, a value above 40 predicts that the protein may be unstable
<sup>4</sup>It is the relative volume occupied by aliphatic side chains (valine, isoleucine, alanine and leucine)
<sup>5</sup>Ancestral sequence by reconstruction
\*Highest isoelectric point values

**Supplementary Table 5.** Parameters of branch-models.

| Model | $\omega 1$ | $\omega 2$ | $-\ln L^4$ | LRT<sup>5</sup> | P-value |
| --- | --- | --- | --- | --- | --- |
| One $\omega$ (one-ratio) | 0.05348 | NA | -31199.102911 | NA | NA |
| Two $\omega$ (two-ratio)<sup>1</sup> | 0.05367 | 999 | -31197.315923 | 3.573976 | 0.0587 |
| Two $\omega$ (two-ratio)<sup>2</sup> | 0.05399 | 0.00 | -31197.838823 | 2.528176 | 0.1118 |
| Two $\omega$ (two-ratio)<sup>3</sup> | 0.05362 | 999 | -31197.199937 | 0.000012 | 0.9972 |

<sup>1</sup>Two $\omega$ , one for the ancestral DmdA gene and another for the rest of genes.
<sup>2</sup>Two $\omega$ , one for the ancestral non-DmdA gene and another for the rest of genes.
<sup>3</sup>Two $\omega$ , one for the two ancestral genes (DmdA and non-DmdA) and another for the rest of genes
<sup>4</sup>Log-likelihood score under the model
<sup>5</sup>Likelihood ratio test

## ADDITIONAL INFORMATION

**Supplementary Data 1.** Details of structural information collected by I-TASSER for each sequence used on the evolutionary study of DmdA gene family (Fig. 2).

