## Supplementary figures and images for "Evolutionary history of dimethylsulfoniopropionate (DMSP) demethylation enzyme DmdA in marine bacteria"

### supplementary figure 1

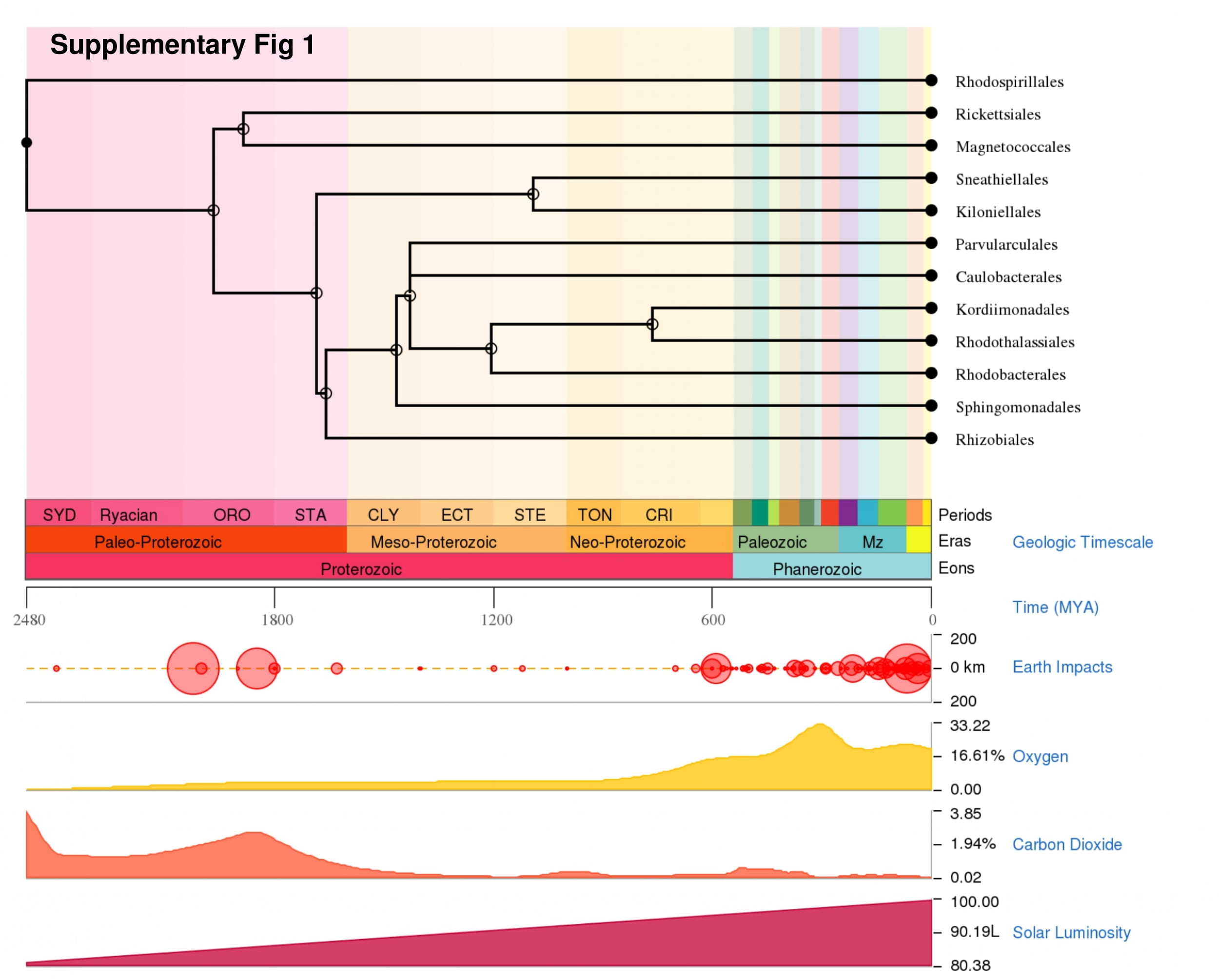

### supplementary figure 2

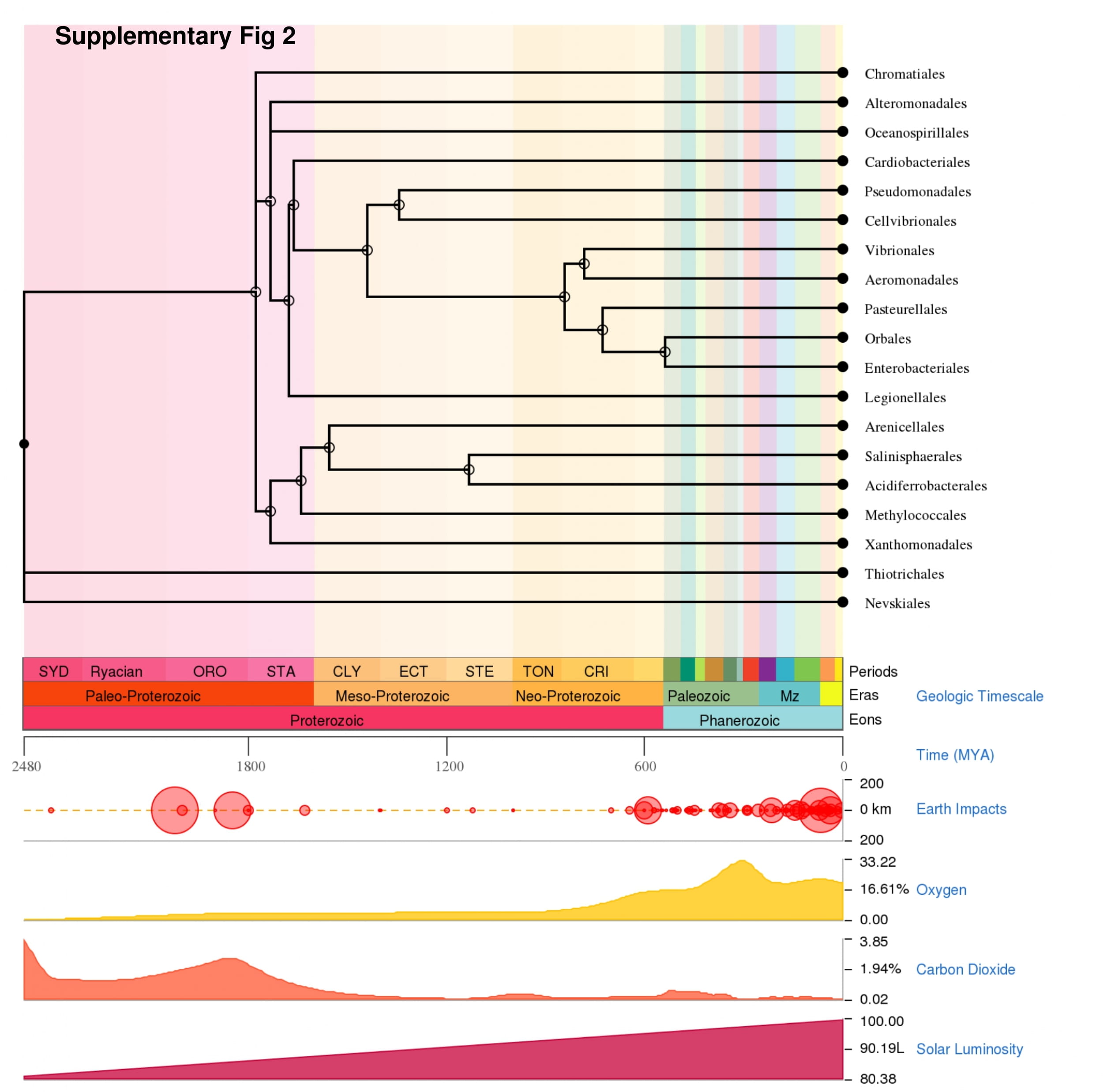

### supplementary figure 3

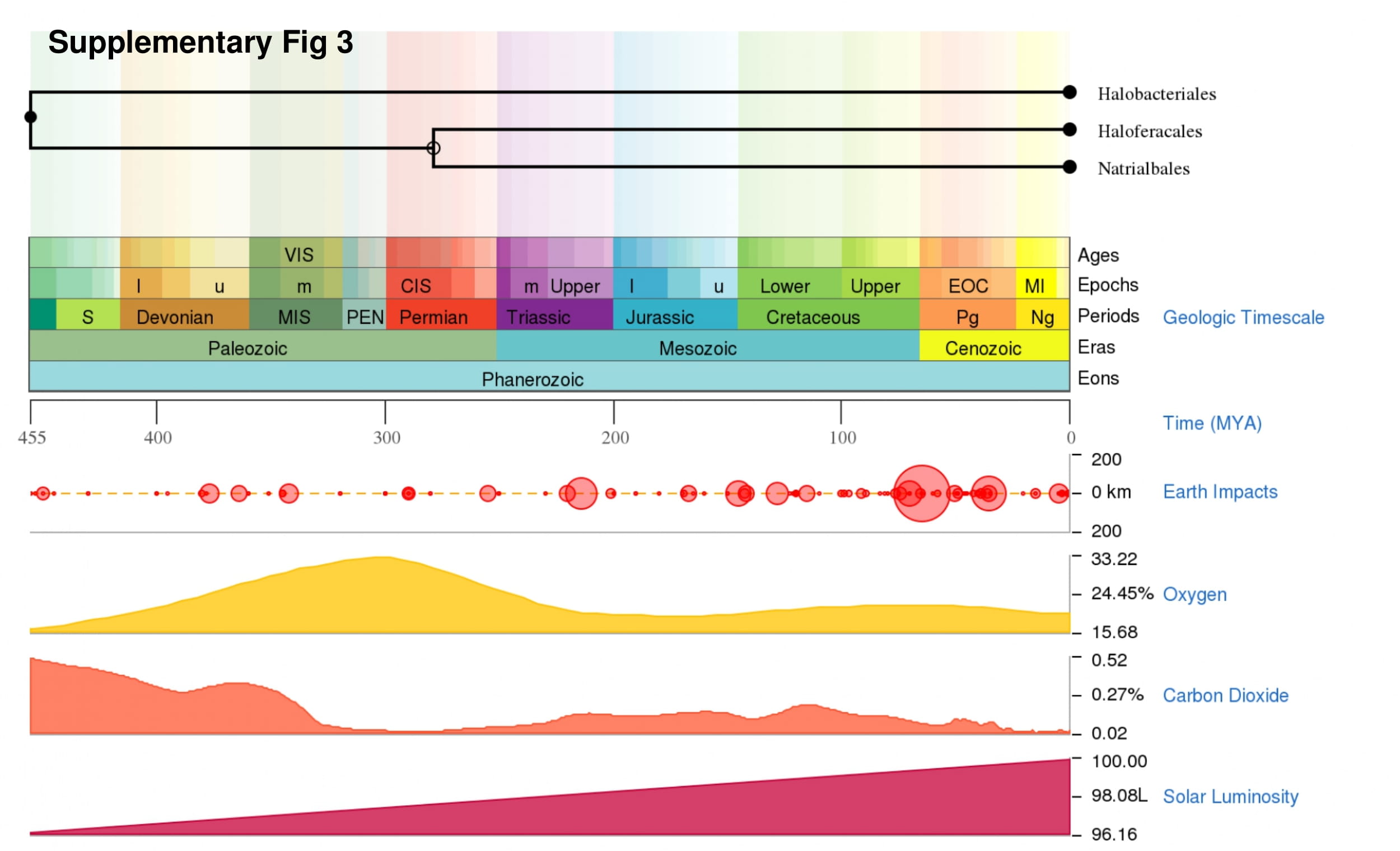

### supplementary figure 4

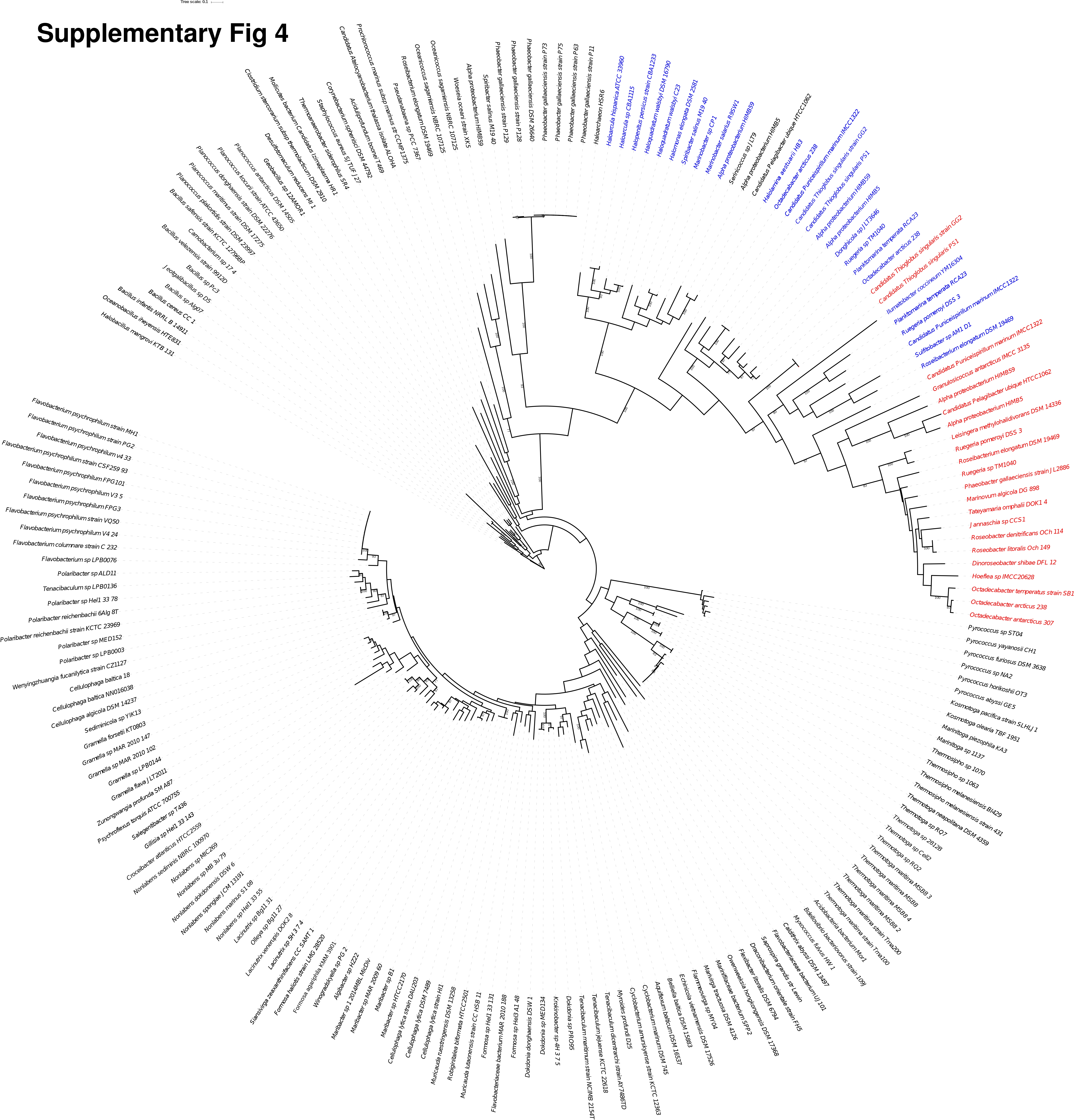

### supplementary figure 5

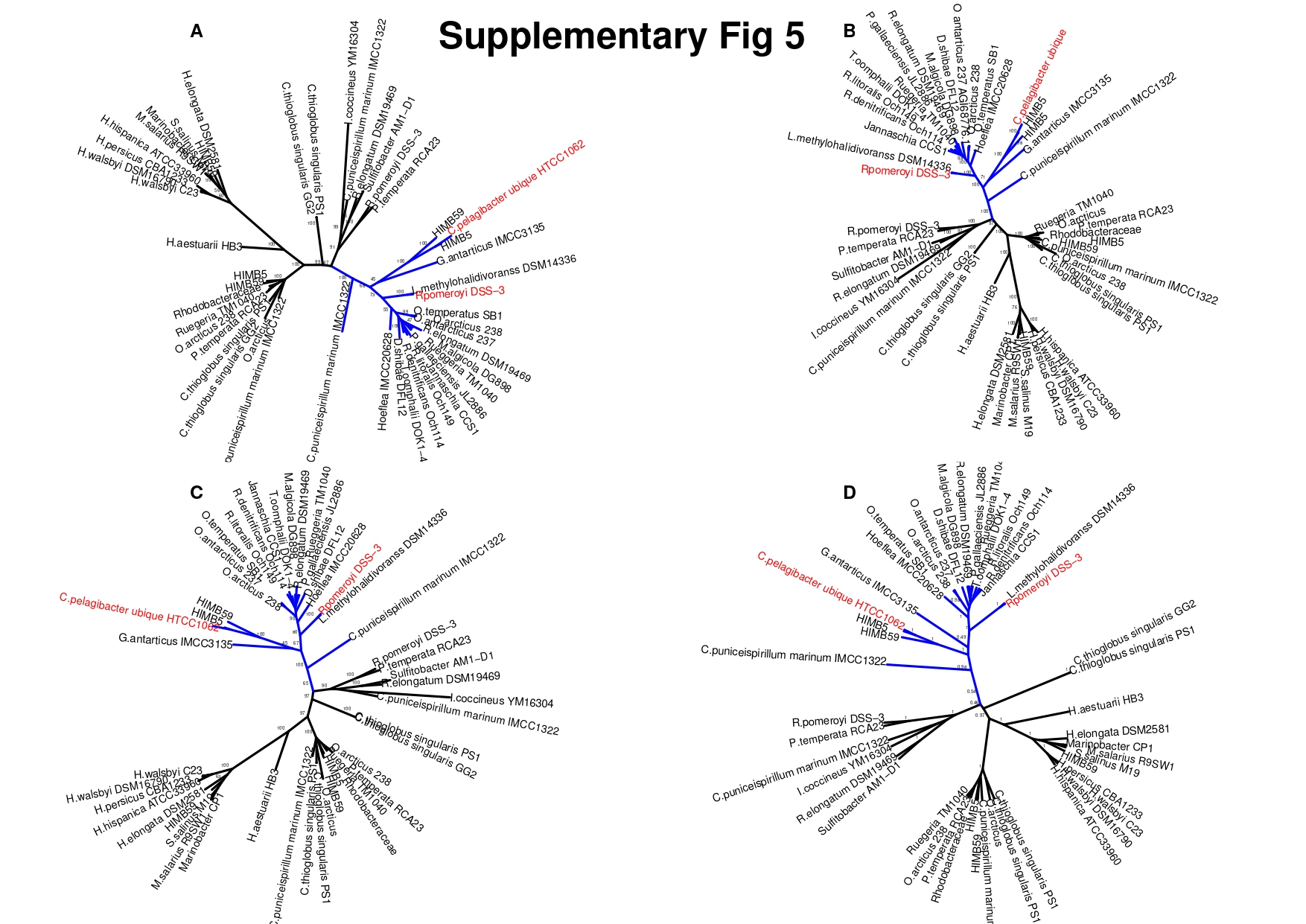

### supplementary figure 6

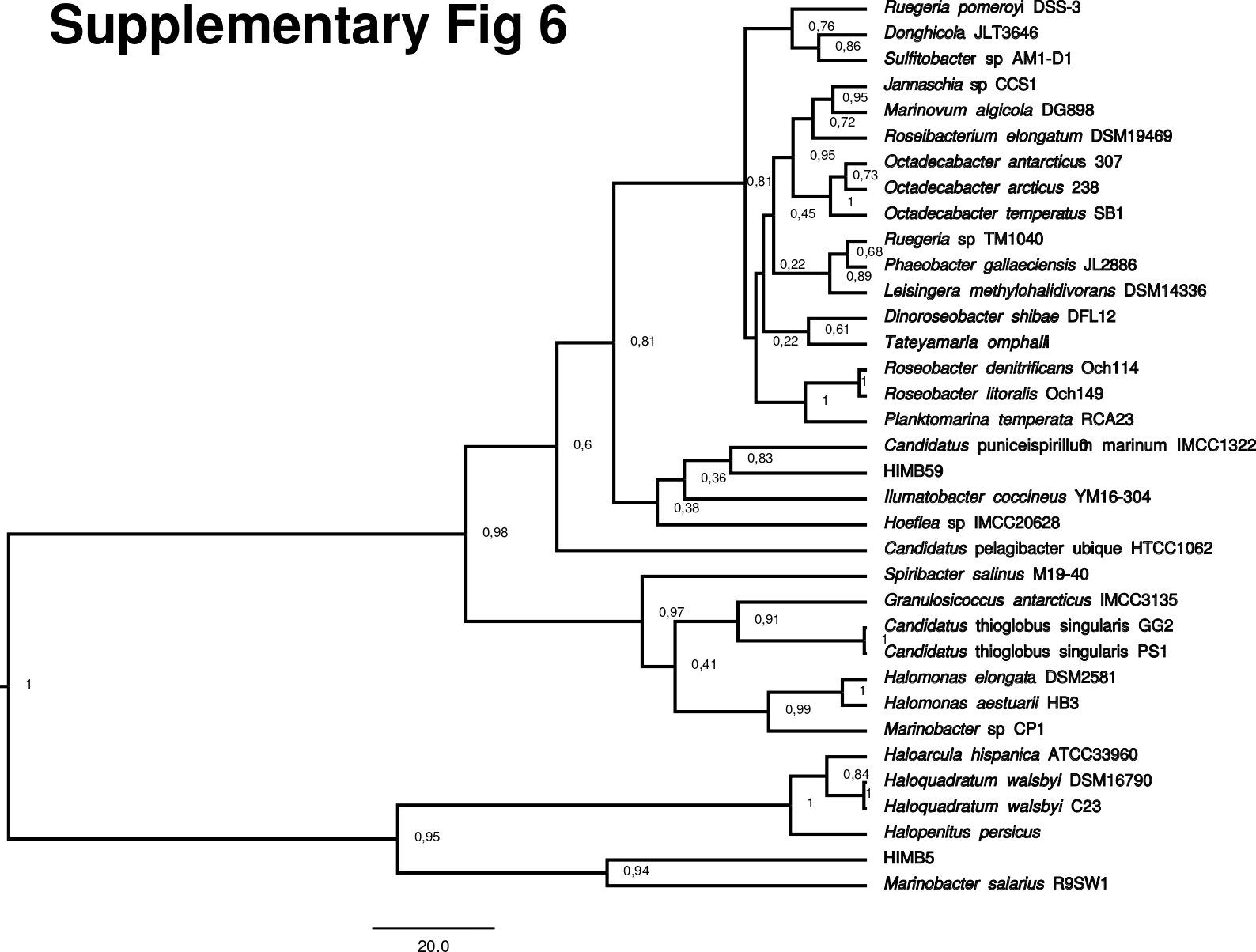

### supplementary figure 7

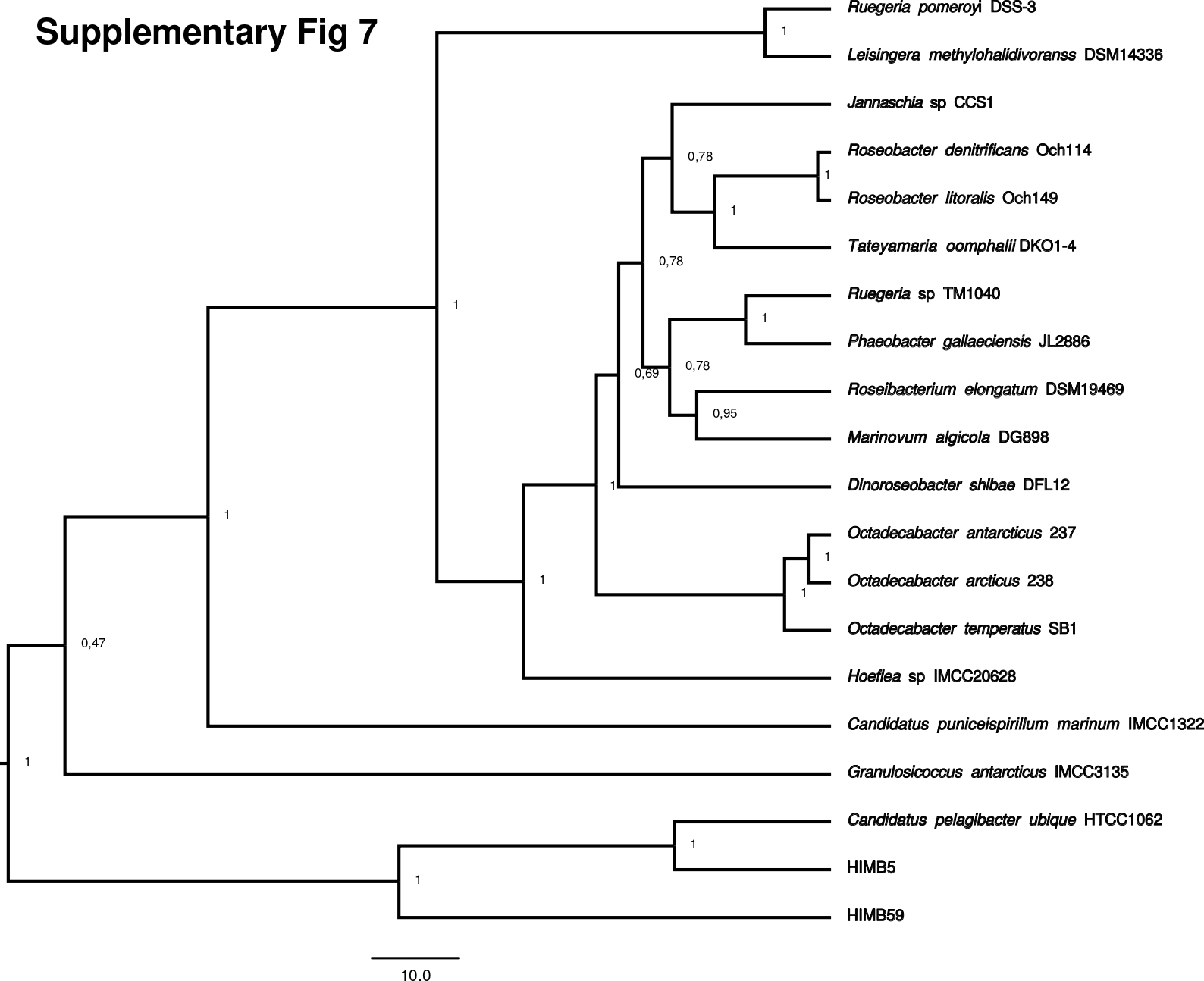

### supplementary figure 8

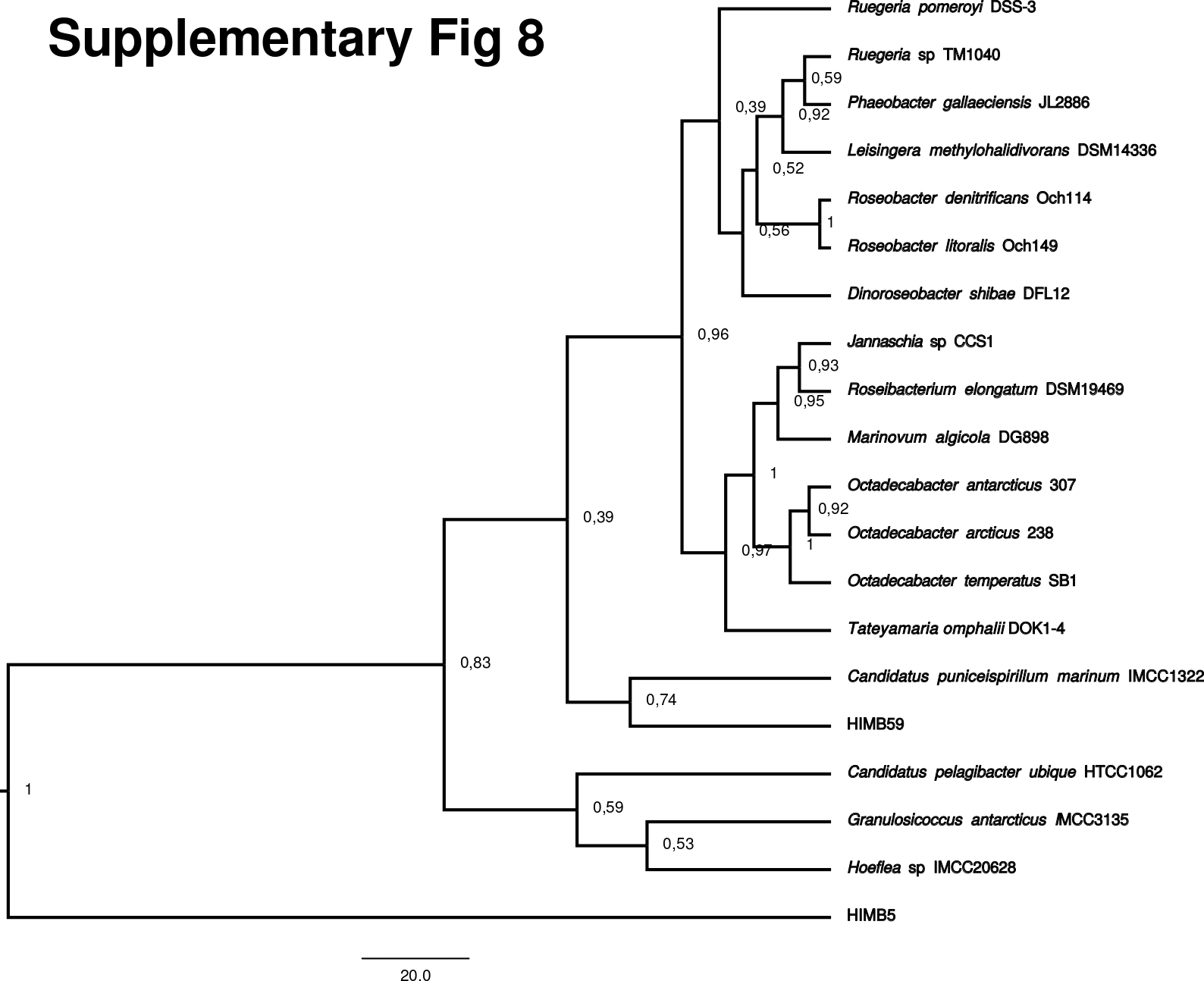

### supplementary figure 9

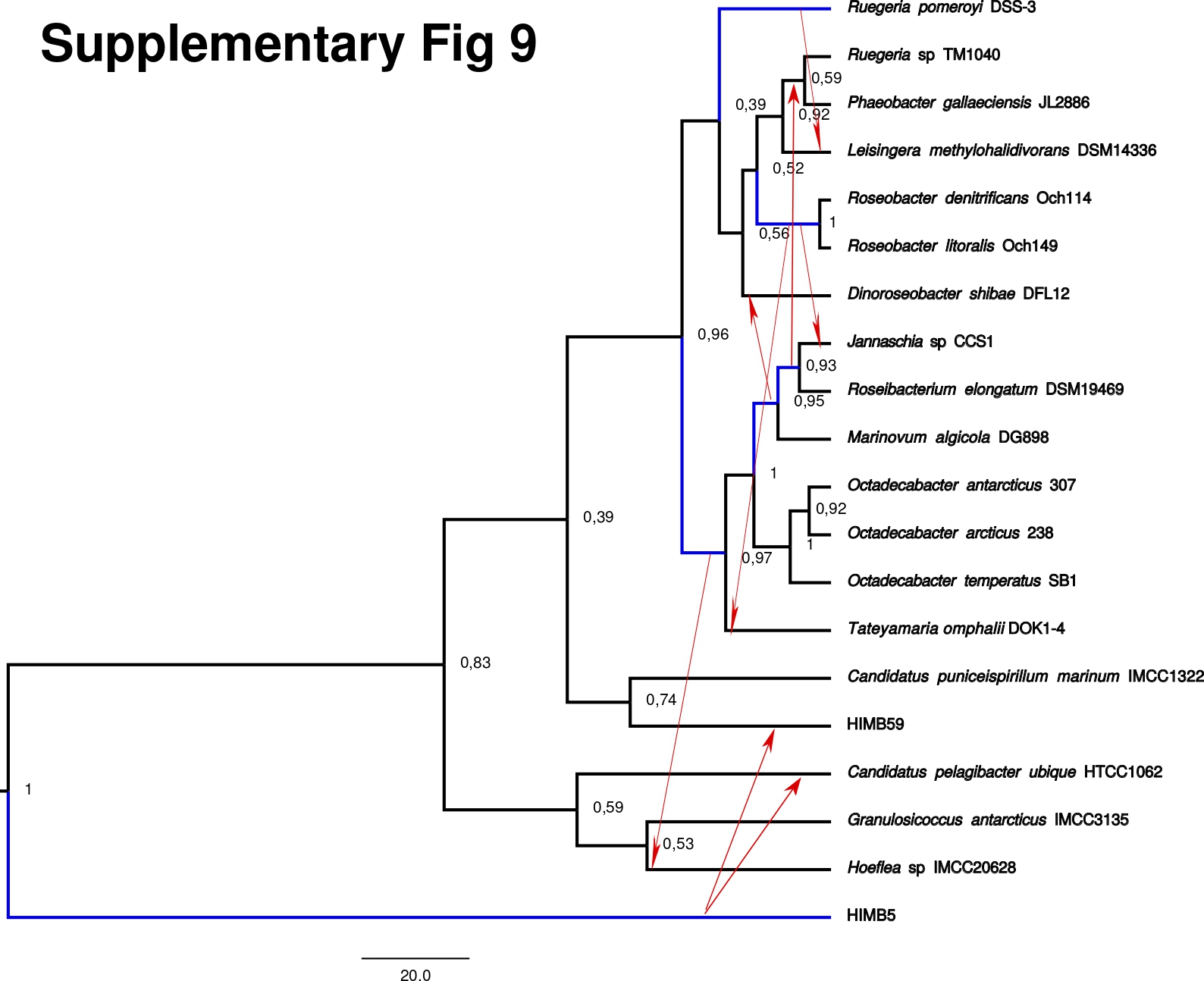

### supplementary figure 10a

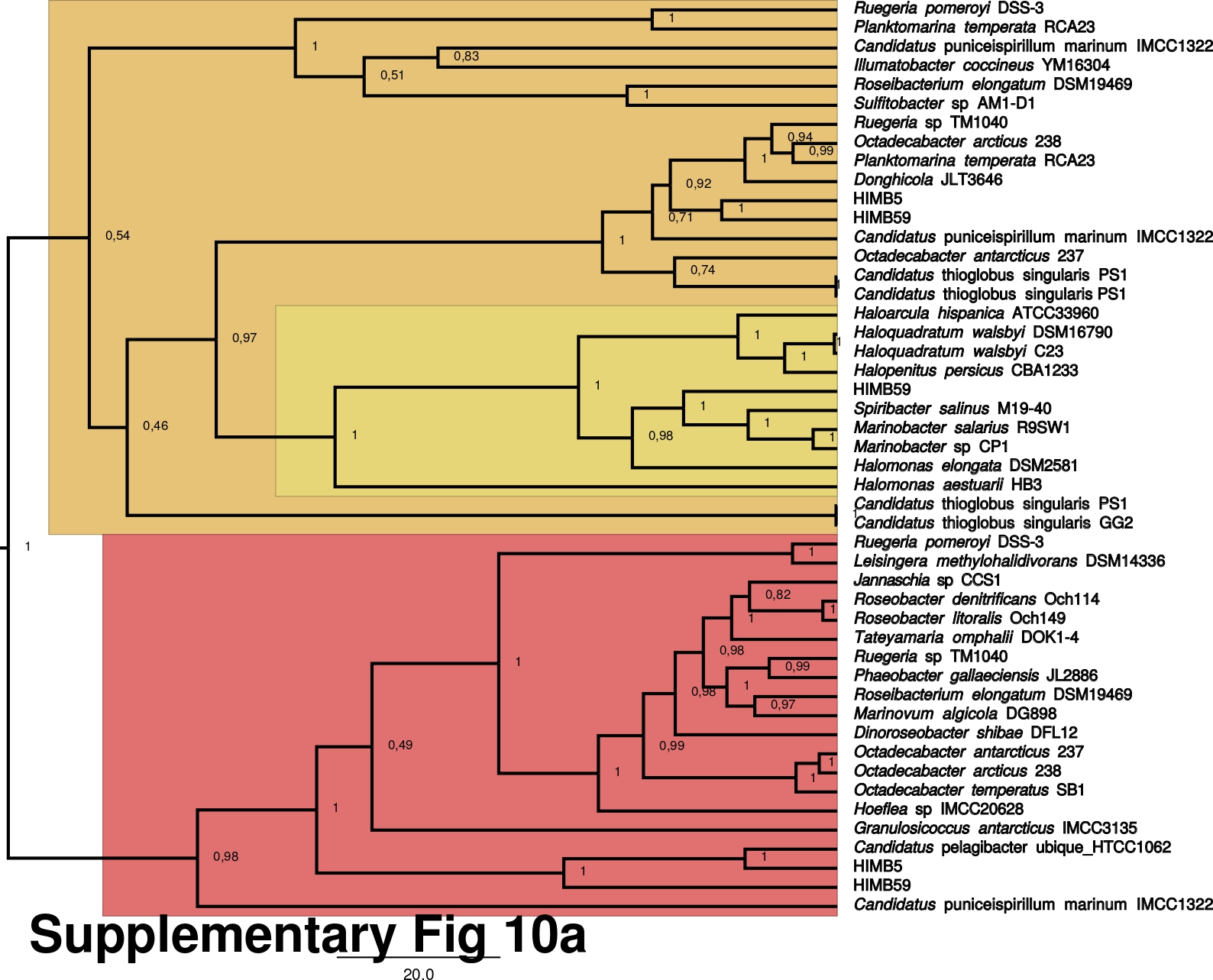

### supplementary figure 10b

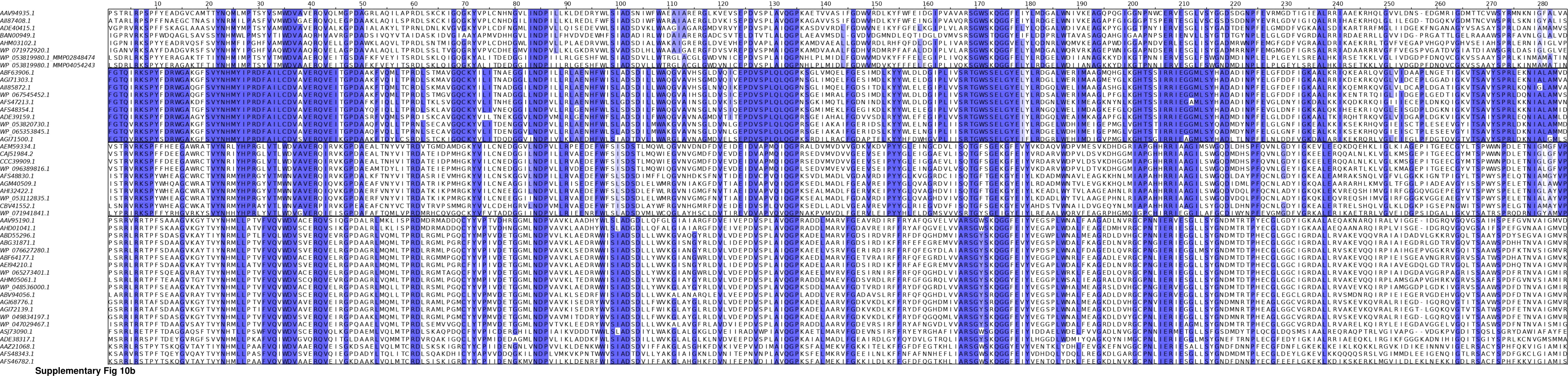

### supplementary figure 11

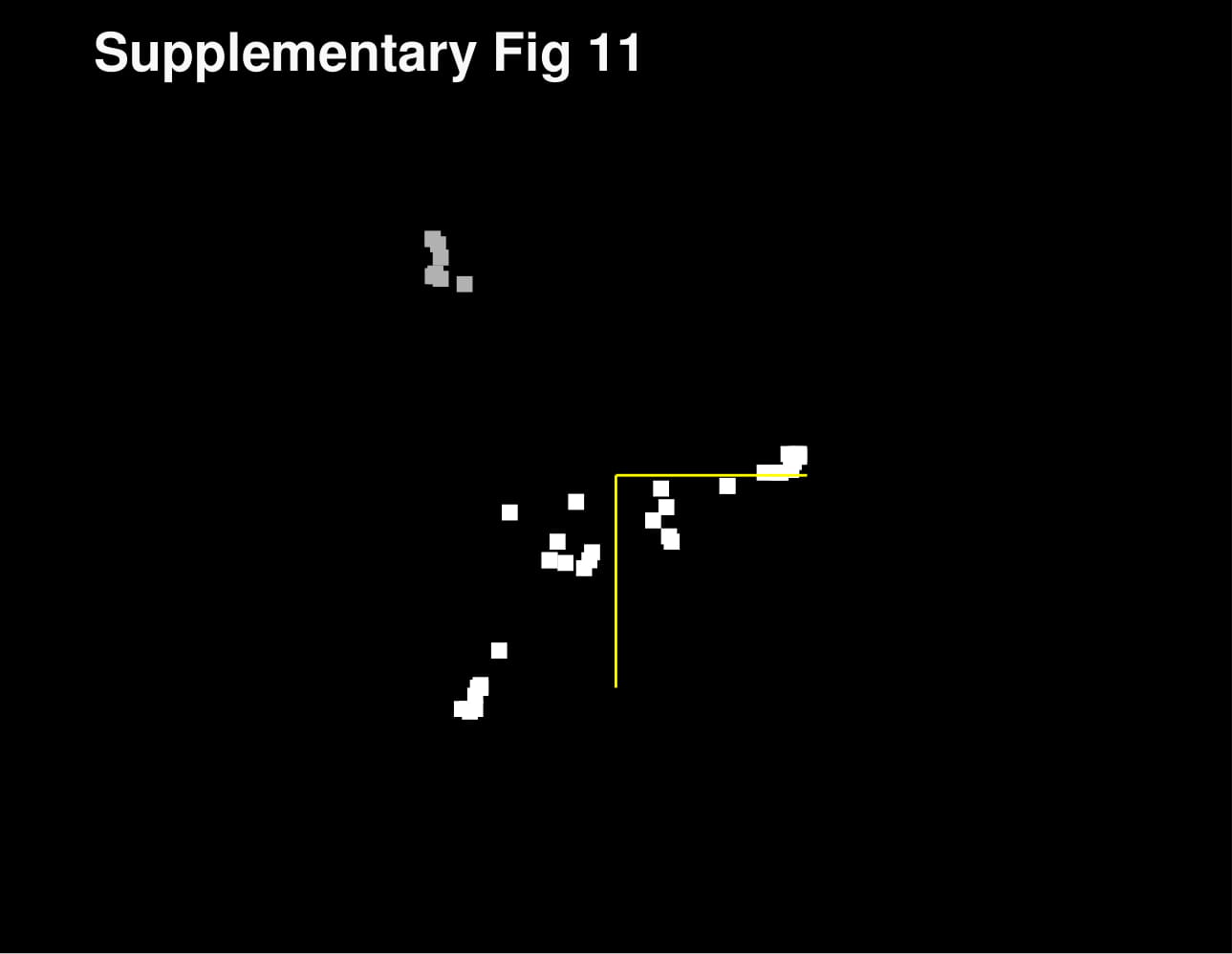

### supplementary figure 12

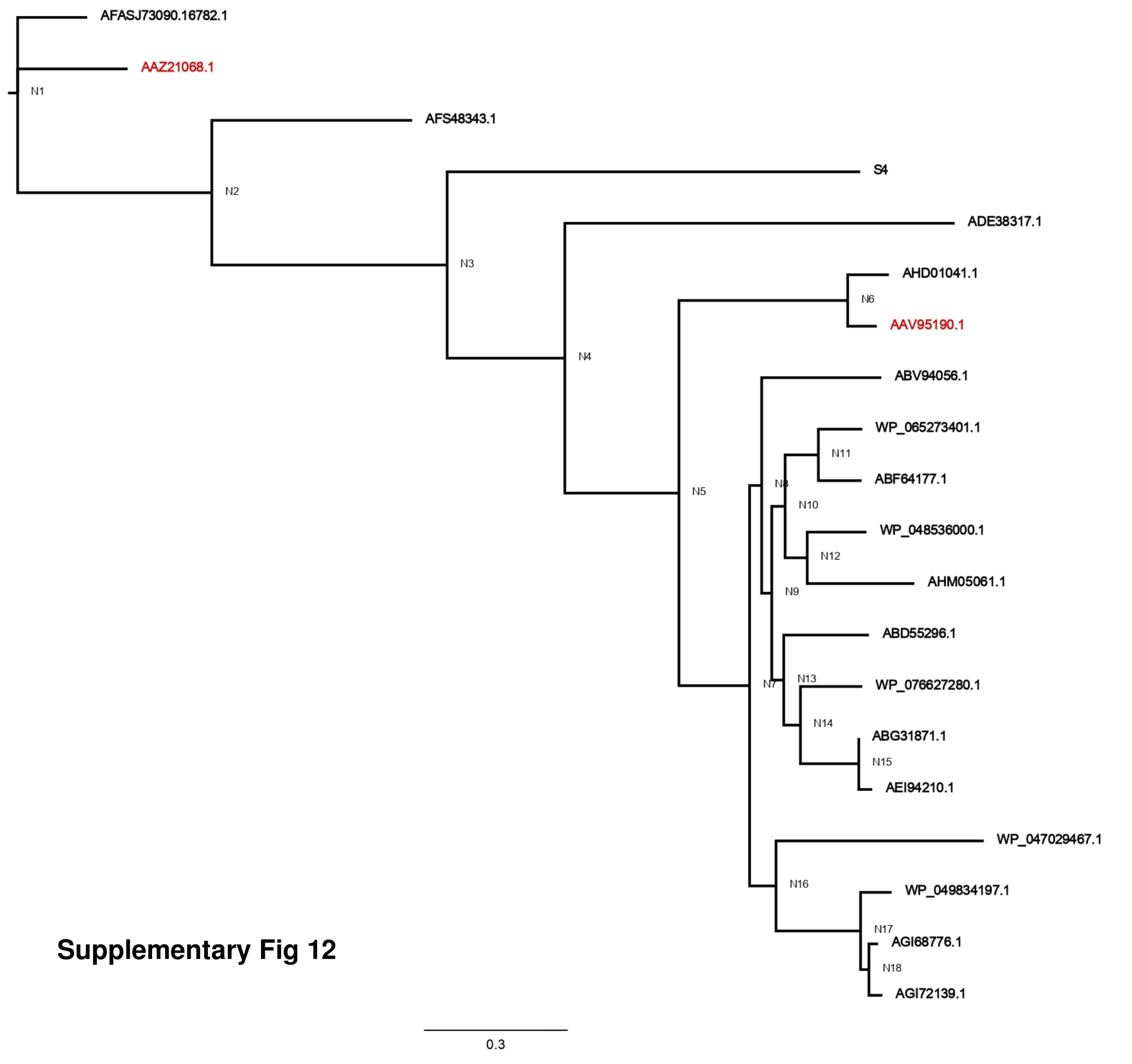

### supplementary figure 13

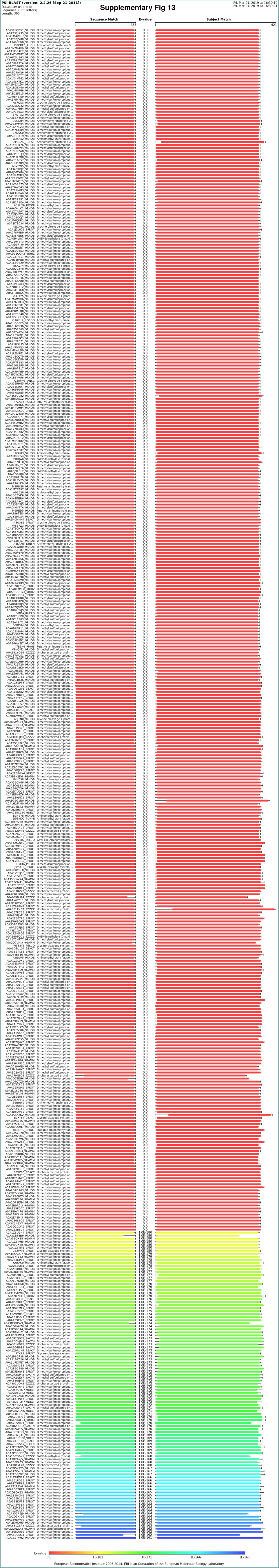

### supplementary figure 14

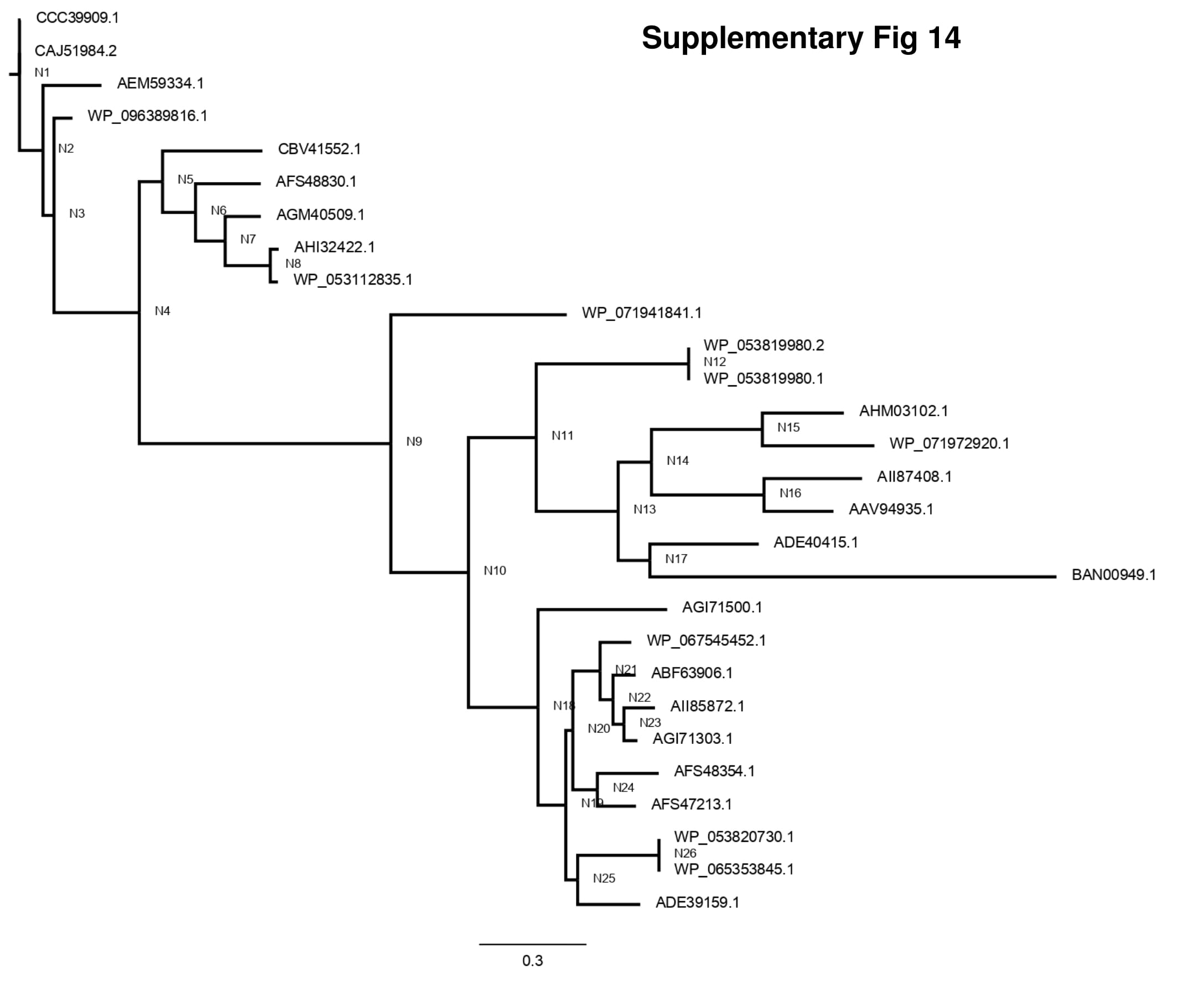

### supplementary figure 15

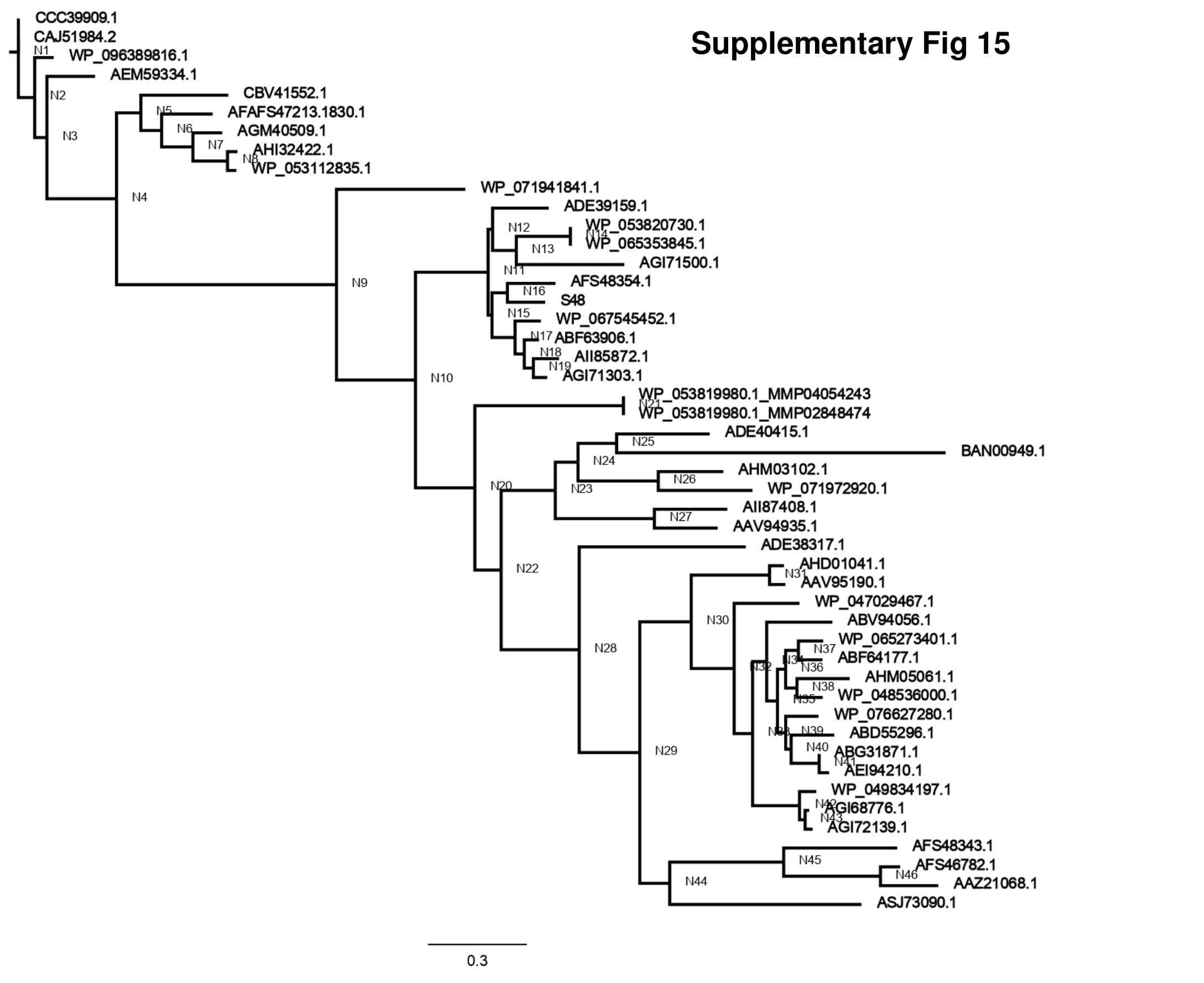

### supplementary figure 16

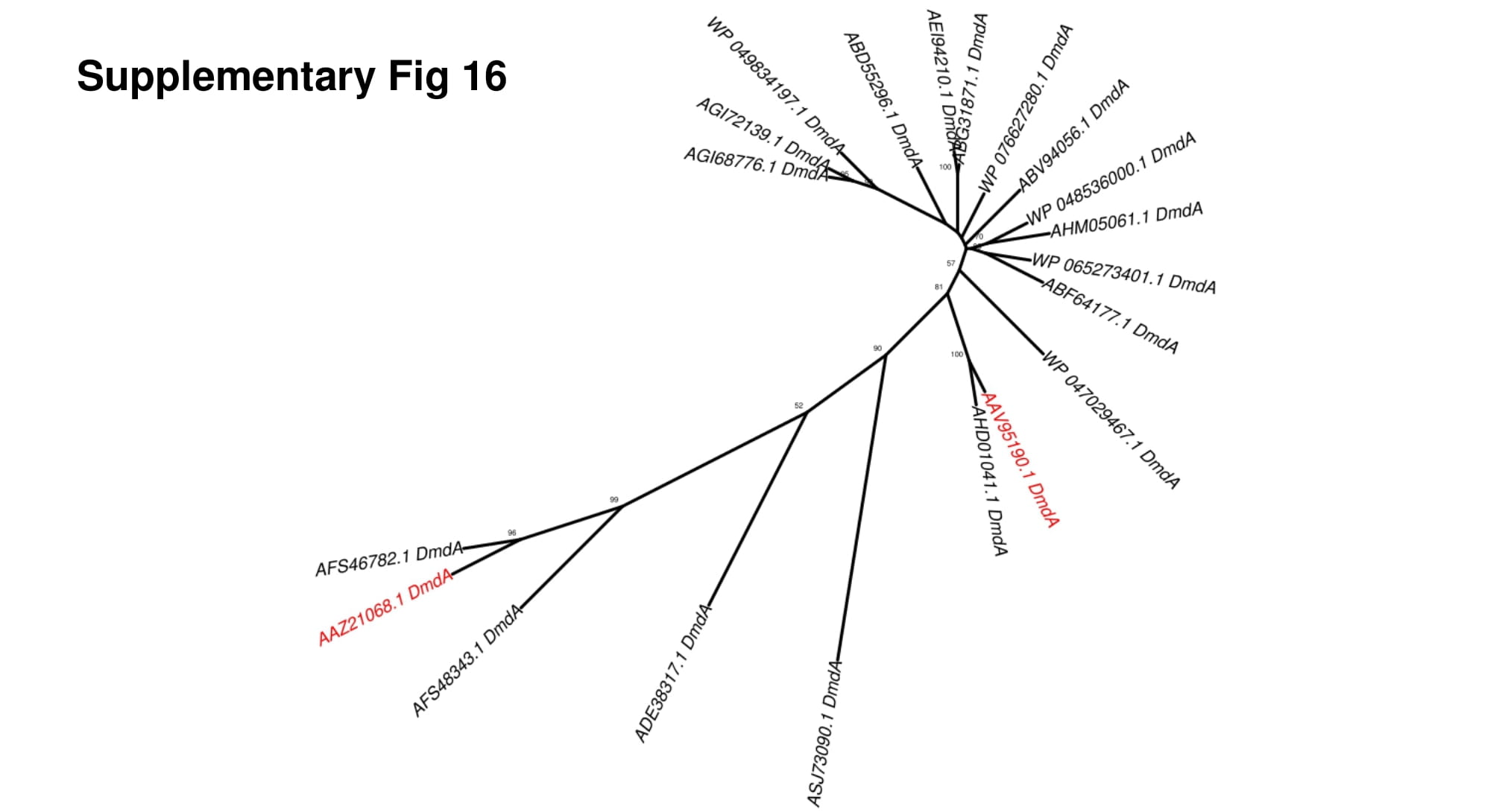

### supplementary figure 17

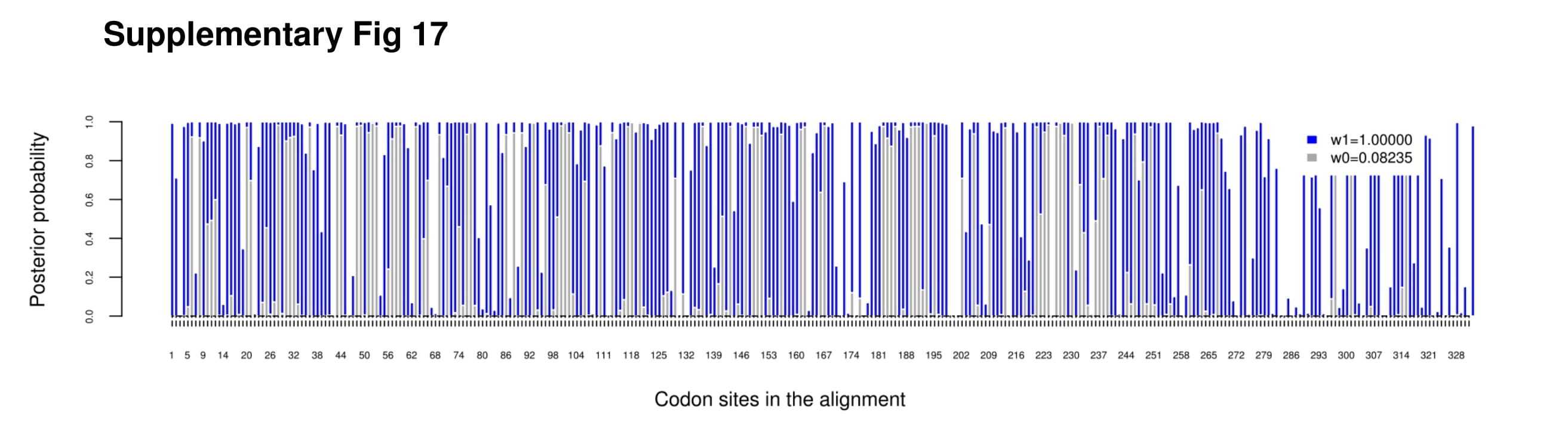

### supplementary figure 18

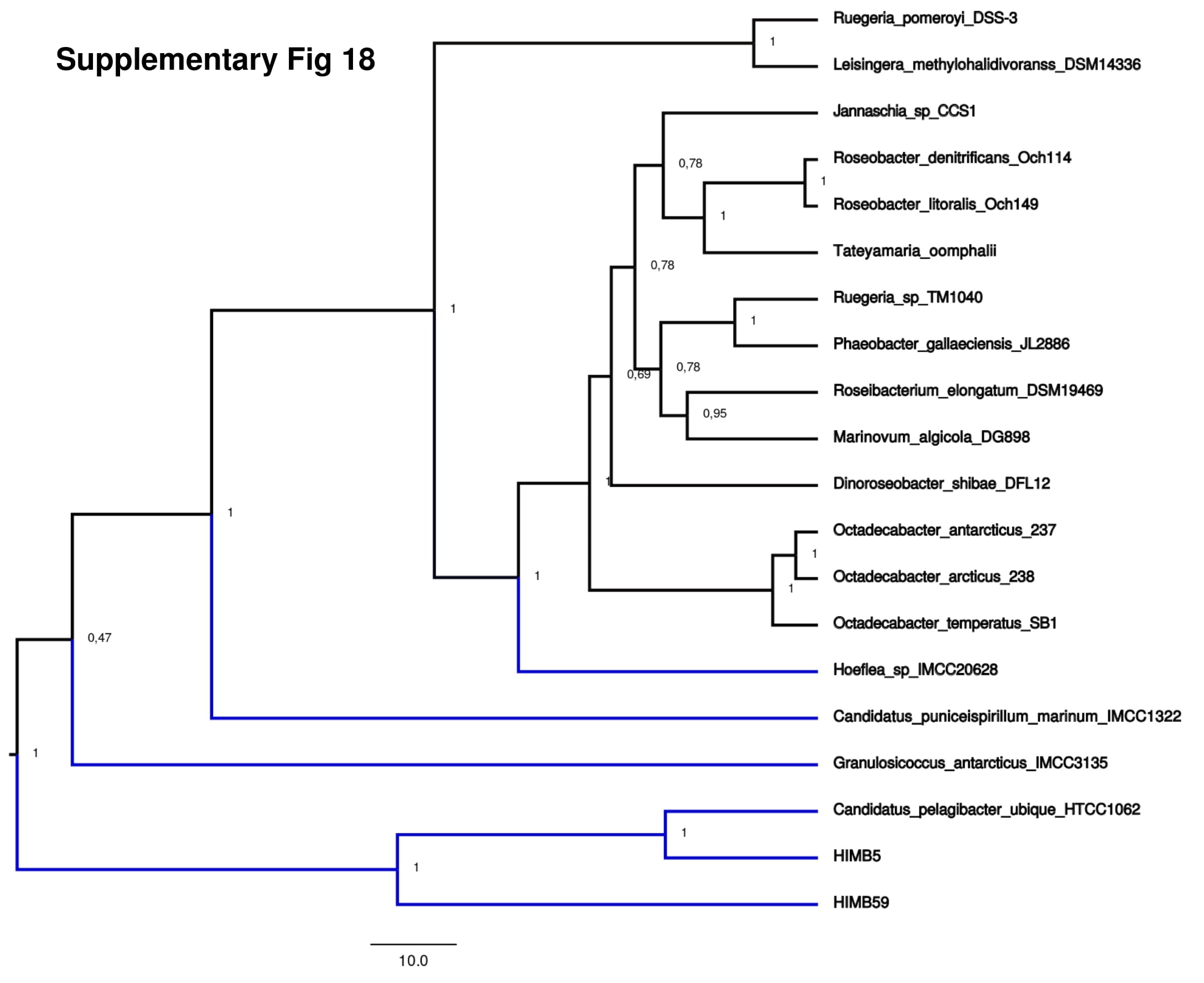

### supplementary figure 19

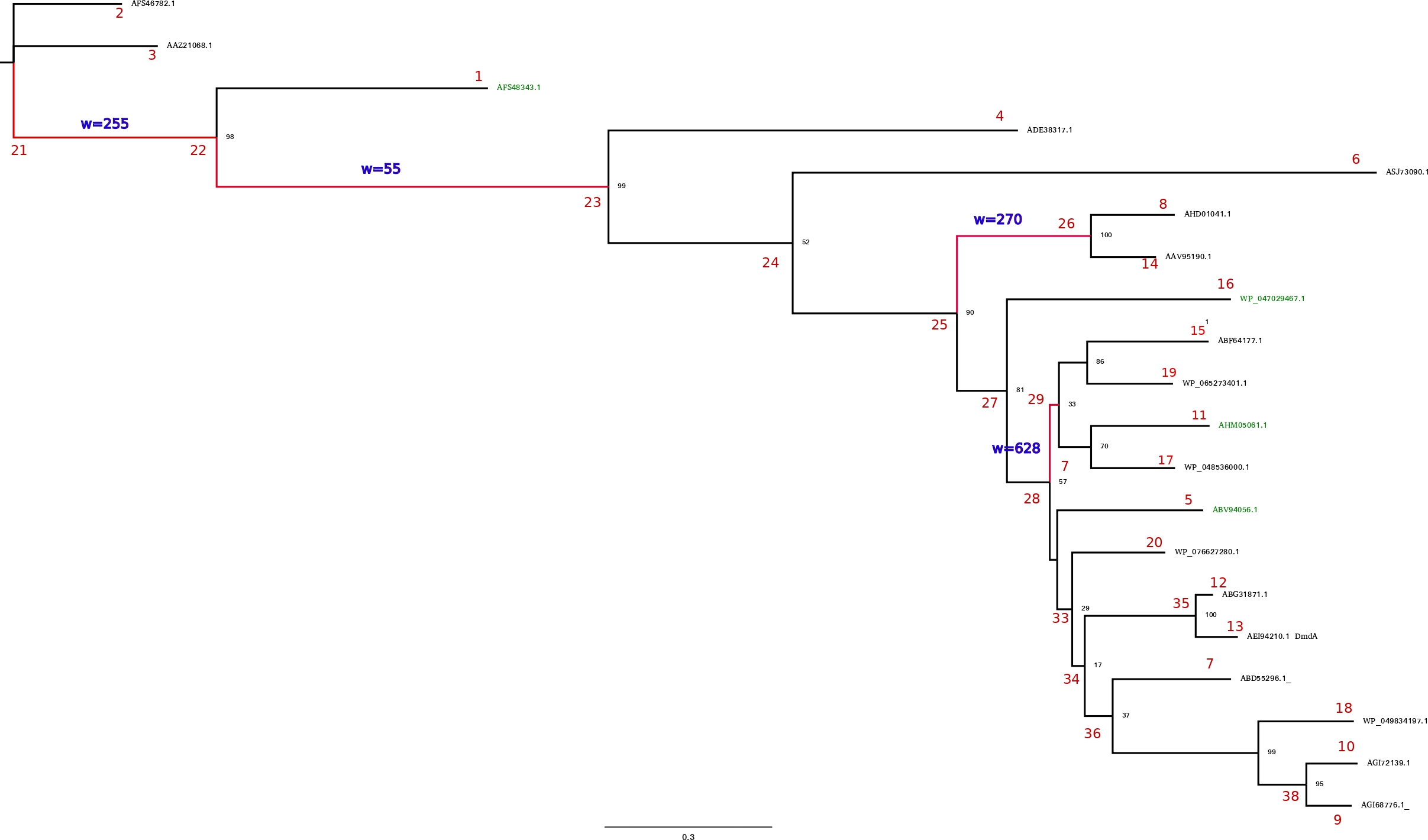

### supplementary figure 20

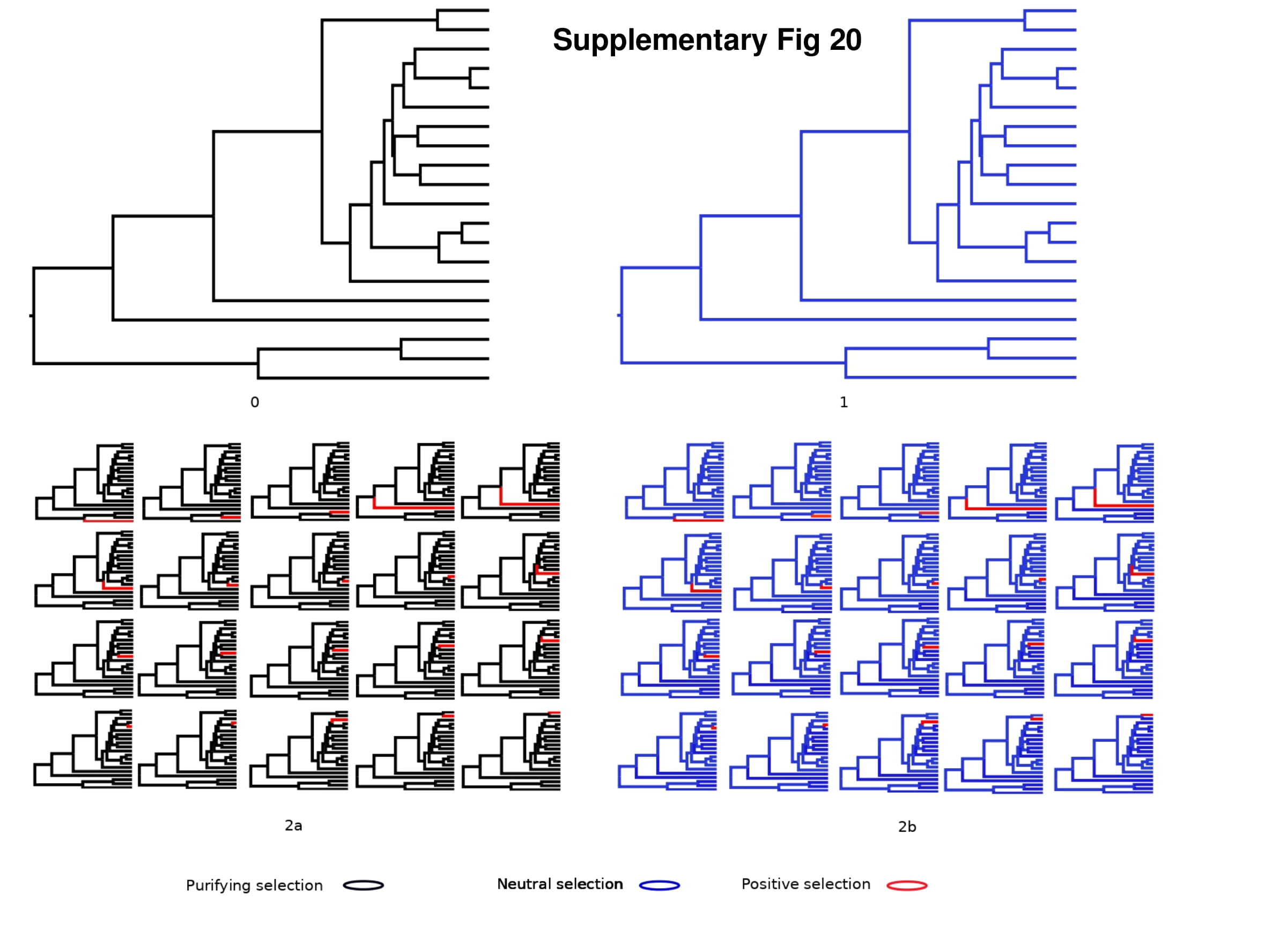

### supplementary figure 21

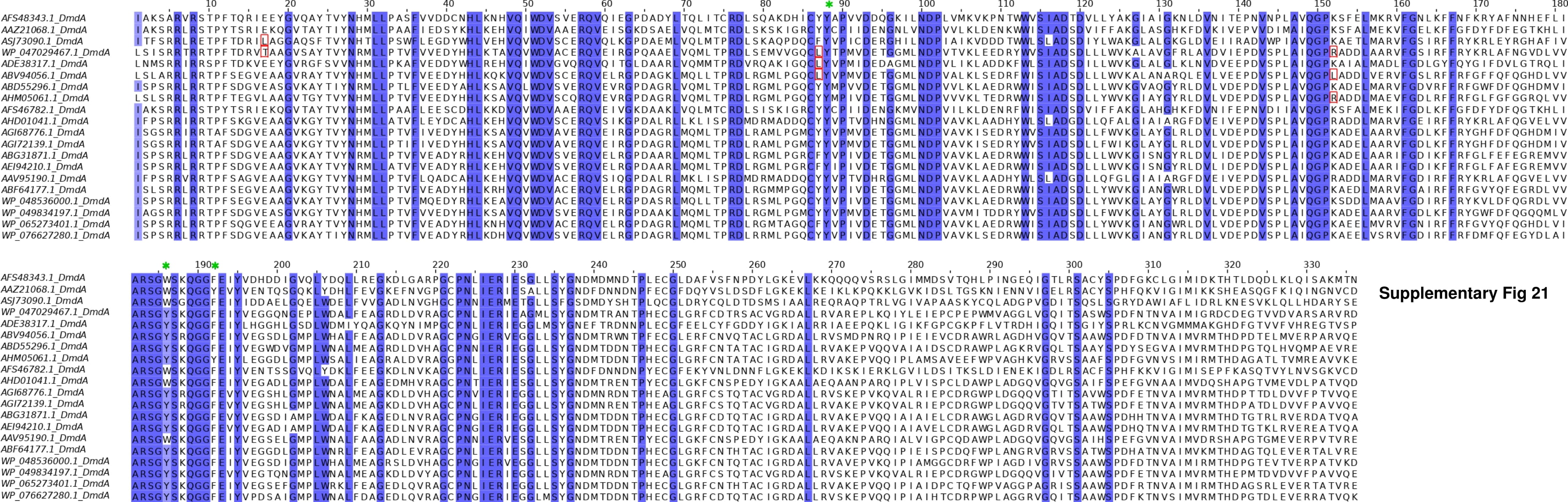

### supplementary figure 22

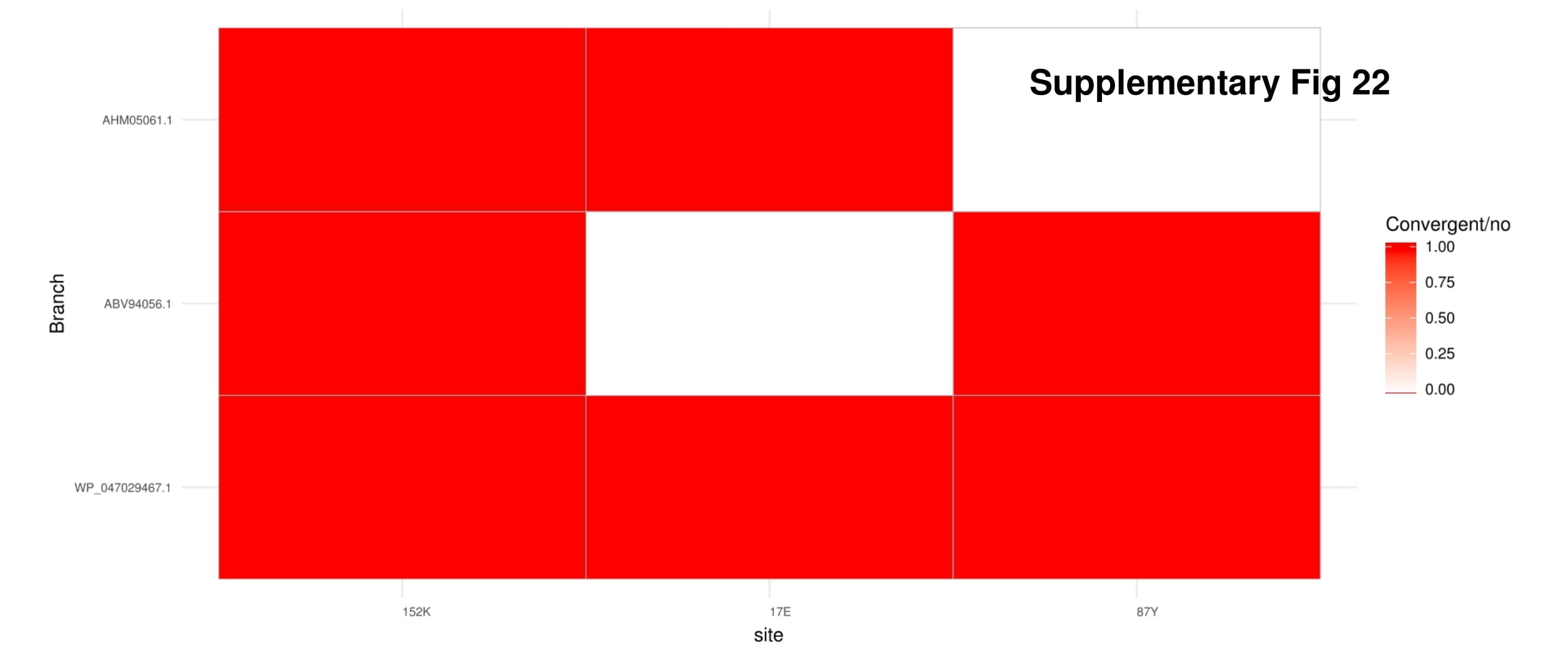

### supplementary figure 23

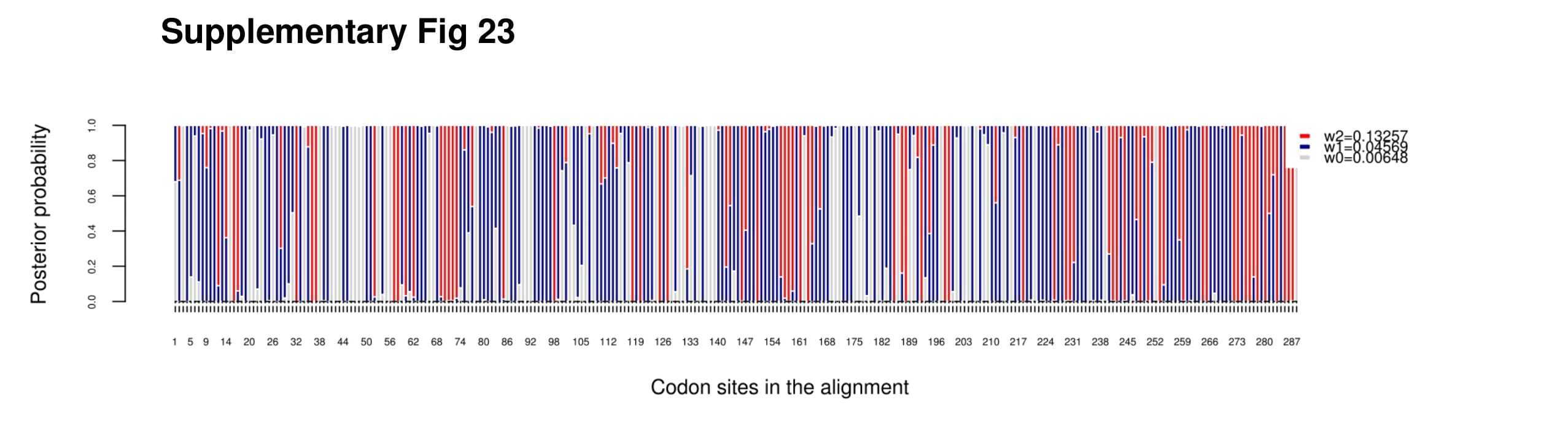

### supplementary figure 24

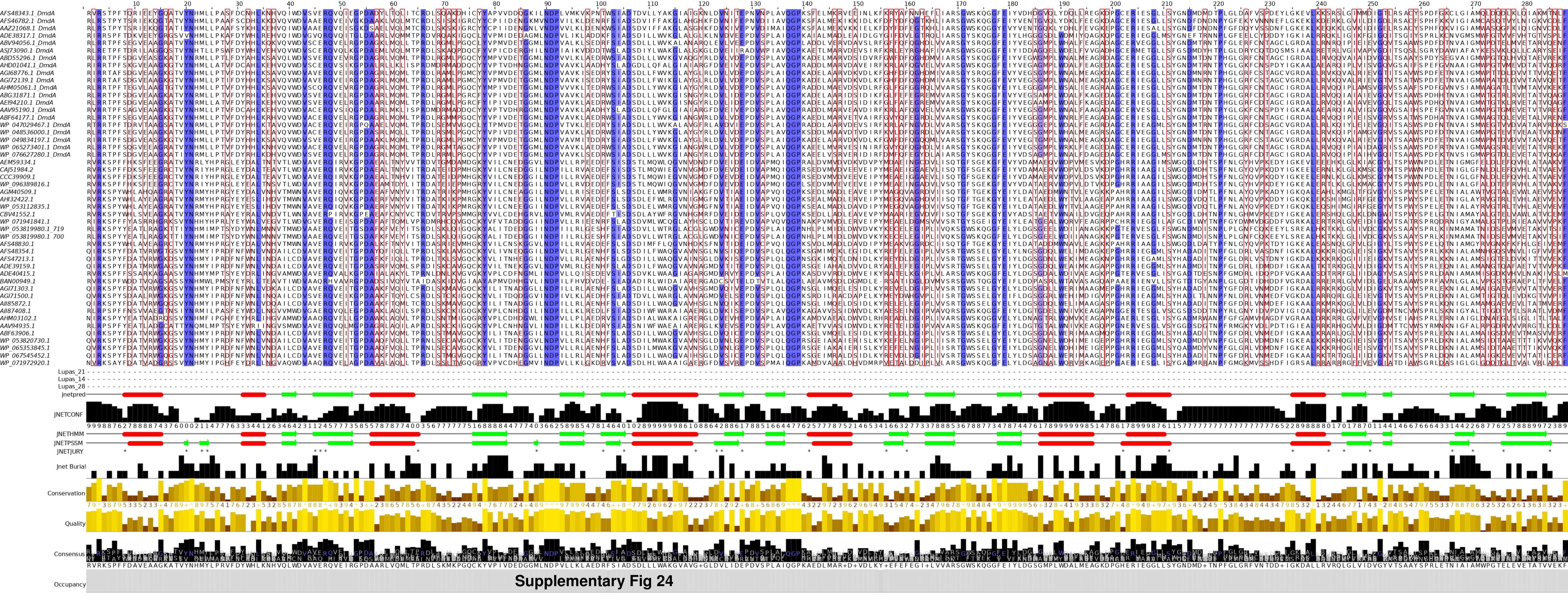

### supplementary figure 25

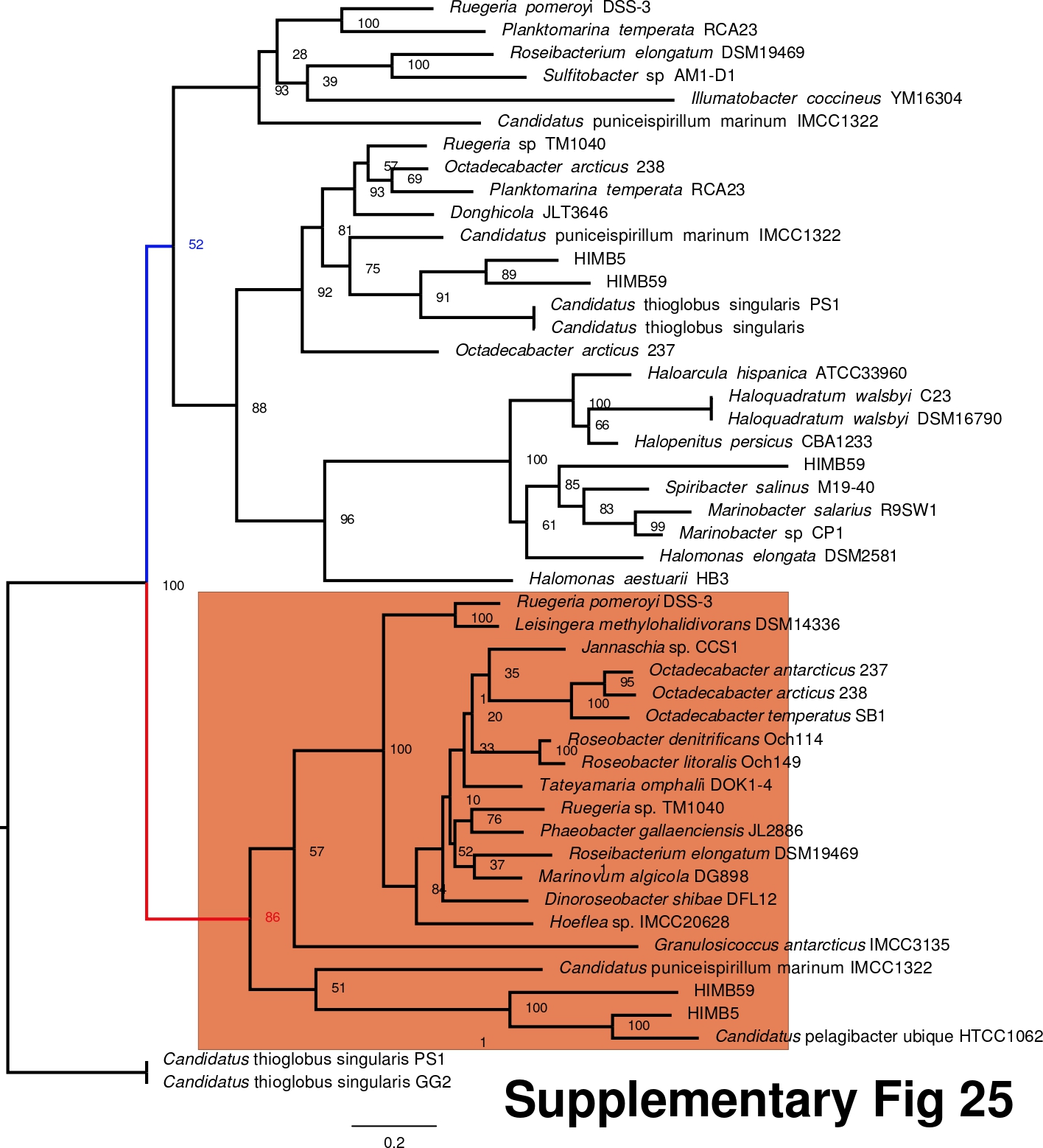
