## supplementary data 1 for "Evolutionary history of dimethylsulfoniopropionate (DMSP) demethylation enzyme DmdA in marine bacteria"

**Table 1**. Up: Top ten analogs identified by LOMETS for threading alignments. Case of AAV94935.1. In the middle: Model predicted by I-TASSER. Down: Top ten identified structural analogs in PDB by TM-align

| **Rank** | **Class** | **Gene name** | **Organism** | **PDB ID** | **Iden1 (%)^a^** | **Iden2 (%)^b^** | **Cov^c^** | **Norm. Z-score^d^** |
| --- | --- | --- | --- | --- | --- | --- | --- | --- |
| 1 | AMT^e^ | *dmdA^f^* | *Pelagibacter ubique (HTCC1062)* | 3tfhA | 40 | 40 | 1.00 | 4.58 |
| 2 | AMT | *dmdA* | *Pelagibacter ubique (HTCC1062)* | 3tfhA | 39 | 40 | 1.00 | 5.33 |
| 3 | AMT | *dmdA* | *Pelagibacter ubique (HTCC1062)* | 3tfhA | 38 | 40 | 0.99 | 5.44 |
| 4 | AMT | *dmdA* | *Pelagibacter ubique (HTCC1062)* | 3tfh | 39 | 40 | 0.99 | 3.25 |
| 5 | Oxidoreductase | *dmgdH*^h^ | *Rattus norvegicus* | 4p9sA | 21 | 25 | 0.97 | 2.40 |
| 6 | AMT | *dmdA* | *Pelagibacter ubique (HTCC1062)* | 3tfhA | 38 | 40 | 0.98 | 5.58 |
| 7 | AMT | *dmdA* | *Pelagibacter ubique (HTCC1062)* | 3tfh | 40 | 40 | 0.98 | 3.61 |
| 8 | AMT | *gcvT^g^* | *Bacillus subtilis* | 1yx2A | 24 | 24 | 0.96 | 6.66 |
| 9 | AMT | *dmdA* | *Pelagibacter ubique (HTCC1062)* | 3tfhA | 39 | 40 | 0.99 | 7.29 |
| 10 | AMT | *dmdA* | *Pelagibacter ubique (HTCC1062)* | 3tfhA | 39 | 40 | 1.00 | 6.72 |

^a^Iden1 is the percentage sequence identity of the templates in the threading aligned region with the query sequence

^b^Iden2 is the percentage sequence identity of the whole template chains with query sequence

^c^Cov-Represents the coverage of the threading alignment and is equal to the number of aligned residues divided by the length of the query protein

^d^Norm. Z-score is the normalized Z-score of the threading alignments. Alignment with a normalized Z-score>1 means a good alignment and vice versa.

^e^Aminomethyltransferase

^f^DmdA DMSP-dependent demethylase

^g^Glycine cleavage system T protein

^h^Dimethylglycine dehydrogenase complexed with tetrahydrofolate

**Predicted model: C-sore= 1.96; TM-score=0.99 dev= 0.04**

| **Rank** | **Class** | **Gene name** | **Organism** | **PDB ID** | **TM-score^a^** | **RMSD^b^** | **IDEN^c^** | **Cov^d^** |
| --- | --- | --- | --- | --- | --- | --- | --- | --- |
| 1 | AMT | *dmdA* | *Pelagibacter ubique (HTCC1062)* | 3tfhA | 0.994 | 0.43 | 0.393 | 0.997 |
| 2 | Oxidoreductase | *dmgdH* | *Rattus norvegicus* | 4p9sA2 | 0.927 | 1.80 | 0.217 | 0.981 |
| 3 | Oxidoreductase | *dmg^e^* | *Arthrobacter globiformis* | 1pj6A | 0.914 | 2.16 | 0.249 | 0.984 |
| 4 | AMT | *gcvT* | *Pyrococcus horikoshii* | 1v5vA | 0.901 | 1.96 | 0.240 | 0.965 |
| 5 | AMT | *gcvT* | *Thermotoga maritima* | 1wooA | 0.898 | 1.73 | 0.261 | 0.948 |
| 6 | Oxidoreductase | *soxA^f^* | *Stenotrophomonas maltophilia* | 2gagA | 0.898 | 2.30 | 0.224 | 0.976 |
| 7 | Oxidoreductase | *soxA* | *Corynebacterium sp.* | 1vrqA | 0.897 | 2.28 | 0.235 | 0.976 |
| 8 | AMT | *AMT* | *Homo sapiens* | 1wsrA | 0.888 | 2.09 | 0.222 | 0.956 |
| 9 | AMT | *gcvT* | *Bartonella henselae* | 3girA | 0.882 | 1.96 | 0.237 | 0.943 |
| 10 | AMT | *gcvT* | *Bacillus subtilis* | 1yx2B | 0.877 | 2.09 | 0.245 | 0.946 |

^ab^It is a standard for measuring structural similarity between two structures

^c^It is the percentage sequence identity in the structurally aligned region

^d^It represents the coverage of the alignment by TM-align and is equal to the number of structurally aligned residues divided by length of the query protein

^e^Dimethylglycine oxidase

^f^Heterotetrameric sarcosine oxidase alpha-subunit

**Table 2**. Up: Top ten analogs identified by LOMETS for threading alignments. Case of AII87408.1. In the middle: Model predicted by I-TASSER. Down: Top ten identified structural analogs in PDB by TM-align

| **Rank** | **Class** | **Gene name** | **Organism** | **PDB ID** | **Iden1 (%)^a^** | **Iden2 (%)^b^** | **Cov^c^** | **Norm. Z-score^d^** |
| --- | --- | --- | --- | --- | --- | --- | --- | --- |
| 1 | AMT^e^ | *dmdA^f^* | *Pelagibacter ubique (HTCC1062)* | 3tfhA | 36 | 37 | 0.99 | 4.64 |
| 2 | AMT | *dmdA* | *Pelagibacter ubique (HTCC1062)* | 3tfhA | 36 | 37 | 1.00 | 5.27 |
| 3 | AMT | *dmdA* | *Pelagibacter ubique (HTCC1062)* | 3tfhA | 36 | 37 | 0.99 | 5.57 |
| 4 | Oxidoreductase | *dmgdH^h^* | *Rattus norvegicus* | 4p9s | 22 | 31 | 0.96 | 3.26 |
| 5 | Oxidoreductase | *dmgdH* | *Rattus norvegicus* | 4p9sA | 21 | 24 | 0.98 | 2.42 |
| 6 | AMT | *dmdA* | *Pelagibacter ubique (HTCC1062)* | 3tfhA | 36 | 37 | 0.99 | 5.74 |
| 7 | AMT | *dmdA* | *Pelagibacter ubique (HTCC1062)* | 3tfh | 36 | 37 | 0.98 | 3.65 |
| 8 | AMT | *gcvT^g^* | *Bacillus subtilis* | 1yx2A | 26 | 28 | 0.96 | 6.47 |
| 9 | AMT | *dmdA* | *Pelagibacter ubique (HTCC1062)* | 3tfhA | 37 | 37 | 0.99 | 7.38 |
| 10 | AMT | *dmdA* | *Pelagibacter ubique (HTCC1062)* | 3tfhA | 36 | 37 | 0.99 | 6.67 |

^a^Iden1 is the percentage sequence identity of the templates in the threading aligned region with the query sequence

^b^Iden2 is the percentage sequence identity of the whole template chains with query sequence

^c^Cov-Represents the coverage of the threading alignment and is equal to the number of aligned residues divided by the length of the query protein

^d^Norm. Z-score is the normalized Z-score of the threading alignments. Alignment with a normalized Z-score>1 means a good alignment and vice versa.

^e^Aminomethyltransferase

^f^DmdA DMSP-dependent demethylase

^g^Glycine cleavage system T protein

^h^Dimethylglycine dehydrogenase complexed with tetrahydrofolate

**Predicted model: C-sore= 1.64; TM-score=0.94 dev= 0.05**

| **Rank** | **Class** | **Gene name** | **Organism** | **PDB ID** | **TM-score^a^** | **RMSD^b^** | **IDEN^c^** | **Cov^d^** |
| --- | --- | --- | --- | --- | --- | --- | --- | --- |
| 1 | AMT | *dmdA* | *Pelagibacter ubique (HTCC1062)* | 3tfhA | 0.985 | 0.81 | 0.365 | 0.997 |
| 2 | Oxidoreductase | *dmgdH* | *Rattus norvegicus* | 4p9sA2 | 0.938 | 1.60 | 0.230 | 0.981 |
| 3 | Oxidoreductase | *dmg^e^* | *Arthrobacter globiformis* | 1pj6A | 0.920 | 2.01 | 0.260 | 0.981 |
| 4 | AMT | *gcvT* | *Pyrococcus horikoshii* | 1v5vA | 0.908 | 1.86 | 0.225 | 0.965 |
| 5 | AMT | *gcvT* | *Thermotoga maritima* | 1wooA | 0.902 | 1.68 | 0.244 | 0.948 |
| 6 | Oxidoreductase | *soxA^f^* | *Stenotrophomonas maltophilia* | 2gagA | 0.899 | 2.28 | 0.213 | 0.973 |
| 7 | Oxidoreductase | *soxA* | *Corynebacterium sp.* | 1vrqA | 0.898 | 2.25 | 0.221 | 0.973 |
| 8 | AMT | *AMT* | *Homo sapiens* | 1wsrA | 0.893 | 1.96 | 0.231 | 0.954 |
| 9 | AMT | *gcvT* | *Bartonella henselae* | 3girA | 0.886 | 1.89 | 0.216 | 0.943 |
| 10 | AMT | *gcvT* | *Bacillus subtilis* | 1yx2B | 0.882 | 2.07 | 0.266 | 0.948 |

^ab^It is a standard for measuring structural similarity between two structures

^c^It is the percentage sequence identity in the structurally aligned region

^d^It represents the coverage of the alignment by TM-align and is equal to the number of structurally aligned residues divided by length of the query protein

^e^Dimethylglycine oxidase

^f^Heterotetrameric sarcosine oxidase alpha-subunit

**Table 3**. Up: Top ten analogs identified by LOMETS for threading alignments. Case of ADE40415.1. In the middle: Model predicted by I-TASSER. Down: Top ten identified structural analogs in PDB by TM-align

| **Rank** | **Class** | **Gene name** | **Organism** | **PDB ID** | **Iden1 (%)^a^** | **Iden2 (%)^b^** | **Cov^c^** | **Norm. Z-score^d^** |
| --- | --- | --- | --- | --- | --- | --- | --- | --- |
| 1 | AMT^e^ | *dmdA^f^* | *Pelagibacter ubique (HTCC1062)* | 3tfhA | 38 | 39 | 1.00 | 4.72 |
| 2 | AMT | *dmdA* | *Pelagibacter ubique (HTCC1062)* | 3tfhA | 38 | 39 | 1.00 | 5.45 |
| 3 | AMT | *dmdA* | *Pelagibacter ubique (HTCC1062)* | 3tfhA | 37 | 39 | 0.99 | 5.59 |
| 4 | AMT | *dmdA* | *Pelagibacter ubique (HTCC1062)* | 3tfh | 38 | 39 | 0.99 | 3.28 |
| 5 | Oxidoreductase | *dmgdH*^h^ | *Rattus norvegicus* | 4p9sA | 24 | 27 | 0.98 | 2.41 |
| 6 | AMT | *dmdA* | *Pelagibacter ubique (HTCC1062)* | 3tfhA | 38 | 39 | 0.99 | 5.78 |
| 7 | AMT | *dmdA* | *Pelagibacter ubique (HTCC1062)* | 3tfh | 37 | 39 | 0.98 | 3.63 |
| 8 | AMT | *gcvT^g^* | *Bacillus subtilis* | 1yx2A | 24 | 25 | 0.95 | 6.60 |
| 9 | AMT | *dmdA* | *Pelagibacter ubique (HTCC1062)* | 3tfhA | 38 | 39 | 1.00 | 7.52 |
| 10 | AMT | *dmdA* | *Pelagibacter ubique (HTCC1062)* | 3tfhA | 38 | 39 | 1.00 | 6.89 |

^a^Iden1 is the percentage sequence identity of the templates in the threading aligned region with the query sequence

^b^Iden2 is the percentage sequence identity of the whole template chains with query sequence

^c^Cov-Represents the coverage of the threading alignment and is equal to the number of aligned residues divided by the length of the query protein

^d^Norm. Z-score is the normalized Z-score of the threading alignments. Alignment with a normalized Z-score>1 means a good alignment and vice versa.

^e^Aminomethyltransferase

^f^DmdA DMSP-dependent demethylase

^g^Glycine cleavage system T protein

^h^Dimethylglycine dehydrogenase complexed with tetrahydrofolate

**Predicted model: C-sore= 2; TM-score=0.99 dev= 0.04**

| **Rank** | **Class** | **Gene name** | **Organism** | **PDB ID** | **TM-score^a^** | **RMSD^b^** | **IDEN^c^** | **Cov^d^** |
| --- | --- | --- | --- | --- | --- | --- | --- | --- |
| 1 | AMT | *dmdA* | *Pelagibacter ubique (HTCC1062)* | 3tfhA | 0.995 | 0.37 | 0.375 | 0.997 |
| 2 | Oxidoreductase | *dmgdH* | *Rattus norvegicus* | 4p9sA2 | 0.925 | 1.88 | 0.235 | 0.981 |
| 3 | Oxidoreductase | *dmg^e^* | *Arthrobacter globiformis* | 1pj6A | 0.908 | 2.25 | 0.254 | 0.981 |
| 4 | AMT | *gcvT* | *Pyrococcus horikoshii* | 1v5vA | 0.900 | 1.94 | 0.262 | 0.962 |
| 6 | Oxidoreductase | *soxA^f^* | *Stenotrophomonas maltophilia* | 2gagA | 0.895 | 2.31 | 0.212 | 0.970 |
| 7 | Oxidoreductase | *soxA* | *Corynebacterium sp.* | 1vrqA | 0.894 | 2.41 | 0.220 | 0.976 |
|  | AMT | *gcvT* | *Thermotoga maritima* | 1wooA | 0.893 | 1.74 | 0.276 | 0.943 |
| 8 | AMT | *AMT* | *Homo sapiens* | 1wsrA | 0.887 | 2.08 | 0.199 | 0.954 |
| 9 | AMT | *gcvT* | *Bartonella henselae* | 3girA | 0.881 | 1.93 | 0.219 | 0.940 |
| 10 | AMT | *gcvT* | *Bacillus subtilis* | 1yx2B | 0.875 | 2.09 | 0.247 | 0.943 |

^ab^It is a standard for measuring structural similarity between two structures

^c^It is the percentage sequence identity in the structurally aligned region

^d^It represents the coverage of the alignment by TM-align and is equal to the number of structurally aligned residues divided by length of the query protein

^e^Dimethylglycine oxidase

^f^Heterotetrameric sarcosine oxidase alpha-subunit

**Table 4**. Up: Top ten analogs identified by LOMETS for threading alignments. Case of BAN00949.1. In the middle: Model predicted by I-TASSER. Down: Top ten identified structural analogs in PDB by TM-align

| **Rank** | **Class** | **Gene name** | **Organism** | **PDB ID** | **Iden1 (%)^a^** | **Iden2 (%)^b^** | **Cov^c^** | **Norm. Z-score^d^** |
| --- | --- | --- | --- | --- | --- | --- | --- | --- |
| 1 | AMT^e^ | *gcvT* | *Thermotoga maritima (ATCC 43589)* | 1wopA | 25 | 25 | 0.88 | 4.29 |
| 2 | AMT^e^ | *dmdA^f^* | *Pelagibacter ubique (HTCC1062)* | 3tfhA | 33 | 30 | 0.90 | 5.18 |
| 3 | AMT^e^ | *dmdA* | *Pelagibacter ubique (HTCC1062)* | 3tfhA | 31 | 30 | 0.90 | 5.03 |
| 4 | Oxidoreductase | *dmg^i^* | *Arthrobacter globiformis* | 1pj6 | 23 | 27 | 0.97 | 3.32 |
| 5 | Oxidoreductase | *dmgdH*^h^ | *Rattus norvegicus* | 4p9sA | 20 | 23 | 0.92 | 2.40 |
| 6 | AMT^e^ | *dmdA* | *Pelagibacter ubique (HTCC1062)* | 3tfhA | 31 | 30 | 0.90 | 5.14 |
| 7 | *Oxidoreductase* | *dmg* | *Arthrobacter globiformis* | 1pj6 | 23 | 27 | 0.95 | 3.62 |
| 8 | AMT^e^ | *gcvT^g^* | *Bacillus subtilis* | 1yx2A | 25 | 22 | 0.87 | 6.32 |
| 9 | AMT^e^ | *dmdA* | *Pelagibacter ubique (HTCC1062)* | 3tfhA | 32 | 30 | 0.91 | 6.84 |
| 10 | AMT^e^ | *dmdA* | *Pelagibacter ubique (HTCC1062)* | 3tfhA | 32 | 30 | 0.91 | 6.17 |

^a^Iden1 is the percentage sequence identity of the templates in the threading aligned region with the query sequence

^b^Iden2 is the percentage sequence identity of the whole template chains with query sequence

^c^Cov-Represents the coverage of the threading alignment and is equal to the number of aligned residues divided by the length of the query protein

^d^Norm. Z-score is the normalized Z-score of the threading alignments. Alignment with a normalized Z-score>1 means a good alignment and vice versa.

^e^Aminomethyltransferase

^f^DmdA DMSP-dependent demethylase

^g^Glycine cleavage system T protein

^h^Dimethylglycine dehydrogenase complexed with tetrahydrofolate

^i^Dimethylglycine oxidase

**Predicted model: C-sore= 1.13; TM-score=0.87 dev= 0.07**

| **Rank** | **Class** | **Gene name** | **Organism** | **PDB ID** | **TM-score^a^** | **RMSD^b^** | **IDEN^c^** | **Cov^d^** |
| --- | --- | --- | --- | --- | --- | --- | --- | --- |
| 1 | Oxidoreductase | *dmg* | *Arthrobacter globiformis* | 1pj6A | 0.948 | 1.62 | 0.229 | 0.988 |
| 2 | Oxidoreductase | *dmgdH* | *Rattus norvegicus* | 4p9sA2 | 0.943 | 1.63 | 0.192 | 0.985 |
| 3 | AMT | *dmdA* | *Pelagibacter ubique HTCC1062* | 3tfhA | 0.879 | 1.38 | 0.326 | 0.908 |
| 6 | Oxidoreductase | *soxA^e^* | *Stenotrophomonas maltophilia* | 2gagA | 0.859 | 2.18 | 0.180 | 0.920 |
| 7 | Oxidoreductase | *soxA* | *Corynebacterium sp.* | 1vrqA | 0.858 | 2.21 | 0.185 | 0.920 |
|  | AMT | *gcvT* | *Thermotoga maritima* | 1wooA | 0.853 | 1.43 | 0.254 | 0.883 |
|  | AMT | *gcvT* | *Pyrococcus horikoshii* | 1v5vA | 0.850 | 1.96 | 0.238 | 0.898 |
| 8 | AMT | *AMT* | *Homo sapiens* | 1wsrA | 0.839 | 1.77 | 0.208 | 0.883 |
| 9 | AMT | *gcvT* | *Bartonella henselae* | 3girA | 0.831 | 1.85 | 0.207 | 0.878 |
| 10 | AMT | *gcvT* | *Bacillus subtilis* | 1yx2B | 0.825 | 1.70 | 0.247 | 0.866 |

^ab^It is a standard for measuring structural similarity between two structures

^c^It is the percentage sequence identity in the structurally aligned region

^d^It represents the coverage of the alignment by TM-align and is equal to the number of structurally aligned residues divided by length of the query protein

^e^Heterotetrameric sarcosine oxidase alpha-subunit

**Table 5**. Up: Top ten analogs identified by LOMETS for threading alignments. Case of AII85872.1. In the middle: Model predicted by I-TASSER. Down: Top ten identified structural analogs in PDB by TM-align

| **Rank** | **Class** | **Gene name** | **Organism** | **PDB ID** | **Iden1 (%)^a^** | **Iden2 (%)^b^** | **Cov^c^** | **Norm. Z-score^d^** |
| --- | --- | --- | --- | --- | --- | --- | --- | --- |
| 1 | AMT^e^ | *dmdA^f^* | *Pelagibacter ubique (HTCC1062)* | 3tfhA | 36 | 36 | 0.97 | 4.77 |
| 2 | AMT | *dmdA* | *Pelagibacter ubique (HTCC1062)* | 3tfhA | 36 | 36 | 0.97 | 5.42 |
| 3 | AMT | *dmdA* | *Pelagibacter ubique (HTCC1062)* | 3tfhA | 36 | 36 | 0.97 | 5.49 |
| 4 | Oxidoreductase | *dmg^i^* | *Arthrobacter globiformis* | 1pj6 | 26 | 30 | 0.98 | 3.25 |
| 5 | Oxidoreductase | *dmgdH*^h^ | *Rattus norvegicus* | 4p9sA | 22 | 26 | 0.97 | 2.40 |
| 6 | AMT | *dmdA* | *Pelagibacter ubique (HTCC1062)* | 3tfhA | 36 | 36 | 0.97 | 5.73 |
| 7 | *Oxidoreductase* | *dmg* | *Arthrobacter globiformis* | 1pj6 | 25 | 30 | 0.97 | 3.63 |
| 8 | AMT | *gcvT^g^* | *Bacillus subtilis* | 1yx2A | 28 | 28 | 0.93 | 6.47 |
| 9 | AMT | *dmdA* | *Pelagibacter ubique (HTCC1062)* | 3tfhA | 36 | 36 | 0.97 | 7.77 |
| 10 | AMT | *dmdA* | *Pelagibacter ubique (HTCC1062)* | 3tfhA | 36 | 36 | 0.97 | 6.72 |

^a^Iden1 is the percentage sequence identity of the templates in the threading aligned region with the query sequence

^b^Iden2 is the percentage sequence identity of the whole template chains with query sequence

^c^Cov-Represents the coverage of the threading alignment and is equal to the number of aligned residues divided by the length of the query protein

^d^Norm. Z-score is the normalized Z-score of the threading alignments. Alignment with a normalized Z-score>1 means a good alignment and vice versa.

^e^Aminomethyltransferase

^f^DmdA DMSP-dependent demethylase

^g^Glycine cleavage system T protein

^h^Dimethylglycine dehydrogenase complexed with tetrahydrofolate

^i^Dimethylglycine oxidase

**Predicted model: C-sore= 1.52; TM-score=0.93 dev= 0.06**

| **Rank** | **Class** | **Gene name** | **Organism** | **PDB ID** | **TM-score^a^** | **RMSD^b^** | **IDEN^c^** | **Cov^d^** |
| --- | --- | --- | --- | --- | --- | --- | --- | --- |
| 1 | AMT | *dmdA* | *Pelagibacter ubique (HTCC1062)* | 3tfhA | 0.960 | 0.76 | 0.358 | 0.971 |
| 2 | Oxidoreductase | *dmgdH* | *Rattus norvegicus* | 4p9sA2 | 0.929 | 1.85 | 0.228 | 0.982 |
| 3 | Oxidoreductase | *dmg* | *Arthrobacter globiformis* | 1pj6A | 0.926 | 1.95 | 0.255 | 0.982 |
| 6 | Oxidoreductase | *soxA^e^* | *Stenotrophomonas maltophilia* | 2gagA | 0.896 | 2.25 | 0.224 | 0.968 |
| 7 | AMT | *gcvT* | *Pyrococcus horikoshii ATCC700860* | 1v5vA | 0.895 | 2.10 | 0.269 | 0.958 |
|  | Oxidoreductase | *soxA* | *Corynebacterium sp.* | 1vrqA | 0.893 | 2.25 | 0.222 | 0.966 |
|  | AMT | *gcvT* | *Thermotoga maritima* | 1wooA | 0.888 | 1.73 | 0.256 | 0.934 |
| 8 | AMT | *AMT* | *Homo sapiens* | 1wsrA | 0.884 | 2.02 | 0.226 | 0.945 |
| 9 | AMT | *gcvT* | *Bartonella henselae* | 3girA | 0.875 | 1.86 | 0.218 | 0.929 |
| 10 | AMT | *gcvT* | *Bacillus subtilis* | 1yx2B | 0.864 | 1.99 | 0.279 | 0.924 |

^ab^It is a standard for measuring structural similarity between two structures

^c^It is the percentage sequence identity in the structurally aligned region

^d^It represents the coverage of the alignment by TM-align and is equal to the number of structurally aligned residues divided by length of the query protein

^e^Heterotetrameric sarcosine oxidase alpha-subunit

**Table 6**. Up: Top ten analogs identified by LOMETS for threading alignments. Case of WP_071972920. In the middle: Model predicted by I-TASSER. Down: Top ten identified structural analogs in PDB by TM-align

| **Rank** | **Class** | **Gene name** | **Organism** | **PDB ID** | **Iden1 (%)^a^** | **Iden2 (%)^b^** | **Cov^c^** | **Norm. Z-score^d^** |
| --- | --- | --- | --- | --- | --- | --- | --- | --- |
| 1 | AMT^e^ | *dmdA^f^* | *Pelagibacter ubique (HTCC1062)* | 3tfhA | 35 | 34 | 0.99 | 4.54 |
| 2 | AMT | *dmdA* | *Pelagibacter ubique (HTCC1062)* | 3tfhA | 35 | 34 | 0.99 | 5.33 |
| 3 | AMT | *dmdA* | *Pelagibacter ubique (HTCC1062)* | 3tfhA | 34 | 34 | 0.99 | 5.59 |
| 4 | Oxidoreductase | *dmg^i^* | *Arthrobacter globiformis* | 3tfh | 34 | 34 | 0.99 | 3.31 |
| 5 | Oxidoreductase | *dmgdH*^h^ | *Rattus norvegicus* | 4p9sA | 19 | 23 | 0.97 | 2.40 |
| 6 | AMT | *dmdA* | *Pelagibacter ubique (HTCC1062)* | 3tfhA | 34 | 34 | 0.99 | 5.62 |
| 7 | *Oxidoreductase* | *mgdH^h^* | *Rattus norvegicus* | 4p9sA | 19 | 23 | 0.96 | 3.59 |
| 8 | AMT | *gcvT^g^* | *Bacillus subtilis* | 1yx2A | 24 | 25 | 0.95 | 6.58 |
| 9 | AMT | *dmdA* | *Pelagibacter ubique (HTCC1062)* | 3tfhA | 35 | 34 | 0.99 | 7.07 |
| 10 | AMT | *dmdA* | *Pelagibacter ubique (HTCC1062)* | 3tfhA | 34 | 34 | 0.99 | 6.73 |

^a^Iden1 is the percentage sequence identity of the templates in the threading aligned region with the query sequence

^b^Iden2 is the percentage sequence identity of the whole template chains with query sequence

^c^Cov-Represents the coverage of the threading alignment and is equal to the number of aligned residues divided by the length of the query protein

^d^Norm. Z-score is the normalized Z-score of the threading alignments. Alignment with a normalized Z-score>1 means a good alignment and vice versa.

^e^Aminomethyltransferase

^f^DmdA DMSP-dependent demethylase

^g^Glycine cleavage system T protein

^h^Dimethylglycine dehydrogenase complexed with tetrahydrofolate

^i^Dimethylglycine oxidase

**Predicted model: C-sore= 1.99; TM-score=0.99 dev= 0.04**

| **Rank** | **Class** | **Gene name** | **Organism** | **PDB ID** | **TM-score^a^** | **RMSD^b^** | **IDEN^c^** | **Cov^d^** |
| --- | --- | --- | --- | --- | --- | --- | --- | --- |
| 1 | AMT | *dmdA* | *Pelagibacter ubique (HTCC1062)* | 3tfhA | 0.988 | 0.42 | 0.346 | 0.992 |
| 2 | Oxidoreductase | *dmgdH* | *Rattus norvegicus* | 4p9sA2 | 0.921 | 1.82 | 0.197 | 0.976 |
| 3 | Oxidoreductase | *dmg* | *Arthrobacter globiformis* | 1pj6A | 0.908 | 2.18 | 0.224 | 0.978 |
| 4 | AMT | *gcvT* | *Pyrococcus horikoshii ATCC700860* | 1v5vA | 0.901 | 2.06 | 0.240 | 0.968 |
| 5 | Oxidoreductase | *soxA^e^* | *Stenotrophomonas maltophilia* | 2gagA | 0.893 | 2.31 | 0.242 | 0.968 |
| 6 | Oxidoreductase | *soxA* | *Corynebacterium sp.* | 1vrqA | 0.892 | 2.27 | 0.237 | 0.968 |
| 7 | AMT | *gcvT* | *Thermotoga maritima* | 1wooA | 0.890 | 1.80 | 0.244 | 0.943 |
| 8 | AMT | *AMT* | *Homo sapiens* | 1wsrA | 0.884 | 2.02 | 0.208 | 0.949 |
| 9 | AMT | *gcvT* | *Bartonella henselae* | 3girA | 0.878 | 1.90 | 0.202 | 0.935 |
| 10 | AMT | *gcvT* | *Bacillus subtilis* | 1yx2B | 0.869 | 2.16 | 0.239 | 0.941 |

^ab^It is a standard for measuring structural similarity between two structures

^c^It is the percentage sequence identity in the structurally aligned region

^d^It represents the coverage of the alignment by TM-align and is equal to the number of structurally aligned residues divided by length of the query protein

^e^Heterotetrameric sarcosine oxidase alpha-subunit

**Table 7**. Up: Top ten analogs identified by LOMETS for threading alignments. Case of AHM03102.1. In the middle: Model predicted by I-TASSER. Down: Top ten identified structural analogs in PDB by TM-align

| **Rank** | **Class** | **Gene name** | **Organism** | **PDB ID** | **Iden1 (%)^a^** | **Iden2 (%)^b^** | **Cov^c^** | **Norm. Z-score^d^** |
| --- | --- | --- | --- | --- | --- | --- | --- | --- |
| 1 | AMT^e^ | *dmdA^f^* | *Pelagibacter ubique (HTCC1062)* | 3tfhA | 37 | 36 | 0.99 | 4.64 |
| 2 | AMT | *dmdA* | *Pelagibacter ubique (HTCC1062)* | 3tfhA | 37 | 36 | 0.99 | 5.36 |
| 3 | AMT | *dmdA* | *Pelagibacter ubique (HTCC1062)* | 3tfhA | 36 | 36 | 0.99 | 5.47 |
| 4 | Oxidoreductase | *dmg^i^* | *Rattus norvegicus* | 4p9s | 21 | 30 | 0.97 | 3.25 |
| 5 | Oxidoreductase | *dmgdH*^h^ | *Rattus norvegicus* | 4p9sA | 20 | 24 | 0.98 | 2.42 |
| 6 | AMT | *dmdA* | *Pelagibacter ubique (HTCC1062)* | 3tfhA | 36 | 36 | 0.98 | 5.68 |
| 7 | *Oxidoreductase* | *dmgdH^h^* | *Rattus norvegicus* | 4p9s | 19 | 30 | 0.97 | 3.59 |
| 8 | AMT | *gcvT^g^* | *Bacillus subtilis* | 1yx2A | 24 | 26 | 0.95 | 6.66 |
| 9 | AMT | *dmdA* | *Pelagibacter ubique (HTCC1062)* | 3tfhA | 37 | 36 | 0.99 | 7.38 |
| 10 | AMT | *dmdA* | *Pelagibacter ubique (HTCC1062)* | 3tfhA | 36 | 36 | 0.99 | 6.61 |

^a^Iden1 is the percentage sequence identity of the templates in the threading aligned region with the query sequence

^b^Iden2 is the percentage sequence identity of the whole template chains with query sequence

^c^Cov-Represents the coverage of the threading alignment and is equal to the number of aligned residues divided by the length of the query protein

^d^Norm. Z-score is the normalized Z-score of the threading alignments. Alignment with a normalized Z-score>1 means a good alignment and vice versa.

^e^Aminomethyltransferase

^f^DmdA DMSP-dependent demethylase

^g^Glycine cleavage system T protein

^h^Dimethylglycine dehydrogenase complexed with tetrahydrofolate

^i^Dimethylglycine oxidase

**Predicted model: C-sore= 1.69; TM-score=0.95 dev= 0.05**

| **Rank** | **Class** | **Gene name** | **Organism** | **PDB ID** | **TM-score^a^** | **RMSD^b^** | **IDEN^c^** | **Cov^d^** |
| --- | --- | --- | --- | --- | --- | --- | --- | --- |
| 1 | AMT | *dmdA* | *Pelagibacter ubique (HTCC1062)* | 3tfhA | 0.981 | 0.64 | 0.367 | 0.989 |
| 2 | Oxidoreductase | *dmgdH* | *Rattus norvegicus* | 4p9sA2 | 0.931 | 1.65 | 0.209 | 0.976 |
| 3 | Oxidoreductase | *dmg* | *Arthrobacter globiformis* | 1pj6A | 0.912 | 2.10 | 0.259 | 0.976 |
| 4 | AMT | *gcvT* | *Pyrococcus horikoshii ATCC700860* | 1v5vA | 0.906 | 1.94 | 0.226 | 0.965 |
| 5 | Oxidoreductase | *soxA^e^* | *Stenotrophomonas maltophilia* | 2gagA | 0.894 | 2.24 | 0.217 | 0.965 |
| 6 | Oxidoreductase | *soxA* | *Corynebacterium sp.* | 1vrqA | 0.894 | 2.22 | 0.228 | 0.965 |
| 7 | AMT | *gcvT* | *Thermotoga maritima* | 1wooA | 0.894 | 1.69 | 0.251 | 0.941 |
| 8 | AMT | *AMT* | *Homo sapiens* | 1wsrA | 0.888 | 1.98 | 0.232 | 0.949 |
| 9 | AMT | *gcvT* | *Bartonella henselae* | 3girA | 0.880 | 1.87 | 0.210 | 0.935 |
| 10 | AMT | *gcvT* | *Bacillus subtilis* | 1yx2B | 0.875 | 2.09 | 0.243 | 0.941 |

^ab^It is a standard for measuring structural similarity between two structures

^c^It is the percentage sequence identity in the structurally aligned region

^d^It represents the coverage of the alignment by TM-align and is equal to the number of structurally aligned residues divided by length of the query protein

^e^Heterotetrameric sarcosine oxidase alpha-subunit

**Table 8**. Up: Top ten analogs identified by LOMETS for threading alignments. Case of AGI71500. In the middle: Model predicted by I-TASSER. Down: Top ten identified structural analogs in PDB by TM-align

| **Rank** | **Class** | **Gene name** | **Organism** | **PDB ID** | **Iden1 (%)^a^** | **Iden2 (%)^b^** | **Cov^c^** | **Norm. Z-score^d^** |
| --- | --- | --- | --- | --- | --- | --- | --- | --- |
| 1 | AMT^e^ | *dmdA^f^* | *Pelagibacter ubique (HTCC1062)* | 3tfhA | 36 | 35 | 0.96 | 4.55 |
| 2 | AMT | *dmdA* | *Pelagibacter ubique (HTCC1062)* | 3tfhA | 36 | 35 | 0.96 | 5.33 |
| 3 | AMT | *dmdA* | *Pelagibacter ubique (HTCC1062)* | 3tfhA | 36 | 35 | 0.96 | 5.44 |
| 4 | Oxidoreductase | *dmg^i^* | *Rattus norvegicus* | 4p9sA | 22 | 27 | 0.97 | 3.28 |
| 5 | Oxidoreductase | *dmgdH*^h^ | *Rattus norvegicus* | 4p9sA | 36 | 35 | 0.96 | 2.41 |
| 6 | AMT | *dmdA* | *Pelagibacter ubique (HTCC1062)* | 3tfhA | 36 | 35 | 0.96 | 5.66 |
| 7 | *Oxidoreductase* | *dmgdH* | *Rattus norvegicus* | 4p9s | 22 | 29 | 0.98 | 3.64 |
| 8 | AMT | *gcvT^g^* | *Bacillus subtilis* | 1yx2A | 22 | 29 | 0.92 | 6.64 |
| 9 | AMT | *dmdA* | *Pelagibacter ubique (HTCC1062)* | 3tfhA | 36 | 35 | 0.95 | 7.57 |
| 10 | AMT | *dmdA* | *Pelagibacter ubique (HTCC1062)* | 3tfhA | 36 | 35 | 0.96 | 6.53 |

^a^Iden1 is the percentage sequence identity of the templates in the threading aligned region with the query sequence

^b^Iden2 is the percentage sequence identity of the whole template chains with query sequence

^c^Cov-Represents the coverage of the threading alignment and is equal to the number of aligned residues divided by the length of the query protein

^d^Norm. Z-score is the normalized Z-score of the threading alignments. Alignment with a normalized Z-score>1 means a good alignment and vice versa.

^e^Aminomethyltransferase

^f^DmdA DMSP-dependent demethylase

^g^Glycine cleavage system T protein

^h^Dimethylglycine dehydrogenase complexed with tetrahydrofolate

^i^Dimethylglycine oxidase

**Predicted model: C-sore= 1.47; TM-score=0.92 dev= 0.06**

| **Rank** | **Class** | **Gene name** | **Organism** | **PDB ID** | **TM-score^a^** | **RMSD^b^** | **IDEN^c^** | **Cov^d^** |
| --- | --- | --- | --- | --- | --- | --- | --- | --- |
| 1 | AMT | *dmdA* | *Pelagibacter ubique (HTCC1062)* | 3tfhA | 0.949 | 0.81 | 0.360 | 0.961 |
| 2 | Oxidoreductase | *dmgdH* | *Rattus norvegicus* | 4p9sA | 0.937 | 1.74 | 0.228 | 0.982 |
| 3 | Oxidoreductase | *dmg* | *Arthrobacter globiformis* | 1pj6A | 0.907 | 2.34 | 0.264 | 0.977 |
| 4 | AMT | *gcvT* | *Pyrococcus horikoshii ATCC700860* | 1v5vA | 0.886 | 2.09 | 0.245 | 0.948 |
| 5 | Oxidoreductase | *soxA^e^* | *Stenotrophomonas maltophilia* | 2gagA | 0.882 | 2.38 | 0.212 | 0.956 |
| 6 | Oxidoreductase | *soxA* | *Corynebacterium sp.* | 1vrqA | 0.881 | 2.37 | 0.231 | 0.956 |
| 7 | AMT | *gcvT* | *Thermotoga maritima* | 1wooA | 0.877 | 1.75 | 0.279 | 0.924 |
| 8 | AMT | *AMT* | *Homo sapiens* | 1wsrA | 0.874 | 2.05 | 0.256 | 0.935 |
| 9 | AMT | *gcvT* | *Bartonella henselae* | 3girA | 0.865 | 1.91 | 0.235 | 0.919 |
| 10 | AMT | *gcvT* | *Bacillus subtilis* | 1yx2B | 0.853 | 2.06 | 0.302 | 0.914 |

^ab^It is a standard for measuring structural similarity between two structures

^c^It is the percentage sequence identity in the structurally aligned region

^d^It represents the coverage of the alignment by TM-align and is equal to the number of structurally aligned residues divided by length of the query protein

^e^Heterotetrameric sarcosine oxidase alpha-subunit

**Table 9**. Up: Top ten analogs identified by LOMETS for threading alignments. Case of WP_067545452. In the middle: Model predicted by I-TASSER. Down: Top ten identified structural analogs in PDB by TM-align

| **Rank** | **Class** | **Gene name** | **Organism** | **PDB ID** | **Iden1 (%)^a^** | **Iden2 (%)^b^** | **Cov^c^** | **Norm. Z-score^d^** |
| --- | --- | --- | --- | --- | --- | --- | --- | --- |
| 1 | AMT^e^ | *dmdA^f^* | *Pelagibacter ubique (HTCC1062)* | 3tfhA | 36 | 35 | 0.97 | 4.66 |
| 2 | AMT | *dmdA* | *Pelagibacter ubique (HTCC1062)* | 3tfhA | 36 | 35 | 0.97 | 5.36 |
| 3 | AMT | *dmdA* | *Pelagibacter ubique (HTCC1062)* | 3tfhA | 35 | 35 | 0.97 | 5.56 |
| 4 | Oxidoreductase | *dmg^i^* | *Rattus norvegicus* | 4p9s | 24 | 33 | 0.97 | 3.26 |
| 5 | Oxidoreductase | *dmgdH*^h^ | *Rattus norvegicus* | 4p9sA | 21 | 26 | 0.98 | 2.40 |
| 6 | AMT | *dmdA* | *Pelagibacter ubique (HTCC1062)* | 3tfhA | 35 | 35 | 0.97 | 5.67 |
| 7 | *Oxidoreductase* | *dmgdH* | *Rattus norvegicus* | 4p9s | 22 | 33 | 0.97 | 3.61 |
| 8 | AMT | *gcvT^g^* | *Bacillus subtilis* | 1yx2A | 28 | 29 | 0.92 | 6.72 |
| 9 | AMT | *dmdA* | *Pelagibacter ubique (HTCC1062)* | 3tfhA | 36 | 35 | 0.97 | 7.48 |
| 10 | AMT | *dmdA* | *Pelagibacter ubique (HTCC1062)* | 3tfhA | 36 | 35 | 0.97 | 6.67 |

^a^Iden1 is the percentage sequence identity of the templates in the threading aligned region with the query sequence

^b^Iden2 is the percentage sequence identity of the whole template chains with query sequence

^c^Cov-Represents the coverage of the threading alignment and is equal to the number of aligned residues divided by the length of the query protein

^d^Norm. Z-score is the normalized Z-score of the threading alignments. Alignment with a normalized Z-score>1 means a good alignment and vice versa.

^e^Aminomethyltransferase

^f^DmdA DMSP-dependent demethylase

^g^Glycine cleavage system T protein

^h^Dimethylglycine dehydrogenase complexed with tetrahydrofolate

^i^Dimethylglycine oxidase

**Predicted model: C-sore= 1.59; TM-score=0.94 dev= 0.05**

| **Rank** | **Class** | **Gene name** | **Organism** | **PDB ID** | **TM-score^a^** | **RMSD^b^** | **IDEN^c^** | **Cov^d^** |
| --- | --- | --- | --- | --- | --- | --- | --- | --- |
| 1 | AMT | *dmdA* | *Pelagibacter ubique (HTCC1062)* | 3tfhA | 0.961 | 0.85 | 0.355 | 0.974 |
| 2 | Oxidoreductase | *dmgdH* | *Rattus norvegicus* | 4p9sA | 0.942 | 1.53 | 0.226 | 0.982 |
| 3 | Oxidoreductase | *dmg* | *Arthrobacter globiformis* | 1pj6A | 0.916 | 2.11 | 0.264 | 0.979 |
| 4 | AMT | *gcvT* | *Pyrococcus horikoshii ATCC700860* | 1v5vA | 0.898 | 2.07 | 0.266 | 0.960 |
| 5 | Oxidoreductase | *soxA^e^* | *Stenotrophomonas maltophilia* | 2gagA | 0.895 | 2.35 | 0.233 | 0.968 |
| 6 | Oxidoreductase | *soxA* | *Corynebacterium sp.* | 1vrqA | 0.894 | 2.27 | 0.231 | 0.966 |
| 7 | AMT | *gcvT* | *Thermotoga maritima* | 1wooA | 0.892 | 1.67 | 0.268 | 0.937 |
| 8 | AMT | *AMT* | *Homo sapiens* | 1wsrA | 0.887 | 2.02 | 0.234 | 0.947 |
| 9 | AMT | *gcvT* | *Bartonella henselae* | 3girA | 0.877 | 1.89 | 0.215 | 0.931 |
| 10 | AMT | *gcvT* | *Bacillus subtilis* | 1yx2B | 0.864 | 2.03 | 0.282 | 0.926 |

^ab^It is a standard for measuring structural similarity between two structures

^c^It is the percentage sequence identity in the structurally aligned region

^d^It represents the coverage of the alignment by TM-align and is equal to the number of structurally aligned residues divided by length of the query protein

^e^Heterotetrameric sarcosine oxidase alpha-subunit

**Table 10**. Up: Top ten analogs identified by LOMETS for threading alignments. Case of ABF63906.1. In the middle: Model predicted by I-TASSER. Down: Top ten identified structural analogs in PDB by TM-align

| **Rank** | **Class** | **Gene name** | **Organism** | **PDB ID** | **Iden1 (%)^a^** | **Iden2 (%)^b^** | **Cov^c^** | **Norm. Z-score^d^** |
| --- | --- | --- | --- | --- | --- | --- | --- | --- |
| 1 | AMT^e^ | *dmdA^f^* | *Pelagibacter ubique (HTCC1062)* | 3tfhA | 37 | 35 | 0.97 | 4.67 |
| 2 | AMT | *dmdA* | *Pelagibacter ubique (HTCC1062)* | 3tfhA | 36 | 35 | 0.97 | 5.39 |
| 3 | AMT | *dmdA* | *Pelagibacter ubique (HTCC1062)* | 3tfhA | 36 | 35 | 0.97 | 5.50 |
| 4 | Oxidoreductase | *dmg^i^* | *Rattus norvegicus* | 4p9s | 25 | 34 | 0.97 | 3.27 |
| 5 | Oxidoreductase | *dmgdH*^h^ | *Rattus norvegicus* | 4p9sA | 22 | 27 | 0.98 | 2.41 |
| 6 | AMT | *dmdA* | *Pelagibacter ubique (HTCC1062)* | 3tfhA | 36 | 35 | 0.96 | 5.62 |
| 7 | *Oxidoreductase* | *dmg* | *Arthrobacter globiformis* | 1pj6 | 25 | 32 | 0.97 | 3.62 |
| 8 | AMT | *gcvT^g^* | *Bacillus subtilis* | 1yx2A | 29 | 29 | 0.93 | 6.64 |
| 9 | AMT | *dmdA* | *Pelagibacter ubique (HTCC1062)* | 3tfhA | 37 | 35 | 0.97 | 7.50 |
| 10 | AMT | *dmdA* | *Pelagibacter ubique (HTCC1062)* | 3tfhA | 37 | 35 | 0.97 | 6.64 |

^a^Iden1 is the percentage sequence identity of the templates in the threading aligned region with the query sequence

^b^Iden2 is the percentage sequence identity of the whole template chains with query sequence

^c^Cov-Represents the coverage of the threading alignment and is equal to the number of aligned residues divided by the length of the query protein

^d^Norm. Z-score is the normalized Z-score of the threading alignments. Alignment with a normalized Z-score>1 means a good alignment and vice versa.

^e^Aminomethyltransferase

^f^DmdA DMSP-dependent demethylase

^g^Glycine cleavage system T protein

^h^Dimethylglycine dehydrogenase complexed with tetrahydrofolate

^i^Dimethylglycine oxidase

**Predicted model: C-sore= 1.53; TM-score=0.93 dev= 0.06**

| **Rank** | **Class** | **Gene name** | **Organism** | **PDB ID** | **TM-score^a^** | **RMSD^b^** | **IDEN^c^** | **Cov^d^** |
| --- | --- | --- | --- | --- | --- | --- | --- | --- |
| 1 | AMT | *dmdA* | *Pelagibacter ubique (HTCC1062)* | 3tfhA | 0.960 | 0.68 | 0.361 | 0.968 |
| 2 | Oxidoreductase | *dmgdH* | *Rattus norvegicus* | 4p9sA | 0.935 | 1.72 | 0.244 | 0.982 |
| 3 | Oxidoreductase | *dmg* | *Arthrobacter globiformis* | 1pj6A | 0.916 | 2.14 | 0.260 | 0.982 |
| 4 | AMT | *gcvT* | *Pyrococcus horikoshii ATCC700860* | 1v5vA | 0.895 | 2.03 | 0.267 | 0.955 |
| 5 | Oxidoreductase | *soxA^e^* | *Stenotrophomonas maltophilia* | 2gagA | 0.891 | 2.37 | 0.235 | 0.966 |
| 6 | Oxidoreductase | *soxA* | *Corynebacterium sp.* | 1vrqA | 0.889 | 2.40 | 0.233 | 0.966 |
| 7 | AMT | *gcvT* | *Thermotoga maritima* | 1wooA | 0.886 | 1.75 | 0.268 | 0.934 |
| 8 | AMT | *AMT* | *Homo sapiens* | 1wsrA | 0.881 | 2.03 | 0.254 | 0.942 |
| 9 | AMT | *gcvT* | *Bartonella henselae* | 3girA | 0.873 | 1.92 | 0.235 | 0.929 |
| 10 | AMT | *gcvT* | *Bacillus subtilis* | 1yx2B | 0.861 | 2.06 | 0.288 | 0.924 |

^ab^It is a standard for measuring structural similarity between two structures

^c^It is the percentage sequence identity in the structurally aligned region

^d^It represents the coverage of the alignment by TM-align and is equal to the number of structurally aligned residues divided by length of the query protein

^e^Heterotetrameric sarcosine oxidase alpha-subunit

**Table 11**. Up: Top ten analogs identified by LOMETS for threading alignments. Case of AGI71303.1. In the middle: Model predicted by I-TASSER. Down: Top ten identified structural analogs in PDB by TM-align

| **Rank** | **Class** | **Gene name** | **Organism** | **PDB ID** | **Iden1 (%)^a^** | **Iden2 (%)^b^** | **Cov^c^** | **Norm. Z-score^d^** |
| --- | --- | --- | --- | --- | --- | --- | --- | --- |
| 1 | AMT^e^ | *dmdA^f^* | *Pelagibacter ubique (HTCC1062)* | 3tfhA | 36 | 36 | 0.97 | 4.74 |
| 2 | AMT | *dmdA* | *Pelagibacter ubique (HTCC1062)* | 3tfhA | 36 | 36 | 0.97 | 5.39 |
| 3 | AMT | *dmdA* | *Pelagibacter ubique (HTCC1062)* | 3tfhA | 36 | 36 | 0.97 | 5.60 |
| 4 | Oxidoreductase | *dmg^i^* | *Rattus norvegicus* | 4p9s | 25 | 33 | 0.97 | 3.27 |
| 5 | Oxidoreductase | *dmgdH*^h^ | *Rattus norvegicus* | 4p9sA | 22 | 26 | 0.97 | 2.39 |
| 6 | AMT | *dmdA* | *Pelagibacter ubique (HTCC1062)* | 3tfhA | 36 | 36 | 0.97 | 5.72 |
| 7 | *Oxidoreductase* | *dmg* | *Arthrobacter globiformis* | 1pj6 | 26 | 32 | 0.97 | 3.62 |
| 8 | AMT | *gcvT^g^* | *Bacillus subtilis* | 1yx2A | 29 | 29 | 0.92 | 6.58 |
| 9 | AMT | *dmdA* | *Pelagibacter ubique (HTCC1062)* | 3tfhA | 36 | 36 | 0.97 | 7.67 |
| 10 | AMT | *dmdA* | *Pelagibacter ubique (HTCC1062)* | 3tfhA | 36 | 36 | 0.97 | 6.70 |

^a^Iden1 is the percentage sequence identity of the templates in the threading aligned region with the query sequence

^b^Iden2 is the percentage sequence identity of the whole template chains with query sequence

^c^Cov-Represents the coverage of the threading alignment and is equal to the number of aligned residues divided by the length of the query protein

^d^Norm. Z-score is the normalized Z-score of the threading alignments. Alignment with a normalized Z-score>1 means a good alignment and vice versa.

^e^Aminomethyltransferase

^f^DmdA DMSP-dependent demethylase

^g^Glycine cleavage system T protein

^h^Dimethylglycine dehydrogenase complexed with tetrahydrofolate

^i^Dimethylglycine oxidase

**Predicted model: C-sore= 1.65; TM-score=0.95 dev= 0.05**

| **Rank** | **Class** | **Gene name** | **Organism** | **PDB ID** | **TM-score^a^** | **RMSD^b^** | **IDEN^c^** | **Cov^d^** |
| --- | --- | --- | --- | --- | --- | --- | --- | --- |
| 1 | AMT | *dmdA* | *Pelagibacter ubique (HTCC1062)* | 3tfhA | 0.960 | 0.68 | 0.363 | 0.971 |
| 2 | Oxidoreductase | *dmgdH* | *Rattus norvegicus* | 4p9sA | 0.931 | 1.84 | 0.239 | 0.982 |
| 3 | Oxidoreductase | *dmg* | *Arthrobacter globiformis* | 1pj6A | 0.916 | 2.220 | 0.260 | 0.982 |
| 4 | Oxidoreductase | *soxA^e^* | *Stenotrophomonas maltophilia* | 2gagA | 0.899 | 2.31 | 0.229 | 0.971 |
| 5 | AMT | *gcvT* | *Pyrococcus horikoshii ATCC700860* | 1v5vA | 0.896 | 2.09 | 0.277 | 0.958 |
| 6 | Oxidoreductase | *soxA* | *Corynebacterium sp.* | 1vrqA | 0.896 | 2.28 | 0.230 | 0.968 |
| 7 | AMT | *gcvT* | *Thermotoga maritima* | 1wooA | 0.888 | 1.68 | 0.268 | 0.932 |
| 8 | AMT | *AMT* | *Homo sapiens* | 1wsrA | 0.883 | 2.05 | 0.240 | 0.945 |
| 9 | AMT | *gcvT* | *Bartonella henselae* | 3girA | 0.874 | 1.89 | 0.224 | 0.929 |
| 10 | AMT | *gcvT* | *Bacillus subtilis* | 1yx2B | 0.861 | 2.07 | 0.291 | 0.924 |

^ab^It is a standard for measuring structural similarity between two structures

^c^It is the percentage sequence identity in the structurally aligned region

^d^It represents the coverage of the alignment by TM-align and is equal to the number of structurally aligned residues divided by length of the query protein

^e^Heterotetrameric sarcosine oxidase alpha-subunit

**Table 12**. Up: Top ten analogs identified by LOMETS for threading alignments. Case of ADE39159.1. In the middle: Model predicted by I-TASSER. Down: Top ten identified structural analogs in PDB by TM-align

| **Rank** | **Class** | **Gene name** | **Organism** | **PDB ID** | **Iden1 (%)^a^** | **Iden2 (%)^b^** | **Cov^c^** | **Norm. Z-score^d^** |
| --- | --- | --- | --- | --- | --- | --- | --- | --- |
| 1 | AMT^e^ | *dmdA^f^* | *Pelagibacter ubique (HTCC1062)* | 3tfhA | 34 | 34 | 0.96 | 4.55 |
| 2 | AMT | *dmdA* | *Pelagibacter ubique (HTCC1062)* | 3tfhA | 34 | 34 | 0.96 | 5.36 |
| 3 | AMT | *dmdA* | *Pelagibacter ubique (HTCC1062)* | 3tfhA | 34 | 34 | 0.96 | 5.57 |
| 4 | *Oxidoreductase* | *dmg^i^* | *Arthrobacter globiformis* | 1pj6 | 26 | 32 | 0.97 | 3.24 |
| 5 | Oxidoreductase | *dmgdH*^h^ | *Rattus norvegicus* | 4p9sA | 22 | 26 | 0.97 | 2.41 |
| 6 | AMT | *dmdA* | *Pelagibacter ubique (HTCC1062)* | 3tfhA | 34 | 34 | 0.95 | 5.56 |
| 7 | *Oxidoreductase* | *dmg* | *Arthrobacter globiformis* | 1pj6 | 27 | 32 | 0.96 | 3.60 |
| 8 | AMT | *gcvT^g^* | *Pyrococcus horikoshii* | 1v5vA | 26 | 29 | 0.95 | 6.32 |
| 9 | AMT | *dmdA* | *Pelagibacter ubique (HTCC1062)* | 3tfhA | 34 | 34 | 0.96 | 7.48 |
| 10 | AMT | *dmdA* | *Pelagibacter ubique (HTCC1062)* | 3tfhA | 34 | 34 | 0.96 | 6.56 |

^a^Iden1 is the percentage sequence identity of the templates in the threading aligned region with the query sequence

^b^Iden2 is the percentage sequence identity of the whole template chains with query sequence

^c^Cov-Represents the coverage of the threading alignment and is equal to the number of aligned residues divided by the length of the query protein

^d^Norm. Z-score is the normalized Z-score of the threading alignments. Alignment with a normalized Z-score>1 means a good alignment and vice versa.

^e^Aminomethyltransferase

^f^DmdA DMSP-dependent demethylase

^g^Glycine cleavage system T protein

^h^Dimethylglycine dehydrogenase complexed with tetrahydrofolate

^i^Dimethylglycine oxidase

**Predicted model: C-sore= 1.50; TM-score=0.92 dev= 0.06**

| **Rank** | **Class** | **Gene name** | **Organism** | **PDB ID** | **TM-score^a^** | **RMSD^b^** | **IDEN^c^** | **Cov^d^** |
| --- | --- | --- | --- | --- | --- | --- | --- | --- |
| 1 | AMT | *dmdA* | *Pelagibacter ubique (HTCC1062)* | 3tfhA | 0.950 | 0.79 | 0.341 | 0.961 |
| 2 | Oxidoreductase | *dmgdH* | *Rattus norvegicus* | 4p9sA | 0.933 | 1.77 | 0.231 | 0.982 |
| 3 | Oxidoreductase | *dmg* | *Arthrobacter globiformis* | 1pj6A | 0.928 | 1.89 | 0.249 | 0.982 |
| 4 | Oxidoreductase | *soxA^e^* | *Stenotrophomonas maltophilia* | 2gagA | 0.888 | 2.23 | 0.206 | 0.958 |
| 5 | AMT | *gcvT* | *Pyrococcus horikoshii ATCC700860* | 1v5vA | 0.887 | 2.08 | 0.261 | 0.948 |
| 6 | Oxidoreductase | *soxA* | *Corynebacterium sp.* | 1vrqA | 0.886 | 2.22 | 0.222 | 0.956 |
| 7 | AMT | *gcvT* | *Thermotoga maritima* | 1wooA | 0.880 | 1.70 | 0.265 | 0.924 |
| 8 | AMT | *AMT* | *Homo sapiens* | 1wsrA | 0.877 | 1.99 | 0.248 | 0.935 |
| 9 | AMT | *gcvT* | *Bartonella henselae* | 3girA | 0.868 | 1.84 | 0.218 | 0.919 |
| 10 | AMT | *gcvT* | *Bacillus subtilis* | 1yx2B | 0.857 | 1.97 | 0.279 | 0.914 |

^ab^It is a standard for measuring structural similarity between two structures

^c^It is the percentage sequence identity in the structurally aligned region

^d^It represents the coverage of the alignment by TM-align and is equal to the number of structurally aligned residues divided by length of the query protein

^e^Heterotetrameric sarcosine oxidase alpha-subunit

**Table 13**. Up: Top ten analogs identified by LOMETS for threading alignments. Case of AFS47213.1. In the middle: Model predicted by I-TASSER. Down: Top ten identified structural analogs in PDB by TM-align

| **Rank** | **Class** | **Gene name** | **Organism** | **PDB ID** | **Iden1 (%)^a^** | **Iden2 (%)^b^** | **Cov^c^** | **Norm. Z-score^d^** |
| --- | --- | --- | --- | --- | --- | --- | --- | --- |
| 1 | AMT^e^ | *dmdA^f^* | *Pelagibacter ubique (HTCC1062)* | 3tfhA | 38 | 37 | 0.97 | 4.75 |
| 2 | AMT | *dmdA* | *Pelagibacter ubique (HTCC1062)* | 3tfhA | 37 | 37 | 0.97 | 5.39 |
| 3 | AMT | *dmdA* | *Pelagibacter ubique (HTCC1062)* | 3tfhA | 38 | 37 | 0.97 | 5.54 |
| 4 | *AMT* | *dmdA* | *Pelagibacter ubique (HTCC1062)* | 3tfh | 38 | 37 | 0.97 | 3.26 |
| 5 | Oxidoreductase | *dmgdH*^h^ | *Rattus norvegicus* | 4p9sA | 22 | 25 | 0.98 | 2.40 |
| 6 | AMT | *dmdA* | *Pelagibacter ubique (HTCC1062)* | 3tfhA | 38 | 37 | 0.97 | 5.82 |
| 7 | *Oxidoreductase* | *dmgdH* | *Rattus norvegicus* | 4p9s | 23 | 32 | 0.97 | 3.61 |
| 8 | AMT | *gcvT^g^* | *Bacillus subtilis* | 1yx2A | 30 | 31 | 0.93 | 6.83 |
| 9 | AMT | *dmdA* | *Pelagibacter ubique (HTCC1062)* | 3tfhA | 38 | 37 | 0.97 | 7.70 |
| 10 | AMT | *dmdA* | *Pelagibacter ubique (HTCC1062)* | 3tfhA | 38 | 37 | 0.97 | 6.75 |

^a^Iden1 is the percentage sequence identity of the templates in the threading aligned region with the query sequence

^b^Iden2 is the percentage sequence identity of the whole template chains with query sequence

^c^Cov-Represents the coverage of the threading alignment and is equal to the number of aligned residues divided by the length of the query protein

^d^Norm. Z-score is the normalized Z-score of the threading alignments. Alignment with a normalized Z-score>1 means a good alignment and vice versa.

^e^Aminomethyltransferase

^f^DmdA DMSP-dependent demethylase

^g^Glycine cleavage system T protein

^h^Dimethylglycine dehydrogenase complexed with tetrahydrofolate

**Predicted model: C-sore= 1.66; TM-score=0.95 dev= 0.05**

| **Rank** | **Class** | **Gene name** | **Organism** | **PDB ID** | **TM-score^a^** | **RMSD^b^** | **IDEN^c^** | **Cov^d^** |
| --- | --- | --- | --- | --- | --- | --- | --- | --- |
| 1 | AMT | *dmdA* | *Pelagibacter ubique (HTCC1062)* | 3tfhA | 0.966 | 0.67 | 0.374 | 0.974 |
| 2 | Oxidoreductase | *dmgdH* | *Rattus norvegicus* | 4p9sA | 0.928 | 1.83 | 0.231 | 0.982 |
| 3 | Oxidoreductase | *dmg^e^* | *Arthrobacter globiformis* | 1pj6A | 0.905 | 2.34 | 0.247 | 0.982 |
| 4 | AMT | *gcvT* | *Pyrococcus horikoshii ATCC700860* | 1v5vA | 0.891 | 2.11 | 0.275 | 0.958 |
| 5 | Oxidoreductase | *soxA^f^* | *Stenotrophomonas maltophilia* | 2gagA | 0.888 | 2.52 | 0.241 | 0.971 |
| 6 | Oxidoreductase | *soxA* | *Corynebacterium sp.* | 1vrqA | 0.886 | 2.44 | 0.228 | 0.968 |
| 7 | AMT | *gcvT* | *Thermotoga maritima* | 1wooA | 0.884 | 1.85 | 0.273 | 0.937 |
| 8 | AMT | *AMT* | *Homo sapiens* | 1wsrA | 0.876 | 2.19 | 0.248 | 0.947 |
| 9 | AMT | *gcvT* | *Bartonella henselae* | 3girA | 0.873 | 1.95 | 0.221 | 0.931 |
| 10 | AMT | *gcvT* | *Bacillus subtilis* | 1yx2B | 0.858 | 2.15 | 0.308 | 0.926 |

^ab^It is a standard for measuring structural similarity between two structures

^c^It is the percentage sequence identity in the structurally aligned region

^d^It represents the coverage of the alignment by TM-align and is equal to the number of structurally aligned residues divided by length of the query protein

^e^Dimethylglycine oxidase

^f^Heterotetrameric sarcosine oxidase alpha-subunit

**Table 14**. Up: Top ten analogs identified by LOMETS for threading alignments. Case of AFS48354. In the middle: Model predicted by I-TASSER. Down: Top ten identified structural analogs in PDB by TM-align

| **Rank** | **Class** | **Gene name** | **Organism** | **PDB ID** | **Iden1 (%)^a^** | **Iden2 (%)^b^** | **Cov^c^** | **Norm. Z-score^d^** |
| --- | --- | --- | --- | --- | --- | --- | --- | --- |
| 1 | AMT^e^ | *dmdA^f^* | *Pelagibacter ubique (HTCC1062)* | 3tfhA | 40 | 40 | 0.97 | 4.69 |
| 2 | AMT | *dmdA* | *Pelagibacter ubique (HTCC1062)* | 3tfhA | 40 | 40 | 0.97 | 5.33 |
| 3 | AMT | *dmdA* | *Pelagibacter ubique (HTCC1062)* | 3tfhA | 39 | 40 | 0.97 | 5.38 |
| 4 | Oxidoreductase | *dmgdH*^h^ | *Rattus norvegicus* | 4p9s | 24 | 33 | 0.97 | 3.27 |
| 5 | Oxidoreductase | *dmgdH* | *Rattus norvegicus* | 4p9sA | 23 | 27 | 0.98 | 2.40 |
| 6 | AMT | *dmdA* | *Pelagibacter ubique (HTCC1062)* | 3tfhA | 39 | 40 | 0.97 | 5.81 |
| 7 | *Oxidoreductase* | *dmgdH* | *Rattus norvegicus* | 4p9s | 24 | 33 | 0.97 | 3.61 |
| 8 | AMT | *gcvT^g^* | *Bacillus subtilis* | 1yx2A | 29 | 29 | 0.93 | 6.58 |
| 9 | AMT | *dmdA* | *Pelagibacter ubique (HTCC1062)* | 3tfhA | 40 | 40 | 0.97 | 7.47 |
| 10 | AMT | *dmdA* | *Pelagibacter ubique (HTCC1062)* | 3tfhA | 40 | 40 | 0.97 | 6.62 |

^a^Iden1 is the percentage sequence identity of the templates in the threading aligned region with the query sequence

^b^Iden2 is the percentage sequence identity of the whole template chains with query sequence

^c^Cov-Represents the coverage of the threading alignment and is equal to the number of aligned residues divided by the length of the query protein

^d^Norm. Z-score is the normalized Z-score of the threading alignments. Alignment with a normalized Z-score>1 means a good alignment and vice versa.

^e^Aminomethyltransferase

^f^DmdA DMSP-dependent demethylase

^g^Glycine cleavage system T protein

^h^Dimethylglycine dehydrogenase complexed with tetrahydrofolate

**Predicted model: C-sore= 1.60; TM-score=0.94 dev= 0.05**

| **Rank** | **Class** | **Gene name** | **Organism** | **PDB ID** | **TM-score^a^** | **RMSD^b^** | **IDEN^c^** | **Cov^d^** |
| --- | --- | --- | --- | --- | --- | --- | --- | --- |
| 1 | AMT | *dmdA* | *Pelagibacter ubique (HTCC1062)* | 3tfhA | 0.963 | 0.77 | 0.398 | 0.974 |
| 2 | Oxidoreductase | *dmgdH* | *Rattus norvegicus* | 4p9sA | 0.938 | 1.63 | 0.239 | 0.982 |
| 3 | Oxidoreductase | *dmg^e^* | *Arthrobacter globiformis* | 1pj6A | 0.913 | 2.16 | 0.261 | 0.979 |
| 4 | AMT | *gcvT* | *Pyrococcus horikoshii ATCC700860* | 1v5vA | 0.897 | 2.09 | 0.258 | 0.960 |
| 5 | Oxidoreductase | *soxA^f^* | *Stenotrophomonas maltophilia* | 2gagA | 0.890 | 2.43 | 0.231 | 0.968 |
| 6 | Oxidoreductase | *soxA* | *Corynebacterium sp.* | 1vrqA | 0.888 | 2.35 | 0.228 | 0.966 |
| 7 | AMT | *gcvT* | *Thermotoga maritima* | 1wooA | 0.888 | 1.76 | 0.282 | 0.937 |
| 8 | AMT | *AMT* | *Homo sapiens* | 1wsrA | 0.885 | 2.05 | 0.234 | 0.947 |
| 9 | AMT | *gcvT* | *Bartonella henselae* | 3girA | 0.875 | 1.92 | 0.221 | 0.931 |
| 10 | AMT | *gcvT* | *Bacillus subtilis* | 1yx2B | 0.862 | 2.07 | 0.302 | 0.926 |

^ab^It is a standard for measuring structural similarity between two structures

^c^It is the percentage sequence identity in the structurally aligned region

^d^It represents the coverage of the alignment by TM-align and is equal to the number of structurally aligned residues divided by length of the query protein

^e^Dimethylglycine oxidase

^f^Heterotetrameric sarcosine oxidase alpha-subunit

**Table 15**. Up: Top ten analogs identified by LOMETS for threading alignments. Case of WP_065353845. In the middle: Model predicted by I-TASSER. Down: Top ten identified structural analogs in PDB by TM-align

| **Rank** | **Class** | **Gene name** | **Organism** | **PDB ID** | **Iden1 (%)^a^** | **Iden2 (%)^b^** | **Cov^c^** | **Norm. Z-score^d^** |
| --- | --- | --- | --- | --- | --- | --- | --- | --- |
| 1 | AMT^e^ | *dmdA^f^* | *Pelagibacter ubique (HTCC1062)* | 3tfhA | 37 | 38 | 0.97 | 4.72 |
| 2 | AMT | *dmdA* | *Pelagibacter ubique (HTCC1062)* | 3tfhA | 36 | 38 | 0.97 | 5.36 |
| 3 | AMT | *dmdA* | *Pelagibacter ubique (HTCC1062)* | 3tfhA | 36 | 38 | 0.97 | 5.59 |
| 4 Oxidoreductase | Oxidoreductase | *dmgdH* | *Rattus norvegicus* | 4p9s | 23 | 30 | 0.97 | 3.27 |
| 5 | Oxidoreductase | *dmgdH*^h^ | *Rattus norvegicus* | 4p9sA | 22 | 24 | 0.98 | 2.40 |
| 6 | AMT | *dmdA* | *Pelagibacter ubique (HTCC1062)* | 3tfhA | 36 | 38 | 0.97 | 5.82 |
| 7 | *Oxidoreductase* | *dmg^i^* | *Arthrobacter globiformis* | 1pj6 | 26 | 30 | 0.97 | 3.62 |
| 8 | AMT | *gcvT^g^* | *Bacillus subtilis* | 1yx2A | 27 | 27 | 0.93 | 6.58 |
| 9 | AMT | *dmdA* | *Pelagibacter ubique (HTCC1062)* | 3tfhA | 37 | 38 | 0.97 | 7.59 |
| 10 | AMT | *dmdA* | *Pelagibacter ubique (HTCC1062)* | 3tfhA | 37 | 38 | 0.97 | 6.56 |

^a^Iden1 is the percentage sequence identity of the templates in the threading aligned region with the query sequence

^b^Iden2 is the percentage sequence identity of the whole template chains with query sequence

^c^Cov-Represents the coverage of the threading alignment and is equal to the number of aligned residues divided by the length of the query protein

^d^Norm. Z-score is the normalized Z-score of the threading alignments. Alignment with a normalized Z-score>1 means a good alignment and vice versa.

^e^Aminomethyltransferase

^f^DmdA DMSP-dependent demethylase

^g^Glycine cleavage system T protein

^h^Dimethylglycine dehydrogenase complexed with tetrahydrofolate

^i^Dimethylglycine oxidase

**Predicted model: C-sore= 1.56; TM-score=0.93 dev= 0.06**

| **Rank** | **Class** | **Gene name** | **Organism** | **PDB ID** | **TM-score^a^** | **RMSD^b^** | **IDEN^c^** | **Cov^d^** |
| --- | --- | --- | --- | --- | --- | --- | --- | --- |
| 1 | AMT | *dmdA* | *Pelagibacter ubique (HTCC1062)* | 3tfhA | 0.961 | 0.74 | 0.361 | 0.971 |
| 2 | Oxidoreductase | *dmgdH* | *Rattus norvegicus* | 4p9sA | 0.939 | 1.60 | 0.226 | 0.982 |
| 3 | Oxidoreductase | *dmg* | *Arthrobacter globiformis* | 1pj6A | 0.915 | 2.12 | 0.251 | 0.979 |
| 4 | AMT | *gcvT* | *Pyrococcus horikoshii ATCC700860* | 1v5vA | 0.898 | 2.15 | 0.238 | 0.963 |
| 5 | Oxidoreductase | *soxA^e^* | *Stenotrophomonas maltophilia* | 2gagA | 0.895 | 2.34 | 0.220 | 0.968 |
| 6 | Oxidoreductase | *soxA* | *Corynebacterium sp.* | 1vrqA | 0.893 | 2.27 | 0.218 | 0.966 |
| 7 | AMT | *gcvT* | *Thermotoga maritima* | 1wooA | 0.888 | 1.76 | 0.265 | 0.937 |
| 8 | AMT | *AMT* | *Homo sapiens* | 1wsrA | 0.886 | 2.05 | 0.217 | 0.947 |
| 9 | AMT | *gcvT* | *Bartonella henselae* | 3girA | 0.875 | 1.92 | 0.207 | 0.931 |
| 10 | AMT | *gcvT* | *Bacillus subtilis* | 1yx2B | 0.863 | 2.07 | 0.276 | 0.926 |

^ab^It is a standard for measuring structural similarity between two structures

^c^It is the percentage sequence identity in the structurally aligned region

^d^It represents the coverage of the alignment by TM-align and is equal to the number of structurally aligned residues divided by length of the query protein

^e^Heterotetrameric sarcosine oxidase alpha-subunit

**Table 16**. Up: Top ten analogs identified by LOMETS for threading alignments. Case of WP_053820730.1. In the middle: Model predicted by I-TASSER. Down: Top ten identified structural analogs in PDB by TM-align

| **Rank** | **Class** | **Gene name** | **Organism** | **PDB ID** | **Iden1 (%)^a^** | **Iden2 (%)^b^** | **Cov^c^** | **Norm. Z-score^d^** |
| --- | --- | --- | --- | --- | --- | --- | --- | --- |
| 1 | AMT^e^ | *dmdA^f^* | *Pelagibacter ubique (HTCC1062)* | 3tfhA | 37 | 21 | 0.53 | 2.51 |
| 2 | Oxidoreductase | *dmgdH^h^* | *Rattus norvegicus* | 4pabA | 17 | 22 | 0.90 | 4.36 |
| 3 | AMT | *dmdA* | *Pelagibacter ubique (HTCC1062)* | 3tfhA | 36 | 21 | 0.54 | 4.25 |
| 4 | Oxidoreductase | *dmgdH* | *Rattus norvegicus* | 4p9sA | 24 | 17 | 0.54 | 3.30 |
| 5 | Oxidoreductase | *dmg* | *Arthrobacter globiformis* | 1pj6 | 20 | 25 | 0.94 | 2.65 |
| 6 | Oxidoreductase | *dmgdH* | *Rattus norvegicus* | 4p9sA | 17 | 22 | 0.91 | 4.79 |
| 7 | *Oxidoreductase* | *dmg^i^* | *Arthrobacter globiformis* | 1pj6 | 26 | 25 | 0.54 | 3.59 |
| 8 | AMT | *gcvT^g^* | *Bacillus subtilis* | 1yx2A | 27 | 15 | 0.51 | 6.47 |
| 9 | AMT | *dmdA* | *Pelagibacter ubique (HTCC1062)* | 3tfhA | 37 | 21 | 0.53 | 2.78 |
| 10 | AMT | *dmdA* | *Pelagibacter ubique (HTCC1062)* | 3tfhA | 36 | 21 | 0.54 | 5.61 |

^a^Iden1 is the percentage sequence identity of the templates in the threading aligned region with the query sequence

^b^Iden2 is the percentage sequence identity of the whole template chains with query sequence

^c^Cov-Represents the coverage of the threading alignment and is equal to the number of aligned residues divided by the length of the query protein

^d^Norm. Z-score is the normalized Z-score of the threading alignments. Alignment with a normalized Z-score>1 means a good alignment and vice versa.

^e^Aminomethyltransferase

^f^DmdA DMSP-dependent demethylase

^g^Glycine cleavage system T protein

^h^Dimethylglycine dehydrogenase complexed with tetrahydrofolate

^i^Dimethylglycine oxidase

**Predicted model: C-sore= 0.34; TM-score=0.67 dev= 0.13**

| **Rank** | **Class** | **Gene name** | **Organism** | **PDB ID** | **TM-score^a^** | **RMSD^b^** | **IDEN^c^** | **Cov^d^** |
| --- | --- | --- | --- | --- | --- | --- | --- | --- |
| 1 | Oxidoreductase | *dmgdH* | *Rattus norvegicus* | 4p9sA | 0.874 | 2.56 | 0.157 | 0.921 |
| 2 | Oxidoreductase | *dmg* | *Arthrobacter globiformis* | 1pj6A | 0.867 | 2.72 | 0.184 | 0.923 |
| 3 | Oxidoreductase | *soxA^e^* | *Stenotrophomonas maltophilia* | 2gagA | 0.536 | 3.70 | 0.146 | 0.590 |
| 4 | Oxidoreductase | *soxA* | *Corynebacterium sp.* | 1vrqA | 0.534 | 3.65 | 0.141 | 0.587 |
| 5 | AMT | *dmdA* | *Pelagibacter ubique (HTCC1062)* | 3tfhA | 0.520 | 1.57 | 0.360 | 0.534 |
| 6 | AMT | *gcvT* | *Pyrococcus horikoshii* | 1v5vA | 0.510 | 2.05 | 0.238 | 0.531 |
| 7 | AMT | *gcvT* | *Thermotoga maritima* | 1wooA | 0.506 | 1.47 | 0.272 | 0.518 |
| 8 | AMT | *AMT* | *Homo sapiens* | 1wsrA | 0.503 | 1.73 | 0.216 | 0.520 |
| 9 | AMT | *gcvT* | *Bartonella henselae* | 3girA | 0.495 | 1.88 | 0.204 | 0.514 |
| 10 | AMT | *gcvT* | *Bacillus subtilis* | 1yx2B | 0.494 | 1.67 | 0.274 | 0.509 |

^ab^It is a standard for measuring structural similarity between two structures

^c^It is the percentage sequence identity in the structurally aligned region

^d^It represents the coverage of the alignment by TM-align and is equal to the number of structurally aligned residues divided by length of the query protein

^e^Heterotetrameric sarcosine oxidase alpha-subunit

**Table 17**. Up: Top ten analogs identified by LOMETS for threading alignments. Case of CCC39909.1. In the middle: Model predicted by I-TASSER. Down: Top ten identified structural analogs in PDB by TM-align

| **Rank** | **Class** | **Gene name** | **Organism** | **PDB ID** | **Iden1 (%)^a^** | **Iden2 (%)^b^** | **Cov^c^** | **Norm. Z-score^d^** |
| --- | --- | --- | --- | --- | --- | --- | --- | --- |
| 1 | AMT^e^ | *gcvT^g^* | *Thermotoga maritima* | 1wopA | 26 | 25 | 0.78 | 3.50 |
| 2 | AMT | *gcvT* | *Pyrococcus horikoshii* | 1v5vA | 25 | 24 | 0.80 | 4.91 |
| 3 | AMT | *gcvT* | *Thermotoga maritima* | 1worA | 27 | 25 | 0.78 | 4.66 |
| 4 | Oxidoreductase | *dmgdH*^h^ | *Rattus norvegicus* | 4p9s | 17 | 19 | 0.88 | 3.44 |
| 5 | Oxidoreductase | *dmgdH* | *Rattus norvegicus* | 4p9sA | 16 | 19 | 0.83 | 2.44 |
| 6 | AMT | *dmdA^f^* | *Pelagibacter ubique (HTCC1062)* | 3tfhA | 25 | 22 | 0.80 | 4.87 |
| 7 | *Oxidoreductase* | *dmgdH* | *Rattus norvegicus* | 4p9s | 18 | 19 | 0.82 | 3.63 |
| 8 | AMT | *gcvT* | *Bacillus subtilis* | 1yx2A | 26 | 22 | 0.77 | 5.74 |
| 9 | AMT | *gcvT* | *Thermotoga maritima* | 1wopA | 26 | 25 | 0.78 | 5.21 |
| 10 | Oxidoreductase | *dmgdH* | *Rattus norvegicus* | 4p9sA | 17 | 19 | 0.84 | 5.80 |

^a^Iden1 is the percentage sequence identity of the templates in the threading aligned region with the query sequence

^b^Iden2 is the percentage sequence identity of the whole template chains with query sequence

^c^Cov-Represents the coverage of the threading alignment and is equal to the number of aligned residues divided by the length of the query protein

^d^Norm. Z-score is the normalized Z-score of the threading alignments. Alignment with a normalized Z-score>1 means a good alignment and vice versa.

^e^Aminomethyltransferase

^f^DmdA DMSP-dependent demethylase

^g^Glycine cleavage system T protein

^h^Dimethylglycine dehydrogenase complexed with tetrahydrofolate

**Predicted model: C-sore= -0.06; TM-score=0.71 dev= 0.12**

| **Rank** | **Class** | **Gene name** | **Organism** | **PDB ID** | **TM-score^a^** | **RMSD^b^** | **IDEN^c^** | **Cov^d^** |
| --- | --- | --- | --- | --- | --- | --- | --- | --- |
| 1 | Oxidoreductase | *dmgdH* | *Rattus norvegicus* | 4p9sA | 0.865 | 1.32 | 0.152 | 0.889 |
| 2 | Oxidoreductase | *dmg^e^* | *Arthrobacter globiformis* | 1pj6A | 0.838 | 1.93 | 0.193 | 0.882 |
| 3 | Oxidoreductase | *soxA^f^* | *Stenotrophomonas maltophilia* | 2gagA | 0.779 | 2.65 | 0.203 | 0.841 |
| 4 | Oxidoreductase | *soxA* | *Corynebacterium sp.* | 1vrqA | 0.776 | 2.68 | 0.195 | 0.838 |
| 5 | AMT | *gcvT* | *Pyrococcus horikoshii* | 1v5vA | 0.773 | 1.66 | 0.237 | 0.801 |
| 6 | AMT | *gcvT* | *Thermotoga maritima* | 1worA | 0.763 | 1.11 | 0.261 | 0.777 |
| 7 | AMT | *AMT* | *Homo sapiens* | 1wsvB | 0.754 | 1.71 | 0.200 | 0.786 |
| 8 | AMT | *dmdA* | *Pelagibacter ubique (HTCC1062)* | 3tfiA | 0.752 | 2..02 | 0.246 | 0.799 |
| 9 | AMT | *gcvT* | *Bartonella henselae* | 3girA | 0.740 | 1.86 | 0.202 | 0.777 |
| 10 | AMT | *gcvT* | *Escherichia coli BL21* | 3a8iA | 0.739 | 1.82 | 0.203 | 0.773 |

^ab^It is a standard for measuring structural similarity between two structures

^c^It is the percentage sequence identity in the structurally aligned region

^d^It represents the coverage of the alignment by TM-align and is equal to the number of structurally aligned residues divided by length of the query protein

^e^Dimethylglycine oxidase

^f^Heterotetrameric sarcosine oxidase alpha-subunit

**Table 18**. Up: Top ten analogs identified by LOMETS for threading alignments. Case of CAJ51984.2. In the middle: Model predicted by I-TASSER. Down: Top ten identified structural analogs in PDB by TM-align

| **Rank** | **Class** | **Gene name** | **Organism** | **PDB ID** | **Iden1 (%)^a^** | **Iden2 (%)^b^** | **Cov^c^** | **Norm. Z-score^d^** |
| --- | --- | --- | --- | --- | --- | --- | --- | --- |
| 1 | AMT^e^ | *gcvT^g^* | *Thermotoga maritima* | 1wopA | 26 | 25 | 0.78 | 3.57 |
| 2 | AMT | *gcvT* | *Pyrococcus horikoshii* | 1v5vA | 25 | 24 | 0.80 | 4.79 |
| 3 | AMT | *gcvT* | *Thermotoga maritima* | 1worA | 27 | 25 | 0.78 | 4.65 |
| 4 | Oxidoreductase | *dmgdH*^h^ | *Rattus norvegicus* | 4p9s | 17 | 19 | 0.88 | 3.45 |
| 5 | Oxidoreductase | *dmgdH* | *Rattus norvegicus* | 4p9sA | 16 | 19 | 0.83 | 2.43 |
| 6 | AMT | *dmdA^f^* | *Pelagibacter ubique (HTCC1062)* | 3tfhA | 25 | 22 | 0.80 | 4.89 |
| 7 | *Oxidoreductase* | *dmgdH* | *Rattus norvegicus* | 4p9s | 17 | 19 | 0.83 | 3.62 |
| 8 | AMT | *gcvT* | *Bacillus subtilis* | 1yx2A | 26 | 22 | 0.77 | 5.74 |
| 9 | AMT | *gcvT* | *Thermotoga maritima* | 1wopA | 26 | 25 | 0.78 | 5.26 |
| 10 | AMT | *dmdA* | *Pelagibacter ubique (HTCC1062)* | 3tfhA | 27 | 22 | 0.80 | 5.83 |

^a^Iden1 is the percentage sequence identity of the templates in the threading aligned region with the query sequence

^b^Iden2 is the percentage sequence identity of the whole template chains with query sequence

^c^Cov-Represents the coverage of the threading alignment and is equal to the number of aligned residues divided by the length of the query protein

^d^Norm. Z-score is the normalized Z-score of the threading alignments. Alignment with a normalized Z-score>1 means a good alignment and vice versa.

^e^Aminomethyltransferase

^f^DmdA DMSP-dependent demethylase

^g^Glycine cleavage system T protein

^h^Dimethylglycine dehydrogenase complexed with tetrahydrofolate

**Predicted model: C-sore= 0.23; TM-score=0.68 dev= 0.12**

| **Rank** | **Class** | **Gene name** | **Organism** | **PDB ID** | **TM-score^a^** | **RMSD^b^** | **IDEN^c^** | **Cov^d^** |
| --- | --- | --- | --- | --- | --- | --- | --- | --- |
| 1 | Oxidoreductase | *dmgdH* | *Rattus norvegicus* | 4p9sA | 0.855 | 1.70 | 0.140 | 0.889 |
| 2 | Oxidoreductase | *dmg^e^* | *Arthrobacter globiformis* | 1pj7A | 0.825 | 2.29 | 0.193 | 0.882 |
| 3 | Oxidoreductase | *soxA^f^* | *Stenotrophomonas maltophilia* | 2gagA | 0.781 | 2.46 | 0.198 | 0.836 |
| 4 | Oxidoreductase | *soxA* | *Corynebacterium sp.* | 1vrqA | 0.778 | 2.39 | 0.193 | 0.832 |
| 5 | AMT | *gcvT* | *Pyrococcus horikoshii* | 1v5vA | 0.774 | 1.67 | 0.240 | 0.801 |
| 6 | AMT | *gcvT* | *Thermotoga maritima* | 1worA | 0.765 | 1.01 | 0.258 | 0.777 |
| 7 | AMT | *AMT* | *Homo sapiens* | 1wsvB | 0.756 | 1.57 | 0.206 | 0.784 |
| 8 | AMT | *dmdA* | *Pelagibacter ubique (HTCC1062)* | 3tfiA | 0.752 | 2.03 | 0.238 | 0.799 |
| 9 | AMT | *gcvT* | *Bartonella henselae* | 3girA | 0.742 | 1.77 | 0.208 | 0.775 |
| 10 | AMT | *gcvT* | *Escherichia coli BL21* | 3a8iA | 0.739 | 1.81 | 0.209 | 0.773 |

^ab^It is a standard for measuring structural similarity between two structures

^c^It is the percentage sequence identity in the structurally aligned region

^d^It represents the coverage of the alignment by TM-align and is equal to the number of structurally aligned residues divided by length of the query protein

^e^Dimethylglycine oxidase

^f^Heterotetrameric sarcosine oxidase alpha-subunit

**Table 19**. Up: Top ten analogs identified by LOMETS for threading alignments. Case of AEM59334.1. In the middle: Model predicted by I-TASSER. Down: Top ten identified structural analogs in PDB by TM-align

| **Rank** | **Class** | **Gene name** | **Organism** | **PDB ID** | **Iden1 (%)^a^** | **Iden2 (%)^b^** | **Cov^c^** | **Norm. Z-score^d^** |
| --- | --- | --- | --- | --- | --- | --- | --- | --- |
| 1 | AMT^e^ | *dmdA^f^* | *Pelagibacter ubique (HTCC1062)* | 3tfhA | 27 | 18 | 0.59 | 2.15 |
| 2 | AMT | *gcvT^g^* | *Pyrococcus horikoshii* | 1v5vA | 26 | 19 | 0.58 | 4.37 |
| 3 | AMT | *gcvT* | *Thermotoga maritima* | 1worA | 27 | 18 | 0.57 | 3.64 |
| 4 | Oxidoreductase | *dmgdH*^h^ | *Rattus norvegicus* | 4p9s | 17 | 16 | 0.65 | 2.89 |
| 5 | Oxidoreductase | *dmgdH* | *Rattus norvegicus* | 4p9sA | 17 | 15 | 0.60 | 2.26 |
| 6 | AMT | *dmdA* | *Pelagibacter ubique (HTCC1062)* | 3tfhA | 27 | 18 | 0.59 | 3.22 |
| 7 | *Oxidoreductase* | *dmgdH* | *Rattus norvegicus* | 4p9s | 17 | 16 | 0.62 | 3.03 |
| 8 | AMT | *gcvT* | *Bacillus subtilis* | 1yx2A | 24 | 16 | 0.56 | 5.85 |
| 9 | AMT | *dmdA* | *Pelagibacter ubique (HTCC1062)* | 3tfhA | 27 | 18 | 0.59 | 3.32 |
| 10 | AMT | *dmdA* | *Pelagibacter ubique (HTCC1062)* | 3tfhA | 27 | 18 | 0.59 | 4.46 |

^a^Iden1 is the percentage sequence identity of the templates in the threading aligned region with the query sequence

^b^Iden2 is the percentage sequence identity of the whole template chains with query sequence

^c^Cov-Represents the coverage of the threading alignment and is equal to the number of aligned residues divided by the length of the query protein

^d^Norm. Z-score is the normalized Z-score of the threading alignments. Alignment with a normalized Z-score>1 means a good alignment and vice versa.

^e^Aminomethyltransferase

^f^DmdA DMSP-dependent demethylase

^g^Glycine cleavage system T protein

^h^Dimethylglycine dehydrogenase complexed with tetrahydrofolate

**Predicted model: C-sore=-2.53; TM-score=0.42 dev= 0.14**

| **Rank** | **Class** | **Gene name** | **Organism** | **PDB ID** | **TM-score^a^** | **RMSD^b^** | **IDEN^c^** | **Cov^d^** |
| --- | --- | --- | --- | --- | --- | --- | --- | --- |
| 1 | Oxidoreductase | *dmgdH* | *Rattus norvegicus* | 4p9sA | 0.637 | 1.14 | 0.151 | 0.647 |
| 2 | Oxidoreductase | *dmg^e^* | *Arthrobacter globiformis* | 1pj6A | 0.613 | 1.91 | 0.204 | 0.639 |
| 3 | Oxidoreductase | *soxA^f^* | *Stenotrophomonas maltophilia* | 2gagA | 0.562 | 2.99 | 0.192 | 0.610 |
| 4 | AMT | *gcvT* | *Pyrococcus horikoshii* | 1v5vA | 0.562 | 2.01 | 0.249 | 0.588 |
| 5 | Oxidoreductase | *soxA* | *Corynebacterium sp.* | 1vrqA | 0.561 | 3.03 | 0.190 | 0.608 |
| 6 | AMT | *dmdA* | *Pelagibacter ubique (HTCC1062)* | 3tfiA | 0.554 | 2.33 | 0.259 | 0.589 |
| 7 | AMT | *gcvT* | *Thermotoga maritima* | 1wopA | 0.553 | 1.57 | 0.265 | 0.570 |
| 8 | AMT | *AMT* | *Homo sapiens* | 1wsvB | 0.551 | 1.76 | 0.219 | 0.571 |
| 9 | AMT | *gcvT* | *Bacillus subtilis (168)* | 1yx2B | 0.541 | 1.77 | 0.249 | 0.562 |
| 10 | AMT | *gcvT* | *Bartonella henselae* | 3girA | 0.540 | 2.00 | 0.216 | 0.565 |

^ab^It is a standard for measuring structural similarity between two structures

^c^It is the percentage sequence identity in the structurally aligned region

^d^It represents the coverage of the alignment by TM-align and is equal to the number of structurally aligned residues divided by length of the query protein

^e^Dimethylglycine oxidase

^f^Heterotetrameric sarcosine oxidase alpha-subunit

**Table 20**. Up: Top ten analogs identified by LOMETS for threading alignments. Case of AFS48830. In the middle: Model predicted by I-TASSER. Down: Top ten identified structural analogs in PDB by TM-align

| **Rank** | **Class** | **Gene name** | **Organism** | **PDB ID** | **Iden1 (%)^a^** | **Iden2 (%)^b^** | **Cov^c^** | **Norm. Z-score^d^** |
| --- | --- | --- | --- | --- | --- | --- | --- | --- |
| 1 | AMT^e^ | *gcvT* | *Thermotoga maritima* | 1wopA | 29 | 26 | 0.82 | 3.78 |
| 2 | AMT | *gcvT^g^* | *Pyrococcus horikoshii* | 1v5vA | 25 | 24 | 0.84 | 4.85 |
| 3 | AMT | *gcvT* | *Thermotoga maritima* | 1worA | 28 | 26 | 0.82 | 4.85 |
| 4 | Oxidoreductase | *dmgdH*^h^ | *Rattus norvegicus* | 4p9sA | 18 | 19 | 0.86 | 3.31 |
| 5 | Oxidoreductase | *dmgdH* | *Rattus norvegicus* | 4p9sA | 19 | 19 | 0.86 | 2.40 |
| 6 | AMT | *gcvT* | *Thermotoga maritima* | 1wopA | 28 | 26 | 0.82 | 4.97 |
| 7 | *Oxidoreductase* | *dmgdH* | *Rattus norvegicus* | 4p9s | 18 | 20 | 0.87 | 3.67 |
| 8 | AMT | *gcvT* | *Bacillus subtilis* | 1yx2A | 25 | 22 | 0.81 | 6.32 |
| 9 | AMT | *gcvT* | *Thermotoga maritima* | 1wopA | 28 | 26 | 0.82 | 5.48 |
| 10 | AMT | *dmdA^f^* | *Pelagibacter ubique (HTCC1062)* | 3tfhA | 27 | 25 | 0.85 | 6.04 |

^a^Iden1 is the percentage sequence identity of the templates in the threading aligned region with the query sequence

^b^Iden2 is the percentage sequence identity of the whole template chains with query sequence

^c^Cov-Represents the coverage of the threading alignment and is equal to the number of aligned residues divided by the length of the query protein

^d^Norm. Z-score is the normalized Z-score of the threading alignments. Alignment with a normalized Z-score>1 means a good alignment and vice versa.

^e^Aminomethyltransferase

^f^DmdA DMSP-dependent demethylase

^g^Glycine cleavage system T protein

^h^Dimethylglycine dehydrogenase complexed with tetrahydrofolate

**Predicted model: C-sore=0.64; TM-score=0.80 dev= 0.09**

| **Rank** | **Class** | **Gene name** | **Organism** | **PDB ID** | **TM-score^a^** | **RMSD^b^** | **IDEN^c^** | **Cov^d^** |
| --- | --- | --- | --- | --- | --- | --- | --- | --- |
| 1 | Oxidoreductase | *dmgdH* | *Rattus norvegicus* | 4p9sA | 0.894 | 1.17 | 0.184 | 0.915 |
| 2 | Oxidoreductase | *dmg^e^* | *Arthrobacter globiformis* | 1pj6A | 0.858 | 2.06 | 0.221 | 0.908 |
| 3 | AMT | *gcvT* | *Pyrococcus horikoshii* | 1v5vA | 0.813 | 1.70 | 0.249 | 0.843 |
| 4 | AMT | *gcvT* | *Thermotoga maritima* | 1worA | 0.809 | 0.89 | 0.284 | 0.820 |
| 5 | Oxidoreductase | *soxA^f^* | *Stenotrophomonas maltophilia* | 2gagA | 0.804 | 2.47 | 0.207 | 0.864 |
| 6 | Oxidoreductase | *soxA* | *Corynebacterium sp. U-96* | 1vrqA | 0.801 | 2.49 | 0.202 | 0.862 |
| 7 | AMT | *AMT* | *Homo sapiens* | 1wsvB | 0.799 | 1.52 | 0.228 | 0.827 |
| 8 | AMT | *dmdA* | *Pelagibacter ubique (HTCC1062)* | 3tfiA | 0.792 | 1.97 | 0.258 | 0.839 |
| 9 | AMT | *gcvT* | *Bartonella henselae* | 3girA | 0.787 | 1.61 | 0.237 | 0.818 |
| 10 | AMT | *gcvT* | *Escherichia coli K12* | 3a8iA | 0.778 | 1.90 | 0.251 | 0.816 |

^ab^It is a standard for measuring structural similarity between two structures

^c^It is the percentage sequence identity in the structurally aligned region

^d^It represents the coverage of the alignment by TM-align and is equal to the number of structurally aligned residues divided by length of the query protein

^e^Dimethylglycine oxidase

^f^Heterotetrameric sarcosine oxidase alpha-subunit

**Table 21**. Up: Top ten analogs identified by LOMETS for threading alignments. Case of AGI68776.1. In the middle: Model predicted by I-TASSER. Down: Top ten identified structural analogs in PDB by TM-align

| **Rank** | **Class** | **Gene name** | **Organism** | **PDB ID** | **Iden1 (%)^a^** | **Iden2 (%)^b^** | **Cov^c^** | **Norm. Z-score^d^** |
| --- | --- | --- | --- | --- | --- | --- | --- | --- |
| 1 | AMT^e^ | *dmdA^f^* | *Pelagibacter ubique (HTCC1062)* | 3tfhA | 43 | 43 | 1.00 | 4.80 |
| 2 | AMT | *dmdA* | *Pelagibacter ubique (HTCC1062)* | 3tfhA | 43 | 43 | 1.00 | 5.33 |
| 3 | AMT | *dmdA* | *Pelagibacter ubique (HTCC1062)* | 3tfhA | 43 | 43 | 1.00 | 5.72 |
| 4 | AMT | *dmdA* | *Pelagibacter ubique (HTCC1062)* | 3tfh | 44 | 43 | 0.99 | 3.27 |
| 5 | Oxidoreductase | *dmgdH^h^* | *Rattus norvegicus* | 4p9sA | 17 | 21 | 0.98 | 2.42 |
| 6 | AMT | *dmdA* | *Pelagibacter ubique (HTCC1062)* | 3tfhA | 43 | 43 | 1.00 | 5.80 |
| 7 | *Oxidoreductase* | *dmgdH* | *Rattus norvegicus* | 4p9sA | 17 | 21 | 0.98 | 3.59 |
| 8 | AMT | *gcvT^g^* | *Bacillus subtilis* | 1yx2A | 25 | 28 | 0.96 | 6.64 |
| 9 | AMT | *dmdA* | *Pelagibacter ubique (HTCC1062)* | 3tfhA | 43 | 43 | 1.00 | 6.79 |
| 10 | AMT | *dmdA* | *Pelagibacter ubique (HTCC1062)* | 3tfhA | 43 | 43 | 1.00 | 6.84 |

^a^Iden1 is the percentage sequence identity of the templates in the threading aligned region with the query sequence

^b^Iden2 is the percentage sequence identity of the whole template chains with query sequence

^c^Cov-Represents the coverage of the threading alignment and is equal to the number of aligned residues divided by the length of the query protein

^d^Norm. Z-score is the normalized Z-score of the threading alignments. Alignment with a normalized Z-score>1 means a good alignment and vice versa.

^e^Aminomethyltransferase

^f^DmdA DMSP-dependent demethylase

^g^Glycine cleavage system T protein

^h^Dimethylglycine dehydrogenase complexed with tetrahydrofolate

**Predicted model: C-sore=2; TM-score=0.99 dev= 0.04**

| **Rank** | **Class** | **Gene name** | **Organism** | **PDB ID** | **TM-score^a^** | **RMSD^b^** | **IDEN^c^** | **Cov^d^** |
| --- | --- | --- | --- | --- | --- | --- | --- | --- |
| 1 | AMT | *dmdA* | *Pelagibacter ubique (HTCC1062)* | 3tfhA | 0.997 | 0.40 | 0.429 | 1.000 |
| 2 | Oxidoreductase | *dmgdH* | *Rattus norvegicus* | 4p9sA | 0.932 | 1.80 | 0.180 | 0.986 |
| 3 | Oxidoreductase | *dmg^e^* | *Arthrobacter globiformes* | 1pj6A2 | 0.915 | 2.18 | 0.244 | 0.986 |
| 4 | Oxidoreductase | *dmg* | *Arthrobacter globiformes* | 1pj7A | 0.914 | 2.18 | 0.244 | 0.986 |
| 5 | AMT | *gcvT* | *Pyrococcus horikoshii* | 1v5vA | 0.906 | 1.97 | 0.237 | 0.970 |
| 6 | Oxidoreductase | *soxA^f^* | *Stenotrophomonas maltophilia* | 2gagA | 0.900 | 2.29 | 0.194 | 0.975 |
| 7 | Oxidoreductase | *soxA* | *Corynebacterium sp. U-96* | 3ad7A | 0.900 | 2.26 | 0.197 | 0.975 |
| 8 | AMT | *gcvT* | *Thermotoga maritima* | 1wooA | 0.899 | 1.74 | 0.250 | 0.951 |
| 9 | AMT | *AMT* | *Homo sapiens* | 1wsrA | 0.891 | 2.08 | 0.259 | 0.959 |
| 10 | AMT | *gcvT* | *Bartonella henselae* | 3girA | 0.885 | 1.97 | 0.210 | 0.948 |

^ab^It is a standard for measuring structural similarity between two structures

^c^It is the percentage sequence identity in the structurally aligned region

^d^It represents the coverage of the alignment by TM-align and is equal to the number of structurally aligned residues divided by length of the query protein

^e^Dimethylglycine oxidase

^f^Heterotetrameric sarcosine oxidase alpha-subunit

**Table 22.** Up: Top ten analogs identified by LOMETS for threading alignments. Case of AGI72139.1. In the middle: Model predicted by I-TASSER. Down: Top ten identified structural analogs in PDB by TM-align

| **Rank** | **Class** | **Gene name** | **Organism** | **PDB ID** | **Iden1 (%)^a^** | **Iden2 (%)^b^** | **Cov^c^** | **Norm. Z-score^d^** |
| --- | --- | --- | --- | --- | --- | --- | --- | --- |
| 1 | AMT^e^ | *dmdA^f^* | *Pelagibacter ubique (HTCC1062)* | 3tfhA | 43 | 43 | 1.00 | 4.79 |
| 2 | AMT | *dmdA* | *Pelagibacter ubique (HTCC1062)* | 3tfhA | 43 | 43 | 1.00 | 5.36 |
| 3 | AMT | *dmdA* | *Pelagibacter ubique (HTCC1062)* | 3tfhA | 43 | 43 | 1.00 | 5.77 |
| 4 | AMT | *dmdA* | *Pelagibacter ubique (HTCC1062)* | 3tfh | 44 | 43 | 0.99 | 3.26 |
| 5 | Oxidoreductase | *dmgdH^h^* | *Rattus norvegicus* | 4p9sA | 17 | 21 | 0.98 | 2.42 |
| 6 | AMT | *dmdA* | *Pelagibacter ubique (HTCC1062)* | 3thA | 43 | 43 | 1.00 | 5.77 |
| 7 | *Oxidoreductase* | *dmgdH* | *Rattus norvegicus* | 4p9sA | 17 | 21 | 0.97 | 3.57 |
| 8 | AMT | *gcvT^g^* | *Bacillus subtilis* | 1yx2A | 26 | 28 | 0.96 | 6.72 |
| 9 | AMT | *dmdA* | *Pelagibacter ubique (HTCC1062)* | 3tfhA | 43 | 43 | 1.00 | 6.75 |
| 10 | AMT | *dmdA* | *Pelagibacter ubique (HTCC1062)* | 3tfhA | 44 | 43 | 1.00 | 6.85 |

^a^Iden1 is the percentage sequence identity of the templates in the threading aligned region with the query sequence

^b^Iden2 is the percentage sequence identity of the whole template chains with query sequence

^c^Cov-Represents the coverage of the threading alignment and is equal to the number of aligned residues divided by the length of the query protein

^d^Norm. Z-score is the normalized Z-score of the threading alignments. Alignment with a normalized Z-score>1 means a good alignment and vice versa.

^e^Aminomethyltransferase

^f^DmdA DMSP-dependent demethylase

^g^Glycine cleavage system T protein

^h^Dimethylglycine dehydrogenase complexed with tetrahydrofolate

**Predicted model: C-sore=2; TM-score=0.99 dev= 0.04**

| **Rank** | **Class** | **Gene name** | **Organism** | **PDB ID** | **TM-score^a^** | **RMSD^b^** | **IDEN^c^** | **Cov^d^** |
| --- | --- | --- | --- | --- | --- | --- | --- | --- |
| 1 | AMT | *dmdA* | *Pelagibacter ubique (HTCC1062)* | 3tfhA | 0.997 | 0.39 | 0.432 | 1.000 |
| 2 | Oxidoreductase | *dmgdH* | *Rattus norvegicus* | 4p9sA | 0.932 | 1.79 | 0.183 | 0.986 |
| 3 | Oxidoreductase | *dmg^e^* | *Arthrobacter globiformes* | 1pj6A2 | 0.915 | 2.18 | 0.252 | 0.986 |
| 4 | Oxidoreductase | *dmg* | *Arthrobacter globiformes* | 1pj7A | 0.914 | 2.18 | 0.252 | 0.986 |
| 5 | AMT | *gcvT* | *Pyrococcus horikoshii* | 1v5vA | 0.906 | 1.97 | 0.248 | 0.970 |
| 6 | Oxidoreductase | *soxA^f^* | *Stenotrophomonas maltophilia* | 2gagA | 0.899 | 2.30 | 0.199 | 0.975 |
| 7 | AMT | *gcvT* | *Thermotoga maritima* | 1wooA | 0.899 | 1.75 | 0.253 | 0.951 |
| 8 | Oxidoreductase | *soxA* | *Corynebacterium sp. U-96* | 3ad7A | 0.899 | 2.28 | 0.202 | 0.975 |
| 9 | AMT | *AMT* | *Homo sapiens* | 1wsrA | 0.891 | 2.08 | 0.265 | 0.959 |
| 10 | AMT | *gcvT* | *Bartonella henselae* | 3girA | 0.885 | 1.99 | 0.210 | 0.948 |

^ab^It is a standard for measuring structural similarity between two structures

^c^It is the percentage sequence identity in the structurally aligned region

^d^It represents the coverage of the alignment by TM-align and is equal to the number of structurally aligned residues divided by length of the query protein

^e^Dimethylglycine oxidase

^f^Heterotetrameric sarcosine oxidase alpha-subunit

**Table 23.** Up: Top ten analogs identified by LOMETS for threading alignments. Case of AGM10509.1 In the middle: Model predicted by I-TASSER. Down: Top ten identified structural analogs in PDB by TM-align

| **Rank** | **Class** | **Gene name** | **Organism** | **PDB ID** | **Iden1 (%)^a^** | **Iden2 (%)^b^** | **Cov^c^** | **Norm. Z-score^d^** |
| --- | --- | --- | --- | --- | --- | --- | --- | --- |
| 1 | AMT^e^ | *gcvT* | *Thermotoga maritima* | 1wopA | 30 | 27 | 0.79 | 3.83 |
| 2 | AMT | *gcvT* | *Pyrococcus horikoshii* | 1v5vA | 26 | 26 | 0.81 | 4.85 |
| 3 | AMT | *gcvT* | *Thermotoga maritima* | 1worA | 29 | 27 | 0.79 | 4.59 |
| 4 | Oxidoreductase | *dmgdH* | *Rattus norvegicus* | 4p9s | 18 | 21 | 0.92 | 3.43 |
| 5 | Oxidoreductase | *dmgdH^h^* | *Rattus norvegicus* | 4p9sA | 19 | 20 | 0.83 | 2.41 |
| 6 | AMT | *gcvT* | *Thermotoga maritima* | 1wopA | 30 | 27 | 0.79 | 4.95 |
| 7 | *Oxidoreductase* | *dmgdH* | *Rattus norvegicus* | 4p9s | 17 | 21 | 0.92 | 3.90 |
| 8 | AMT | *gcvT^g^* | *Bacillus subtilis* | 1yx2A | 25 | 21 | 0.79 | 6.29 |
| 9 | AMT | *dmdA^f^* | *Pelagibacter ubique (HTCC1062)* | 3tfhA | 28 | 24 | 0.82 | 6.09 |
| 10 | AMT | *dmdA* | *Pelagibacter ubique (HTCC1062)* | 3tfhA | 27 | 24 | 0.82 | 5.93 |

^a^Iden1 is the percentage sequence identity of the templates in the threading aligned region with the query sequence

^b^Iden2 is the percentage sequence identity of the whole template chains with query sequence

^c^Cov-Represents the coverage of the threading alignment and is equal to the number of aligned residues divided by the length of the query protein

^d^Norm. Z-score is the normalized Z-score of the threading alignments. Alignment with a normalized Z-score>1 means a good alignment and vice versa.

^e^Aminomethyltransferase

^f^DmdA DMSP-dependent demethylase

^g^Glycine cleavage system T protein

^h^Dimethylglycine dehydrogenase complexed with tetrahydrofolate

**Predicted model: C-sore=0.55; TM-score=0.79 dev= 0.09**

| **Rank** | **Class** | **Gene name** | **Organism** | **PDB ID** | **TM-score^a^** | **RMSD^b^** | **IDEN^c^** | **Cov^d^** |
| --- | --- | --- | --- | --- | --- | --- | --- | --- |
| 1 | Oxidoreductase | *dmgdH* | *Rattus norvegicus* | 4p9sA | 0.887 | 0.40 | 0.186 | 0.913 |
| 2 | Oxidoreductase | *dmg^e^* | *Arthrobacter globiformis* | 1pj6A | 0.855 | 2.07 | 0.214 | 0.906 |
| 3 | AMT | *gcvT* | *Pyrococcus horikoshii* | 1v5vA | 0.788 | 1.55 | 0.260 | 0.815 |
| 4 | AMT | *gcvT* | *Thermotoga maritima* | 1worA | 0.784 | 0.91 | 0.301 | 0.795 |
| 5 | Oxidoreductase | *soxA^f^* | *Stenotrophomonas maltophilia* | 2gagA | 0.783 | 2.52 | 0.204 | 0.842 |
| 6 | Oxidoreductase | *soxA* | *Corynebacterium sp. U-96* | 1vrqA | 0.780 | 2.46 | 0.200 | 0.837 |
| 7 | AMT | *AMT(GCST)* | *Homo sapiens* | 1wsvB | 0.769 | 1.56 | 0.221 | 0.797 |
| 8 | AMT | *dmdA* | *Pelagibacter ubique HTCC1062* | 3tfiA | 0.768 | 2.02 | 0.257 | 0.817 |
| 9 | AMT | *gcvT* | *Bartonella henselae* | 3girA | 0.757 | 1.66 | 0.241 | 0.788 |
| 10 | AMT | *gcvT* | *Bacillus subtilis 168* | 1yx2B | 0.753 | 1.58 | 0.246 | 0.781 |

^ab^It is a standard for measuring structural similarity between two structures

^c^It is the percentage sequence identity in the structurally aligned region

^d^It represents the coverage of the alignment by TM-align and is equal to the number of structurally aligned residues divided by length of the query protein

^e^Dimethylglycine oxidase

^f^Heterotetrameric sarcosine oxidase alpha-subunit

**Table 24**. Up: Top ten analogs identified by LOMETS for threading alignments. Case of AHD01041.1. In the middle: Model predicted by I-TASSER. Down: Top ten identified structural analogs in PDB by TM-align

| **Rank** | **Class** | **Gene name** | **Organism** | **PDB ID** | **Iden1 (%)^a^** | **Iden2 (%)^b^** | **Cov^c^** | **Norm. Z-score^d^** |
| --- | --- | --- | --- | --- | --- | --- | --- | --- |
| 1 | AMT^e^ | *dmdA^f^* | *Pelagibacter ubique (HTCC1062)* | 3tfhA | 42 | 42 | 1.00 | 4.76 |
| 2 | AMT | *dmdA* | *Pelagibacter ubique (HTCC1062)* | 3tfhA | 41 | 42 | 1.00 | 5.33 |
| 3 | AMT | *dmdA* | *Pelagibacter ubique (HTCC1062)* | 3tfhA | 42 | 42 | 1.00 | 5.73 |
| 4 | Oxidoreductase | *dmgdH* | *Rattus norvegicus* | 4p9sA | 20 | 22 | 0.98 | 3.27 |
| 5 | Oxidoreductase | *dmgdH^h^* | *Rattus norvegicus* | 4p9sA | 18 | 22 | 0.98 | 2.42 |
| 6 | AMT | *gcvT* | *Pelagibacter ubique (HTCC1062)* | 3tfhA | 41 | 42 | 1.00 | 5.67 |
| 7 | *Oxidoreductase* | *dmgdH* | *Rattus norvegicus* | 4p9sA | 18 | 22 | 0.98 | 3.60 |
| 8 | AMT | *gcvT^g^* | *Bacillus subtilis* | 1yx2A | 24 | 25 | 0.96 | 6.32 |
| 9 | AMT | *dmdA* | *Pelagibacter ubique (HTCC1062)* | 3tfhA | 42 | 42 | 1.00 | 7.74 |
| 10 | AMT | *dmdA* | *Pelagibacter ubique (HTCC1062)* | 3tfhA | 41 | 42 | 1.00 | 6.82 |

^a^Iden1 is the percentage sequence identity of the templates in the threading aligned region with the query sequence

^b^Iden2 is the percentage sequence identity of the whole template chains with query sequence

^c^Cov-Represents the coverage of the threading alignment and is equal to the number of aligned residues divided by the length of the query protein

^d^Norm. Z-score is the normalized Z-score of the threading alignments. Alignment with a normalized Z-score>1 means a good alignment and vice versa.

^e^Aminomethyltransferase

^f^DmdA DMSP-dependent demethylase

^g^Glycine cleavage system T protein

^h^Dimethylglycine dehydrogenase complexed with tetrahydrofolate

**Predicted model: C-sore=1.69; TM-score=0.95 dev= 0.05**

| **Rank** | **Class** | **Gene name** | **Organism** | **PDB ID** | **TM-score^a^** | **RMSD^b^** | **IDEN^c^** | **Cov^d^** |
| --- | --- | --- | --- | --- | --- | --- | --- | --- |
| 1 | Oxidoreductase | *dmdA* | *Pelagibacter ubique HTCC1062* | 3tfhA | 0.990 | 0.71 | 0.412 | 1.000 |
| 2 | Oxidoreductase | *dmg^e^* | *Arthrobacter globiformis* | 1pj6A | 0.943 | 1.59 | 0.189 | 0.986 |
| 3 | AMT | *gcvT* | *Pyrococcus horikoshii* | 1v5vA | 0.927 | 2.01 | 0.253 | 0.989 |
| 4 | AMT | *gcvT* | *Thermotoga maritima* | 1worA | 0.926 | 2.02 | 0.253 | 0.989 |
| 5 | Oxidoreductase | *soxA^f^* | *Stenotrophomonas maltophilia* | 2gagA | 0.916 | 1.89 | 0.228 | 0.975 |
| 6 | Oxidoreductase | *soxA* | *Corynebacterium sp. U-96* | 1vrqA | 0.910 | 1.69 | 0.235 | 0.959 |
| 7 | AMT | *AMT(GCST)* | *Homo sapiens* | 1wsvB | 0.909 | 2.22 | 0.199 | 0.981 |
| 8 | AMT | *dmdA* | *Pelagibacter ubique HTCC1062* | 3tfiA | 0.908 | 2.19 | 0.191 | 0.981 |
| 9 | AMT | *gcvT* | *Bartonella henselae* | 3girA | 0.901 | 1.87 | 0.244 | 0.959 |
| 10 | AMT | *gcvT* | *Bacillus subtilis 168* | 1yx2B | 0.891 | 1.82 | 0.230 | 0.945 |

^ab^It is a standard for measuring structural similarity between two structures

^c^It is the percentage sequence identity in the structurally aligned region

^d^It represents the coverage of the alignment by TM-align and is equal to the number of structurally aligned residues divided by length of the query protein

^e^Dimethylglycine oxidase

^f^Heterotetrameric sarcosine oxidase alpha-subunit

**Table 25.** Up: Top ten analogs identified by LOMETS for threading alignments. Case of AHI32422.1. In the middle: Model predicted by I-TASSER. Down: Top ten identified structural analogs in PDB by TM-align

| **Rank** | **Class** | **Gene name** | **Organism** | **PDB ID** | **Iden1 (%)^a^** | **Iden2 (%)^b^** | **Cov^c^** | **Norm. Z-score^d^** |
| --- | --- | --- | --- | --- | --- | --- | --- | --- |
| 1 | AMT^e^ | *gcvT* | *Thermotoga maritima* | 1wopA | 29 | 25 | 0.79 | 3.70 |
| 2 | AMT | *gcvT* | *Pyrococcus horikoshii* | 1v5vA | 24 | 24 | 0.81 | 4.82 |
| 3 | AMT | *gcvT* | *Thermotoga maritima* | 1worA | 29 | 25 | 0.79 | 4.59 |
| 4 | Oxidoreductase | *dmgdH* | *Rattus norvegicus* | 4p9s | 17 | 19 | 0.92 | 3.51 |
| 5 | Oxidoreductase | *dmgdH^h^* | *Rattus norvegicus* | 4p9sA | 19 | 19 | 0.82 | 2.42 |
| 6 | AMT | *gcvT* | *Thermotoga maritima* | 1wopA | 29 | 25 | 0.79 | 4.90 |
| 7 | *Oxidoreductase* | *dmgdH* | *Rattus norvegicus* | 4p9s | 18 | 19 | 0.83 | 3.64 |
| 8 | AMT | *gcvT^g^* | *Bacillus subtilis* | 1yx2A | 25 | 20 | 0.78 | 6.21 |
| 9 | AMT | *dmdA^f^* | *Pelagibacter ubique (HTCC1062)* | 3tfhA | 28 | 24 | 0.81 | 6.13 |
| 10 | AMT | *dmdA* | *Pelagibacter ubique (HTCC1062)* | 3tfhA | 28 | 24 | 0.81 | 5.96 |

^a^Iden1 is the percentage sequence identity of the templates in the threading aligned region with the query sequence

^b^Iden2 is the percentage sequence identity of the whole template chains with query sequence

^c^Cov-Represents the coverage of the threading alignment and is equal to the number of aligned residues divided by the length of the query protein

^d^Norm. Z-score is the normalized Z-score of the threading alignments. Alignment with a normalized Z-score>1 means a good alignment and vice versa.

^e^Aminomethyltransferase

^f^DmdA DMSP-dependent demethylase

^g^Glycine cleavage system T protein

^h^Dimethylglycine dehydrogenase complexed with tetrahydrofolate

**Predicted model: C-sore=0.61; TM-score=0.80 dev= 0.09**

| **Rank** | **Class** | **Gene name** | **Organism** | **PDB ID** | **TM-score^a^** | **RMSD^b^** | **IDEN^c^** | **Cov^d^** |
| --- | --- | --- | --- | --- | --- | --- | --- | --- |
| 1 | Oxidoreductase | *dmgdH* | *Rattus norvegicus* | 4p9sA | 0.896 | 1.16 | 0.176 | 0.914 |
| 2 | Oxidoreductase | *dmg^e^* | *Arthrobacter globiformis* | 1pj6A | 0.860 | 1.97 | 0.207 | 0.907 |
| 3 | AMT | *gcvT* | *Pyrococcus horikoshii* | 1v5vA | 0.776 | 1.78 | 0.238 | 0.808 |
| 4 | AMT | *gcvT* | *Thermotoga maritima* | 1worA | 0.774 | 0.96 | 0.289 | 0.786 |
| 5 | Oxidoreductase | *soxA^f^* | *Stenotrophomonas maltophilia* | 2gagA | 0.774 | 2.71 | 0.211 | 0.837 |
| 6 | Oxidoreductase | *soxA* | *Corynebacterium sp. U-96* | 3ad7A | 0.771 | 2.46 | 0.202 | 0.828 |
| 7 | AMT | *dmdA* | *Pelagibacter ubique HTCC1062* | 3tfiA | 0.758 | 2.09 | 0.262 | 0.808 |
| 8 | AMT | *AMT (GCST)* | *Homo sapiens* | 1wsvB | 0.758 | 1.55 | 0.216 | 0.786 |
| 9 | AMT | *gcvT* | *Bartonella henselae* | 3girA | 0.747 | 1.67 | 0.247 | 0.777 |
| 10 | AMT | *gcvT* | *Bacillus subtilis 168* | 1yx2B | 0.743 | 1.65 | 0.251 | 0.773 |

^ab^It is a standard for measuring structural similarity between two structures

^c^It is the percentage sequence identity in the structurally aligned region

^d^It represents the coverage of the alignment by TM-align and is equal to the number of structurally aligned residues divided by length of the query protein

^e^Dimethylglycine oxidase

^f^Heterotetrameric sarcosine oxidase alpha-subunit

**Table 26.** Up: Top ten analogs identified by LOMETS for threading alignments. Case of AHM03102.1. In the middle: Model predicted by I-TASSER. Down: Top ten identified structural analogs in PDB by TM-align

| **Rank** | **Class** | **Gene name** | **Organism** | **PDB ID** | **Iden1 (%)^a^** | **Iden2 (%)^b^** | **Cov^c^** | **Norm. Z-score^d^** |
| --- | --- | --- | --- | --- | --- | --- | --- | --- |
| 1 | AMT^e^ | *dmdA^f^* | *Pelagibacter ubique (HTCC1062)* | 3tfhA | 37 | 36 | 0.99 | 4.64 |
| 2 | AMT | *dmdA* | *Pelagibacter ubique (HTCC1062)* | 3tfhA | 37 | 36 | 0.99 | 5.36 |
| 3 | AMT | *dmdA* | *Pelagibacter ubique (HTCC1062)* | 3tfhA | 36 | 36 | 0.99 | 5.48 |
| 4 | Oxidoreductase | *dmgdH* | *Rattus norvegicus* | 4p9s | 21 | 24 | 0.97 | 3.25 |
| 5 | Oxidoreductase | *dmgdH^h^* | *Rattus norvegicus* | 4p9sA | 20 | 24 | 0.98 | 2.42 |
| 6 | AMT | *dmdA* | *Pelagibacter ubique (HTCC1062)* | 3tfhA | 36 | 36 | 0.98 | 5.68 |
| 7 | *Oxidoreductase* | *dmgdH* | *Rattus norvegicus* | 4p9s | 19 | 24 | 0.97 | 3.59 |
| 8 | AMT | *gcvT^g^* | *Bacillus subtilis* | 1yx2A | 24 | 26 | 0.95 | 6.66 |
| 9 | AMT | *dmdA* | *Pelagibacter ubique (HTCC1062)* | 3tfhA | 37 | 36 | 0.99 | 7.41 |
| 10 | AMT | *dmdA* | *Pelagibacter ubique (HTCC1062)* | 3tfhA | 36 | 36 | 0.99 | 6.61 |

^a^Iden1 is the percentage sequence identity of the templates in the threading aligned region with the query sequence

^b^Iden2 is the percentage sequence identity of the whole template chains with query sequence

^c^Cov-Represents the coverage of the threading alignment and is equal to the number of aligned residues divided by the length of the query protein

^d^Norm. Z-score is the normalized Z-score of the threading alignments. Alignment with a normalized Z-score>1 means a good alignment and vice versa.

^e^Aminomethyltransferase

^f^DmdA DMSP-dependent demethylase

^g^Glycine cleavage system T protein

^h^Dimethylglycine dehydrogenase complexed with tetrahydrofolate

**Predicted model: C-sore=1.69; TM-score=0.95 dev= 0.05**

| **Rank** | **Class** | **Gene name** | **Organism** | **PDB ID** | **TM-score^a^** | **RMSD^b^** | **IDEN^c^** | **Cov^d^** |
| --- | --- | --- | --- | --- | --- | --- | --- | --- |
| 1 | AMT | *dmdA* | *Pelagibacter ubique HTCC1062* | 3tfiA | 0.981 | 0.63 | 0.367 | 0.989 |
| 2 | Oxidoreductase | *dmgdH* | *Rattus norvegicus* | 4p9sA | 0.931 | 1.65 | 0.209 | 0.976 |
| 3 | Oxidoreductase | *dmg^e^* | *Arthrobacter globiformis* | 1pj6A | 0.912 | 2.10 | 0.259 | 0.976 |
| 4 | AMT | *gcvT* | *Pyrococcus horikoshii* | 1v5vA | 0.906 | 1.94 | 0.226 | 0.965 |
| 5 | Oxidoreductase | *soxA^f^* | *Stenotrophomonas maltophilia* | 2gagA | 0.894 | 2.24 | 0.217 | 0.965 |
| 6 | Oxidoreductase | *soxA* | *Corynebacterium sp. U-96* | 3ad7A | 0.894 | 2.21 | 0.228 | 0.965 |
| 7 | AMT | *gcvT* | *Thermotoga maritima* | 1worA | 0.894 | 1.69 | 0.251 | 0.941 |
| 8 | AMT | *AMT (GCST)* | *Homo sapiens* | 1wsvA | 0.888 | 1.98 | 0.232 | 0.949 |
| 9 | AMT | *gcvT* | *Bartonella henselae* | 3girA | 0.880 | 1.87 | 0.210 | 0.935 |
| 10 | AMT | *gcvT* | *Bacillus subtilis 168* | 1yx2B | 0.875 | 2.09 | 0.243 | 0.941 |

^ab^It is a standard for measuring structural similarity between two structures

^c^It is the percentage sequence identity in the structurally aligned region

^d^It represents the coverage of the alignment by TM-align and is equal to the number of structurally aligned residues divided by length of the query protein

^e^Dimethylglycine oxidase

^f^Heterotetrameric sarcosine oxidase alpha-subunit

**Table 27**. Up: Top ten analogs identified by LOMETS for threading alignments. Case of AHM05061.1. In the middle: Model predicted by I-TASSER. Down: Top ten identified structural analogs in PDB by TM-align

| **Rank** | **Class** | **Gene name** | **Organism** | **PDB ID** | **Iden1 (%)^a^** | **Iden2 (%)^b^** | **Cov^c^** | **Norm. Z-score^d^** |
| --- | --- | --- | --- | --- | --- | --- | --- | --- |
| 1 | AMT^e^ | *dmdA^f^* | *Pelagibacter ubique (HTCC1062)* | 3tfhA | 43 | 43 | 0.99 | 4.59 |
| 2 | AMT | *dmdA* | *Pelagibacter ubique (HTCC1062)* | 3tfhA | 43 | 43 | 0.99 | 5.24 |
| 3 | AMT | *dmdA* | *Pelagibacter ubique (HTCC1062)* | 3tfhA | 42 | 43 | 0.99 | 5.45 |
| 4 | AMT | *dmdA* | *Pelagibacter ubique HTCC1062* | 3tfh | 43 | 43 | 0.99 | 3.29 |
| 5 | Oxidoreductase | *dmgdH^h^* | *Rattus norvegicus* | 4p9sA | 17 | 19 | 0.98 | 2.42 |
| 6 | AMT | *dmdA* | *Pelagibacter ubique (HTCC1062)* | 3tfhA | 42 | 43 | 0.99 | 5.59 |
| 7 | *AMT* | *dmdA* | *Pelagibacter ubique HTCC1062* | 3tfh | 42 | 43 | 0.98 | 3.63 |
| 8 | AMT | *gcvT^g^* | *Bacillus subtilis* | 1yx2A | 27 | 28 | 0.96 | 6.47 |
| 9 | AMT | *dmdA* | *Pelagibacter ubique (HTCC1062)* | 3tfhA | 43 | 43 | 1.00 | 7.59 |
| 10 | AMT | *dmdA* | *Pelagibacter ubique (HTCC1062)* | 3tfhA | 42 | 43 | 1.00 | 6.55 |

^a^Iden1 is the percentage sequence identity of the templates in the threading aligned region with the query sequence

^b^Iden2 is the percentage sequence identity of the whole template chains with query sequence

^c^Cov-Represents the coverage of the threading alignment and is equal to the number of aligned residues divided by the length of the query protein

^d^Norm. Z-score is the normalized Z-score of the threading alignments. Alignment with a normalized Z-score>1 means a good alignment and vice versa.

^e^Aminomethyltransferase

^f^DmdA DMSP-dependent demethylase

^g^Glycine cleavage system T protein

^h^Dimethylglycine dehydrogenase complexed with tetrahydrofolate

**Predicted model: C-sore=2.00; TM-score=0.99 dev= 0.04**

| **Rank** | **Class** | **Gene name** | **Organism** | **PDB ID** | **TM-score^a^** | **RMSD^b^** | **IDEN^c^** | **Cov^d^** |
| --- | --- | --- | --- | --- | --- | --- | --- | --- |
| 1 | AMT | *dmdA* | *Pelagibacter ubique HTCC1062* | 3tfhA | 0.981 | 0.63 | 0.367 | 0.989 |
| 2 | Oxidoreductase | *dmgdH* | *Rattus norvegicus* | 4p9sA | 0.931 | 1.65 | 0.209 | 0.976 |
| 3 | Oxidoreductase | *dmg^e^* | *Arthrobacter globiformis* | 1pj6A2 | 0.912 | 2.10 | 0.259 | 0.976 |
| 4 | Oxidoreductase | *dmg* | *Arthrobacter globiformis* | 1pj7A | 0.906 | 1.94 | 0.226 | 0.965 |
| 5 | AMT | *gcvT* | *Pyrococcus horikoshii* | 1v5vA | 0.894 | 2.24 | 0.217 | 0.965 |
| 6 | Oxidoreductase | *soxA* | *Corynebacterium sp. U-96* | 3ad7A | 0.894 | 2.21 | 0.228 | 0.965 |
| 7 | Oxidoreductase | *soxA^f^* | *Stenotrophomonas maltophilia* | 2gagA | 0.894 | 1.69 | 0.251 | 0.941 |
| 8 | AMT | *gcvT* | *Thermotoga maritima* | 1wooA | 0.888 | 1.98 | 0.232 | 0.949 |
| 9 | AMT | *AMT(GCST)* | *Homo sapiens* | 1wsrA | 0.880 | 1.87 | 0.210 | 0.935 |
| 10 | AMT | *gcvT* | *Bartonella henselae* | 3girA | 0.875 | 2.09 | 0.243 | 0.941 |

^ab^It is a standard for measuring structural similarity between two structures

^c^It is the percentage sequence identity in the structurally aligned region

^d^It represents the coverage of the alignment by TM-align and is equal to the number of structurally aligned residues divided by length of the query protein

^e^Dimethylglycine oxidase

^f^Heterotetrameric sarcosine oxidase alpha-subunit

**Table 28**. Up: Top ten analogs identified by LOMETS for threading alignments. Case of ASJ73090.1. In the middle: Model predicted by I-TASSER. Down: Top ten identified structural analogs in PDB by TM-align

| **Rank** | **Class** | **Gene name** | **Organism** | **PDB ID** | **Iden1 (%)^a^** | **Iden2 (%)^b^** | **Cov^c^** | **Norm. Z-score^d^** |
| --- | --- | --- | --- | --- | --- | --- | --- | --- |
| 1 | AMT^e^ | *dmdA^f^* | *Pelagibacter ubique (HTCC1062)* | 3tfhA | 41 | 40 | 0.96 | 4.69 |
| 2 | AMT | *dmdA* | *Pelagibacter ubique (HTCC1062)* | 3tfhA | 41 | 40 | 0.96 | 5.45 |
| 3 | AMT | *dmdA* | *Pelagibacter ubique (HTCC1062)* | 3tfhA | 41 | 40 | 0.96 | 5.56 |
| 4 | AMT | *dmdA* | *Pelagibacter ubique HTCC1062* | 3tfh | 41 | 40 | 0.95 | 3.24 |
| 5 | Oxidoreductase | *dmgdH^h^* | *Rattus norvegicus* | 4p9sA | 20 | 24 | 0.96 | 2.40 |
| 6 | AMT | *dmdA* | *Pelagibacter ubique (HTCC1062)* | 3tfhA | 41 | 40 | 0.96 | 5.72 |
| 7 | *Oxidoreductase* | *dmgdH* | *Rattus norvegicus* | 4p9s | 19 | 24 | 0.95 | 3.58 |
| 8 | AMT | *gcvT^g^* | *Bacillus subtilis* | 1yx2A | 21 | 23 | 0.92 | 6.47 |
| 9 | AMT | *dmdA* | *Pelagibacter ubique (HTCC1062)* | 3tfhA | 41 | 40 | 0.96 | 7.97 |
| 10 | AMT | *dmdA* | *Pelagibacter ubique (HTCC1062)* | 3tfhA | 41 | 40 | 0.96 | 6.69 |

^a^Iden1 is the percentage sequence identity of the templates in the threading aligned region with the query sequence

^b^Iden2 is the percentage sequence identity of the whole template chains with query sequence

^c^Cov-Represents the coverage of the threading alignment and is equal to the number of aligned residues divided by the length of the query protein

^d^Norm. Z-score is the normalized Z-score of the threading alignments. Alignment with a normalized Z-score>1 means a good alignment and vice versa.

^e^Aminomethyltransferase

^f^DmdA DMSP-dependent demethylase

^g^Glycine cleavage system T protein

^h^Dimethylglycine dehydrogenase complexed with tetrahydrofolate

**Predicted model: C-sore=1.77; TM-score=0.96 dev= 0.05**

| **Rank** | **Class** | **Gene name** | **Organism** | **PDB ID** | **TM-score^a^** | **RMSD^b^** | **IDEN^c^** | **Cov^d^** |
| --- | --- | --- | --- | --- | --- | --- | --- | --- |
| 1 | AMT | *dmdA* | *Pelagibacter ubique HTCC1062* | 3tfhA | 0.951 | 0.51 | 0.409 | 0.956 |
| 2 | Oxidoreductase | *dmgdH* | *Rattus norvegicus* | 4p9sA | 0.914 | 1.90 | 0.199 | 0.969 |
| 3 | Oxidoreductase | *dmg^e^* | *Arthrobacter globiformis* | 1pj6A2 | 0.886 | 2.51 | 0.226 | 0.966 |
| 4 | AMT | *gcvT* | *Pyrococcus horikoshii* | 1v5vA | 0.877 | 2.24 | 0.245 | 0.948 |
| 5 | Oxidoreductase | *soxA^f^* | *Stenotrophomonas maltophilia* | 2gagA | 0.874 | 2.52 | 0.175 | 0.953 |
| 6 | Oxidoreductase | *soxA* | *Corynebacterium sp. U-96* | 3ad7A | 0.873 | 2.49 | 0.183 | 0.953 |
| 7 | AMT | *gcvT* | *Thermotoga maritima* | 1wooA | 0.867 | 1.87 | 0.244 | 0.919 |
| 8 | AMT | *AMT(GCST)* | *Homo sapiens* | 1wsrA | 0.860 | 2.27 | 0.221 | 0.932 |
| 9 | AMT | *gcvT* | *Bartonella henselae* | 3girA | 0.856 | 2.02 | 0.205 | 0.917 |
| 10 | AMT | *gcvT* | *Bacillus subtilis* | 1yx2B | 0.846 | 2.11 | 0.211 | 0.911 |

^ab^It is a standard for measuring structural similarity between two structures

^c^It is the percentage sequence identity in the structurally aligned region

^d^It represents the coverage of the alignment by TM-align and is equal to the number of structurally aligned residues divided by length of the query protein

^e^Dimethylglycine oxidase

^f^Heterotetrameric sarcosine oxidase alpha-subunit

**Table 29**. Up: Top ten analogs identified by LOMETS for threading alignments. Case of CBV41552.1. In the middle: Model predicted by I-TASSER. Down: Top ten identified structural analogs in PDB by TM-align

| **Rank** | **Class** | **Gene name** | **Organism** | **PDB ID** | **Iden1 (%)^a^** | **Iden2 (%)^b^** | **Cov^c^** | **Norm. Z-score^d^** |
| --- | --- | --- | --- | --- | --- | --- | --- | --- |
| 1 | AMT^e^ | *gcvT* | *Thermotoga maritima* | 1wopA | 32 | 26 | 0.75 | 3.52 |
| 2 | AMT | *dmdA^f^* | *Pelagibacter ubique (HTCC1062)* | 3tfhA | 29 | 24 | 0.77 | 4.82 |
| 3 | AMT | *gcvT* | *Thermotoga maritima* | 1worA | 31 | 26 | 0.75 | 4.65 |
| 4 | AMT | *dmgdH* | *Rattus norvegicus* | 4p9s | 19 | 22 | 0.92 | 3.50 |
| 5 | Oxidoreductase | *dmgdH^h^* | *Rattus norvegicus* | 4p9sA | 20 | 20 | 0.80 | 2.42 |
| 6 | AMT | *gcvT* | *Thermotoga maritima* | 1wopA | 32 | 26 | 0.75 | 4.81 |
| 7 | *Oxidoreductase* | *dmgdH* | *Rattus norvegicus* | 4p9s | 18 | 22 | 0.88 | 3.91 |
| 8 | AMT | *gcvT^g^* | *Bacillus subtilis* | 1yx2A | 27 | 21 | 0.74 | 6.40 |
| 9 | AMT | *dmdA* | *Pelagibacter ubique (HTCC1062)* | 3tfhA | 29 | 24 | 0.77 | 5.62 |
| 10 | AMT | *dmdA* | *Pelagibacter ubique (HTCC1062)* | 3tfhA | 29 | 24 | 0.77 | 5.96 |

^a^Iden1 is the percentage sequence identity of the templates in the threading aligned region with the query sequence

^b^Iden2 is the percentage sequence identity of the whole template chains with query sequence

^c^Cov-Represents the coverage of the threading alignment and is equal to the number of aligned residues divided by the length of the query protein

^d^Norm. Z-score is the normalized Z-score of the threading alignments. Alignment with a normalized Z-score>1 means a good alignment and vice versa.

^e^Aminomethyltransferase

^f^DmdA DMSP-dependent demethylase

^g^Glycine cleavage system T protein

^h^Dimethylglycine dehydrogenase complexed with tetrahydrofolate

**Predicted model: C-sore=0.68; TM-score=0.81 dev= 0.09**

| **Rank** | **Class** | **Gene name** | **Organism** | **PDB ID** | **TM-score^a^** | **RMSD^b^** | **IDEN^c^** | **Cov^d^** |
| --- | --- | --- | --- | --- | --- | --- | --- | --- |
| 1 | AMT | *dmgdH* | *Rattus norvegicus* | 4p9sA | 0.906 | 1.31 | 0.184 | 0.928 |
| 2 | Oxidoreductase | *dmg^e^* | *Arthrobacter globiformis* | 1pj6A2 | 0.878 | 2.00 | 0.200 | 0.924 |
| 3 | Oxidoreductase | *soxA^f^* | *Stenotrophomonas maltophilia* | 2gagA | 0.762 | 2.78 | 0.209 | 0.825 |
| 4 | Oxidoreductase | *soxA* | *Corynebacterium sp.* | 3ad8A | 0.760 | 2.88 | 0.202 | 0.825 |
| 5 | AMT | *dmdA* | *Pelagibacter ubique HTCC1062* | 3tfiA | 0.742 | 1.81 | 0.285 | 0.777 |
| 6 | AMT | *gcvT* | *Thermotoga maritima* | 1worA | 0.738 | 1.10 | 0.314 | 0.752 |
| 7 | AMT | *gcvT* | *Pyrococcus horikoshii* | 1v5vA | 0.737 | 1.63 | 0.266 | 0.766 |
| 8 | AMT | *AMT(GCST)* | *Homo sapiens* | 1wsrA | 0.731 | 1.66 | 0.202 | 0.760 |
| 9 | AMT | *gcvT* | *Bartonella henselae* | 3girA | 0.722 | 1.70 | 0.246 | 0.752 |
| 10 | AMT | *gcvT* | *Bacillus subtilis* | 1yx2B | 0.712 | 1.52 | 0.269 | 0.737 |

^ab^It is a standard for measuring structural similarity between two structures

^c^It is the percentage sequence identity in the structurally aligned region

^d^It represents the coverage of the alignment by TM-align and is equal to the number of structurally aligned residues divided by length of the query protein

^e^Dimethylglycine oxidase

^f^Heterotetrameric sarcosine oxidase alpha-subunit

**Table 30**. Up: Top ten analogs identified by LOMETS for threading alignments. Case of WP_047029467. In the middle: Model predicted by I-TASSER. Down: Top ten identified structural analogs in PDB by TM-align

| **Rank** | **Class** | **Gene name** | **Organism** | **PDB ID** | **Iden1 (%)^a^** | **Iden2 (%)^b^** | **Cov^c^** | **Norm. Z-score^d^** |
| --- | --- | --- | --- | --- | --- | --- | --- | --- |
| 1 | AMT^e^ | *dmdA^f^* | *Pelagibacter ubique (HTCC1062)* | 3tfhA | 42 | 42 | 1.00 | 4.90 |
| 2 | AMT | *dmdA* | *Pelagibacter ubique (HTCC1062)* | 3tfhA | 42 | 42 | 1.00 | 5.45 |
| 3 | AMT | *dmdA* | *Pelagibacter ubique (HTCC1062)* | 3tfhA | 42 | 42 | 1.00 | 5.87 |
| 4 | AMT | *dmdA* | *Pelagibacter ubique (HTCC1062)* | 3tfh | 42 | 42 | 0.99 | 3.30 |
| 5 | Oxidoreductase | *dmgdH^h^* | *Rattus norvegicus* | 4p9sA | 19 | 20 | 0.98 | 2.42 |
| 6 | AMT | *dmdA* | *Pelagibacter ubique (HTCC1062)* | 3tfhA | 42 | 42 | 0.99 | 5.76 |
| 7 | *Oxidoreductase* | *dmdA* | *Pelagibacter ubique (HTCC1062)* | 3tfh | 42 | 42 | 0.98 | 3.62 |
| 8 | AMT | *gcvT^g^* | *Bacillus subtilis* | 1yx2A | 25 | 26 | 0.96 | 6.40 |
| 9 | AMT | *dmdA* | *Pelagibacter ubique (HTCC1062)* | 3tfhA | 42 | 42 | 1.00 | 8.06 |
| 10 | AMT | *dmdA* | *Pelagibacter ubique (HTCC1062)* | 3tfhA | 42 | 42 | 1.00 | 6.93 |

^a^Iden1 is the percentage sequence identity of the templates in the threading aligned region with the query sequence

^b^Iden2 is the percentage sequence identity of the whole template chains with query sequence

^c^Cov-Represents the coverage of the threading alignment and is equal to the number of aligned residues divided by the length of the query protein

^d^Norm. Z-score is the normalized Z-score of the threading alignments. Alignment with a normalized Z-score>1 means a good alignment and vice versa.

^e^Aminomethyltransferase

^f^DmdA DMSP-dependent demethylase

^g^Glycine cleavage system T protein

^h^Dimethylglycine dehydrogenase complexed with tetrahydrofolate

**Predicted model: C-sore=2; TM-score=0.99 dev= 0.04**

| **Rank** | **Class** | **Gene name** | **Organism** | **PDB ID** | **TM-score^a^** | **RMSD^b^** | **IDEN^c^** | **Cov^d^** |
| --- | --- | --- | --- | --- | --- | --- | --- | --- |
| 1 | AMT | *dmdA* | *Pelagibacter ubique (HTCC1062)* | 3tfhA | 0.997 | 0.36 | 0.421 | 1.000 |
| 2 | Oxidoreductase | *dmgdH* | *Rattus norvegicus* | 4p9sA | 0.928 | 1.86 | 0.185 | 0.984 |
| 3 | Oxidoreductase | *dmg^e^* | *Arthrobacter globiformis* | 1pj6A | 0.911 | 2.23 | 0.260 | 0.984 |
| 4 | AMT | *gcvT* | *Pyrococcus horikoshii* | 1v5vA | 0.902 | 1.95 | 0.225 | 0.965 |
| 5 | Oxidoreductase | *soxA^f^* | *Stenotrophomonas maltophilia* | 2gagA | 0.897 | 2.31 | 0.209 | 0.973 |
| 6 | Oxidoreductase | *soxA* | *Corynebacterium sp. U-96* | 3ad7A | 0.897 | 2.37 | 0.207 | 0.976 |
| 7 | AMT | *gcvT* | *Thermotoga maritima* | 1wooA | 0.895 | 1.74 | 0.261 | 0.946 |
| 8 | AMT | *AMT(GCST)* | *Homo sapiens* | 1wsrA | 0.889 | 2.08 | 0.239 | 0.957 |
| 9 | AMT | *gcvT* | *Bartonella henselae* | 3girA | 0.883 | 1.94 | 0.222 | 0.943 |
| 10 | AMT | *gcvT* | *Bacillus subtilis* | 1yx2B | 0.877 | 2.10 | 0.253 | 0.946 |

^ab^It is a standard for measuring structural similarity between two structures

^c^It is the percentage sequence identity in the structurally aligned region

^d^It represents the coverage of the alignment by TM-align and is equal to the number of structurally aligned residues divided by length of the query protein

^e^Dimethylglycine oxidase

^f^Heterotetrameric sarcosine oxidase alpha-subunit

**Table 31**. Up: Top ten analogs identified by LOMETS for threading alignments. Case of WP_048536000. In the middle: Model predicted by I-TASSER. Down: Top ten identified structural analogs in PDB by TM-align

| **Rank** | **Class** | **Gene name** | **Organism** | **PDB ID** | **Iden1 (%)^a^** | **Iden2 (%)^b^** | **Cov^c^** | **Norm. Z-score^d^** |
| --- | --- | --- | --- | --- | --- | --- | --- | --- |
| 1 | AMT^e^ | *dmdA^f^* | *Pelagibacter ubique (HTCC1062)* | 3tfhA | 44 | 44 | 1.00 | 4.77 |
| 2 | AMT | *dmdA* | *Pelagibacter ubique (HTCC1062)* | 3tfhA | 44 | 44 | 1.00 | 5.36 |
| 3 | AMT | *dmdA* | *Pelagibacter ubique (HTCC1062)* | 3tfhA | 44 | 44 | 1.00 | 5.80 |
| 4 | AMT | *dmdA* | *Pelagibacter ubique (HTCC1062)* | 3tfh | 44 | 44 | 0.99 | 3.19 |
| 5 | Oxidoreductase | *dmgdH^h^* | *Rattus norvegicus* | 4p9sA | 17 | 20 | 0.97 | 2.35 |
| 6 | AMT^e^ | *dmdA* | *Pelagibacter ubique (HTCC1062)* | 3tfhA | 44 | 44 | 0.99 | 5.70 |
| 7 | *Oxidoreductase* | *dmdA* | *Pelagibacter ubique (HTCC1062)* | 3tfh | 43 | 44 | 0.98 | 3.52 |
| 8 | AMT | *gcvT^g^* | *Bacillus subtilis* | 1yx2A | 25 | 27 | 0.96 | 6.47 |
| 9 | AMT | *dmdA* | *Pelagibacter ubique (HTCC1062)* | 3tfhA | 44 | 44 | 1.00 | 7.58 |
| 10 | AMT | *dmdA* | *Pelagibacter ubique (HTCC1062)* | 3tfhA | 44 | 44 | 1.00 | 6.78 |

^a^Iden1 is the percentage sequence identity of the templates in the threading aligned region with the query sequence

^b^Iden2 is the percentage sequence identity of the whole template chains with query sequence

^c^Cov-Represents the coverage of the threading alignment and is equal to the number of aligned residues divided by the length of the query protein

^d^Norm. Z-score is the normalized Z-score of the threading alignments. Alignment with a normalized Z-score>1 means a good alignment and vice versa.

^e^Aminomethyltransferase

^f^DmdA DMSP-dependent demethylase

^g^Glycine cleavage system T protein

^h^Dimethylglycine dehydrogenase complexed with tetrahydrofolate

**Predicted model: C-sore=2; TM-score=0.99 dev= 0.04**

| **Rank** | **Class** | **Gene name** | **Organism** | **PDB ID** | **TM-score^a^** | **RMSD^b^** | **IDEN^c^** | **Cov^d^** |
| --- | --- | --- | --- | --- | --- | --- | --- | --- |
| 1 | AMT | *dmdA* | *Pelagibacter ubique (HTCC1062)* | 3tfhA | 0.997 | 0.36 | 0.421 | 1.000 |
| 2 | Oxidoreductase | *dmgdH* | *Rattus norvegicus* | 4p9sA | 0.928 | 1.86 | 0.185 | 0.984 |
| 3 | Oxidoreductase | *dmg^e^* | *Arthrobacter globiformis* | 1pj6A2 | 0.911 | 2.23 | 0.260 | 0.984 |
| 4 | Oxidoreductase | *dmg* | *Arthrobacter globiformis* | 1pj7A | 0.902 | 1.95 | 0.225 | 0.965 |
| 5 | AMT | *gcvT* | *Pyrococcus horikoshii* | 1v5vA | 0.897 | 2.31 | 0.209 | 0.973 |
| 6 | Oxidoreductase | *soxA^f^* | *Stenotrophomonas maltophilia* | 2gagA | 0.897 | 2.37 | 0.207 | 0.976 |
| 7 | Oxidoreductase | *soxA* | *Corynebacterium sp. U-96* | 1vrqA | 0.895 | 1.74 | 0.261 | 0.946 |
| 8 | AMT | *gcvT* | *Thermotoga maritima* | 1wooA | 0.889 | 2.08 | 0.239 | 0.957 |
| 9 | AMT | *AMT(GCST)* | *Homo sapiens* | 1wsrA | 0.883 | 1.94 | 0.222 | 0.943 |
| 10 | AMT | *gcvT* | *Bartonella henselae* | 3girA | 0.877 | 2.10 | 0.253 | 0.946 |

^ab^It is a standard for measuring structural similarity between two structures

^c^It is the percentage sequence identity in the structurally aligned region

^d^It represents the coverage of the alignment by TM-align and is equal to the number of structurally aligned residues divided by length of the query protein

^e^Dimethylglycine oxidase

^f^Heterotetrameric sarcosine oxidase alpha-subunit

**Table 32**. Up: Top ten analogs identified by LOMETS for threading alignments. Case of WP_049834197. In the middle: Model predicted by I-TASSER. Down: Top ten identified structural analogs in PDB by TM-align

| **Rank** | **Class** | **Gene name** | **Organism** | **PDB ID** | **Iden1 (%)^a^** | **Iden2 (%)^b^** | **Cov^c^** | **Norm. Z-score^d^** |
| --- | --- | --- | --- | --- | --- | --- | --- | --- |
| 1 | AMT^e^ | *dmdA^f^* | *Pelagibacter ubique (HTCC1062)* | 3tfhA | 44 | 45 | 1.00 | 4.83 |
| 2 | AMT | *dmdA* | *Pelagibacter ubique (HTCC1062)* | 3tfhA | 44 | 45 | 1.00 | 5.42 |
| 3 | AMT | *dmdA* | *Pelagibacter ubique (HTCC1062)* | 3tfhA | 44 | 45 | 1.00 | 5.76 |
| 4 | AMT | *dmdA* | *Pelagibacter ubique (HTCC1062)* | 3tfh | 44 | 45 | 0.99 | 3.20 |
| 5 | Oxidoreductase | *dmgdH^h^* | *Rattus norvegicus* | 4p9sA | 18 | 22 | 0.98 | 2.36 |
| 6 | AMT | *dmdA* | *Pelagibacter ubique (HTCC1062)* | 3tfhA | 44 | 45 | 0.99 | 5.87 |
| 7 | *Oxidoreductase* | *dmgdH* | *Rattus norvegicus* | 4p9sA | 18 | 22 | 0.98 | 3.51 |
| 8 | AMT | *gcvT^g^* | *Bacillus subtilis* | 1yx2A | 25 | 28 | 0.96 | 6.72 |
| 9 | AMT | *dmdA* | *Pelagibacter ubique (HTCC1062)* | 3tfhA | 45 | 45 | 1.00 | 7.85 |
| 10 | AMT | *dmdA* | *Pelagibacter ubique (HTCC1062)* | 3tfhA | 45 | 45 | 1.00 | 6.80 |

^a^Iden1 is the percentage sequence identity of the templates in the threading aligned region with the query sequence

^b^Iden2 is the percentage sequence identity of the whole template chains with query sequence

^c^Cov-Represents the coverage of the threading alignment and is equal to the number of aligned residues divided by the length of the query protein

^d^Norm. Z-score is the normalized Z-score of the threading alignments. Alignment with a normalized Z-score>1 means a good alignment and vice versa.

^e^Aminomethyltransferase

^f^DmdA DMSP-dependent demethylase

^g^Glycine cleavage system T protein

^h^Dimethylglycine dehydrogenase complexed with tetrahydrofolate

**Predicted model: C-sore=2; TM-score=0.99 dev= 0.04**

| **Rank** | **Class** | **Gene name** | **Organism** | **PDB ID** | **TM-score^a^** | **RMSD^b^** | **IDEN^c^** | **Cov^d^** |
| --- | --- | --- | --- | --- | --- | --- | --- | --- |
| 1 | AMT | *dmdA* | *Pelagibacter ubique (HTCC1062)* | 3tfhA | 0.997 | 0.39 | 0.443 | 1.000 |
| 2 | Oxidoreductase | *dmgdH* | *Rattus norvegicus* | 4p9sA | 0.932 | 1.79 | 0.194 | 0.986 |
| 3 | Oxidoreductase | *dmg^e^* | *Arthrobacter globiformis* | 1pj6A2 | 0.915 | 2.18 | 0.258 | 0.986 |
| 4 | Oxidoreductase | *dmg* | *Arthrobacter globiformis* | 1pj7A | 0.914 | 2.18 | 0.258 | 0.986 |
| 5 | AMT | *gcvT* | *Pyrococcus horikoshii* | 1v5vA | 0.906 | 1.97 | 0.251 | 0.970 |
| 6 | AMT | *gcvT* | *Thermotoga maritima* | 1wooA | 0.899 | 1.74 | 0.250 | 0.951 |
| 7 | Oxidoreductase | *soxA^f^* | *Stenotrophomonas maltophilia* | 2gagA | 0.899 | 2.30 | 0.197 | 0.975 |
| 8 | Oxidoreductase | *soxA* | *Corynebacterium sp. U-96* | 3ad7A | 0.899 | 2.36 | 0.202 | 0.978 |
| 9 | AMT | *AMT(GCST)* | *Homo sapiens* | 1wsrA | 0.891 | 2.07 | 0.259 | 0.959 |
| 10 | AMT | *gcvT* | *Bartonella henselae* | 3girA | 0.885 | 1.99 | 0.219 | 0.948 |

^ab^It is a standard for measuring structural similarity between two structures

^c^It is the percentage sequence identity in the structurally aligned region

^d^It represents the coverage of the alignment by TM-align and is equal to the number of structurally aligned residues divided by length of the query protein

^e^Dimethylglycine oxidase

^f^Heterotetrameric sarcosine oxidase alpha-subunit

**Table 33**. Up: Top ten analogs identified by LOMETS for threading alignments. Case of WP_053112834. In the middle: Model predicted by I-TASSER. Down: Top ten identified structural analogs in PDB by TM-align

| **Rank** | **Class** | **Gene name** | **Organism** | **PDB ID** | **Iden1 (%)^a^** | **Iden2 (%)^b^** | **Cov^c^** | **Norm. Z-score^d^** |
| --- | --- | --- | --- | --- | --- | --- | --- | --- |
| 1 | AMT^e^ | *gcvT* | *Thermotoga maritima* | 1wopA | 30 | 26 | 0.79 | 3.70 |
| 2 | AMT | *gcvT* | *Pyrococcus horikoshii* | 1v5vA | 23 | 23 | 0.81 | 4.82 |
| 3 | AMT | *gcvT* | *Thermotoga maritima* | 1worA | 29 | 26 | 0.79 | 4.52 |
| 4 | Oxidoreductase | *dmgdH* | *Rattus norvegicus* | 4p9s | 17 | 20 | 0.91 | 3.45 |
| 5 | Oxidoreductase | *dmgdH^h^* | *Rattus norvegicus* | 4p9sA | 18 | 20 | 0.82 | 2.36 |
| 6 | AMT | *gcvT* | *Thermotoga maritima* | 1wopA | 30 | 26 | 0.79 | 4.90 |
| 7 | *Oxidoreductase* | *dmgdH* | *Rattus norvegicus* | 4p9s | 18 | 20 | 0.85 | 3.59 |
| 8 | AMT | *gcvT^g^* | *Bacillus subtilis* | 1yx2A | 24 | 20 | 0.78 | 6.27 |
| 9 | AMT | *dmdA^f^* | *Pelagibacter ubique (HTCC1062)* | 3tfhA | 26 | 23 | 0.81 | 6.16 |
| 10 | AMT | *dmdA* | *Pelagibacter ubique (HTCC1062)* | 3tfhA | 26 | 23 | 0.81 | 5.91 |

^a^Iden1 is the percentage sequence identity of the templates in the threading aligned region with the query sequence

^b^Iden2 is the percentage sequence identity of the whole template chains with query sequence

^c^Cov-Represents the coverage of the threading alignment and is equal to the number of aligned residues divided by the length of the query protein

^d^Norm. Z-score is the normalized Z-score of the threading alignments. Alignment with a normalized Z-score>1 means a good alignment and vice versa.

^e^Aminomethyltransferase

^f^DmdA DMSP-dependent demethylase

^g^Glycine cleavage system T protein

^h^Dimethylglycine dehydrogenase complexed with tetrahydrofolate

**Predicted model: C-sore=0.56; TM-score=0.79 dev= 0.09**

| **Rank** | **Class** | **Gene name** | **Organism** | **PDB ID** | **TM-score^a^** | **RMSD^b^** | **IDEN^c^** | **Cov^d^** |
| --- | --- | --- | --- | --- | --- | --- | --- | --- |
| 1 | Oxidoreductase | *dmgdH* | *Rattus norvegicus* | 4p9sA | 0.997 | 0.39 | 0.443 | 1.000 |
| 2 | Oxidoreductase | *dmg^e^* | *Arthrobacter globiformis* | 1pj6A | 0.932 | 1.79 | 0.194 | 0.986 |
| 3 | AMT | *gcvT* | *Pyrococcus horikoshii* | 1v5vA | 0.915 | 2.18 | 0.258 | 0.986 |
| 4 | AMT | *gcvT* | *Thermotoga maritima* | 1worA | 0.914 | 2.18 | 0.258 | 0.986 |
| 5 | Oxidoreductase | *soxA^f^* | *Stenotrophomonas maltophilia* | 2gagA | 0.906 | 1.97 | 0.251 | 0.970 |
| 6 | Oxidoreductase | *soxA* | *Corynebacterium sp. U-96* | 1vrqA | 0.899 | 1.74 | 0.250 | 0.951 |
| 7 | AMT | *AMT(GCST)* | *Homo sapiens* | 1wsvB | 0.899 | 2.30 | 0.197 | 0.975 |
| 8 | AMT | *dmdA* | *Pelagibacter ubique (HTCC1062)* | 3tfiA | 0.899 | 2.36 | 0.202 | 0.978 |
| 9 | AMT | *gcvT* | *Bartonella henselae* | 3girA | 0.891 | 2.07 | 0.259 | 0.959 |
| 10 | AMT | *gcvT* | *Bacillus subtilis* | 1yx2B | 0.885 | 1.99 | 0.219 | 0.948 |

^ab^It is a standard for measuring structural similarity between two structures

^c^It is the percentage sequence identity in the structurally aligned region

^d^It represents the coverage of the alignment by TM-align and is equal to the number of structurally aligned residues divided by length of the query protein

^e^Dimethylglycine oxidase

^f^Heterotetrameric sarcosine oxidase alpha-subunit

**Table 34**. Up: Top ten analogs identified by LOMETS for threading alignments. Case of WP_053819980_PS1. In the middle: Model predicted by I-TASSER. Down: Top ten identified structural analogs in PDB by TM-align

| **Rank** | **Class** | **Gene name** | **Organism** | **PDB ID** | **Iden1 (%)^a^** | **Iden2 (%)^b^** | **Cov^c^** | **Norm. Z-score^d^** |
| --- | --- | --- | --- | --- | --- | --- | --- | --- |
| 1 | AMT^e^ | *dmdA^f^* | *Pelagibacter ubique (HTCC1062)* | 3tfhA | 40 | 40 | 0.99 | 4.74 |
| 2 | AMT | *dmdA* | *Pelagibacter ubique (HTCC1062)* | 3tfhA | 40 | 40 | 0.99 | 5.36 |
| 3 | AMT | *dmdA* | *Pelagibacter ubique (HTCC1062)* | 3tfhA | 40 | 40 | 0.99 | 5.62 |
| 4 | AMT | *dmdA* | *Pelagibacter ubique (HTCC1062)* | 3tfh | 40 | 40 | 0.99 | 3.23 |
| 5 | Oxidoreductase | *dmgdH^h^* | *Rattus norvegicus* | 4p9sA | 20 | 24 | 0.98 | 2.35 |
| 6 | AMT | *dmdA* | *Pelagibacter ubique (HTCC1062)* | 3tfhA | 39 | 40 | 0.99 | 5.74 |
| 7 | *AMT* | *dmdA* | *Pelagibacter ubique (HTCC1062)* | 3tfh | 39 | 40 | 0.98 | 3.55 |
| 8 | AMT | *gcvT^g^* | *Bacillus subtilis* | 1yx2A | 24 | 26 | 0.95 | 6.47 |
| 9 | AMT | *dmdA* | *Pelagibacter ubique (HTCC1062)* | 3tfhA | 41 | 40 | 0.99 | 8.01 |
| 10 | AMT | *dmdA* | *Pelagibacter ubique (HTCC1062)* | 3tfhA | 40 | 40 | 0.99 | 6.81 |

^a^Iden1 is the percentage sequence identity of the templates in the threading aligned region with the query sequence

^b^Iden2 is the percentage sequence identity of the whole template chains with query sequence

^c^Cov-Represents the coverage of the threading alignment and is equal to the number of aligned residues divided by the length of the query protein

^d^Norm. Z-score is the normalized Z-score of the threading alignments. Alignment with a normalized Z-score>1 means a good alignment and vice versa.

^e^Aminomethyltransferase

^f^DmdA DMSP-dependent demethylase

^g^Glycine cleavage system T protein

^h^Dimethylglycine dehydrogenase complexed with tetrahydrofolate

**Predicted model: C-sore=1.71; TM-score=0.95 dev= 0.05**

| **Rank** | **Class** | **Gene name** | **Organism** | **PDB ID** | **TM-score^a^** | **RMSD^b^** | **IDEN^c^** | **Cov^d^** |
| --- | --- | --- | --- | --- | --- | --- | --- | --- |
| 1 | AMT | *dmdA* | *Pelagibacter ubique (HTCC1062)* | 3tfhA | 0.988 | 0.60 | 0.404 | 0.995 |
| 2 | Oxidoreductase | *dmgdH* | *Rattus norvegicus* | 4p9sA | 0.924 | 1.89 | 0.214 | 0.981 |
| 3 | Oxidoreductase | *dmg^e^* | *Arthrobacter globiformis* | 1pj6A | 0.907 | 2.26 | 0.247 | 0.981 |
| 4 | AMT | *gcvT* | *Pyrococcus horikoshii* | 1v5vA | 0.894 | 1.97 | 0.254 | 0.957 |
| 5 | Oxidoreductase | *soxA^f^* | *Corynebacterium sp. U-96* | 3ad7A | 0.890 | 2.41 | 0.216 | 0.968 |
| 6 | Oxidoreductase | *soxA* | *Stenotrophomonas maltophilia* | 2gagA | 0.890 | 2.30 | 0.219 | 0.965 |
| 7 | AMT | *gcvT* | *Thermotoga maritima* | 1wooA | 0.887 | 1.83 | 0.246 | 0.941 |
| 8 | AMT | *AMT(GCST)* | *Homo sapiens* | 1wsrA | 0.879 | 2.13 | 0.236 | 0.949 |
| 9 | AMT | *gcvT* | *Bartonella henselae* | 3girA | 0.876 | 1.95 | 0.231 | 0.935 |
| 10 | AMT | *gcvT* | *Bacillus subtilis* | 1yx2B | 0.868 | 2.18 | 0.246 | 0.941 |

^ab^It is a standard for measuring structural similarity between two structures

^c^It is the percentage sequence identity in the structurally aligned region

^d^It represents the coverage of the alignment by TM-align and is equal to the number of structurally aligned residues divided by length of the query protein

^e^Dimethylglycine oxidase

^f^Heterotetrameric sarcosine oxidase alpha-subunit

**Table 35**. Up: Top ten analogs identified by LOMETS for threading alignments. Case of WP_065273401.1. In the middle: Model predicted by I-TASSER. Down: Top ten identified structural analogs in PDB by TM-align

| **Rank** | **Class** | **Gene name** | **Organism** | **PDB ID** | **Iden1 (%)^a^** | **Iden2 (%)^b^** | **Cov^c^** | **Norm. Z-score^d^** |
| --- | --- | --- | --- | --- | --- | --- | --- | --- |
| 1 | AMT^e^ | *dmdA^f^* | *Pelagibacter ubique (HTCC1062)* | 3tfhA | 45 | 44 | 1.00 | 4.79 |
| 2 | AMT | *dmdA* | *Pelagibacter ubique (HTCC1062)* | 3tfhA | 44 | 44 | 1.00 | 5.36 |
| 3 | AMT | *dmdA* | *Pelagibacter ubique (HTCC1062)* | 3tfhA | 44 | 44 | 1.00 | 5.69 |
| 4 | AMT | *dmdA* | *Pelagibacter ubique (HTCC1062)* | 3tfh | 45 | 44 | 0.99 | 3.19 |
| 5 | Oxidoreductase | *dmgdH^h^* | *Rattus norvegicus* | 4p9sA | 17 | 20 | 0.98 | 2.36 |
| 6 | AMT | *dmdA* | *Pelagibacter ubique (HTCC1062)* | 3tfhA | 44 | 44 | 1.00 | 5.81 |
| 7 | *AMT* | *dmdA* | *Pelagibacter ubique (HTCC1062)* | 3tfh | 44 | 44 | 0.98 | 3.53 |
| 8 | AMT | *gcvT^g^* | *Bacillus subtilis* | 1yx2A | 24 | 26 | 0.96 | 6.64 |
| 9 | AMT | *dmdA* | *Pelagibacter ubique (HTCC1062)* | 3tfhA | 44 | 44 | 1.00 | 7.77 |
| 10 | AMT | *dmdA* | *Pelagibacter ubique (HTCC1062)* | 3tfhA | 44 | 44 | 1.00 | 6.76 |

^a^Iden1 is the percentage sequence identity of the templates in the threading aligned region with the query sequence

^b^Iden2 is the percentage sequence identity of the whole template chains with query sequence

^c^Cov-Represents the coverage of the threading alignment and is equal to the number of aligned residues divided by the length of the query protein

^d^Norm. Z-score is the normalized Z-score of the threading alignments. Alignment with a normalized Z-score>1 means a good alignment and vice versa.

^e^Aminomethyltransferase

^f^DmdA DMSP-dependent demethylase

^g^Glycine cleavage system T protein

^h^Dimethylglycine dehydrogenase complexed with tetrahydrofolate

**Predicted model: C-sore=2; TM-score=0.99 dev= 0.04**

| **Rank** | **Class** | **Gene name** | **Organism** | **PDB ID** | **TM-score^a^** | **RMSD^b^** | **IDEN^c^** | **Cov^d^** |
| --- | --- | --- | --- | --- | --- | --- | --- | --- |
| 1 | AMT | *dmdA* | *Pelagibacter ubique (HTCC1062)* | 3tfhA | 0.997 | 0.39 | 0.439 | 1.00 |
| 2 | Oxidoreductase | *dmgdH* | *Rattus norvegicus* | 4p9sA | 0.929 | 1.81 | 0.180 | 0.984 |
| 3 | Oxidoreductase | *dmg^e^* | *Arthrobacter globiformis* | 1pj6A | 0.913 | 2.17 | 0.249 | 0.984 |
| 4 | Oxidoreductase | *dmg* | *Arthrobacter globiformis* | 1pj7A | 0.913 | 2.17 | 0.249 | 0.984 |
| 5 | AMT | *gcvT* | *Pyrococcus horikoshii* | 1v5vA | 0.904 | 1.96 | 0.223 | 0.967 |
| 6 | Oxidoreductase | *soxA^f^* | *Stenotrophomonas maltophilia* | 2gagA | 0.899 | 2.31 | 0.199 | 0.976 |
| 7 | Oxidoreductase | *soxA* | *Corynebacterium sp. U-96* | 3ad7A | 0.899 | 2.28 | 0.182 | 0.976 |
| 8 | AMT | *gcvT* | *Thermotoga maritima* | 1wooA | 0.897 | 1.76 | 0.227 | 0.948 |
| 9 | AMT | *AMT(GCST)* | *Homo sapiens* | 1wsrA | 0.890 | 2.10 | 0.230 | 0.959 |
| 10 | AMT | *gcvT* | *Bartonella henselae* | 3girA | 0.885 | 1.94 | 0.228 | 0.946 |

^ab^It is a standard for measuring structural similarity between two structures

^c^It is the percentage sequence identity in the structurally aligned region

^d^It represents the coverage of the alignment by TM-align and is equal to the number of structurally aligned residues divided by length of the query protein

^e^Dimethylglycine oxidase

^f^Heterotetrameric sarcosine oxidase alpha-subunit

**Table 36**. Up: Top ten analogs identified by LOMETS for threading alignments. Case of WP_071941841.1. In the middle: Model predicted by I-TASSER. Down: Top ten identified structural analogs in PDB by TM-align

| **Rank** | **Class** | **Gene name** | **Organism** | **PDB ID** | **Iden1 (%)^a^** | **Iden2 (%)^b^** | **Cov^c^** | **Norm. Z-score^d^** |
| --- | --- | --- | --- | --- | --- | --- | --- | --- |
| 1 | AMT^e^ | *dmdA^f^* | *Pelagibacter ubique (HTCC1062)* | 3tfhA | 33 | 32 | 0.92 | 4.23 |
| 2 | AMT | *gcvT* | *Pyrococcus horikoshii* | 1v5vA | 27 | 28 | 0.91 | 5.28 |
| 3 | AMT | *dmdA* | *Pelagibacter ubique (HTCC1062)* | 3tfhA | 34 | 32 | 0.92 | 5.23 |
| 4 | Oxidoreductase | *dmgdH* | *Rattus norvegicus* | 4p9s | 24 | 25 | 0.95 | 3.20 |
| 5 | Oxidoreductase | *dmgdH^h^* | *Rattus norvegicus* | 4p9sA | 21 | 25 | 0.93 | 2.34 |
| 6 | AMT | *dmdA* | *Pelagibacter ubique (HTCC1062)* | 3tfhA | 33 | 32 | 0.92 | 5.36 |
| 7 | *Oxidoreductase* | *dmg* | *Arthrobacter globiformis* | 1pj6 | 23 | 27 | 0.94 | 3.53 |
| 8 | AMT | *gcvT^g^* | *Bacillus subtilis* | 1yx2A | 29 | 28 | 0.89 | 6.40 |
| 9 | AMT | *dmdA* | *Pelagibacter ubique (HTCC1062)* | 3tfhA | 34 | 32 | 0.92 | 6.94 |
| 10 | AMT | *dmdA* | *Pelagibacter ubique (HTCC1062)* | 3tfhA | 34 | 32 | 0.92 | 6.35 |

^a^Iden1 is the percentage sequence identity of the templates in the threading aligned region with the query sequence

^b^Iden2 is the percentage sequence identity of the whole template chains with query sequence

^c^Cov-Represents the coverage of the threading alignment and is equal to the number of aligned residues divided by the length of the query protein

^d^Norm. Z-score is the normalized Z-score of the threading alignments. Alignment with a normalized Z-score>1 means a good alignment and vice versa.

^e^Aminomethyltransferase

^f^DmdA DMSP-dependent demethylase

^g^Glycine cleavage system T protein

^h^Dimethylglycine dehydrogenase complexed with tetrahydrofolate

**Predicted model: C-sore=1.11; TM-score=0.87 dev= 0.07**

| **Rank** | **Class** | **Gene name** | **Organism** | **PDB ID** | **TM-score^a^** | **RMSD^b^** | **IDEN^c^** | **Cov^d^** |
| --- | --- | --- | --- | --- | --- | --- | --- | --- |
| 1 | Oxidoreductase | *dmgdH* | *Rattus norvegicus* | 4p9sA | 0.997 | 0.39 | 0.439 | 1.00 |
| 2 | AMT | *dmdA* | *Pelagibacter ubique HTCC1062* | 3tfiA | 0.929 | 1.81 | 0.180 | 0.984 |
| 3 | Oxidoreductase | *dmg^e^* | *Arthrobacter globiformis* | 1pj6A | 0.913 | 2.17 | 0.249 | 0.984 |
| 4 | AMT | *gcvT* | *Pyrococcus horikoshii* | 1v5vA | 0.913 | 2.17 | 0.249 | 0.984 |
| 5 | Oxidoreductase | *soxA^f^* | *Stenotrophomonas maltophilia* | 2gagA | 0.904 | 1.96 | 0.223 | 0.967 |
| 6 | Oxidoreductase | *soxA* | *Corynebacterium sp. U-96* | 1vrqA | 0.899 | 2.31 | 0.199 | 0.976 |
| 7 | AMT | *gcvT* | *Thermotoga maritima* | 1worA | 0.899 | 2.28 | 0.182 | 0.976 |
| 8 | AMT | *AMT(GCST)* | *Homo sapiens* | 1wsrA | 0.897 | 1.76 | 0.227 | 0.948 |
| 9 | AMT | *gcvT* | *Bartonella henselae* | 3girA | 0.890 | 2.10 | 0.230 | 0.959 |
| 10 | AMT | *gcvT* | *Bacillus subtilis* | 1yx2B | 0.885 | 1.94 | 0.228 | 0.946 |

^ab^It is a standard for measuring structural similarity between two structures

^c^It is the percentage sequence identity in the structurally aligned region

^d^It represents the coverage of the alignment by TM-align and is equal to the number of structurally aligned residues divided by length of the query protein

^e^Dimethylglycine oxidase

^f^Heterotetrameric sarcosine oxidase alpha-subunit

**Table 37**. Up: Top ten analogs identified by LOMETS for threading alignments. Case of WP_076627280.1. In the middle: Model predicted by I-TASSER. Down: Top ten identified structural analogs in PDB by TM-align

| **Rank** | **Class** | **Gene name** | **Organism** | **PDB ID** | **Iden1 (%)^a^** | **Iden2 (%)^b^** | **Cov^c^** | **Norm. Z-score^d^** |
| --- | --- | --- | --- | --- | --- | --- | --- | --- |
| 1 | AMT^e^ | *dmdA^f^* | *Pelagibacter ubique (HTCC1062)* | 3tfhA | 42 | 43 | 1.00 | 4.70 |
| 2 | AMT | *dmdA* | *Pelagibacter ubique (HTCC1062)* | 3tfhA | 43 | 43 | 1.00 | 5.33 |
| 3 | AMT | *dmdA* | *Pelagibacter ubique (HTCC1062)* | 3tfhA | 42 | 43 | 1.00 | 5.69 |
| 4 | AMT | *dmdA* | *Pelagibacter ubique (HTCC1062)* | 3tfh | 44 | 43 | 0.99 | 3.19 |
| 5 | Oxidoreductase | *dmgdH^h^* | *Rattus norvegicus* | 4p9sA | 18 | 21 | 0.98 | 2.36 |
| 6 | AMT | *dmdA* | *Pelagibacter ubique (HTCC1062)* | 3tfhA | 41 | 43 | 0.99 | 5.82 |
| 7 | *AMT* | *dmdA* | *Pelagibacter ubique (HTCC1062)* | 3tfh | 41 | 43 | 0.98 | 3.53 |
| 8 | AMT | *gcvT^g^* | *Bacillus subtilis* | 1yx2A | 25 | 27 | 0.96 | 6.61 |
| 9 | AMT | *dmdA* | *Pelagibacter ubique (HTCC1062)* | 3tfhA | 43 | 43 | 1.00 | 7.43 |
| 10 | AMT | *dmdA* | *Pelagibacter ubique (HTCC1062)* | 3tfhA | 43 | 43 | 1.00 | 6.73 |

^a^Iden1 is the percentage sequence identity of the templates in the threading aligned region with the query sequence

^b^Iden2 is the percentage sequence identity of the whole template chains with query sequence

^c^Cov-Represents the coverage of the threading alignment and is equal to the number of aligned residues divided by the length of the query protein

^d^Norm. Z-score is the normalized Z-score of the threading alignments. Alignment with a normalized Z-score>1 means a good alignment and vice versa.

^e^Aminomethyltransferase

^f^DmdA DMSP-dependent demethylase

^g^Glycine cleavage system T protein

^h^Dimethylglycine dehydrogenase complexed with tetrahydrofolate

**Predicted model: C-sore=2; TM-score=0.99 dev= 0.04**

| **Rank** | **Class** | **Gene name** | **Organism** | **PDB ID** | **TM-score^a^** | **RMSD^b^** | **IDEN^c^** | **Cov^d^** |
| --- | --- | --- | --- | --- | --- | --- | --- | --- |
| 1 | AMT | *dmdA* | *Pelagibacter ubique HTCC1062* | 3tfhA | 0.997 | 0.38 | 0.428 | 1.00 |
| 2 | Oxidoreductase | *dmgdh* | *Rattus norvegicus* | 4p9sA | 0.930 | 1.79 | 0.183 | 0.984 |
| 3 | Oxidoreductase | *dmg^e^* | *Arthrobacter globiformis* | 1pj6A2 | 0.913 | 2.16 | 0.255 | 0.984 |
| 4 | Oxidoreductase | *dmg* | *Arthrobacter globiformis* | 1pj7A | 0.913 | 2.17 | 0.255 | 0.984 |
| 5 | AMT | *gcvT* | *Pyrococcus horikoshii* | 1v5vA | 0.905 | 1.94 | 0.223 | 0.967 |
| 6 | Oxidoreductase | *soxA^f^* | *Stenotrophomonas maltophilia* | 2gagA | 0.900 | 2.27 | 0.188 | 0.976 |
| 7 | Oxidoreductase | *soxA* | *Corynebacterium sp. U-96* | 3ad7A | 0.899 | 2.27 | 0.182 | 0.976 |
| 8 | AMT | *gcvT* | *Thermotoga maritima* | 1wooA | 0.897 | 1.74 | 0.239 | 0.946 |
| 9 | AMT | *AMT(GCST)* | *Homo sapiens* | 1wsrA | 0.891 | 2.08 | 0.264 | 0.959 |
| 10 | AMT | *gcvT* | *Bartonella henselae* | 3girA | 0.886 | 1.92 | 0.239 | 0.946 |

^ab^It is a standard for measuring structural similarity between two structures

^c^It is the percentage sequence identity in the structurally aligned region

^d^It represents the coverage of the alignment by TM-align and is equal to the number of structurally aligned residues divided by length of the query protein

^e^Dimethylglycine oxidase

^f^Heterotetrameric sarcosine oxidase alpha-subunit

**Table 38**. Up: Top ten analogs identified by LOMETS for threading alignments. Case of WP_096389816.1. In the middle: Model predicted by I-TASSER. Down: Top ten identified structural analogs in PDB by TM-align

| **Rank** | **Class** | **Gene name** | **Organism** | **PDB ID** | **Iden1 (%)^a^** | **Iden2 (%)^b^** | **Cov^c^** | **Norm. Z-score^d^** |
| --- | --- | --- | --- | --- | --- | --- | --- | --- |
| 1 | AMT^e^ | *gcvT* | *Thermotoga maritima* | 1wopA | 30 | 27 | 0.79 | 3.58 |
| 2 | AMT | *gcvT* | *Pyrococcus horikoshii* | 1v5vA | 27 | 25 | 0.81 | 4.94 |
| 3 | AMT | *gcvT* | *Rhermotoga maritima* | 1worA | 31 | 27 | 0.79 | 4.63 |
| 4 | Oxidoreductase | *dmgdH* | *Rattus norvegicus* | 4p9s | 18 | 20 | 0.89 | 3.35 |
| 5 | Oxidoreductase | *dmgdH^h^* | *Rattus norvegicus* | 4p9sA | 18 | 20 | 0.84 | 2.38 |
| 6 | AMT | *gcvT* | *Thermotoga maritima* | 1wopA | 30 | 27 | 0.79 | 4.90 |
| 7 | *Oxidoreductase* | *dmgdH* | *Rattus norvegicus* | 4p9s | 18 | 20 | 0.84 | 3.57 |
| 8 | AMT | *gcvT^g^* | *Bacillus subtilis* | 1yx2A | 26 | 23 | 0.78 | 5.96 |
| 9 | AMT | *dmdA^f^* | *Pelagibacter ubique (HTCC1062)* | 3tfhA | 27 | 23 | 0.82 | 5.78 |
| 10 | AMT | *dmdA* | *Pelagibacter ubique (HTCC1062)* | 3tfhA | 28 | 23 | 0.82 | 5.76 |

^a^Iden1 is the percentage sequence identity of the templates in the threading aligned region with the query sequence

^b^Iden2 is the percentage sequence identity of the whole template chains with query sequence

^c^Cov-Represents the coverage of the threading alignment and is equal to the number of aligned residues divided by the length of the query protein

^d^Norm. Z-score is the normalized Z-score of the threading alignments. Alignment with a normalized Z-score>1 means a good alignment and vice versa.

^e^Aminomethyltransferase

^f^DmdA DMSP-dependent demethylase

^g^Glycine cleavage system T protein

^h^Dimethylglycine dehydrogenase complexed with tetrahydrofolate

**Predicted model: C-sore=0.48; TM-score=0.78 dev= 0.10**

| **Rank** | **Class** | **Gene name** | **Organism** | **PDB ID** | **TM-score^a^** | **RMSD^b^** | **IDEN^c^** | **Cov^d^** |
| --- | --- | --- | --- | --- | --- | --- | --- | --- |
| 1 | Oxidoreductase | *dmgdH* | *Rattus norvegicus* | 4p9sA | 0.885 | 1.02 | 0.170 | 0.900 |
| 2 | Oxidoreductase | *dmg^e^* | *Arthrobacter globiformis* | 1pj6A | 0.848 | 2.00 | 0.216 | 0.894 |
| 3 | Oxidoreductase | *soxA^f^* | *Stenotrophomonas maltophilia* | 2gagA | 0.790 | 2.68 | 0.198 | 0.854 |
| 4 | Oxidoreductase | *soxA* | *Corynebacterium sp. U-96* | 1x31A | 0.787 | 2.67 | 0.193 | 0.851 |
| 5 | AMT | *gcvT* | *Thermotoga maritima* | 1wooA | 0.777 | 1.07 | 0.297 | 0.792 |
| 6 | AMT | *gcvT* | *Pyrococcus horikoshii* | 1v5vA | 0.777 | 1.69 | 0.252 | 0.809 |
| 7 | AMT | *AMT(GCST)* | *Homo sapiens* | 1wsvB | 0.766 | 1.56 | 0.229 | 0.794 |
| 8 | AMT | *dmdA* | *Pelagibacter ubique HTCC1062* | 3tfiA | 0.762 | 2.09 | 0.254 | 0.811 |
| 9 | AMT | *gcvT* | *Bartonella henselae* | 3girA | 0.752 | 1.74 | 0.220 | 0.785 |
| 10 | AMT | *gcvT* | *Bacillus subtilis* | 1yx2B | 0.750 | 1.58 | 0.251 | 0.778 |

^ab^It is a standard for measuring structural similarity between two structures

^c^It is the percentage sequence identity in the structurally aligned region

^d^It represents the coverage of the alignment by TM-align and is equal to the number of structurally aligned residues divided by length of the query protein

^e^Dimethylglycine oxidase

^f^Heterotetrameric sarcosine oxidase alpha-subunit

**Table 39.** Up: Top ten analogs identified by LOMETS for threading alignments. Case of ABF64177.1. In the middle: Model predicted by I-TASSER. Down: Top ten identified structural analogs in PDB by TM-align

| **Rank** | **Class** | **Gene name** | **Organism** | **PDB ID** | **Iden1 (%)^a^** | **Iden2 (%)^b^** | **Cov^c^** | **Norm. Z-score^d^** |
| --- | --- | --- | --- | --- | --- | --- | --- | --- |
| 1 | AMT^e^ | *dmdA^f^* | *Pelagibacter ubique (HTCC1062)* | 3tfhA | 44 | 42 | 0.96 | 4.59 |
| 2 | AMT | *dmdA* | *Pelagibacter ubique (HTCC1062)* | 3tfhA | 44 | 42 | 0.95 | 5.21 |
| 3 | AMT | *dmdA* | *Pelagibacter ubique (HTCC1062)* | 3tfhA | 44 | 42 | 0.96 | 5.55 |
| 4 | Oxidoreductase | *dmgdH* | *Rattus norvegicus* | 4p9s | 17 | 22 | 0.97 | 3.28 |
| 5 | Oxidoreductase | *dmgdH^h^* | *Rattus norvegicus* | 4p9sA | 18 | 21 | 0.94 | 2.38 |
| 6 | AMT | *dmdA* | *Pelagibacter ubique (HTCC1062)* | 3tfhA | 44 | 42 | 0.95 | 5.58 |
| 7 | *Oxidoreductase* | *dmgdH* | *Rattus norvegicus* | 4p9s | 17 | 22 | 0.97 | 3.63 |
| 8 | AMT | *gcvT^g^* | *Bacillus subtilis* | 1yx2A | 25 | 26 | 0.91 | 6.61 |
| 9 | AMT | *dmdA* | *Pelagibacter ubique (HTCC1062)* | 3tfhA | 44 | 42 | 0.96 | 7.88 |
| 10 | AMT | *dmdA* | *Pelagibacter ubique (HTCC1062)* | 3tfhA | 44 | 42 | 0.95 | 11.88 |

^a^Iden1 is the percentage sequence identity of the templates in the threading aligned region with the query sequence

^b^Iden2 is the percentage sequence identity of the whole template chains with query sequence

^c^Cov-Represents the coverage of the threading alignment and is equal to the number of aligned residues divided by the length of the query protein

^d^Norm. Z-score is the normalized Z-score of the threading alignments. Alignment with a normalized Z-score>1 means a good alignment and vice versa.

^e^Aminomethyltransferase

^f^DmdA DMSP-dependent demethylase

^g^Glycine cleavage system T protein

^h^Dimethylglycine dehydrogenase complexed with tetrahydrofolate

**Predicted model: C-sore=1.62; TM-score=0.94 dev= 0.05**

| **Rank** | **Class** | **Gene name** | **Organism** | **PDB ID** | **TM-score^a^** | **RMSD^b^** | **IDEN^c^** | **Cov^d^** |
| --- | --- | --- | --- | --- | --- | --- | --- | --- |
| 1 | AMT | *dmdA* | *Pelagibacter ubique HTCC1062* | 3tfiA | 0.947 | 0.85 | 0.436 | 0.958 |
| 2 | Oxidoreductase | *dmgdH* | *Rattus norvegicus* | 4p9sA | 0.939 | 1.62 | 0.175 | 0.982 |
| 3 | Oxidoreductase | *dmg^e^* | *Arthrobacter globiformis* | 1pj6A | 0.921 | 2.06 | 0.246 | 0.982 |
| 4 | AMT | *gcvT* | *Pyrococcus horikoshii* | 1v5vA | 0.870 | 1.84 | 0.228 | 0.922 |
| 5 | Oxidoreductase | *soxA^f^* | *Stenotrophomonas maltophilia* | 2gagA | 0.866 | 2.29 | 0.192 | 0.932 |
| 6 | Oxidoreductase | *soxA* | *Corynebacterium sp. U-96* | 3ad7A | 0.865 | 2.37 | 0.184 | 0.935 |
| 7 | AMT | *gcvT* | *Thermotoga maritima* | 1wooA | 0.864 | 1.60 | 0.247 | 0.904 |
| 8 | AMT | *AMT(GCST)* | *Homo sapiens* | 1wsvA | 0.858 | 1.96 | 0.256 | 0.914 |
| 9 | AMT | *gcvT* | *Bartonella henselae* | 3girA | 0.849 | 1.88 | 0.242 | 0.901 |
| 10 | AMT | *gcvT* | *Bacillus subtilis* | 1yx2B | 0.846 | 2.05 | 0.252 | 0.906 |

^ab^It is a standard for measuring structural similarity between two structures

^c^It is the percentage sequence identity in the structurally aligned region

^d^It represents the coverage of the alignment by TM-align and is equal to the number of structurally aligned residues divided by length of the query protein

^e^Dimethylglycine oxidase

^f^Heterotetrameric sarcosine oxidase alpha-subunit

**Table 40**. Up: Top ten analogs identified by LOMETS for threading alignments. Case of AEI94210.1. In the middle: Model predicted by I-TASSER. Down: Top ten identified structural analogs in PDB by TM-align

| **Rank** | **Class** | **Gene name** | **Organism** | **PDB ID** | **Iden1 (%)^a^** | **Iden2 (%)^b^** | **Cov^c^** | **Norm. Z-score^d^** |
| --- | --- | --- | --- | --- | --- | --- | --- | --- |
| 1 | AMT^e^ | *dmdA^f^* | *Pelagibacter ubique (HTCC1062)* | 3tfhA | 44 | 44 | 1.00 | 4.83 |
| 2 | AMT | *dmdA* | *Pelagibacter ubique (HTCC1062)* | 3tfhA | 44 | 44 | 1.00 | 5.36 |
| 3 | AMT | *dmdA* | *Pelagibacter ubique (HTCC1062)* | 3tfhA | 44 | 44 | 1.00 | 5.76 |
| 4 | AMT | *dmdA* | *Pelagibacter ubique (HTCC1062)* | 3tfh | 44 | 44 | 0.99 | 3.17 |
| 5 | Oxidoreductase | *dmgdH^h^* | *Rattus norvegicus* | 4p9sA | 16 | 23 | 0.98 | 2.36 |
| 6 | AMT | *dmdA* | *Pelagibacter ubique (HTCC1062)* | 3tfhA | 44 | 44 | 1.00 | 5.86 |
| 7 | *AMT* | *dmdA* | *Pelagibacter ubique (HTCC1062)* | 3tfh | 42 | 44 | 0.98 | 3.53 |
| 8 | AMT | *gcvT^g^* | *Bacillus subtilis* | 1yx2A | 24 | 26 | 0.96 | 6.58 |
| 9 | AMT | *dmdA* | *Pelagibacter ubique (HTCC1062)* | 3tfhA | 44 | 44 | 1.00 | 7.87 |
| 10 | AMT | *dmdA* | *Pelagibacter ubique (HTCC1062)* | 3tfhA | 44 | 44 | 1.00 | 6.74 |

^a^Iden1 is the percentage sequence identity of the templates in the threading aligned region with the query sequence

^b^Iden2 is the percentage sequence identity of the whole template chains with query sequence

^c^Cov-Represents the coverage of the threading alignment and is equal to the number of aligned residues divided by the length of the query protein

^d^Norm. Z-score is the normalized Z-score of the threading alignments. Alignment with a normalized Z-score>1 means a good alignment and vice versa.

^e^Aminomethyltransferase

^f^DmdA DMSP-dependent demethylase

^g^Glycine cleavage system T protein

^h^Dimethylglycine dehydrogenase complexed with tetrahydrofolate

**Predicted model: C-sore=2; TM-score=0.99 dev= 0.04**

| **Rank** | **Class** | **Gene name** | **Organism** | **PDB ID** | **TM-score^a^** | **RMSD^b^** | **IDEN^c^** | **Cov^d^** |
| --- | --- | --- | --- | --- | --- | --- | --- | --- |
| 1 | AMT | *dmdA* | *Pelagibacter ubique HTCC1062* | 3tfhA | 0.997 | 0.41 | 0.436 | 1.000 |
| 2 | Oxidoreductase | *dmgdH* | *Rattus norvegicus* | 4p9sA | 0.930 | 1.80 | 0.175 | 0.984 |
| 3 | Oxidoreductase | *dmg^e^* | *Arthrobacter globiformis* | 1pj6A2 | 0.913 | 2.17 | 0.249 | 0.984 |
| 4 | Oxidoreductase | *dmg* | *Arthrobacter globiformis* | 1pj7A | 0.912 | 2.18 | 0.249 | 0.984 |
| 5 | AMT | *gcvT* | *Pyrococcus horikoshii* | 1v5vA | 0.907 | 1.92 | 0.231 | 0.967 |
| 6 | Oxidoreductase | *soxA^f^* | *Stenotrophomonas maltophilia* | 2gagA | 0.901 | 2.28 | 0.191 | 0.976 |
| 7 | Oxidoreductase | *soxA* | *Corynebacterium sp. U-96* | 3ad7A | 0.900 | 2.26 | 0.191 | 0.976 |
| 8 | AMT | *gcvT* | *Thermotoga maritima* | 1worA | 0.898 | 1.79 | 0.255 | 0.951 |
| 9 | AMT | *AMT(GCST)* | *Homo sapiens* | 1wsrA | 0.892 | 2.07 | 0.259 | 0.959 |
| 10 | AMT | *gcvT* | *Bartonella henselae* | 3girA | 0.886 | 1.93 | 0.225 | 0.946 |

^ab^It is a standard for measuring structural similarity between two structures

^c^It is the percentage sequence identity in the structurally aligned region

^d^It represents the coverage of the alignment by TM-align and is equal to the number of structurally aligned residues divided by length of the query protein

^e^Dimethylglycine oxidase

^f^Heterotetrameric sarcosine oxidase alpha-subunit

**Table 41**. Up: Top ten analogs identified by LOMETS for threading alignments. Case of ABG31871. In the middle: Model predicted by I-TASSER. Down: Top ten identified structural analogs in PDB by TM-align

| **Rank** | **Class** | **Gene name** | **Organism** | **PDB ID** | **Iden1 (%)^a^** | **Iden2 (%)^b^** | **Cov^c^** | **Norm. Z-score^d^** |
| --- | --- | --- | --- | --- | --- | --- | --- | --- |
| 1 | AMT^e^ | *dmdA^f^* | *Pelagibacter ubique (HTCC1062)* | 3tfhA | 44 | 44 | 1.00 | 4.86 |
| 2 | AMT | *dmdA* | *Pelagibacter ubique (HTCC1062)* | 3tfhA | 44 | 44 | 1.00 | 5.39 |
| 3 | AMT | *dmdA* | *Pelagibacter ubique (HTCC1062)* | 3tfhA | 44 | 44 | 1.00 | 5.80 |
| 4 | AMT | *dmdA* | *Pelagibacter ubique (HTCC1062)* | 3tfh | 44 | 44 | 0.99 | 3.18 |
| 5 | Oxidoreductase | *dmgdH^h^* | *Rattus norvegicus* | 4p9sA | 16 | 22 | 0.98 | 2.36 |
| 6 | AMT | *dmdA* | *Pelagibacter ubique (HTCC1062)* | 3tfhA | 44 | 44 | 1.00 | 5.93 |
| 7 | *AMT* | *dmdA* | *Pelagibacter ubique (HTCC1062)* | 3tfh | 44 | 44 | 0.98 | 3.53 |
| 8 | AMT | *gcvT^g^* | *Bacillus subtilis* | 1yx2A | 24 | 26 | 0.96 | 6.66 |
| 9 | AMT | *dmdA* | *Pelagibacter ubique (HTCC1062)* | 3tfhA | 44 | 44 | 1.00 | 7.94 |
| 10 | AMT | *dmdA* | *Pelagibacter ubique (HTCC1062)* | 3tfhA | 44 | 44 | 1.00 | 6.82 |

^a^Iden1 is the percentage sequence identity of the templates in the threading aligned region with the query sequence

^b^Iden2 is the percentage sequence identity of the whole template chains with query sequence

^c^Cov-Represents the coverage of the threading alignment and is equal to the number of aligned residues divided by the length of the query protein

^d^Norm. Z-score is the normalized Z-score of the threading alignments. Alignment with a normalized Z-score>1 means a good alignment and vice versa.

^e^Aminomethyltransferase

^f^DmdA DMSP-dependent demethylase

^g^Glycine cleavage system T protein

^h^Dimethylglycine dehydrogenase complexed with tetrahydrofolate

**Predicted model: C-sore=2; TM-score=0.99 dev= 0.04**

| **Rank** | **Class** | **Gene name** | **Organism** | **PDB ID** | **TM-score^a^** | **RMSD^b^** | **IDEN^c^** | **Cov^d^** |
| --- | --- | --- | --- | --- | --- | --- | --- | --- |
| 1 | AMT | *dmdA* | *Pelagibacter ubique HTCC1062* | 3tfhA | 0.997 | 0.41 | 0.436 | 1.000 |
| 2 | Oxidoreductase | *dmgdH* | *Rattus norvegicus* | 4p9sA | 0.931 | 1.79 | 0.175 | 0.984 |
| 3 | Oxidoreductase | *dmg^e^* | *Arthrobacter globiformis* | 1pj6A2 | 0.914 | 2.16 | 0.258 | 0.984 |
| 4 | Oxidoreductase | *dmg* | *Arthrobacter globiformis* | 1pj7A | 0.913 | 2.18 | 0.258 | 0.984 |
| 5 | AMT | *gcvT* | *Pyrococcus horikoshii* | 1v5vA | 0.905 | 1.94 | 0.237 | 0.967 |
| 6 | Oxidoreductase | *soxA^f^* | *Stenotrophomonas maltophilia* | 2gagA | 0.900 | 2.25 | 0.202 | 0.976 |
| 7 | Oxidoreductase | *soxA* | *Corynebacterium sp. U-96* | 3ad7A | 0.900 | 2.26 | 0.196 | 0.976 |
| 8 | AMT | *gcvT* | *Thermotoga maritima* | 1wooA | 0.898 | 1.73 | 0.261 | 0.948 |
| 9 | AMT | *AMT(GCST)* | *Homo sapiens* | 1wsrA | 0.892 | 2.07 | 0.267 | 0.959 |
| 10 | AMT | *gcvT* | *Bartonella henselae* | 3girA | 0.887 | 1.92 | 0.228 | 0.946 |

^ab^It is a standard for measuring structural similarity between two structures

^c^It is the percentage sequence identity in the structurally aligned region

^d^It represents the coverage of the alignment by TM-align and is equal to the number of structurally aligned residues divided by length of the query protein

^e^Dimethylglycine oxidase

^f^Heterotetrameric sarcosine oxidase alpha-subunit

**Table 42**. Up: Top ten analogs identified by LOMETS for threading alignments. Case of ABD55296.1. In the middle: Model predicted by I-TASSER. Down: Top ten identified structural analogs in PDB by TM-align

| **Rank** | **Class** | **Gene name** | **Organism** | **PDB ID** | **Iden1 (%)^a^** | **Iden2 (%)^b^** | **Cov^c^** | **Norm. Z-score^d^** |
| --- | --- | --- | --- | --- | --- | --- | --- | --- |
| 1 | AMT^e^ | *dmdA^f^* | *Pelagibacter ubique (HTCC1062)* | 3tfhA | 43 | 44 | 1.00 | 4.70 |
| 2 | AMT | *dmdA* | *Pelagibacter ubique (HTCC1062)* | 3tfhA | 44 | 44 | 1.00 | 5.33 |
| 3 | AMT | *dmdA* | *Pelagibacter ubique (HTCC1062)* | 3tfhA | 43 | 44 | 1.00 | 5.76 |
| 4 | AMT | *dmdA* | *Pelagibacter ubique (HTCC1062)* | 3tfh | 45 | 44 | 0.99 | 3.18 |
| 5 | Oxidoreductase | *dmgdH^h^* | *Rattus norvegicus* | 4p9sA | 18 | 21 | 0.98 | 2.36 |
| 6 | AMT | *dmdA* | *Pelagibacter ubique (HTCC1062)* | 3tfhA | 43 | 44 | 0.99 | 5.76 |
| 7 | *AMT* | *dmdA* | *Pelagibacter ubique (HTCC1062)* | 3tfh | 43 | 44 | 0.98 | 3.52 |
| 8 | AMT | *gcvT^g^* | *Bacillus subtilis* | 1yx2A | 24 | 26 | 0.96 | 6.61 |
| 9 | AMT | *dmdA* | *Pelagibacter ubique (HTCC1062)* | 3tfhA | 44 | 44 | 1.00 | 7.68 |
| 10 | AMT | *dmdA* | *Pelagibacter ubique (HTCC1062)* | 3tfhA | 43 | 44 | 1.00 | 6.76 |

^a^Iden1 is the percentage sequence identity of the templates in the threading aligned region with the query sequence

^b^Iden2 is the percentage sequence identity of the whole template chains with query sequence

^c^Cov-Represents the coverage of the threading alignment and is equal to the number of aligned residues divided by the length of the query protein

^d^Norm. Z-score is the normalized Z-score of the threading alignments. Alignment with a normalized Z-score>1 means a good alignment and vice versa.

^e^Aminomethyltransferase

^f^DmdA DMSP-dependent demethylase

^g^Glycine cleavage system T protein

^h^Dimethylglycine dehydrogenase complexed with tetrahydrofolate

**Predicted model: C-sore=2; TM-score=0.99 dev= 0.04**

| **Rank** | **Class** | **Gene name** | **Organism** | **PDB ID** | **TM-score^a^** | **RMSD^b^** | **IDEN^c^** | **Cov^d^** |
| --- | --- | --- | --- | --- | --- | --- | --- | --- |
| 1 | AMT | *dmdA* | *Pelagibacter ubique HTCC1062* | 3tfhA | 0.997 | 0.37 | 0.439 | 1.000 |
| 2 | Oxidoreductase | *dmgdH* | *Rattus norvegicus* | 4p9sA | 0.929 | 1.82 | 0.183 | 0.984 |
| 3 | Oxidoreductase | *dmg^e^* | *Arthrobacter globiformis* | 1pj6A2 | 0.913 | 2.16 | 0.263 | 0.984 |
| 4 | Oxidoreductase | *dmg* | *Arthrobacter globiformis* | 1pj7A | 0.913 | 2.17 | 0.263 | 0.984 |
| 5 | AMT | *gcvT* | *Pyrococcus horikoshii* | 1v5vA | 0.904 | 1.95 | 0.239 | 0.967 |
| 6 | Oxidoreductase | *soxA^f^* | *Stenotrophomonas maltophilia* | 2gagA | 0.899 | 2.31 | 0.202 | 0.976 |
| 7 | Oxidoreductase | *soxA* | *Corynebacterium sp. U-96* | 3ad7A | 0.898 | 2.28 | 0.202 | 0.976 |
| 8 | AMT | *gcvT* | *Thermotoga maritima* | 1worA | 0.896 | 1.82 | 0.249 | 0.951 |
| 9 | AMT | *AMT(GCST)* | *Homo sapiens* | 1wsrA | 0.890 | 2.09 | 0.244 | 0.959 |
| 10 | AMT | *gcvT* | *Bartonella henselae* | 3girA | 0.885 | 1.94 | 0.236 | 0.946 |

^ab^It is a standard for measuring structural similarity between two structures

^c^It is the percentage sequence identity in the structurally aligned region

^d^It represents the coverage of the alignment by TM-align and is equal to the number of structurally aligned residues divided by length of the query protein

^e^Dimethylglycine oxidase

^f^Heterotetrameric sarcosine oxidase alpha-subunit

**Table 43**. Up: Top ten analogs identified by LOMETS for threading alignments. Case of ABV94056.1. In the middle: Model predicted by I-TASSER. Down: Top ten identified structural analogs in PDB by TM-align

| **Rank** | **Class** | **Gene name** | **Organism** | **PDB ID** | **Iden1 (%)^a^** | **Iden2 (%)^b^** | **Cov^c^** | **Norm. Z-score^d^** |
| --- | --- | --- | --- | --- | --- | --- | --- | --- |
| 1 | AMT^e^ | *dmdA^f^* | *Pelagibacter ubique (HTCC1062)* | 3tfhA | 41 | 41 | 1.00 | 4.91 |
| 2 | AMT | *dmdA* | *Pelagibacter ubique (HTCC1062)* | 3tfhA | 41 | 41 | 1.00 | 5.45 |
| 3 | AMT | *dmdA* | *Pelagibacter ubique (HTCC1062)* | 3tfhA | 41 | 41 | 1.00 | 5.84 |
| 4 | AMT | *dmdA* | *Pelagibacter ubique (HTCC1062)* | 3tfh | 41 | 41 | 0.99 | 3.20 |
| 5 | Oxidoreductase | *dmgdH^h^* | *Rattus norvegicus* | 4p9sA | 18 | 21 | 0.98 | 2.35 |
| 6 | AMT | *dmdA* | *Pelagibacter ubique (HTCC1062)* | 3tfhA | 41 | 41 | 1.00 | 5.92 |
| 7 | *AMT* | *dmdA* | *Pelagibacter ubique (HTCC1062)* | 3tfh | 40 | 41 | 0.98 | 3.52 |
| 8 | AMT | *gcvT^g^* | *Bacillus subtilis* | 1yx2A | 25 | 25 | 0.96 | 6.40 |
| 9 | AMT | *dmdA* | *Pelagibacter ubique (HTCC1062)* | 3tfhA | 41 | 41 | 1.00 | 8.02 |
| 10 | AMT | *dmdA* | *Pelagibacter ubique (HTCC1062)* | 3tfhA | 41 | 41 | 1.00 | 6.90 |

^a^Iden1 is the percentage sequence identity of the templates in the threading aligned region with the query sequence

^b^Iden2 is the percentage sequence identity of the whole template chains with query sequence

^c^Cov-Represents the coverage of the threading alignment and is equal to the number of aligned residues divided by the length of the query protein

^d^Norm. Z-score is the normalized Z-score of the threading alignments. Alignment with a normalized Z-score>1 means a good alignment and vice versa.

^e^Aminomethyltransferase

^f^DmdA DMSP-dependent demethylase

^g^Glycine cleavage system T protein

^h^Dimethylglycine dehydrogenase complexed with tetrahydrofolate

**Predicted model: C-sore=2; TM-score=0.99 dev= 0.04**

| **Rank** | **Class** | **Gene name** | **Organism** | **PDB ID** | **TM-score^a^** | **RMSD^b^** | **IDEN^c^** | **Cov^d^** |
| --- | --- | --- | --- | --- | --- | --- | --- | --- |
| 1 | AMT | *dmdA* | *Pelagibacter ubique HTCC1062* | 3tfhA | 0.998 | 0.35 | 0.408 | 1.000 |
| 2 | Oxidoreductase | *dmgdH* | *Rattus norvegicus* | 4p9sA | 0.927 | 1.87 | 0.180 | 0.984 |
| 3 | Oxidoreductase | *dmg^e^* | *Arthrobacter globiformis* | 1pj6A2 | 0.910 | 2.25 | 0.276 | 0.984 |
| 4 | Oxidoreductase | *dmg* | *Arthrobacter globiformis* | 1pj7A | 0.910 | 2.25 | 0.276 | 0.984 |
| 5 | AMT | *gcvT* | *Pyrococcus horikoshii* | 1v5vA | 0.902 | 1.94 | 0.237 | 0.965 |
| 6 | Oxidoreductase | *soxA^f^* | *Stenotrophomonas maltophilia* | 2gagA | 0.897 | 2.40 | 0.201 | 0.976 |
| 7 | Oxidoreductase | *soxA* | *Corynebacterium sp. U-96* | 3ad7A | 0.897 | 2.37 | 0.201 | 0.976 |
| 8 | AMT | *gcvT* | *Thermotoga maritima* | 1wooA | 0.895 | 1.75 | 0.253 | 0.946 |
| 9 | AMT | *AMT(GCST)* | *Homo sapiens* | 1wsrA | 0.888 | 2.09 | 0.261 | 0.957 |
| 10 | AMT | *gcvT* | *Bartonella henselae* | 3girA | 0.883 | 1.94 | 0.242 | 0.943 |

^ab^It is a standard for measuring structural similarity between two structures

^c^It is the percentage sequence identity in the structurally aligned region

^d^It represents the coverage of the alignment by TM-align and is equal to the number of structurally aligned residues divided by length of the query protein

^e^Dimethylglycine oxidase

^f^Heterotetrameric sarcosine oxidase alpha-subunit

**Table 44.** Up: Top ten analogs identified by LOMETS for threading alignments. Case of AAZ21068.1. In the middle: Model predicted by I-TASSER. Down: Top ten identified structural analogs in PDB by TM-align

| **Rank** | **Class** | **Gene name** | **Organism** | **PDB ID** | **Iden1 (%)^a^** | **Iden2 (%)^b^** | **Cov^c^** | **Norm. Z-score^d^** |
| --- | --- | --- | --- | --- | --- | --- | --- | --- |
| 1 | AMT^e^ | *dmdA^f^* | *Pelagibacter ubique (HTCC1062)* | 3tfhA | 100 | 100 | 1.00 | 4.99 |
| 2 | AMT | *dmdA* | *Pelagibacter ubique (HTCC1062)* | 3tfhA | 100 | 100 | 1.00 | 5.58 |
| 3 | AMT | *dmdA* | *Pelagibacter ubique (HTCC1062)* | 3tfhA | 100 | 100 | 1.00 | 5.83 |
| 4 | AMT | *dmdA* | *Pelagibacter ubique (HTCC1062)* | 3tfh | 100 | 100 | 1.00 | 3.31 |
| 5 | Oxidoreductase | *dmgdH^g^* | *Rattus norvegicus* | 4p9sA | 18 | 21 | 0.98 | 2.35 |
| 6 | AMT | *dmdA* | *Pelagibacter ubique (HTCC1062)* | 3tfhA | 100 | 100 | 1.00 | 6.20 |
| 7 | *AMT* | *dmdA* | *Pelagibacter ubique (HTCC1062)* | 3tfh | 0.94 | 100 | 0.99 | 3.57 |
| 8 | AMT | *dmdA* | *Pelagibacter ubique (HTCC1062)* | 3tfjA | 100 | 100 | 1.00 | 6.47 |
| 9 | AMT | *dmdA* | *Pelagibacter ubique (HTCC1062)* | 3tfhA | 100 | 100 | 1.00 | 8.10 |
| 10 | AMT | *dmdA* | *Pelagibacter ubique (HTCC1062)* | 3tfhA | 100 | 100 | 1.00 | 7.00 |

^a^Iden1 is the percentage sequence identity of the templates in the threading aligned region with the query sequence

^b^Iden2 is the percentage sequence identity of the whole template chains with query sequence

^c^Cov-Represents the coverage of the threading alignment and is equal to the number of aligned residues divided by the length of the query protein

^d^Norm. Z-score is the normalized Z-score of the threading alignments. Alignment with a normalized Z-score>1 means a good alignment and vice versa.

^e^Aminomethyltransferase

^f^DmdA DMSP-dependent demethylase

^g^Dimethylglycine dehydrogenase complexed with tetrahydrofolate

**Predicted model: C-sore=2; TM-score=0.99 dev= 0.04**

| **Rank** | **Class** | **Gene name** | **Organism** | **PDB ID** | **TM-score^a^** | **RMSD^b^** | **IDEN^c^** | **Cov^d^** |
| --- | --- | --- | --- | --- | --- | --- | --- | --- |
| 1 | AMT | *dmdA* | *Pelagibacter ubique HTCC1062* | 3tfhA | 0.997 | 0.41 | 1.000 | 1.000 |
| 2 | Oxidoreductase | *dmgdH* | *Rattus norvegicus* | 4p9sA | 0.929 | 1.83 | 0.193 | 0.984 |
| 3 | Oxidoreductase | *dmg^e^* | *Arthrobacter globiformis* | 1pj6A | 0.912 | 2.20 | 0.220 | 0.984 |
| 4 | AMT | *gcvT* | *Pyrococcus horikoshii* | 1v5vA | 0.901 | 1.93 | 0.231 | 0.962 |
| 5 | Oxidoreductase | *soxA^f^* | *Corynebacterium sp. U-96* | 3ad7A | 0.896 | 2.34 | 0.206 | 0.973 |
| 6 | Oxidoreductase | *soxA* | *Stenotrophomonas maltophilia* | 2gagA | 0.896 | 2.28 | 0.198 | 0.970 |
| 7 | AMT | *gcvT* | *Thermotoga maritima* | 1wooA | 0.894 | 1.72 | 0.236 | 0.943 |
| 8 | AMT | *AMT(GCST)* | *Homo sapiens* | 1wsrA | 0.887 | 2.07 | 0.185 | 0.954 |
| 9 | AMT | *gcvT* | *Bartonella henselae* | 3girA | 0.882 | 1.91 | 0.196 | 0.940 |
| 10 | AMT | *gcvT* | *Bacillus subtilis* | 1yx2B | 0.875 | 2.15 | 0.246 | 0.946 |

^ab^It is a standard for measuring structural similarity between two structures

^c^It is the percentage sequence identity in the structurally aligned region

^d^It represents the coverage of the alignment by TM-align and is equal to the number of structurally aligned residues divided by length of the query protein

^e^Dimethylglycine oxidase

^f^Heterotetrameric sarcosine oxidase alpha-subunit

**Table 45**. Up: Top ten analogs identified by LOMETS for threading alignments. Case of AFS46782.1. In the middle: Model predicted by I-TASSER. Down: Top ten identified structural analogs in PDB by TM-align

| **Rank** | **Class** | **Gene name** | **Organism** | **PDB ID** | **Iden1 (%)^a^** | **Iden2 (%)^b^** | **Cov^c^** | **Norm. Z-score^d^** |
| --- | --- | --- | --- | --- | --- | --- | --- | --- |
| 1 | AMT^e^ | *dmdA^f^* | *Pelagibacter ubique (HTCC1062)* | 3tfhA | 76 | 76 | 1.00 | 4.94 |
| 2 | AMT | *dmdA* | *Pelagibacter ubique (HTCC1062)* | 3tfhA | 76 | 76 | 1.00 | 5.45 |
| 3 | AMT | *dmdA* | *Pelagibacter ubique (HTCC1062)* | 3tfhA | 74 | 76 | 1.00 | 5.62 |
| 4 | AMT | *dmdA* | *Pelagibacter ubique (HTCC1062)* | 3tfh | 75 | 76 | 0.99 | 3.23 |
| 5 | Oxidoreductase | *dmgdH^h^* | *Rattus norvegicus* | 4p9sA | 17 | 21 | 0.98 | 2.36 |
| 6 | AMT | *dmdA* | *Pelagibacter ubique (HTCC1062)* | 3tfhA | 76 | 76 | 1.00 | 6.08 |
| 7 | *AMT* | *dmdA* | *Pelagibacter ubique (HTCC1062)* | 3tfh | 71 | 76 | 0.99 | 3.57 |
| 8 | AMT | *gcvT^g^* | *Bacillus subtilis* | 1yx2A | 26 | 27 | 0.96 | 6.77 |
| 9 | AMT | *dmdA* | *Pelagibacter ubique (HTCC1062)* | 3tfhA | 76 | 76 | 1.00 | 7.90 |
| 10 | AMT | *dmdA* | *Pelagibacter ubique (HTCC1062)* | 3tfhA | 74 | 76 | 1.00 | 6.85 |

^a^Iden1 is the percentage sequence identity of the templates in the threading aligned region with the query sequence

^b^Iden2 is the percentage sequence identity of the whole template chains with query sequence

^c^Cov-Represents the coverage of the threading alignment and is equal to the number of aligned residues divided by the length of the query protein

^d^Norm. Z-score is the normalized Z-score of the threading alignments. Alignment with a normalized Z-score>1 means a good alignment and vice versa.

^e^Aminomethyltransferase

^f^DmdA DMSP-dependent demethylase

^g^Glycine cleavage system T protein

^h^Dimethylglycine dehydrogenase complexed with tetrahydrofolate

**Predicted model: C-sore=1.95; TM-score=0.99 dev= 0.04**

| **Rank** | **Class** | **Gene name** | **Organism** | **PDB ID** | **TM-score^a^** | **RMSD^b^** | **IDEN^c^** | **Cov^d^** |
| --- | --- | --- | --- | --- | --- | --- | --- | --- |
| 1 | AMT | *dmdA* | *Pelagibacter ubique HTCC1062* | 3tfhA | 0.997 | 0.41 | 0.753 | 1.000 |
| 2 | Oxidoreductase | *dmgdH* | *Rattus norvegicus* | 4p9sA | 0.931 | 1.84 | 0.182 | 0.986 |
| 3 | Oxidoreductase | *dmg^e^* | *Arthrobacter globiformis* | 1pj6A | 0.913 | 2.22 | 0.212 | 0.986 |
| 4 | AMT | *gcvT* | *Pyrococcus horikoshii* | 1v5vA | 0.900 | 1.99 | 0.237 | 0.965 |
| 5 | Oxidoreductase | *soxA^f^* | *Stenotrophomonas maltophilia* | 2gagA | 0.895 | 2.30 | 0.201 | 0.970 |
| 6 | Oxidoreductase | *soxA* | *Corynebacterium sp. U-96* | 3ad7A | 0.895 | 2.36 | 0.218 | 0.973 |
| 7 | AMT | *gcvT* | *Thermotoga maritima* | 1wooA | 0.893 | 1.77 | 0.253 | 0.946 |
| 8 | AMT | *AMT(GCST)* | *Homo sapiens* | 1wsrA | 0.885 | 2.10 | 0.199 | 0.954 |
| 9 | AMT | *gcvT* | *Bartonella henselae* | 3girA | 0.880 | 1.99 | 0.207 | 0.943 |
| 10 | AMT | *gcvT* | *Bacillus subtilis* | 1yx2B | 0.876 | 2.10 | 0.259 | 0.946 |

^ab^It is a standard for measuring structural similarity between two structures

^c^It is the percentage sequence identity in the structurally aligned region

^d^It represents the coverage of the alignment by TM-align and is equal to the number of structurally aligned residues divided by length of the query protein

^e^Dimethylglycine oxidase

^f^Heterotetrameric sarcosine oxidase alpha-subunit

**Table 46**. Up: Top ten analogs identified by LOMETS for threading alignments. Case of AFS48343.1. In the middle: Model predicted by I-TASSER. Down: Top ten identified structural analogs in PDB by TM-align

| **Rank** | **Class** | **Gene name** | **Organism** | **PDB ID** | **Iden1 (%)^a^** | **Iden2 (%)^b^** | **Cov^c^** | **Norm. Z-score^d^** |
| --- | --- | --- | --- | --- | --- | --- | --- | --- |
| 1 | AMT^e^ | *dmdA^f^* | *Pelagibacter ubique (HTCC1062)* | 3tfhA | 56 | 55 | 1.00 | 4.95 |
| 2 | AMT | *dmdA* | *Pelagibacter ubique (HTCC1062)* | 3tfhA | 55 | 55 | 1.00 | 5.39 |
| 3 | AMT | *dmdA* | *Pelagibacter ubique (HTCC1062)* | 3tfhA | 55 | 55 | 1.00 | 5.76 |
| 4 | AMT | *dmdA* | *Pelagibacter ubique (HTCC1062)* | 3tfh | 55 | 55 | 0.99 | 3.19 |
| 5 | Oxidoreductase | *dmgdH^h^* | *Rattus norvegicus* | 4p9sA | 18 | 23 | 0.98 | 2.36 |
| 6 | AMT | *dmdA* | *Pelagibacter ubique (HTCC1062)* | 3tfhA | 55 | 55 | 0.99 | 5.96 |
| 7 | *AMT* | *dmdA* | *Pelagibacter ubique (HTCC1062)* | 3tfh | 54 | 55 | 0.98 | 3.55 |
| 8 | AMT | *gcvT^g^* | *Bacillus subtilis* | 1yx2A | 23 | 25 | 0.96 | 6.47 |
| 9 | AMT | *dmdA* | *Pelagibacter ubique (HTCC1062)* | 3tfhA | 56 | 55 | 1.00 | 7.94 |
| 10 | AMT | *dmdA* | *Pelagibacter ubique (HTCC1062)* | 3tfhA | 56 | 55 | 1.00 | 6.73 |

^a^Iden1 is the percentage sequence identity of the templates in the threading aligned region with the query sequence

^b^Iden2 is the percentage sequence identity of the whole template chains with query sequence

^c^Cov-Represents the coverage of the threading alignment and is equal to the number of aligned residues divided by the length of the query protein

^d^Norm. Z-score is the normalized Z-score of the threading alignments. Alignment with a normalized Z-score>1 means a good alignment and vice versa.

^e^Aminomethyltransferase

^f^DmdA DMSP-dependent demethylase

^g^Glycine cleavage system T protein

^h^Dimethylglycine dehydrogenase complexed with tetrahydrofolate

**Predicted model: C-sore=2; TM-score=0.99 dev= 0.04**

| **Rank** | **Class** | **Gene name** | **Organism** | **PDB ID** | **TM-score^a^** | **RMSD^b^** | **IDEN^c^** | **Cov^d^** |
| --- | --- | --- | --- | --- | --- | --- | --- | --- |
| 1 | AMT | *dmdA* | *Pelagibacter ubique HTCC1062* | 3tfhA | 0.995 | 0.36 | 0.553 | 0.997 |
| 2 | Oxidoreductase | *dmgdH* | *Rattus norvegicus* | 4p9sA | 0.930 | 1.84 | 0.193 | 0.986 |
| 3 | Oxidoreductase | *dmg^e^* | *Arthrobacter globiformis* | 1pj6A | 0.913 | 2.22 | 0.218 | 0.986 |
| 4 | Oxidoreductase | *dmg* | *Arthrobacter globiformis* | 1pj6A2 | 0.913 | 2.22 | 0.218 | 0.986 |
| 5 | AMT | *gcvT* | *Pyrococcus horikoshii* | 1v5vA | 0.903 | 1.99 | 0.247 | 0.967 |
| 6 | Oxidoreductase | *soxA^f^* | *Stenotrophomonas maltophilia* | 2gagA | 0.898 | 2.30 | 0.196 | 0.973 |
| 7 | Oxidoreductase | *soxA* | *Corynebacterium sp. U-96* | 3ad7A | 0.897 | 2.27 | 0.201 | 0.973 |
| 8 | AMT | *gcvT* | *Thermotoga maritima* | 1wooA | 0.896 | 1.77 | 0.244 | 0.948 |
| 9 | AMT | *AMT(GCST)* | *Homo sapiens* | 1wsrA | 0.889 | 2.09 | 0.233 | 0.957 |
| 10 | AMT | *gcvT* | *Bartonella henselae* | 3girA | 0.883 | 1.98 | 0.204 | 0.946 |

^ab^It is a standard for measuring structural similarity between two structures

^c^It is the percentage sequence identity in the structurally aligned region

^d^It represents the coverage of the alignment by TM-align and is equal to the number of structurally aligned residues divided by length of the query protein

^e^Dimethylglycine oxidase

^f^Heterotetrameric sarcosine oxidase alpha-subunit

**Table 47**. Up: Top ten analogs identified by LOMETS for threading alignments. Case of ADE38317.1. In the middle: Model predicted by I-TASSER. Down: Top ten identified structural analogs in PDB by TM-align

| **Rank** | **Class** | **Gene name** | **Organism** | **PDB ID** | **Iden1 (%)^a^** | **Iden2 (%)^b^** | **Cov^c^** | **Norm. Z-score^d^** |
| --- | --- | --- | --- | --- | --- | --- | --- | --- |
| 1 | AMT^e^ | *dmdA^f^* | *Pelagibacter ubique (HTCC1062)* | 3tfhA | 46 | 46 | 0.99 | 4.75 |
| 2 | AMT | *dmdA* | *Pelagibacter ubique (HTCC1062)* | 3tfhA | 46 | 46 | 0.99 | 5.36 |
| 3 | AMT | *dmdA* | *Pelagibacter ubique (HTCC1062)* | 3tfhA | 46 | 46 | 0.99 | 5.59 |
| 4 | AMT | *dmdA* | *Pelagibacter ubique (HTCC1062)* | 3tfh | 46 | 46 | 0.99 | 3.23 |
| 5 | Oxidoreductase | *dmgdH^h^* | *Rattus norvegicus* | 4p9sA | 19 | 22 | 0.97 | 2.35 |
| 6 | AMT | *dmdA* | *Pelagibacter ubique (HTCC1062)* | 3tfhA | 46 | 46 | 0.99 | 5.72 |
| 7 | *AMT* | *dmdA* | *Pelagibacter ubique (HTCC1062)* | 3tfh | 46 | 46 | 0.98 | 3.58 |
| 8 | AMT | *gcvT^g^* | *Thermotoga maritima* | 1wosA | 23 | 24 | 0.94 | 5.99 |
| 9 | AMT | *dmdA* | *Pelagibacter ubique (HTCC1062)* | 3tfhA | 46 | 46 | 0.99 | 7.99 |
| 10 | AMT | *dmdA* | *Pelagibacter ubique (HTCC1062)* | 3tfhA | 46 | 46 | 0.99 | 6.75 |

^a^Iden1 is the percentage sequence identity of the templates in the threading aligned region with the query sequence

^b^Iden2 is the percentage sequence identity of the whole template chains with query sequence

^c^Cov-Represents the coverage of the threading alignment and is equal to the number of aligned residues divided by the length of the query protein

^d^Norm. Z-score is the normalized Z-score of the threading alignments. Alignment with a normalized Z-score>1 means a good alignment and vice versa.

^e^Aminomethyltransferase

^f^DmdA DMSP-dependent demethylase

^g^Glycine cleavage system T protein

^h^Dimethylglycine dehydrogenase complexed with tetrahydrofolate

**Predicted model: C-sore=1.96; TM-score=0.99 dev= 0.04**

| **Rank** | **Class** | **Gene name** | **Organism** | **PDB ID** | **TM-score^a^** | **RMSD^b^** | **IDEN^c^** | **Cov^d^** |
| --- | --- | --- | --- | --- | --- | --- | --- | --- |
| 1 | AMT | *dmdA* | *Pelagibacter ubique HTCC1062* | 3tfhA | 0.988 | 0.47 | 0.462 | 0.992 |
| 2 | Oxidoreductase | *dmgdH* | *Rattus norvegicus* | 4p9sA | 0.924 | 1.83 | 0.196 | 0.978 |
| 3 | Oxidoreductase | *dmg^e^* | *Arthrobacter globiformis* | 1pj6A | 0.907 | 2.21 | 0.231 | 0.978 |
| 4 | AMT | *gcvT* | *Pyrococcus horikoshii* | 1v5vA | 0.895 | 1.92 | 0.254 | 0.954 |
| 5 | Oxidoreductase | *soxA^f^* | *Corynebacterium sp. U-96* | 3ad7A | 0.894 | 2.24 | 0.214 | 0.968 |
| 6 | Oxidoreductase | *soxA* | *Stenotrophomonas maltophilia* | 2gagA | 0.894 | 2.24 | 0.211 | 0.968 |
| 7 | AMT | *gcvT* | *Thermotoga maritima* | 1wooA | 0.888 | 1.74 | 0.227 | 0.938 |
| 8 | AMT | *AMT(GCST)* | *Homo sapiens* | 1wsrA | 0.882 | 2.08 | 0.227 | 0.949 |
| 9 | AMT | *gcvT* | *Bartonella henselae* | 3girA | 0.878 | 1.92 | 0.248 | 0.935 |
| 10 | AMT | *gcvT* | *Bacillus subtilis* | 1yx2B | 0.868 | 2.09 | 0.265 | 0.935 |

^ab^It is a standard for measuring structural similarity between two structures

^c^It is the percentage sequence identity in the structurally aligned region

^d^It represents the coverage of the alignment by TM-align and is equal to the number of structurally aligned residues divided by length of the query protein

^e^Dimethylglycine oxidase

^f^Heterotetrameric sarcosine oxidase alpha-subunit

**Table 48**. Up: Top ten analogs identified by LOMETS for threading alignments. Case of AAV95190.1. In the middle: Model predicted by I-TASSER. Down: Top ten identified structural analogs in PDB by TM-align

| **Rank** | **Class** | **Gene name** | **Organism** | **PDB ID** | **Iden1 (%)^a^** | **Iden2 (%)^b^** | **Cov^c^** | **Norm. Z-score^d^** |
| --- | --- | --- | --- | --- | --- | --- | --- | --- |
| 1 | AMT^e^ | *dmdA^f^* | *Pelagibacter ubique (HTCC1062)* | 3tfhA | 40 | 41 | 0.99 | 4.65 |
| 2 | AMT | *gcvT^g^* | *Pyrococcus horikoshii* | 1v5vA | 22 | 26 | 0.98 | 5.21 |
| 3 | AMT | *gcvT* | *Thermotoga maritima* | 1worA | 22 | 25 | 0.96 | 5.69 |
| 4 | AMT | *dmdA* | *Pelagibacter ubique (HTCC1062)* | 3tfh | 41 | 41 | 1.00 | 3.20 |
| 5 | Oxidoreductase | *dmgdH^h^* | *Rattus norvegicus* | 4p9sA | 18 | 21 | 0.99 | 2.36 |
| 6 | AMT | *dmdA* | *Pelagibacter ubique (HTCC1062)* | 3tfhA | 39 | 41 | 0.99 | 5.60 |
| 7 | *AMT* | *dmgdH* | *Rattus norvegicus* | 4p9sA | 18 | 21 | 0.98 | 3.52 |
| 8 | AMT | *gcvT* | *Bacillus subtilis* | 1yx2A | 23 | 24 | 0.96 | 6.47 |
| 9 | AMT | *dmdA* | *Pelagibacter ubique (HTCC1062)* | 3tfhA | 41 | 41 | 1.00 | 7.46 |
| 10 | AMT | *dmdA* | *Pelagibacter ubique (HTCC1062)* | 3tfhA | 41 | 41 | 1.00 | 6.69 |

^a^Iden1 is the percentage sequence identity of the templates in the threading aligned region with the query sequence

^b^Iden2 is the percentage sequence identity of the whole template chains with query sequence

^c^Cov-Represents the coverage of the threading alignment and is equal to the number of aligned residues divided by the length of the query protein

^d^Norm. Z-score is the normalized Z-score of the threading alignments. Alignment with a normalized Z-score>1 means a good alignment and vice versa.

^e^Aminomethyltransferase

^f^DmdA DMSP-dependent demethylase

^g^Glycine cleavage system T protein

^h^Dimethylglycine dehydrogenase complexed with tetrahydrofolate

**Predicted model: C-sore=1.45; TM-score=0.92 dev= 0.06**

| **Rank** | **Class** | **Gene name** | **Organism** | **PDB ID** | **TM-score^a^** | **RMSD^b^** | **IDEN^c^** | **Cov^d^** |
| --- | --- | --- | --- | --- | --- | --- | --- | --- |
| 1 | AMT | *dmdA* | *Pelagibacter ubique HTCC1062* | 3tfhA | 0.974 | 1.19 | 0.407 | 1.000 |
| 2 | Oxidoreductase | *dmgdH* | *Rattus norvegicus* | 4p9sA | 0.944 | 1.56 | 0.178 | 0.986 |
| 3 | AMT | *gcvT* | *Pyrococcus horikoshii* | 1v5vA | 0.935 | 1.53 | 0.214 | 0.975 |
| 4 | AMT | *gcvT* | *Thermotoga maritima* | 1worA | 0.930 | 1.33 | 0.234 | 0.962 |
| 5 | Oxidoreductase | *dmg^e^* | *Arthrobacter globiformis* | 1pj6A2 | 0.928 | 2.02 | 0.239 | 0.989 |
| 6 | Oxidoreductase | *dmg* | *Arthrobacter globiformis* | 1pj7A | 0.928 | 2.02 | 0.239 | 0.989 |
| 7 | Oxidoreductase | *soxA^f^* | *Stenotrophomonas maltophilia* | 2gagA | 0.922 | 1.99 | 0.193 | 0.984 |
| 8 | Oxidoreductase | *soxA* | *Corynebacterium sp. U-96* | 1x31A | 0.921 | 2.01 | 0.182 | 0.984 |
| 9 | AMT | *AMT(GCST)* | *Homo sapiens* | 1wsrA | 0.910 | 1.66 | 0.256 | 0.956 |
| 10 | AMT | *gcvT* | *Bartonella henselae* | 3girA | 0.903 | 1.71 | 0.228 | 0.951 |

^ab^It is a standard for measuring structural similarity between two structures

^c^It is the percentage sequence identity in the structurally aligned region

^d^It represents the coverage of the alignment by TM-align and is equal to the number of structurally aligned residues divided by length of the query protein

^e^Dimethylglycine oxidase

^f^Heterotetrameric sarcosine oxidase alpha-subunit

**Table 49**. Up: Top ten analogs identified by LOMETS for threading alignments. Case of “ancestral dmda and no-dmda sequence” inferred by FastML. In the middle: Model predicted by I-TASSER. Down: Top ten identified structural analogs in PDB by TM-align

| **Rank** | **Class** | **Gene name** | **Organism** | **PDB ID** | **Iden1 (%)^a^** | **Iden2 (%)^b^** | **Cov^c^** | **Norm. Z-score^d^** |
| --- | --- | --- | --- | --- | --- | --- | --- | --- |
| 1 | AMT^e^ | *gcvT^g^* | *Thermotoga maritima* | 1wopA | 29 | 34 | 1.00 | 3.97 |
| 2 | AMT | *AMT(GCST)* | *Homo sapiens* | 1wsrA | 25 | 30 | 1.00 | 3.91 |
| 3 | AMT | *gcvT* | *Thermotoga maritima* | 1worA | 30 | 34 | 0.99 | 4.77 |
| 4 | AMT | *dmdA^f^* | *Pelagibacter ubique HTCC1062* | 3tfh | 31 | 32 | 1.00 | 3.19 |
| 5 | Oxidoreductase | *dmgdH^h^* | *Rattus norvegicus* | 4p9sA | 16 | 23 | 0.99 | 2.30 |
| 6 | AMT | *gcvT* | *Bartonella henselae* | 3girA | 23 | 29 | 1.00 | 4.49 |
| 7 | *Oxidoreductase* | *dmgdH^h^* | *Rattus norvegicus* | 4p9sA | 19 | 23 | 0.99 | 3.37 |
| 8 | AMT | *gcvT* | *Bacillus subtilis* | 1yx2A | 27 | 31 | 0.99 | 4.64 |
| 9 | AMT | *dmdA* | *Pelagibacter ubique HTCC1062* | 3tfhA | 28 | 32 | 0.99 | 6.12 |
| 10 | AMT | *gcvT* | *Bartonella henselae* | 3girA | 24 | 29 | 0.99 | 5.62 |

^a^Iden1 is the percentage sequence identity of the templates in the threading aligned region with the query sequence

^b^Iden2 is the percentage sequence identity of the whole template chains with query sequence

^c^Cov-Represents the coverage of the threading alignment and is equal to the number of aligned residues divided by the length of the query protein

^d^Norm. Z-score is the normalized Z-score of the threading alignments. Alignment with a normalized Z-score>1 means a good alignment and vice versa.

^e^Aminomethyltransferase

^f^DmdA DMSP-dependent demethylase

^g^Glycine cleavage system T protein

^h^Dimethylglycine dehydrogenase complexed with tetrahydrofolate

**Predicted model: C-sore=1.25; TM-score=0.89 dev= 0.07**

| **Rank** | **Class** | **Gene name** | **Organism** | **PDB ID** | **TM-score^a^** | **RMSD^b^** | **IDEN^c^** | **Cov^d^** |
| --- | --- | --- | --- | --- | --- | --- | --- | --- |
| 1 | AMT | *gcvT* | *Thermotoga maritima* | 1wooA | 0.960 | 1.39 | 0.278 | 0.997 |
| 2 | Oxidoreductase | *dmgdH* | *Rattus norvegicus* | 4p9sA | 0.951 | 1.48 | 0.155 | 0.993 |
| 3 | AMT | *AMT(GCST)* | *Homo sapiens* | 1wsrA | 0.948 | 1.57 | 0.236 | 0.997 |
| 4 | AMT | *gcvT* | *Pyrococcus horikoshii* | 1v5vA | 0.947 | 1.57 | 0.254 | 0.997 |
| 5 | Oxidoreductase | *soxA* | *Corynebacterium sp U-96* | 1vrqA2 | 0.942 | 1.72 | 0.232 | 1.000 |
| 6 | Oxidoreductase | *soxA* | *Corynebacterium sp. U-96* | 1x31A | 0.941 | 1.73 | 0.232 | 1.000 |
| 7 | Oxidoreductase | *soxA^f^* | *Stenotrophomonas maltophilia* | 2gagA | 0.941 | 1.73 | 0.242 | 1.000 |
| 8 | Oxidoreductase | *dmg^e^* | *Arthrobacter globiformis* | 1pj7A2 | 0.928 | 1.89 | 0.225 | 0.997 |
| 9 | Oxidoreductase | *dmg* | *Arthrobacter globiformis* | 1pj6A | 0.928 | 1.89 | 0.225 | 0.997 |
| 10 | AMT | *gcvT* | *Bartonella henselae* | 3girA | 0.927 | 1.87 | 0.234 | 0.990 |

^ab^It is a standard for measuring structural similarity between two structures

^c^It is the percentage sequence identity in the structurally aligned region

^d^It represents the coverage of the alignment by TM-align and is equal to the number of structurally aligned residues divided by length of the query protein

^e^Dimethylglycine oxidase

^f^Heterotetrameric sarcosine oxidase alpha-subunit

**Table 50**. Up: Top ten analogs identified by LOMETS for threading alignments. Case of “ancestral dmda sequence” inferred by FastML. In the middle: Model predicted by I-TASSER. Down: Top ten identified structural analogs in PDB by TM-align

| **Rank** | **Class** | **Gene name** | **Organism** | **PDB ID** | **Iden1 (%)^a^** | **Iden2 (%)^b^** | **Cov^c^** | **Norm. Z-score^d^** |
| --- | --- | --- | --- | --- | --- | --- | --- | --- |
| 1 | AMT^e^ | *dmdA^f^* | *Pelagibacter ubique HTCC1062* | 3tfhA | 83 | 85 | 1.00 | 4.93 |
| 2 | AMT | *dmdA* | *Pelagibacter ubique HTCC1062* | 3tfhA | 83 | 85 | 1.00 | 5.48 |
| 3 | AMT | *dmdA* | *Pelagibacter ubique HTCC1062* | 3tfhA | 83 | 82 | 1.00 | 5.67 |
| 4 | AMT | *dmdA* | *Pelagibacter ubique HTCC1062* | 3tfh | 83 | 85 | 1.00 | 3.19 |
| 5 | Oxidoreductase | *dmgdH^h^* | *Rattus norvegicus* | 4p9sA | 17 | 20 | 0.98 | 2.33 |
| 6 | AMT | *dmdA* | *Pelagibacter ubique HTCC1062* | 3tfhA | 83 | 85 | 1.00 | 6.03 |
| 7 | *Oxidoreductase* | *dmgdH* | *Rattus norvegicus* | 4p9s | 17 | 20 | 0.98 | 3.50 |
| 8 | AMT | *gcvT^g^* | *Bacillus subtilis* | 1yx2A | 26 | 27 | 0.96 | 6.83 |
| 9 | AMT | *dmdA* | *Pelagibacter ubique HTCC1062* | 3tfhA | 83 | 85 | 1.00 | 8.06 |
| 10 | AMT | *dmdA* | *BPelagibacter ubique HTCC1062* | 3tfhA | 83 | 85 | 1.00 | 6.88 |

^a^Iden1 is the percentage sequence identity of the templates in the threading aligned region with the query sequence

^b^Iden2 is the percentage sequence identity of the whole template chains with query sequence

^c^Cov-Represents the coverage of the threading alignment and is equal to the number of aligned residues divided by the length of the query protein

^d^Norm. Z-score is the normalized Z-score of the threading alignments. Alignment with a normalized Z-score>1 means a good alignment and vice versa.

^e^Aminomethyltransferase

^f^DmdA DMSP-dependent demethylase

^g^Glycine cleavage system T protein

^h^Dimethylglycine dehydrogenase complexed with tetrahydrofolate

**Predicted model: C-sore=2; TM-score=0.99 dev= 0.04**

| **Rank** | **Class** | **Gene name** | **Organism** | **PDB ID** | **TM-score^a^** | **RMSD^b^** | **IDEN^c^** | **Cov^d^** |
| --- | --- | --- | --- | --- | --- | --- | --- | --- |
| 1 | AMT^e^ | *dmdA* | *Pelagibacter ubique HTCC1062* | 3tfhA | 0.997 | 0.42 | 0.825 | 1.000 |
| 2 | Oxidoreductase | *dmgdH* | *Rattus norvegicus* | 4p9sA | 0.930 | 1.84 | 0.181 | 0.986 |
| 3 | Oxidoreductase | *dmg^e^* | *Arthrobacter globiformis* | 1pj6A2 | 0.914 | 2.19 | 0.228 | 0.986 |
| 4 | Oxidoreductase | *dmg* | *Arthrobacter globiformis* | 1pj7A | 0.914 | 2.19 | 0.228 | 0.986 |
| 5 | AMT^e^ | *gcvT* | *Pyrococcus horikoshii* | 1v5vA | 0.905 | 2.03 | 0.239 | 0.973 |
| 6 | Oxidoreductase | *soxA^f^* | *Corynebacterium sp. U-96* | 3ad7A | 0.899 | 2.31 | 0.216 | 0.978 |
| 7 | Oxidoreductase | *soxA* | *Stenotrophomonas maltophilia* | 2gagA | 0.899 | 2.34 | 0.194 | 0.978 |
| 8 | AMT | *gcvT* | *Thermotoga maritima* | 1wooA | 0.897 | 1.79 | 0.242 | 0.951 |
| 9 | AMT | *AMT(GCST)* | *Homo sapiens* | 1wsrA | 0.889 | 2.16 | 0.208 | 0.962 |
| 10 | AMT | *gcvT* | *Bartonella henselae* | 3girA | 0.885 | 1.98 | 0.211 | 0.948 |

^ab^It is a standard for measuring structural similarity between two structures

^c^It is the percentage sequence identity in the structurally aligned region

^d^It represents the coverage of the alignment by TM-align and is equal to the number of structurally aligned residues divided by length of the query protein

^e^Dimethylglycine oxidase

^f^Heterotetrameric sarcosine oxidase alpha-subunit

**Table 51**. Up: Top ten analogs identified by LOMETS for threading alignments. Case of “ancestral no-dmda sequence” inferred by FastML. In the middle: Model predicted by I-TASSER. Down: Top ten identified structural analogs in PDB by TM-align

| **Rank** | **Class** | **Gene name** | **Organism** | **PDB ID** | **Iden1 (%)^a^** | **Iden2 (%)^b^** | **Cov^c^** | **Norm. Z-score^d^** |
| --- | --- | --- | --- | --- | --- | --- | --- | --- |
| 1 | AMT^e^ | *gcvT^g^* | *Thermotoga maritima* | 1wopA | 29 | 31 | 0.91 | 3.93 |
| 2 | AMT | *gcvT* | *Pyrococcus horikoshii* | 1v5vA | 26 | 29 | 0.94 | 4.52 |
| 3 | AMT | *gcvT* | *Thermotoga maritima* | 1worA | 29 | 31 | 0.91 | 5.02 |
| 4 | Oxidoreductase | *dmg* | *Arthrobacter globiformis* | 1pj6 | 23 | 26 | 0.96 | 3.21 |
| 5 | Oxidoreductase | *dmgdH^h^* | *Rattus norvegicus* | 4p9sA | 16 | 24 | 0.96 | 2.38 |
| 6 | AMT | *gcvT* | *Thermotoga maritima* | 1wopA | 29 | 31 | 0.91 | 4.81 |
| 7 | Oxidoreductase | *dmg^i^* | *Arthrobacter globiformis* | 1pj6 | 22 | 26 | 0.97 | 3.57 |
| 8 | AMT | *gcvT* | *Bacillus subtilis* | 1yx2A | 26 | 26 | 0.89 | 5.31 |
| 9 | AMT | *dmdA^f^* | *Pelagibacter ubique HTCC1062* | 3tfhA | 26 | 29 | 0.93 | 6.47 |
| 10 | Oxidoreductase | *dmg* | *Arthrobacter globiformis* | 1pj5A | 22 | 26 | 0.97 | 5.80 |

^a^Iden1 is the percentage sequence identity of the templates in the threading aligned region with the query sequence

^b^Iden2 is the percentage sequence identity of the whole template chains with query sequence

^c^Cov-Represents the coverage of the threading alignment and is equal to the number of aligned residues divided by the length of the query protein

^d^Norm. Z-score is the normalized Z-score of the threading alignments. Alignment with a normalized Z-score>1 means a good alignment and vice versa.

^e^Aminomethyltransferase

^f^DmdA DMSP-dependent demethylase

^g^Glycine cleavage system T protein

^h^Dimethylglycine dehydrogenase complexed with tetrahydrofolate

^i^Dimethylglycine oxidase

**Predicted model: C-sore=0.76; TM-score=0.82 dev= 0.09**

| **Rank** | **Class** | **Gene name** | **Organism** | **PDB ID** | **TM-score^a^** | **RMSD^b^** | **IDEN^c^** | **Cov^d^** |
| --- | --- | --- | --- | --- | --- | --- | --- | --- |
| 1 | Oxidoreductase | *dmgdH* | *Rattus norvegicus* | 4p9sA | 0.940 | 1.63 | 0.158 | 0.983 |
| 2 | Oxidoreductase | *dmg* | *Arthrobacter globiformis* | 1pj7A | 0.914 | 2.23 | 0.225 | 0.986 |
| 3 | AMT^e^ | *gcvT* | *Pyrococcus horikoshii* | 1v5vA | 0.892 | 1.68 | 0.250 | 0.933 |
| 4 | AMT^e^ | *gcvT* | *Thermotoga maritima* | 1worA | 0.892 | 1.14 | 0.292 | 0.914 |
| 5 | AMT^e^ | *AMT(GCST)* | *Homo sapiens* | 1wsvB | 0.882 | 1.59 | 0.217 | 0.922 |
| 6 | Oxidoreductase | *soxA^e^* | *Stenotrophomonas maltophilia* | 2gagA | 0.876 | 2.36 | 0.231 | 0.950 |
| 7 | Oxidoreductase | *soxA* | *Corynebacterium_sp_U-96* | 1vrqA | 0.874 | 2.25 | 0.212 | 0.944 |
| 8 | AMT | *gcvT* | *Bartonella henselae* | 3girA | 0.864 | 1.80 | 0.207 | 0.911 |
| 9 | AMT | *dmdA* | *Pelagibacter ubique HTCC1062* | 3tfiA | 0.859 | 2.18 | 0.254 | 0.931 |
| 10 | AMT | *gcvT* | *Escherichia coli* | 3a8iA | 0.859 | 1.82 | 0.208 | 0.903 |

^ab^It is a standard for measuring structural similarity between two structures

^c^It is the percentage sequence identity in the structurally aligned region

^d^It represents the coverage of the alignment by TM-align and is equal to the number of structurally aligned residues divided by length of the query protein

^e^Heterotetrameric sarcosine oxidase alpha-subunit
